# Substrates and Cyclic Peptide Inhibitors of the Oligonucleotide Activated SIRT7

**DOI:** 10.1101/2023.06.16.545261

**Authors:** Julie E. Bolding, Alexander L. Nielsen, Iben Jensen, Tobias N. Hansen, Line A. Ryberg, Samuel T. Jameson, Pernille Harris, Günther H. J. Peters, John M. Denu, Joseph M. Rogers, Christian A. Olsen

## Abstract

The sirtuins are NAD^+^-dependent lysine deacylases, comprising seven isoforms (SIRT1–7) in humans, which are involved in the regulation of a plethora of biology, including gene expression and metabolism. The sirtuins share a common hydrolytic mechanism but display preferences for different ε*-N*-acyllysine substrates. SIRT7 deacetylates targets in nuclei and nucleoli but remains one of the lesser studied of the seven isoforms; in part, because of a lack of chemical tools to specifically probe SIRT7 activity. Here we expressed SIRT7 and, using small-angle X-ray scattering, reveal SIRT7 to be a monomeric enzyme with low degree of globular flexibility in solution. We developed a fluorogenic assay for investigation of the substrate preferences of SIRT7 and to evaluate compounds that modulate its activity. We report several mechanism-based SIRT7 inhibitors as well as *de novo* cyclic peptide inhibitors selected from mRNA-display library screening that exhibit selectivity for SIRT7 over other sirtuin isoforms and stabilize SIRT7 in cells.

## Introduction

The sirtuins are a group of NAD^+^-dependent lysine deacylase enzymes with seven isoforms (SIRT1–7) present in humans. The enzymes share a conserved catalytic site and nucleotide binding pocket but serve distinct cellular roles and have different substrate preference and subcellular localization.^[1]^ The biological roles of SIRT7 remain less well established than for other sirtuin isoforms. SIRT7 is predominantly localized to the nucleus and centers around the nucleoli where it interacts with RNA polymerases and histone proteins.^[2]^ It has been found to remove acetyl,^[3]^ propionyl,^[3b]^ glutaryl,^[4]^ and succinyl^[5]^ posttranslational modifications from the ε-amino groups of lysine residues in a variety of proteins. The acetylated histone 3 lysine 18 (H3K18ac) and lysine 36 (H3K36ac) are among the most studied targets, being involved in a range of regulatory roles.^[3a, 6]^

For example, SIRT7 has been found to play a role in DNA repair,^[5, 6c]^ rDNA stability and pre-rRNA processing,^[2a, 7]^ endoplasmic reticulum stress,^[8]^ mitochondrial homeostasis,^[9]^ energy metabolism,^[10]^ and bone formation.^[3b]^ SIRT7 has also been implicated in different diseases,^[6e, 11]^ including tumorigenesis.^[12]^ Despite its main documented role as a deacetylase, SIRT7 more efficiently catalyzes the removal of ε-*N*-acyllysine modifications of longer length in biochemical assays *in vitro* ^[6f, 13]^ and has been shown to exert improved catalytic efficiency in the presence of various oligonucleotide additives ^[13a, 14]^ or reconstituted nucleosome particles.^[6f, 13b, 14]^ It is well documented that the catalytic activity of both SIRT1 and SIRT6 can be enhanced by allosteric binders^[15]^ and the activity of SIRT6 is increased by binding to nucleosomes^[16]^ with high affinity through its intrinsically disordered C-terminus.^[17]^ SIRT7, on the other hand, has extended N- and C-termini that are positively charged and have been proposed to interact with negatively charged patches of oligonucleotides.^[14]^ Structural elucidation of SIRT7 is challenging and has thus far been limited to computational predictions^[18]^ and an X-ray crystal structure of a fusion protein containing the N-terminal domain of SIRT7.^[19]^ Despite the intriguing biological implications of SIRT7, the development of modulators of its activity has been limited.^[20]^

Here, we establish a protocol for expression of catalytically active SIRT7 enzyme that is amenable to storage and small angle X-ray scattering (SAXS) show that SIRT7 is monomeric in solution and has a low degree of overall flexibility. With this enzyme in hand, we have developed an assay, which allows for semi-high-throughput evaluation of activators and inhibitors of the SIRT7 activity. Using this assay, several SIRT7 inhibitors were designed and, finally, *de novo* screening of large libraries of peptides, led to the discovery of a novel potent cyclic peptide inhibitor.

## Results and discussion

Because the SIRT7 obtained from commercial sources generally lack hydrolytic activity, it has been challenging to develop broadly applicable and efficient screening assays for determining substrate specificity of the enzyme and develop modulators of its activity. One previous assay was developed by Lin and co-workers, using recombinant His-tagged SIRT7 and HPLC-based evaluation of the SIRT7-mediated hydrolysis of PTMs from histone peptides of 12-to 15-mer length.^[13a, 14]^ However, we were interested in improving the throughput of the assay to enable more systematic evaluation of substrates and modulators of SIRT7. Also, because the N- and C-termini of the enzyme have been proposed to be involved in oligonucleotide recognition and be of importance for catalysis, we designed a construct in which the affinity tag could be removed by introducing a tobacco etch virus (TEV) protease cleavage site. We envisioned that SAXS could provide biophysical characterization and lower resolution structural insight into wildtype SIRT7.

Thus, samples of purified SIRT7 ranging from a concentration of 1.02 mg/mL to 3.91 mg/mL were prepared and subjected to SAXS measurement. From visual inspection of the resulting SAXS curves, the overall shapes of the curves did not change with increasing concentration and molecular masses derived from the SAXS data ranged from 40–46 kDa corresponding to a single SIRT7 monomer (45 kDa) (Supporting Figure S1). Further, a Kratky plot indicated that the protein is folded with low flexibility (Supporting Figure S1). A merged curve was generated from the data recorded at low and high concentration and this curve was used for modeling. A structure of human SIRT7 has been predicted using the AlphaFold2 structure prediction (PDB Q9NRC8) (Figure 1).^[18b, 21]^ Overall, the model confidence is high or very high for the N-terminal domain and the Rossman fold containing catalytic domain. For the N-terminal domain this likely reflects that an X-ray crystal structure is available (residues 5–73; PDB 5IQZ)^[19]^ and for the Rossman fold domain there is high sequence similarity to SIRT6, for which several structures are available (*e*.*g*., PDB 3K35,^[22]^ 6XV1,^[23]^ 7CL0,^[24]^ 8F86,^[16b]^ 8G57^[16c]^)

**Figure 1.**
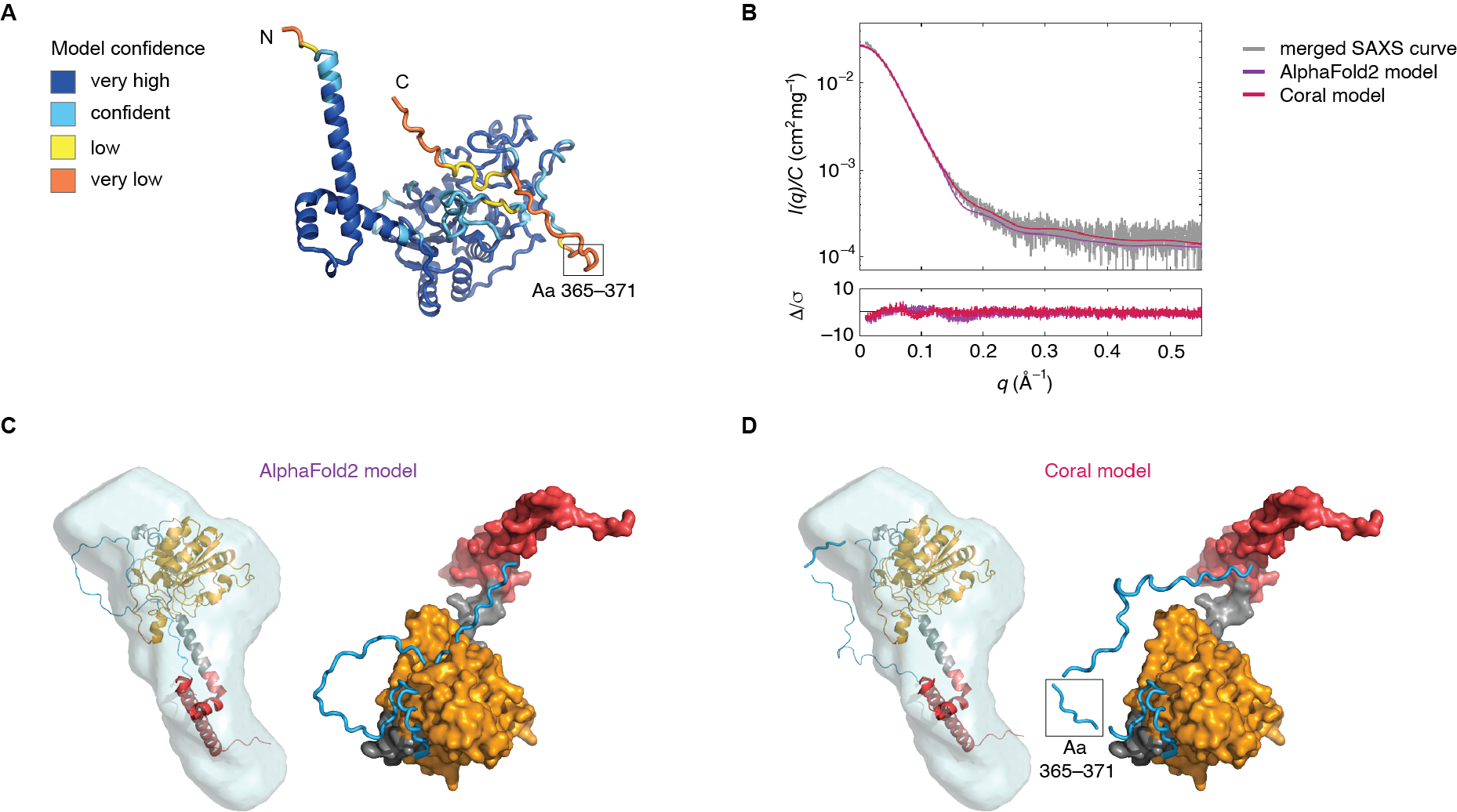
Small angle X-ray scattering. A) The AlphaFold2 model of human SIRT7 colored by model confidence based on the per-residue confidence score (pLDDT): very high (pLDDT > 90), confident (pLDDT = 90–70), low (pLDDT = 70–50), and very low (pLDDT < 50). Residues 365–371 (black box) were flexible in the Coral modeling. B) Crysol fits of the AlphaFold2 and the Coral model to the experimental SAXS data. C,D) Superposition of the SAXS derived envelope with the Alphafold2 model and the Coral model, respectively. Structural representation of the model with N-terminus (red) and Rossman fold and catalytic domain (orange) shown as surface view and the C-terminus (blue) shown as cartoon.

AlphaFold2 confidently predicts the N-terminus to be three a-helices that are very solvent-exposed. AlphaFold2 also predicts the inter-domain accuracy between these helices and the Rossman fold; it has confidence in orientation of N-terminus and how it projects from the catalytic domain. In contrast, the model confidence is either low or very low for the C-terminal domain from residues 364–400. The theoretical scattering curve for the AlphaFold2 model shows a reasonable fit to the merged experimental scattering curve with c^2^ = 2.3 (Figure 1B) and superimpose well with the generated SAXS envelope (Figure 1C). To improve the fit, modeling with Coral was carried out allowing flexibility in the C-terminal residues 365–371 for which the model confidence is very low in the AlphaFold2 prediction. The resulting Coral model has an improved fit to the experimental scattering curve with c^2^ = 1.3 (Figure 1B) and superimpose well with the SAXS envelope (Figure 1D). The Coral model has a different conformation of the C-terminus compared to the AlphaFold2 model, suggesting that the C-terminus does not bind to the Rossman fold (Figure 1C,D), which is in line with the low/very low confidence in the AlphaFold2 model in this area (Figure 1A). Taken together, our data fitting may reflect some degree of flexibility of the C-terminal domain, which is not seen in the generated Kratky plot; likely, because the C-terminal domain is a minor fraction of the protein. Whether the C-terminus may interact with substrate as well as the Rossman domain to bring the substrate into proximity with the active site will require high-resolution structural insight.

While many approaches could be considered for the discovery of novel substrates for sirtuin and HDAC enzymes,^[25]^ past efforts in our laboratory have primarily involved screening of modified peptides based on potential native target sequences,^[26]^ often by applying trypsin-coupled assays with the release of a C-terminally conjugated 7-amino-4-methylcoumarin (AMC) fluorophore.^[27]^ We therefore performed a screening of the activity of our recombinant SIRT7 against an in-house library of 40 selected fluorogenic substrates, based on five different peptide sequences. We further included oligonucleotide additives previously reported by Lin and co-workers to enhance the catalytic efficacy of SIRT7, i.e., double-stranded DNA (salmon sperm DNA sheared to an average size of less than 2000 base pairs) or yeast tRNA (Figure 2A,B).^[13a, 14]^ Generally, we observed a significantly lower activity of SIRT7 compared to the activity observed for other isoforms (SIRT1–3 and SIRT5) against their preferred substrates. We found that SIRT7 displays low activity against our selection of acetylated (Kac) fluorogenic substrates, despite acetylation being the most reported posttranslational modification hydrolyzed by SIRT7.^[28]^ We observed the highest activity of recombinant SIRT7 against hexanoyl (Khex, C6), octanoyl (Koct, C8), decanoyl (Kdec, C_10_), and lauryl (Klau, C_12_) containing substrates. Substantially higher conversions were observed for the substrates based on the QPKKacyl and TARKacyl peptide sequences compared to the alternative peptide sequences bearing the same acyl group, suggesting an increased activity against substrates containing a positively charged amino acid residue in the *i*-1 position. Interestingly, the two reported SIRT7 targets H3K18ac and H3K36ac both have neighbouring positively charged amino acid residues. In agreement with previously reported observations,^[13a, 14]^ we found that hydrolytic activity was enhanced substantially in the presence of the oligonucleotides and particularly by the addition of tRNA. While the maximum degree of hydrolysis was observed for the Ac-QPKKlau-AMC substrate, we found that its low aqueous solubility was prohibitive at the desired substrate concentration (50 μM final concentration). Thus, we proceeded with the assay development using Ac-QPKKdec-AMC, for which the second highest degree of hydrolysis was obtained. First, a titration of SIRT7 was performed in the presence or absence of oligonucleotides, which showed a linear correlation between substrate conversion and SIRT7 concentration (Figure 2C). Next, we performed a titration of DNA and tRNA activation with Ac-QPKKdec-AMC and observed a normal distribution relationship between oligonucleotide concentration and deacylation activity, with a maximum activation of 800% in the presence of SIRT7–tRNA (1:8 mol/mol). A further increase in tRNA concentration resulted in a Hook effect, where activity decreased (Figure 2D).

**Figure 2.**
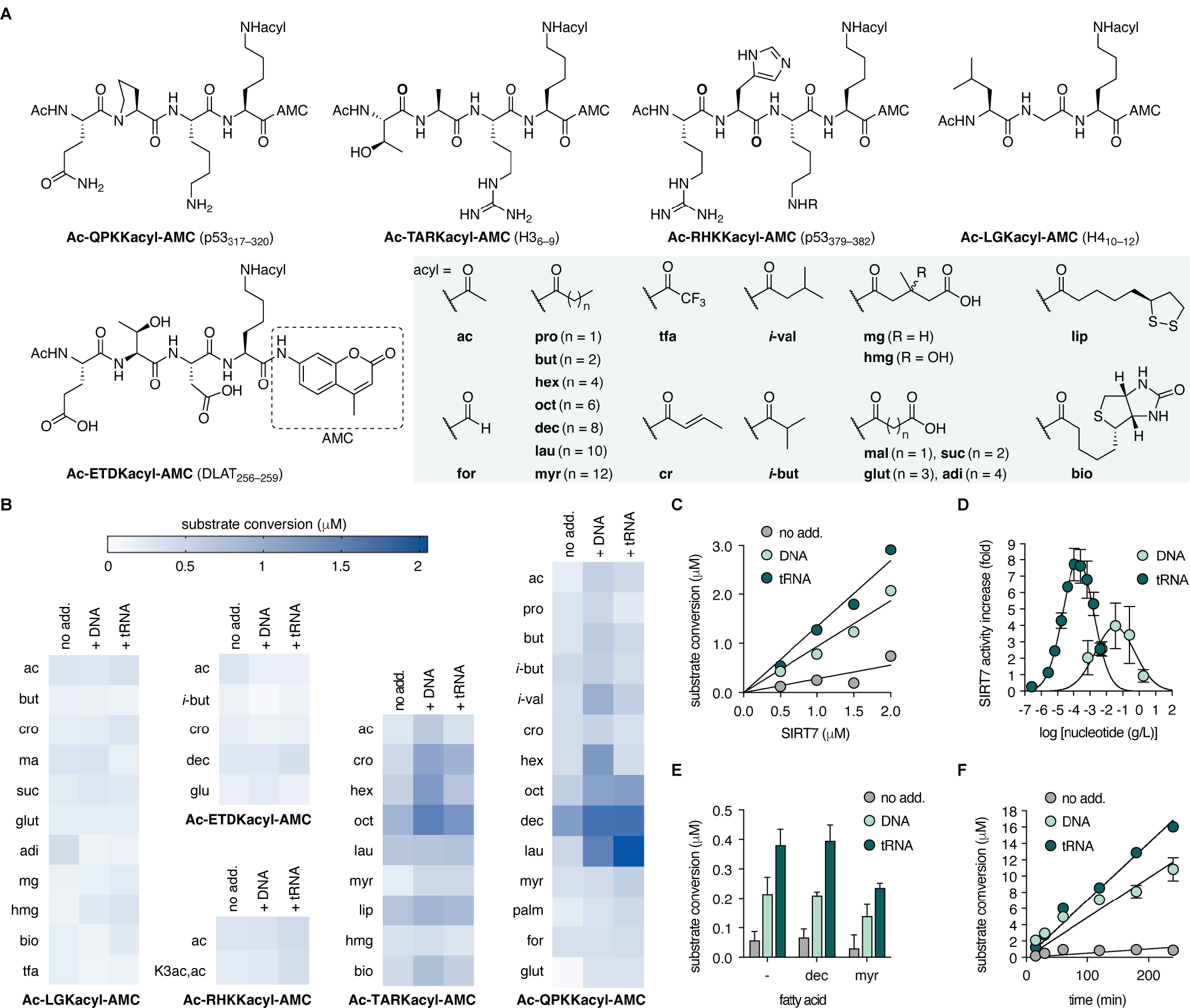
Screening for substrates of SIRT7. A) Substrate sequences and acyl groups (for novel substrates, see Supporting Figure S2). B) Heatmap of SIRT7-dependent deacylase activity [1 h incubation at 37 °C with SIRT7 (500 nM), substrate (50 μM), NAD^+^ (500 μM), and DNA (75 μg/mL) or tRNA (37 μg/mL)]. For data values, see Supporting Table S1. C) Titration of SIRT7 using Ac-QPKKdec-AMC (50 μM) as substrate. D) Titration of DNA and tRNA using Ac-QPKKdec-AMC (50 μM) as substrate. E) Activation of SIRT7-dependent deacetylation of substrate Ac-QPKKac-AMC (50 μM) by fatty acids. F) SIRT7-mediated hydrolysis over time, using Ac-QPKKdec-AMC (50 μM) as substrate. All data are based on two individual experiments performed in duplicate [1 h incubation at 37 °C]. Error bars represent the mean ± SD.

Both SIRT6 and SIRT7 belong to the class IV gene family and because SIRT7 mainly acts as a deacetylase in cells, while displaying a preference for long-chain acyl modifications in our fluorogenic assays, we envisioned that SIRT7 deacetylation could be activated by the presence of fatty acids as observed for SIRT6.^[15a]^ However, no increase in catalytic activity of SIRT7 was observed in the presence of neither capric- nor myristic acid (Figure 2E). Lastly, we tested the assay linearity over time; given the lower catalytic activity of recombinant SIRT7 compared to the catalytic efficiency of SIRT1–3 and SIRT5 in fluorogenic assays.^[26a, 26c, 29]^ The assay progression, and the SIRT7 stability (in presence of either DNA or tRNA), remained linear for up to 4 h at 37 °C (Figure 2F).

After observing that SIRT7 exhibited preference for the Ac-QPKKacyl-AMC substrates, which is based on the p53_317-320_ amino acid sequence, we envisioned that catalysis could be further enhanced by designing a substrate mimicking the sequences of well documented histone target sites. Past reports from Lin and co-workers monitored SIRT7 deacylation using an H3K9 sequence, instead of the validated SIRT7 targets on histone 3, which were H3K18 and H3K36^[3a, 6]^ and more recently also H3K37.^[6f]^ We therefore synthesized the fluorogenic H3_15−18_ (Ac-APRKacyl-AMC) and H3_33−36_ (Ac-GGVKacyl-AMC) substrates (Figure 3A) and monitored the SIRT7-mediated hydrolysis of these in presence of tRNA. Given the resemblance of the QPKK and the APRK sequence, it was not surprising that the catalytic efficiency towards the two were in the same range. Nevertheless, we observed a slight increase in activity of approximately 2-fold when applying the Ac-APRKacyl-AMC substrate. The GGVK motif, on the other hand, gave substantially lower conversion, which we speculate might be due to the lack of the positive charge from the K37 residue in the natural substrate. Again, we observed substantially lower deacetylase and demyristoylase activity compared to dedecanoylase activity (Figure 3B). We next attempted to perform kinetic experiments to determine Michaelis-Menten parameters with tRNA as an additive but, unfortunately, the data did not allow fitting to the equation due to an estimated *K*_M_ of >100 μM (Supporting Figure S3). Because nucleosomes have also been shown to activate SIRT7,^[13b]^ we instead turned to reconstituted nucleosome particles as an additive. Using the best performing substrate from the tRNA-activated tests (Figure 3B), we performed a titration of the concentration of nucleosomes added, showing optimal substrate conversion at 250 nM with a SIRT7 concentration of 500 nM (Figure 3C). The level of conversion of substrate 1b under nucleosome activation enabled determination of Michaelis-Menten parameters, giving a *K*_M_ of 12 μM (Figure 3D), which is somewhat higher than determined for longer histone peptides.^[13b]^ Further, the measured *k*_cat_ value is also low compared the ones reported with longer peptides, which may not be so surprising, given that our system is even more artificial than the previously reported. However, for the purposes of rapid compound screening and kinetic evaluation, we found the developed conditions satisfactory.

**Figure 3.**
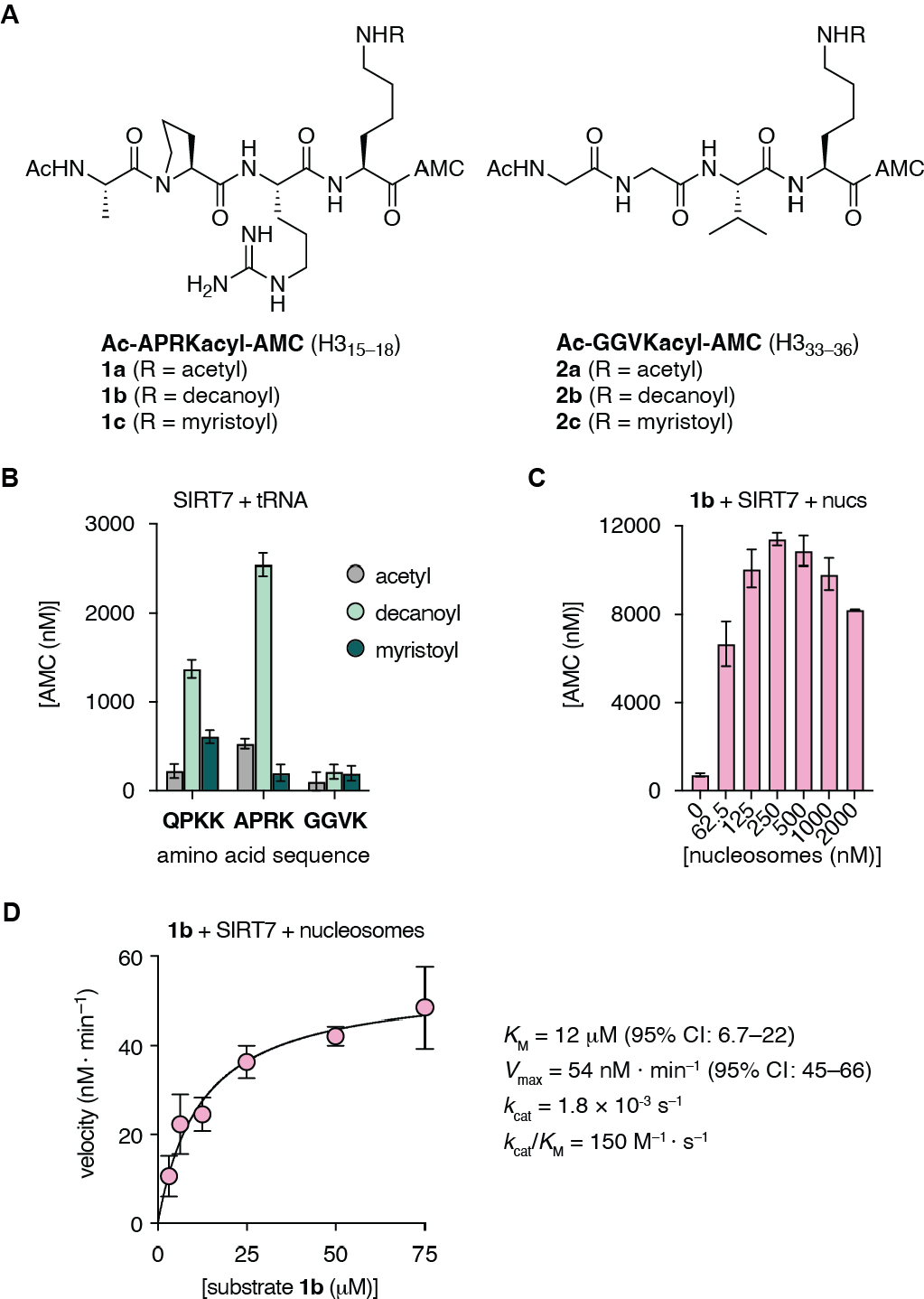
Novel substrates, activation by nucleosomes, and Michaelis-Menten kinetics for SIRT7. A) Structures of fluorogenic substrates based on the sequence of known target sites of SIRT7 in histone 3 (1a–c and 2a–c). For chemical synthesis, see Supporting Figure S2. B) SIRT7-mediated hydrolysis of selected substrates. Conditions: SIRT7 (500 nM), substrate (50 μM), NAD^+^ (500 μM), tRNA (4 μM). All reactions were incubated for 1 h at 37 °C. C) SIRT7-mediated hydrolysis of substrate 1b in the presence of nucleosomes. Conditions: SIRT7 (500 nM), substrate (50 μM), NAD^+^ (500 μM), nucleosomes (62.5–2000 nM). All reactions were incubated for 1 h at 37 °C. D) Michaelis-Menten kinetics for SIRT7-mediated hydrolysis of substrate 1b in the presence of nucleosomes. Conditions: SIRT7 (500 nM), substrate (3.125–75 μM), NAD^+^ (500 μM), nucleosomes (200 nM). All reactions were incubated for 1 h at 37 °C.

With our assay protocol established, we performed a structure–activity relationship (SAR) study to develop a potent inhibitor of SIRT7 as previously achieved for SIRT2, 3, and 5.^[30]^ First, we tested a selection of known sirtuin inhibitors with varying selectivity profiles, including nicotinamide,^[31]^ EX-527,^[32]^ TM,^[33]^ a macrocyclic peptide S2iL5,^[34]^ and three thiourea-containing mechanism-based inhibitors developed in our own lab ^[26f, 30a, 30b]^ (Supporting Figure S4). Compound 3 was the most potent inhibitor of the dedecanoylase activity of SIRT7 by a modest 26% at 100 μM inhibitor concentration (Figure 4A), and we therefore used its scaffold as the starting point for further improvement. We first tested a selection of ε-*N*-acyllysine mimics (4–7 Figure 4B), including a Ksuc mimic due to the reported desuccinylase activity of SIRT7,^[5]^ but none of these compounds, including the inverse thioamide (7), exhibited substantially improved potency. Next, we attempted to improve potency by substituting the amino acids in the *i*+2 position (Figure 4C) and *i*+1 position (Figure 4D), respectively. Compounds 8–13 revealed that an increase in potency could be achieved with a negatively charged side chain in the *i*+2 position and D-glutamate (11) even provided inhibition at 10 μM compound concentration (Figure 4C). The importance of the residue in the *i*+1 position was investigated by synthesizing compounds 14–29, encompassing variations in both size, polarity, and stereochemistry (Figure 4D). Most substitutions caused a decrease in potency with positively charged side chains (20–22) being very poorly tolerated and the introduction of D-amino acids (21 and 24) leading to a decrease in potency as well. Negatively charged and hydrophobic residues were better accommodated and compound 27, containing and L-valine residue, was the most potent of the series.

**Figure 4.**
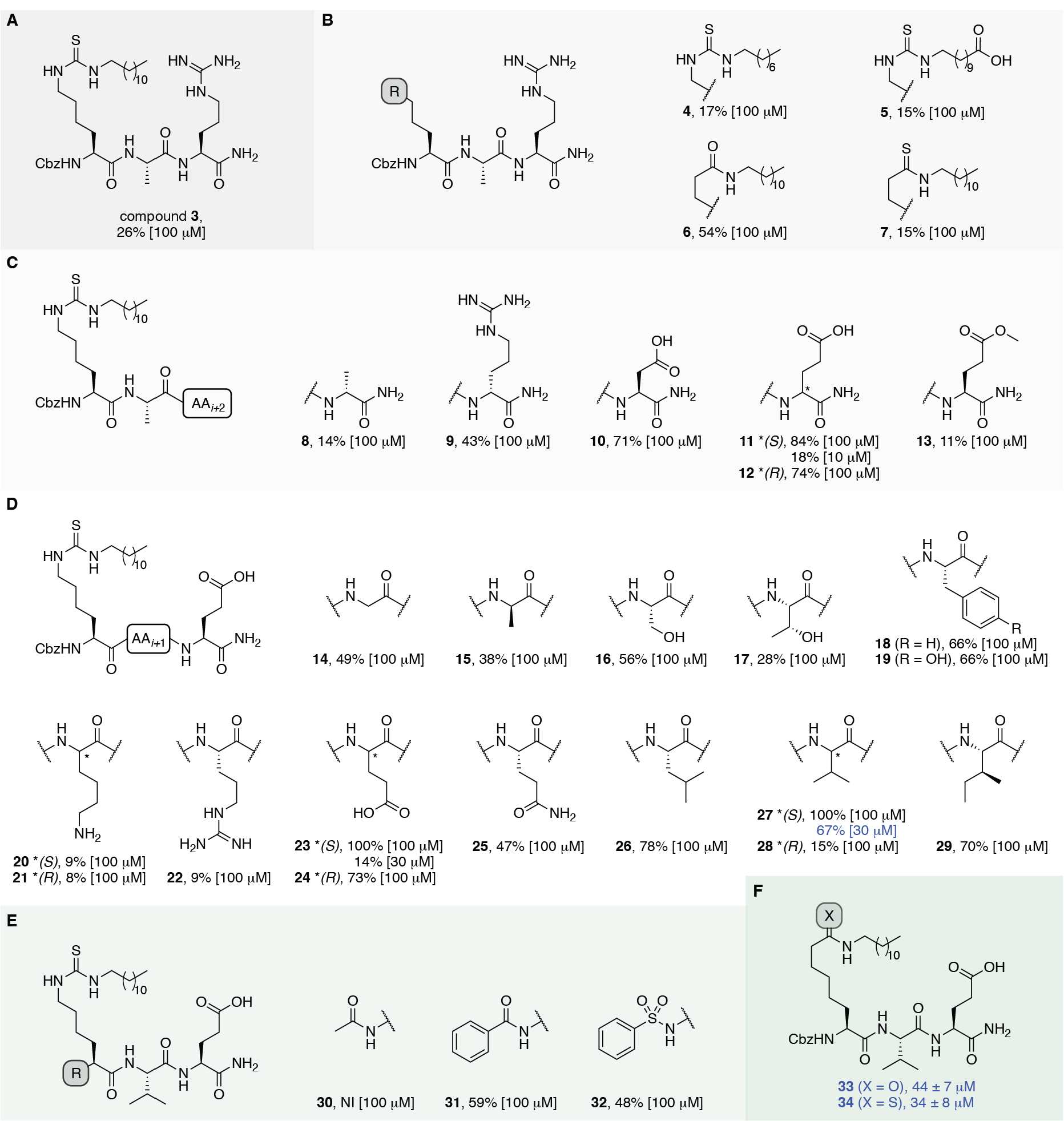
Structure-activity relationship study of SIRT7 inhibitors. A) Lead compound 3.^[30b]^ B) Amide isosteres. For chemical synthesis, see Supporting Figure S5. C) Optimization of the *i*+2 position. For chemical synthesis, see Supporting Figure S5. D) Optimization of the *i*+1 position. For chemical synthesis, see Supporting Figure S5. E) N-terminal derivatization. For chemical synthesis, see Supporting Figure S5. F) Final inverse (thio)amide inhibitors. For chemical synthesis, see Supporting Figure S5. Potencies are given as mean IC_50_-values ± SD or %-inhibition at denoted concentrations,using recombinant SIRT7 (500 nM), tRNA (4 μM), NAD^+^ (500 μM), and Ac-QPKKdec-AMC (50 μM). Data are based on two individual experiments performed in duplicate.

We then briefly investigated the importance of the N-terminal group by synthesizing the acetyl, benzamide, and phenylsulfonamide analogues (30–32; Figure 4E), which all led to a significant loss in potency. Finally, we synthesized inverse amide (33) and inverse thioamide (34) analogues based on the scaffold of the most potent compound (27) in the SAR study (Figure 4F). Unfortunately, the most potent inhibitors (33 and 34), exhibited potencies in the micromolar range (34–44 μM), which is substantially higher than previously achieved with mechanism-based chemotypes against other SIRT isoforms.^[30]^

We therefore used the “RaPID” screening platform, which has successfully furnished high-affinity cyclic peptides for a range of targets,^[35]^ including SIRT2,^[34]^ where binding was aided by a SIRT-targeting mechanism-based warhead. RaPID uses the ribosome to synthesize and screen large numbers (>10^12^) of peptides containing non-canonical amino acids that allow for cyclization. Here, the first amino acid of each peptide was reprogrammed to the non-canonical *N*-chloroacetyl-D-phenylalanine, which reacts spontaneously with a cysteine residue to form stable, thioether-containing macrocyclic structures. An mRNA library was constructed, encoding peptides starting with the *N*-chloroacetyl-D-phenylalanine, followed by 4–12 randomized canonical amino acids and then a cysteine (Figure 5A). We chose to omit a mechanism-based warhead, which could contribute to binding to other sirtuins, relying instead on just the peptide structure for specific interactions with SIRT7. Then, mRNA display was used to covalently attach each cyclic peptide to its encoding mRNA, after synthesis by the ribosome. This mRNA was reverse transcribed to generate the complimentary DNA that, after screening, could be sequenced to identify the SIRT7-binding cyclic peptides. We first incubated our RaPID library with immobilized full-length SIRT7. However, it became clear that SIRT7 was binding the oligonucleotide tag rather than the peptides (Supporting Figure S6A), consistent with the activation of SIRT7 by oligonucleotides (*vide supra*) that presumably bind to the enzyme. To allow selection based on peptide interaction with SIRT7, we followed two strategies. First, we added random DNA and RNA to outcompete any interaction between the full-length SIRT7 and the oligonucleotide tag. Second, we designed an additional SIRT7 construct that retained the enzymatic core but lacked the regions believed to drive nucleic acid binding, namely the N- and C-termini.^[36]^ We removed the N-terminal–lysine and arginine-rich–helical region, which projects from the core domain in the AlphaFold2 prediction. We also removed the C-terminal region that has low AlphaFold2 confidence score (pLDDT) and is likely to be disordered, leaving the core enzymatic domain amino acids 74–363 (Figure 5B). A RaPID selection was carried out using this core domain, including the addition of random DNA and RNA. Both of the RaPID selections, against full-length and the core domain of SIRT7, furnished enriched peptide sequences (Figure 5C, Supporting Figure S6B and S7) and we found that the protein target no longer bound the mRNA displayed peptides through the nucleic acids (Supporting Figure S6C,D). The top 3 sequences from the RaPID selection against full-length SIRT7 contained at least one cysteine residue (Supporting Figure S6B) and so did the top 5 hits from the selection against the catalytic domain (Figure 5C). We therefore set out to re-synthesize all the possible cyclic peptides based on these sequences (Supporting Figure S8A); although, previous reports have suggested that the smaller of the cycles are kinetically preferred.^[37]^ To ensure that only one cyclic structure was formed, and to avoid potential disulfide bond formation, the peptides were synthesized with the cysteines that did not engage in cyclization mutated to alanine (35–47; Supporting Figure S8A and Figure 5D). All successfully prepared peptide lariats and macrocycles (13 of the 16 attempted syntheses) were tested for their ability to inhibit the deacylase activity of SIRT7 (Supporting Figure S8B). The three most potent peptides (39, 41, and 44) were selected for full dose-response experiments where only the lariats 39 and 41 proved to be more potent than 34 (Supporting Figure S8C).

**Figure 5.**
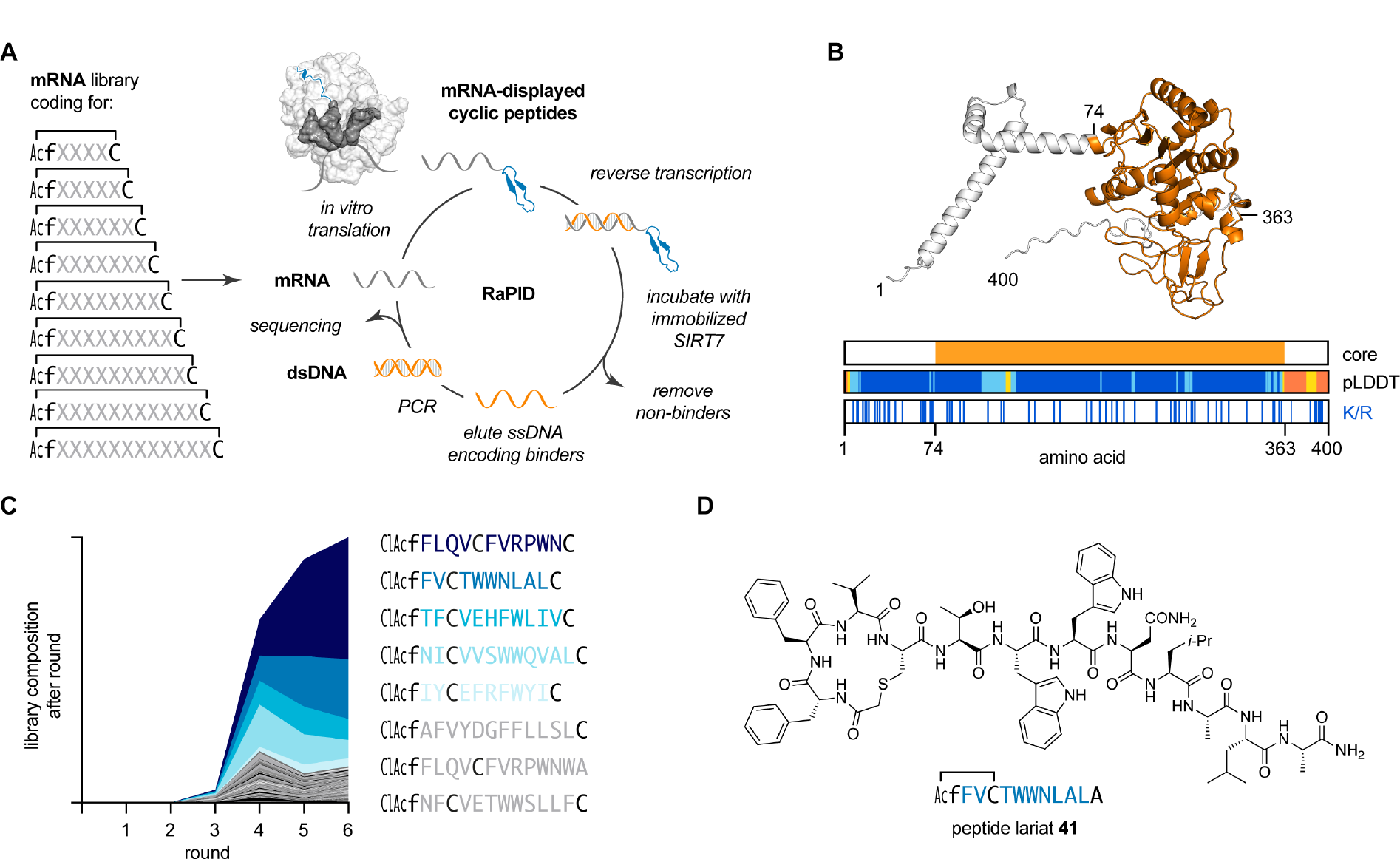
*De novo* cyclic peptides discovered for SIRT7 binding. A) Overview of RaPID selection, starting from a mRNA library encoding cyclic peptides with 4–12 randomized amino acids, where X can be any of canonical amino acids (apart from methionine). B) Design of construct of just the enzymatic core of SIRT7 (orange), guided by the AlphaFold2 structure prediction (shown), confidence measures (pLDDT), and distribution of lysine and arginine (K/R). C) Sequencing of RaPID selection against the core domain of SIRT7, showing the relative degree of enrichment of peptide sequences. D) Structure of the most potent re-synthesized peptide (41) derived from the RaPID selections.

To address selectivity, we tested the three most potent tripeptide inhibitors (27, 33, and 34) and the cyclic peptides 39 and 41 against five different recombinant sirtuin isoforms: SIRT1–3, 5, and 6 (catalytically active SIRT4 is not commercially available). Like the initial parent compound (3), we still observed notable inhibition of class I sirtuins for compound 27 (thiourea). Its inverse thioamide analogue 34 also inhibited SIRT1, 2, and 6 to a substantial extent at 10 μM inhibitor concentration, while the inverse amide (33) exhibited lower degree of inhibition of SIRT2 deacetylation and only limited inhibition of SIRT3 at 10 μM concentration (Figure 6A). Both peptide lariats were devoid of inhibition of SIRT2 but at 10 μM concentration, 39 and 41 exhibited some inhibition of SIRT3 and SIRT6, respectively (Figure 6A). We therefore performed full dose–response experiments with 39 against SIRT7 and SIRT3 (Supporting Figure S8D), which revealed just two-fold selectivity for SIRT7. On the other hand, full dose-response experiments for 41 against SIRT6 and 7 revealed a >50-fold selectivity and a *K*_i_ value against SIRT7 of 0.5 μM (Figure 6B).

**Figure 6.**
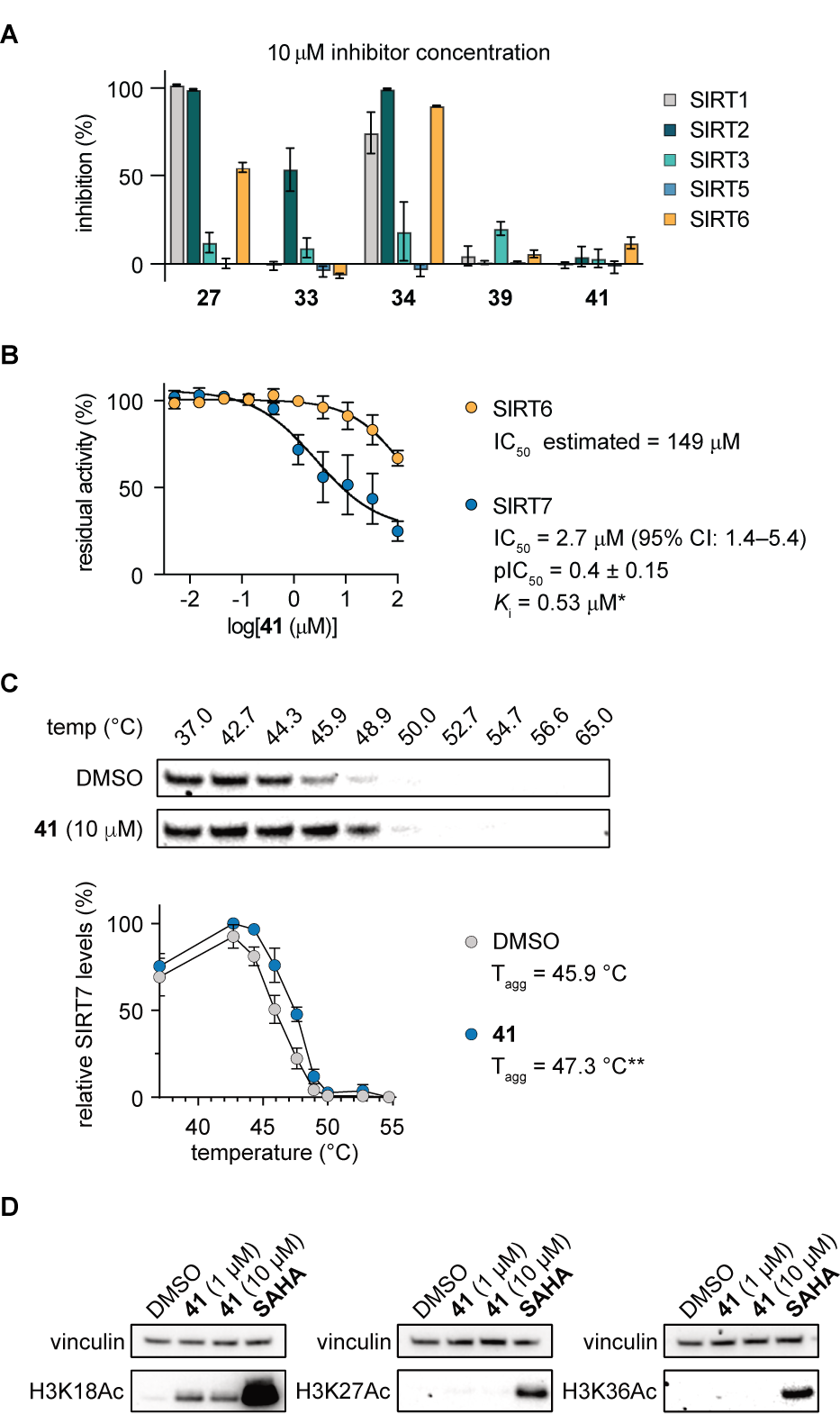
Selectivity of selected inhibitors and evaluation of the peptide lariat 41 SIRT7 inhibitor. A) Assessment of inhibition towards the other sirtuin subtypes using Ac-QPKKac-AMC (SIRT1–3), Ac-LGKsuc-AMC (SIRT5), and Ac-ETDKmyr-AMC (SIRT6) as substrates. B) Dose–response curves, IC_50_ values, and *K*_i_ values [*estimated by the Cheng-Prusoff equation: *K*_i_ = IC_50_/(1+ [S]/*K*_M_)] against recombinant SIRT7 (500 nM) using the Ac-QPKKdec-AMC (1b) substrate (50 μM), NAD^+^ (500 μM), and nucleosomes or recombinant SIRT6 (300 nM), NAD^+^ (500 μM), and Ac-ETDKmyr-AMC substrate (50 μM). C) Representative immunoblots for the thermal shift of SIRT7 in HEK293T cells after 2 h treatment with lariat 41 (10 μM) or DMSO control at temperatures ranging from 37–65 °C, plots of the data, and calculated T_agg_ values (*n* = 3; see Supporting Figure S9 for full blots). Statistical significance was calculated using unpaired *t*-test of the T_agg_ values from independent experiments. Adjusted *p*-value: \*\**p* < 0.01. D) Representative Western blots of histone acetylation (H3K18, H3K27, and H3K36) upon 5 h treatment with lariat 41 (1 or 10 μM), SAHA (10 μM), or DMSO. See Supporting Figure S10 for full blots (*n* = 2).

Finally, we investigated the ability of 41 to engage the intended target in living cells by performing immunoblot-based cellular thermal stability assay (CETSA)^[38]^ for SIRT7 for the first time. This protocol will be useful for the evaluation of future probes that target SIRT7 and, gratifyingly, we observed a significant stabilization of SIRT7 when treating HEK293T cells with peptide lariat 41 at 10 μM concentration (Figure 6C). Further, we found that 41 (10 μM) caused an increase in the acetylation level of H3K18 in HEK293T cells, while we did not observe an effect on neither H3K27 nor H3K36, which may require more sensitive means of detection (Figure 6D).

## Conclusion

In summary, we first expressed catalytically active SIRT7, characterized it biophysically using SAXS, and established conditions for assaying SIRT7 modulators by screening for the activity of our expressed enzyme against a collection of >40 different small fluorophore-conjugated peptides.

Our SAXS data provide experimental insight into the structure of a non-tagged, catalytically functional SIRT7 enzyme for the first time; although, the resolution of SAXS ranges from 50–10 Å.^[39]^ Thus, the SAXS data revealed that SIRT7 is a monomer with low structural flexibility in solution. Further, Coral modeling guided by the SAXS data, and using the AlphaFold2 structure of SIRT7 as starting point, provided a model of the globular structure of SIRT7, which, contrary to the AlphaFold2 structure, presented a C-terminus that does not bind to the Rossman domain. However, AlphaFold2 confidently predicts the full-length SIRT7 to be highly non-globular, including single, solvent-exposed α-helices of the N-terminus projecting at a defined angle from the enzymatic core, and the SAXS data largely support this prediction.

In agreement with previous investigations using longer peptide substrates in an HPLC-based assay format, we found that substrates containing longer acyl modifications to be most efficiently hydrolyzed,^[13a, 14]^ with Kdec-containing substrates being the preferred We also noticed from our screening that peptide sequences containing a positive charge were preferred over neutral and negatively charged sequences, which is in accordance with demonstrated targets of SIRT7 (*i*.*e*., H3K18 and H3K36).^[6a, 6f, 40]^ This also agrees with the previously proposed hypothesis that the negatively charged oligonucleotides enable the formation of a ternary complex (substrate–oligonucleotide–SIRT7),^[6f]^ which was also supported by an observed a Hook effect for both DNA and tRNA as activators. Unlike the behavior of SIRT6, which is known to be activated by fatty acids,^[15a, 15d]^ neither decanoic nor myristic acid activated SIRT7. Reconstituted nucleosomes, on the other hand, activated the catalytic removal of acyl groups substantially more than the tRNA (>4-fold), and whereas Michaelis-Menten parameters could not be determined using tRNA, the addition of nucleosomes allowed for determination of kinetic parameters of the system.

Next, we focused on the development of inhibitors of SIRT7. Only three examples of SIRT7 inhibitors were previously reported, a small molecule and two cyclic peptide analogues.^[20]^ The cyclic peptides also exhibited potent inhibition of SIRT1–3 and 6, with SIRT2 being inhibited 10-fold more potently than SIRT7^[20b]^ and selectivity was not addressed for the small molecule-based inhibitor, leaving ample room for further development of inhibitors. Thus, we performed the first SAR study reported for development of SIRT7 inhibitors, where we iteratively designed and tested 30 tripeptide analogues to optimize the amino acid sequence, the N-terminal functional group, and the ε-*N*-acyllysine mimicking motif. Unfortunately, the most potent compounds [33 (IC_50_ = 44 μM) and 34 (IC_50_ = 34 μM)] exhibited more potent inhibition of SIRT2 than SIRT7.

We therefore finally turned to screening of large libraries of random peptides to find *de novo* inhibitors of SIRT7, using the RaPID system, which allows screening of trillions of cyclic peptides. The top hits from selection for binding to either the full-length SIRT7 or the core enzymatic domain were re-synthesized and characterized. The peptide lariat, 41, had a >10-fold higher potency than the best mechanism-based inhibitor (34) and is the most potent inhibitor of *in vitro* SIRT7 activity reported to the best of our knowledge (*K*_i_ = 0.5 μM). Lariat 41 is also the first compound to convincingly exhibit selectivity for SIRT7 over other sirtuin isoforms (>50-fold), and we were able to demonstrate target engagement in HEK293T cells by CETSA as well as hyper acetylation of H3K18. Together these data, render lariat 41 a promising starting point for the development of inhibitors to fully investigate the SIRT7 *in vivo*. Further, because RaPID selects for protein binders rather than inhibitors, future investigations into the detailed binding mechanism and affinity of 41 against SIRT7 may reveal whether the peptide has potential as a starting point for the development of targeted protein degraders.^[41]^

## Supporting information

Supporting Information includes figures, tables, experimental methods, compound characterization data, copies of HPLC chromatograms, and copies of NMR spectra for synthesized compounds. The authors have cited additional references within the Supporting Information.^[42]^

## Acknowledgements

We thank Drs. Peter Fristrup and Ana Rita Colaço (PhD thesis, DTU Chemistry 2017) for early homology modelling work on SIRT7 that is not included in the present manuscript. We acknowledge DESY (Hamburg, Germany), a member of the Helmholtz Association HGF, for providing beam time for the SAXS experiments and provision of experimental facilities; the data were collected at PETRA III beamline P12 operated by EMBL Hamburg. Beamtime was allocated for proposal SAXS-734. We thank the Independent Research Fund Denmark– Medical Sciences (0134-00435B; C.A.O.) for financial support. This project has received funding from the European Research Council (ERC) under the European Union’s Horizon 2020 Research and Innovation Programme (grant agreement number: 725172–*SIRFUNCT*; C.A.O.).

## TOC Graphic (11 × 2.5 cm)

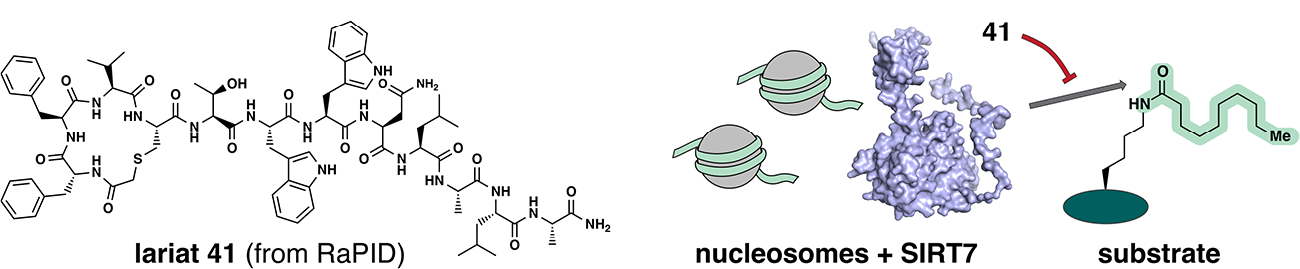

### TOC text (max 450 characters)

Broad screening of the substrate specificity of SIRT7 revealed a strong preference for the cleavage of long chain acyl lysine modifications. Further, the activity was substantially increased in the presence of oligonucleotides or nucleosome particles. Finally, mRNA display selection delivered a cyclopeptide with potency in the sub-micromolar range and selectivity for SIRT7, which binds and stabilizes SIRT7 in cells.

