## Supporting Information for "Substrates and Cyclic Peptide Inhibitors of the Oligonucleotide Activated SIRT7"

#### Supporting figures

A

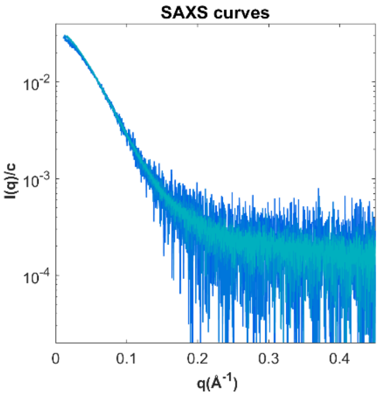

B

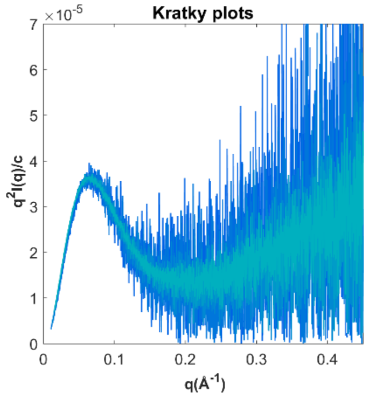

C

| **Conc.** *mg/ml* | **I(0)/conc.** *ml/mg* | **MM** *kDa* | **Rg** *Å* | **Dmax** *Å* | **V** *$nm^3$* |
|---|---|---|---|---|---|
| 1.02 | 0.028 | 39 | 32 | 113 | 86 |
| 1.07 | 0.029 | 40 | 33 | 114 | 89 |
| 2.11 | 0.031 | 43 | 35 | 126 | 94 |
| 3.30 | 0.032 | 44 | 36 | 128 | 96 |
| 3.91 | 0.033 | 46 | 37 | 135 | 99 |

D

| **Instrument** | **P12, PETRA III, DESY** |
|---|---|
| Detector | Pilatus 6M |
| Wavelength (Å) | 1.24 |
| q-range (nm-1) | 0.026-7.263 |
| Exposure time (s) | 30 x 0.095 |
| Temperature (K) | 293 |
| Sample-to-detector distance (mm) | 3000 |

**Figure S1.** Small angle X-ray scattering data.
(A) SAXS curves plotted on a log-scale. (B) Kratky plot. (C) Table showing molecular parameters derived from SAXS data: Forward scattering (I(0)), molecular mass (MM), radius of gyration (Rg), maximum intramolecular distance (Dmax) and Porod volume (V). (D) Table showing the experimental setup of the SAXS experiments.

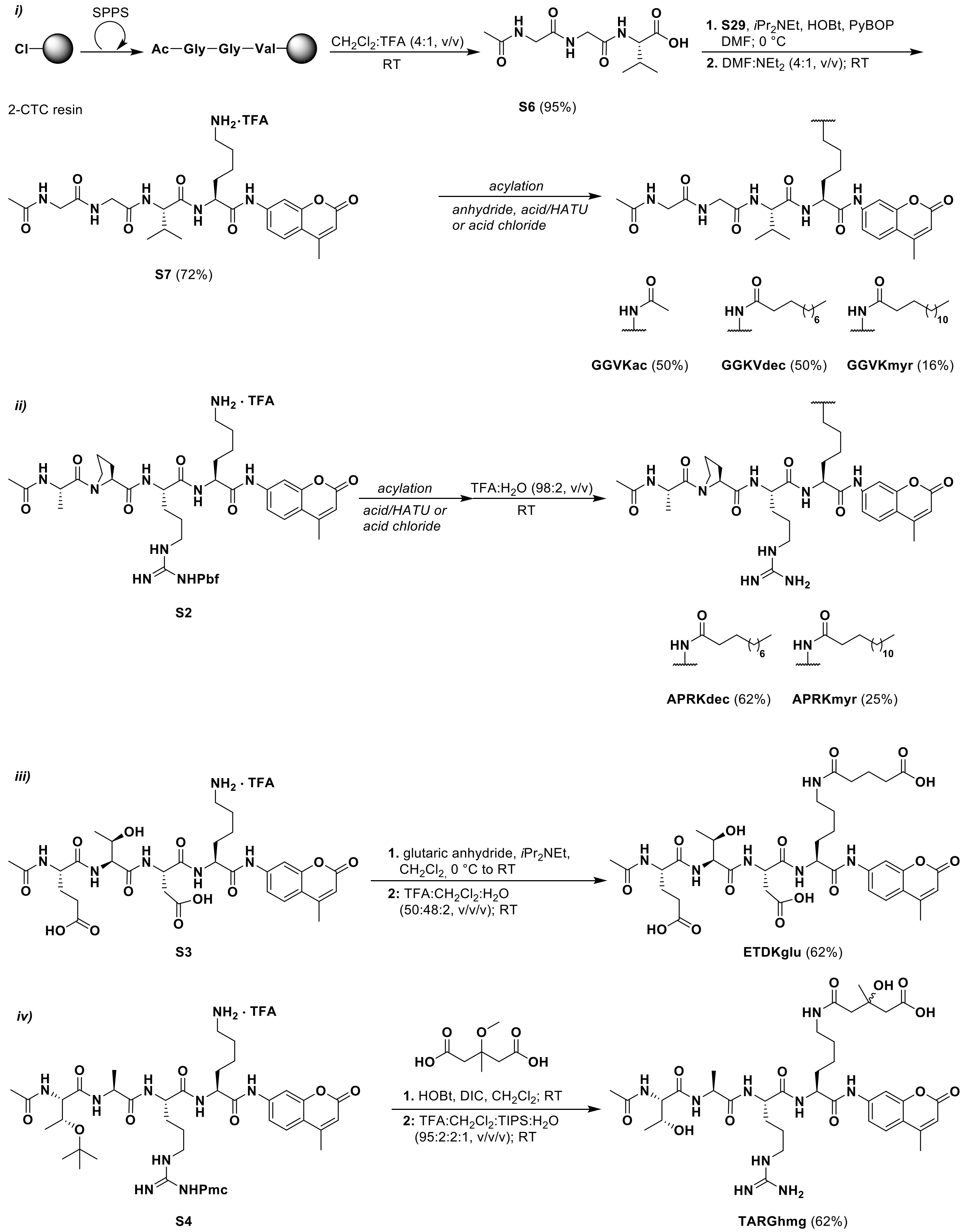

**Figure S2.** Synthesis of fluorogenic substrates.

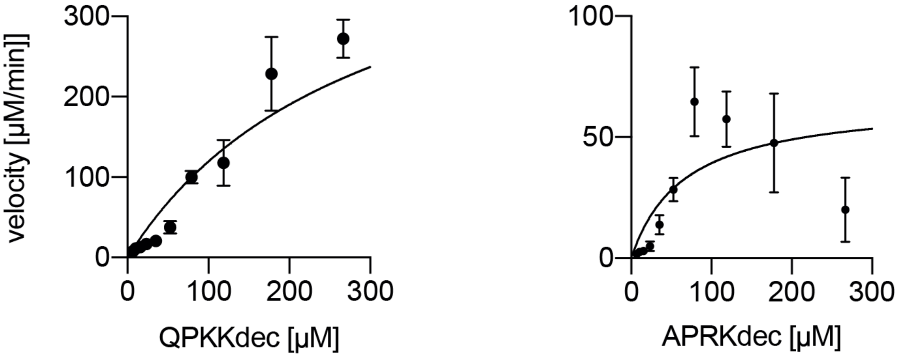

**Figure S3.** Michaelis-Menten kinetic parameters.
Michaelis-Menten plot of steate-state rate experiments for dedecanoylation of SIRT7 (500 nM) against the QPKKdec and APRKdec substrates. Data are presented as mean ± SEM. Note: Data above 100 µM are ambigous due to limited aqueous solubility.

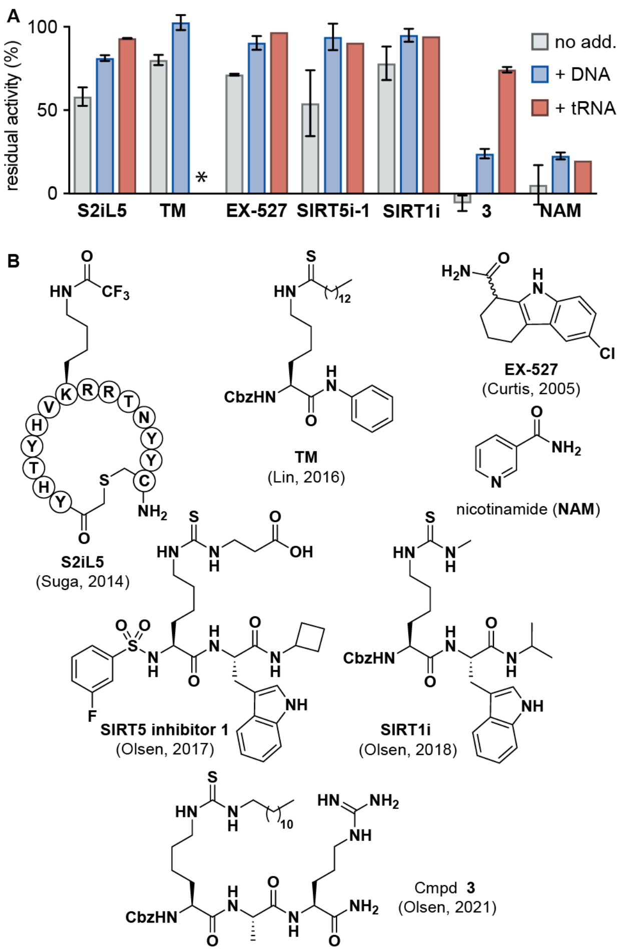

**Figure S4**. Preliminary inhibitor screening.
*Not tested

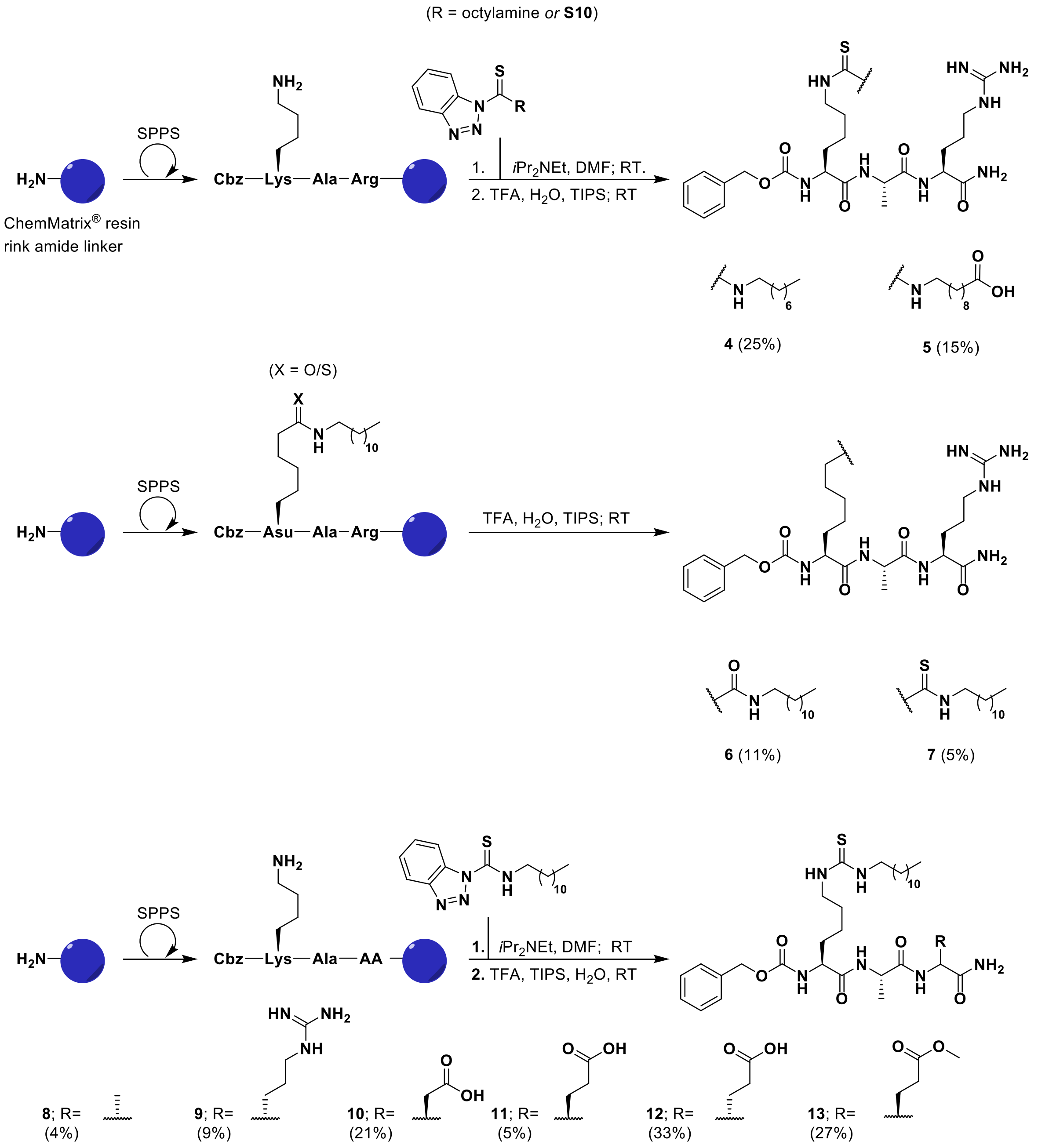

**Figure S5.** Synthesis of inhibitors on solid-phase.

*Continued on next page*

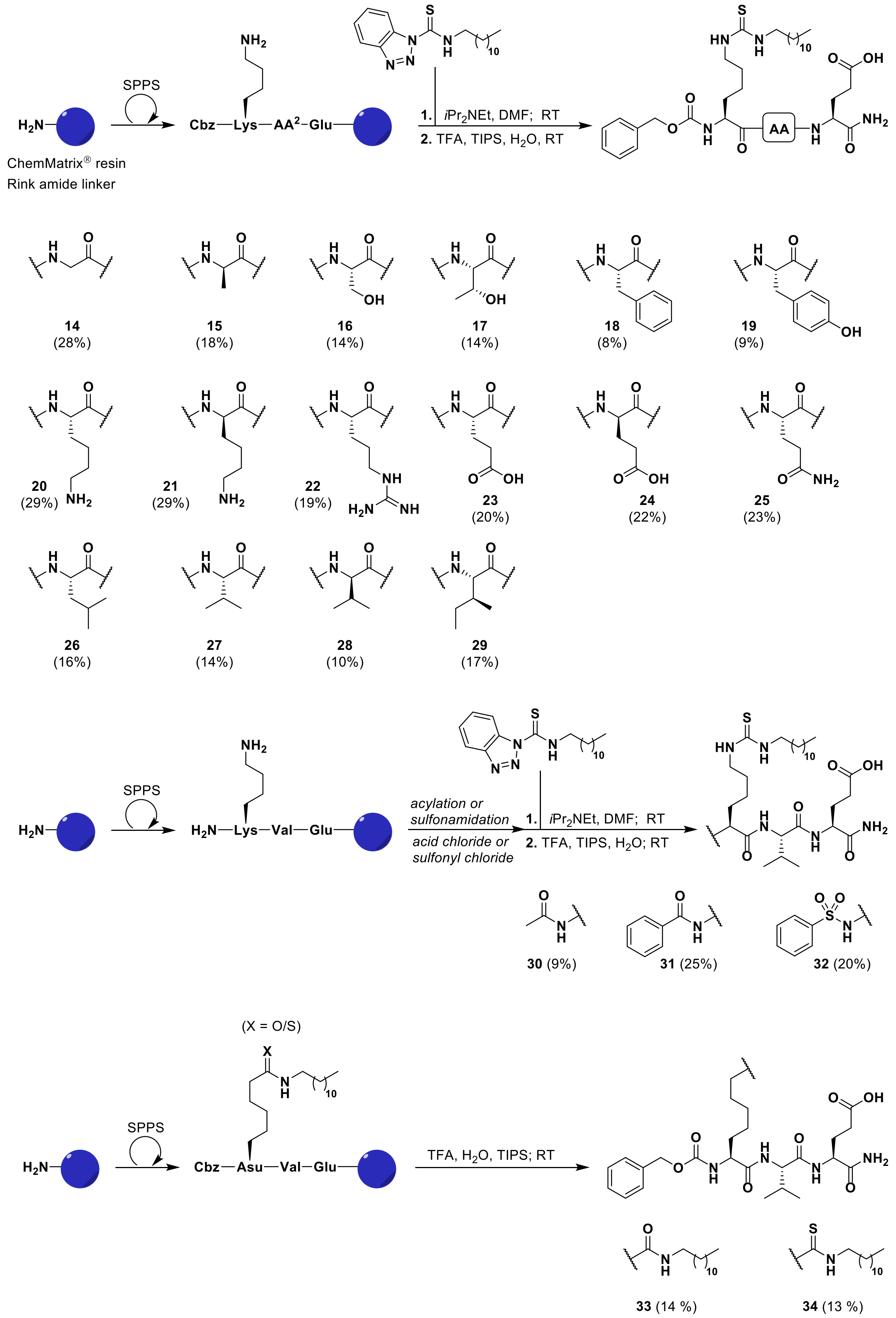

**Figure S5–part II** . Synthesis of inhibitors on solid-phase.

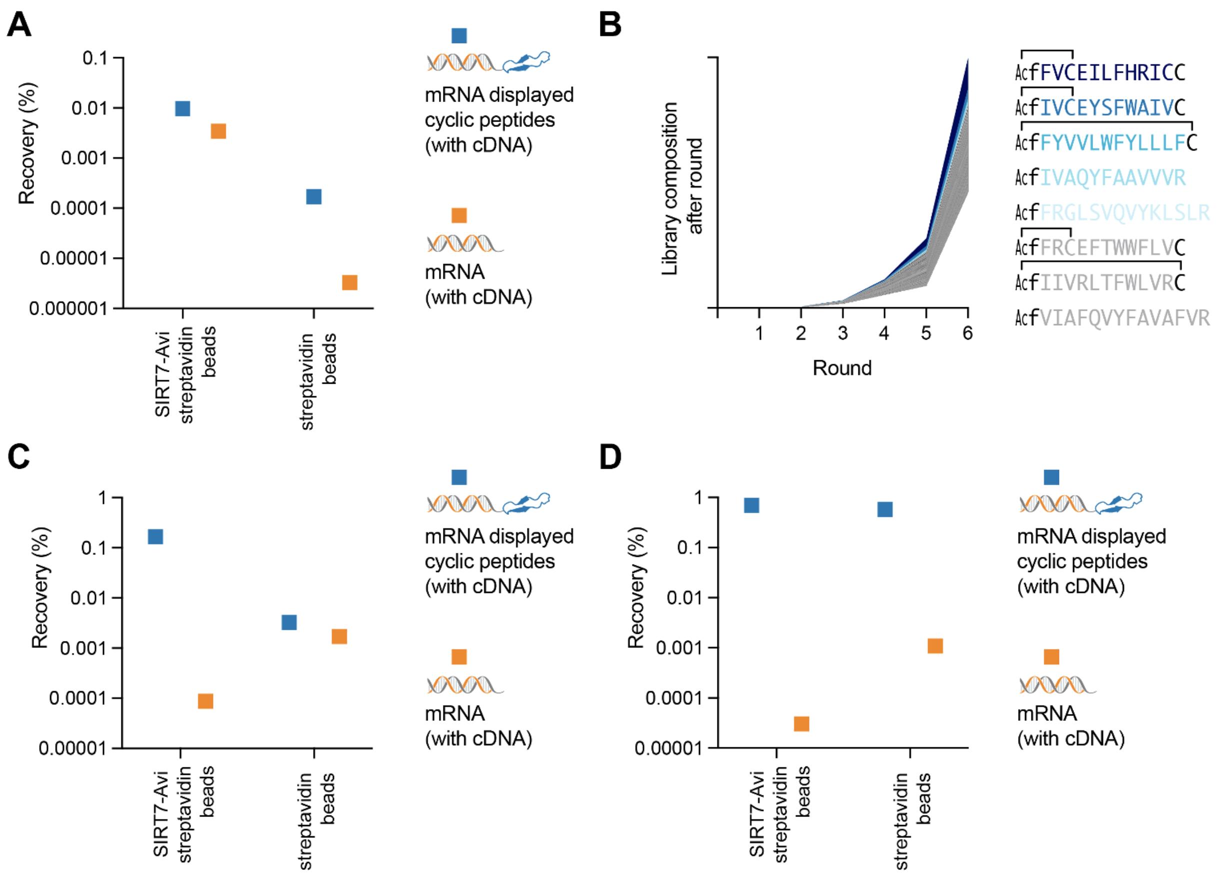

**Figure S6.** RaPID selection data.
(A) Product of round 3 of the first RaPID selection against full-length SIRT7, with N-terminal SUMO tag, and no additional random DNA or RNA, tested for binding to magentic beads with and without immobilized SIRT7, followed using qPCR of recovered DNA. Equivalent recovery with and without the mRNA displayed peptide suggests that SIRT7 binds the oligonucleotide tag rather than the peptide. (B) Sequencing of RaPID selection against the full-length SIRT7 showing enrichment of particular peptide sequences. (C) Product of round 6 of the RaPID selection against the core domain of SIRT7, including random DNA and RNA, tested for oligonucleotide binding shows that the peptide is responsible for the interaction. (D) Product of round 6 of the RaPID selection against full-length SIRT7, including random DNA and RNA, shows that the peptide is responsible for the binding instead of the . However, high recovery in the absence of SIRT7 suggests the majority of peptides have affinity for streptavidin.

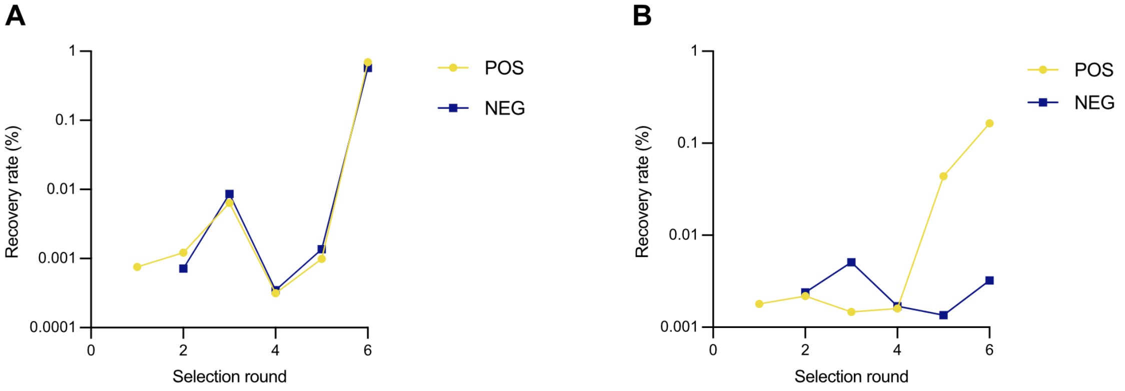

**Figure S7.** Total peptide recovery in RaPID selection rounds.
(A) Sequencing of RaPID selection against the full-length SIRT7 showing enrichment of particular peptide sequences. (B) Sequencing of RaPID selection against the core domain of SIRT7, showing enrichment of peptide sequences

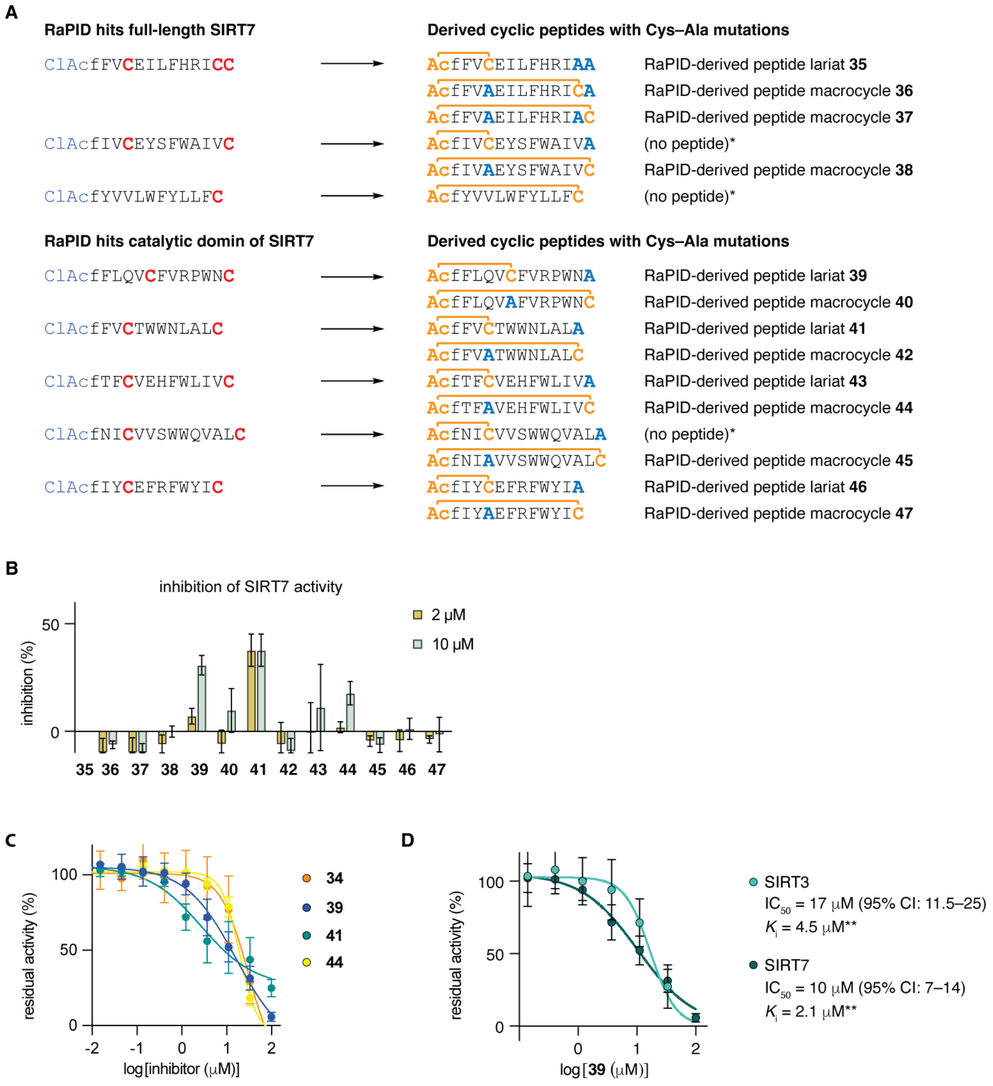

**Figure S8.** Sequences and potencies of RaPID-derived peptide lariats and macrocycles. (A) Hit sequences from RaPID and sequences of designed peptides derived from these. (B) Two-dose screening for activity of successfully synthesized peptides. (C) Dose-response curves for inhibition of SIRT7 by **34**, **39**, **41**, and **44**. (D) Dose–response curves, $IC_{50}$ values, and $K_i$ values (*estimated by the Cheng-Prusoff equation) for the inhibition of the deacylation activity of recombinant SIRT7 (500 nM) against the Ac-QPKKdec-AMC (**1b**) substrate (50 µM) in the presence of $NAD^+$ (500 µM) and nucleosomes or recombinant SIRT3 (300 nM) against $NAD^+$ (500 µM) and Ac-QPKKmyr-AMC substrate [10 µM; $K_M$ value = 3.6 µM].[1]

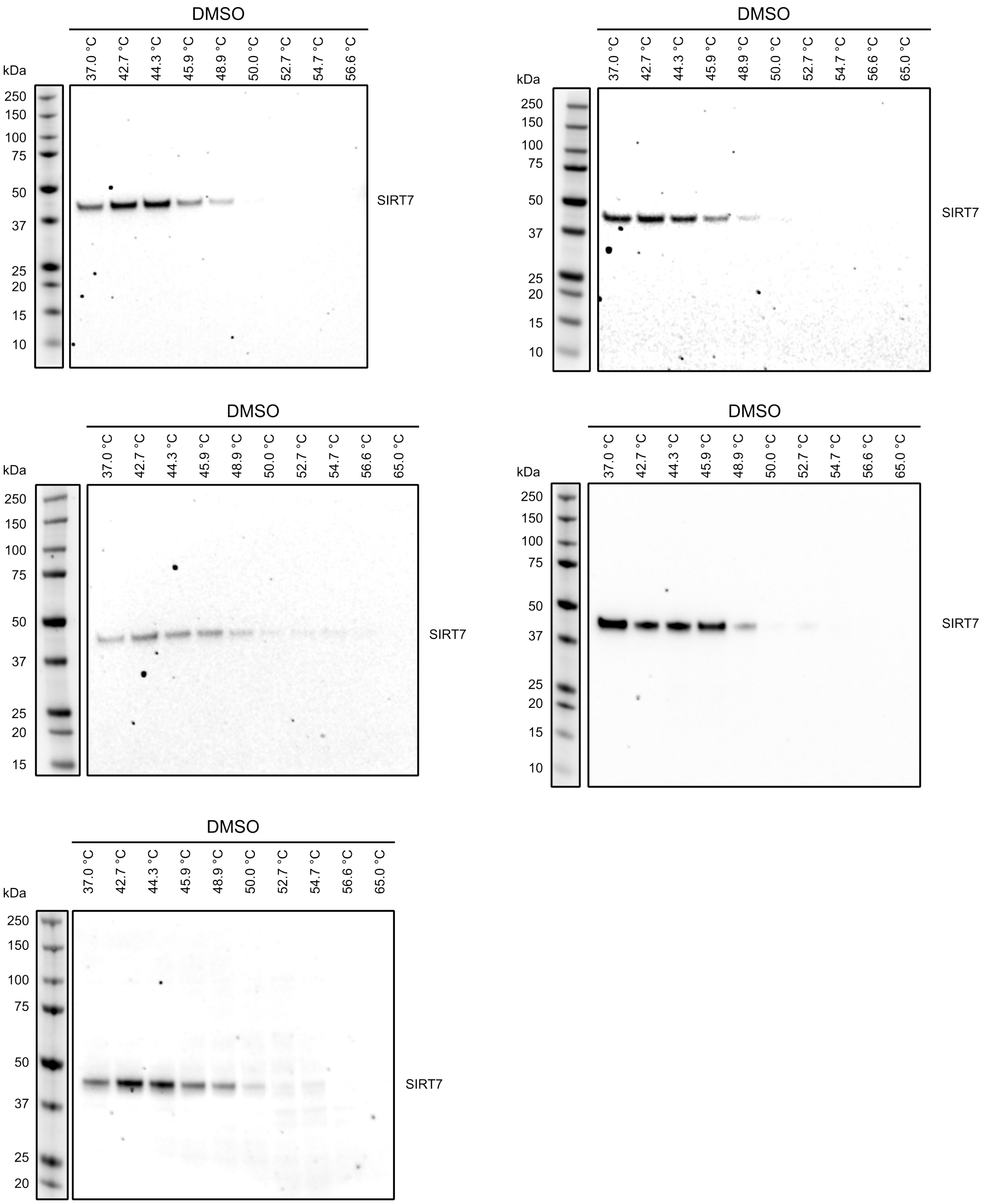

**Figure S9.** Full blots from all replicates of the CETSA assays.
Immunoblots for the thermal shift of SIRT7 in HEK293T cells after 2 h treatment with DMSO control at temperatures ranging from 37–65 °C

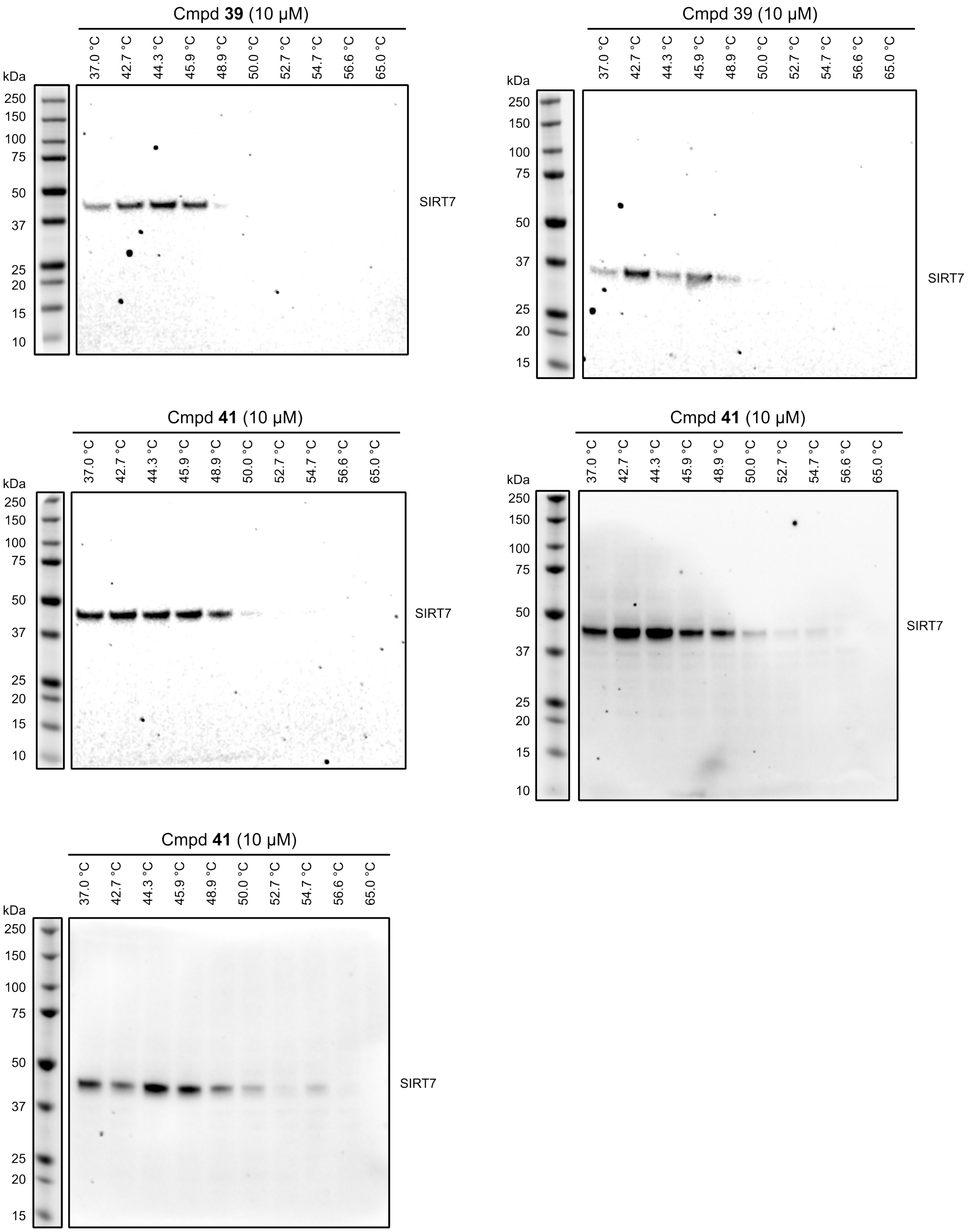

**Figure S9 continued.** Full blots from all replicates of the CETSA assays. immunoblots for the thermal shift of SIRT7 in HEK293T cells after 2 h treatment with peptide **39** (10 µM) or lariat **41** (10 µM) at temperatures ranging from 37–65 °C

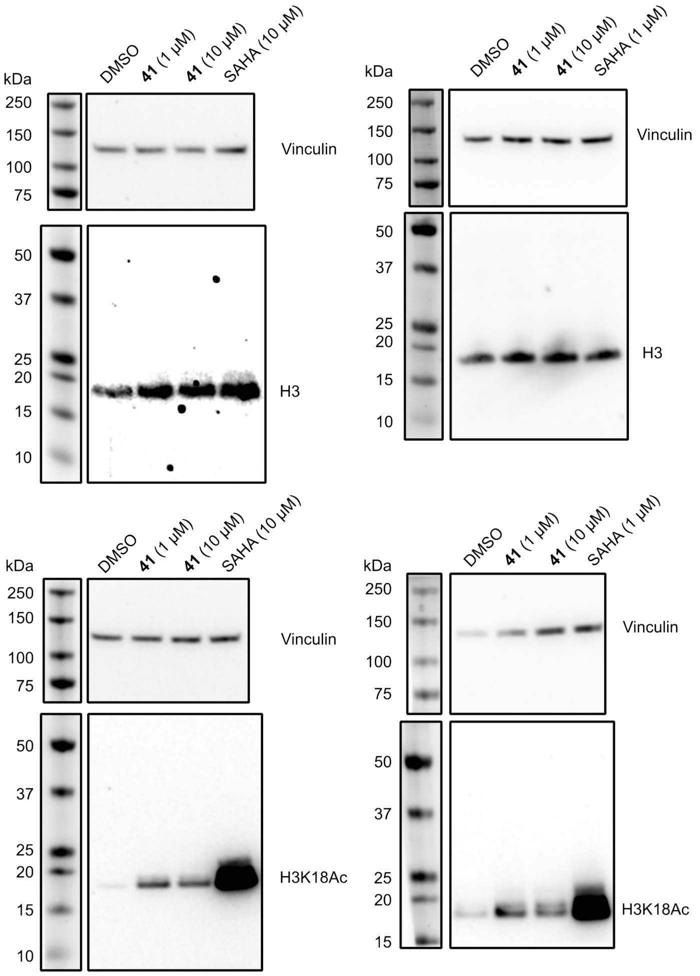

**Figure S10.** Full blots from all replicates of the histone acetylation assays.
Western blots of histone acetylation (H3K18, H3K27, and H3K36) or histone 3 (H3) upon 5 h treatment with lariat 41 (1 or 10 µM), SAHA (10 or 1 µM), or DMSO.

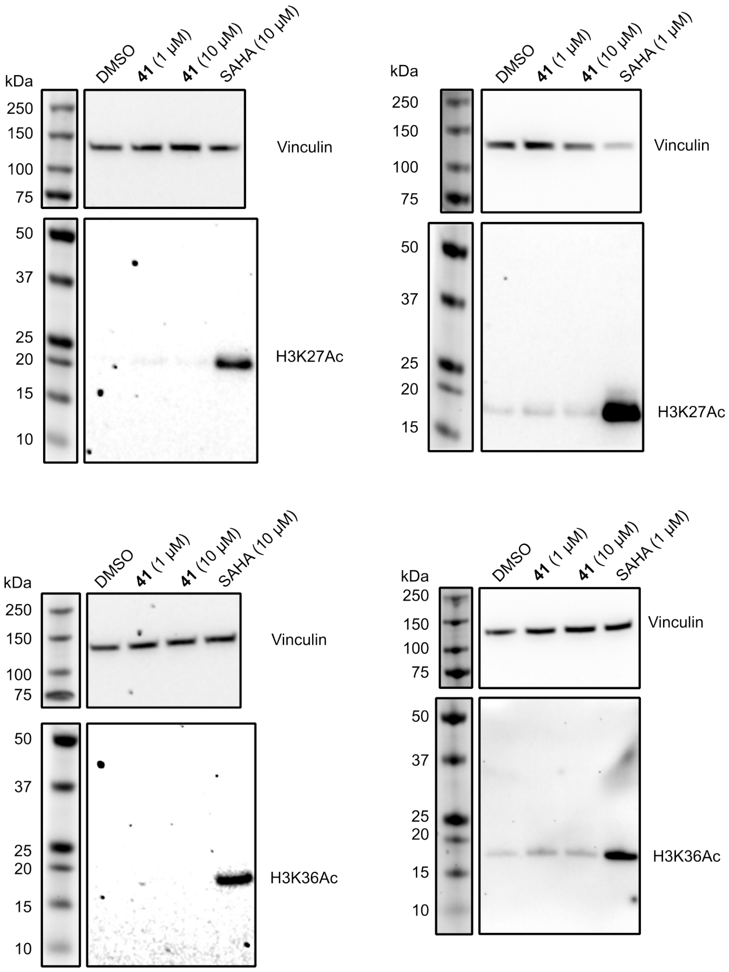

**Figure S10 continued.** Full blots from all replicates of the histone acetylation assays. Western blots of histone acetylation (H3K18, H3K27, and H3K36) or histone 3 (H3) upon 5 h treatment with lariat **41** (1 or 10 μM), SAHA (10 or 1 μM), or DMSO

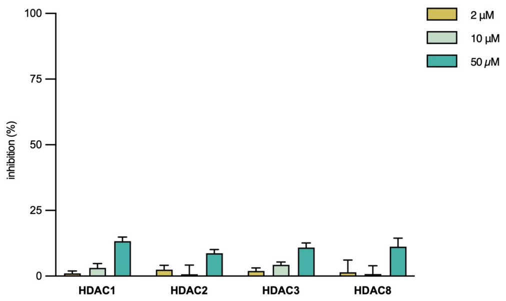

**Figure S11.** Three-dose inhibition assay of peptide **41** against HDAC1-3 and HDAC8
Inhibition at three different concentrations of peptide **41** (50 μM, 10 μM, and 2 μM) of the deacetylation activity of recombinant HDAC1 (5 nM), HDAC2 (2 nM), and HDAC3 (1 nM) against the Ac-LGK(Ac)-AMC substrate (20 μM) and the deacylation acitivity of HDAC8 (0.25 nM) against the substrate Ac-LGK(TFA)-AMC (100 μM). The enzymes were incubated with substrate at 37 °C for 30 minutes followed by development with 0.2 mg·mL$^{-1}$ trypsin for 15 minutes.

**Table S1. Substrate screening data.**

| Substrate | SIRT7 | SIRT7 + DNA | SIRT7 + tRNA |
|---|---|---|---|
| Ac-Leu-Gly-Lys(ac)-AMC | 213 ± 14 | 179 ± 11 | 257 ± 167 |
| Ac-Leu-Gly-Lys(but)-AMC | 96 ± 6 | 79 ± 1 | 62 ± 37 |
| Ac-Leu-Gly-Lys(cro)-AMC | 120 ± 34 | 149 ± 81 | 215 ± 6 |
| Ac-Leu-Gly-Lys(ma)-AMC | 184 ± 47 | 243 ± 37 | 103 ± 9 |
| Ac-Leu-Gly-Lys(suc)-AMC | 137 ± 34 | 191 ± 68 | 161 ± 40 |
| Ac-Leu-Gly-Lys(glut)-AMC | 107 ± 22 | 123 ± 14 | 123 ± 37 |
| Ac-Leu-Gly-Lys(adi)-AMC | 299 ± 147 | 60 ± 14 | 72 ± 17 |
| Ac-Leu-Gly-Lys(mg)-AMC | 75 ± 3 | 91 ± 60 | 143 ± 10 |
| Ac-Leu-Gly-Lys(hmg)-AMC | 61 ± 15 | 152 ± 39 | 240 ± 29 |
| Ac-Leu-Gly-Lys(bio)-AMC | 100 ± 19 | 48 ± 6 | 151 ± 17 |
| Ac-Leu-Gly-Lys(tfa)-AMC | 104 ± 37 | 40 ± 21 | 68 ± 11 |
| Ac-Arg-His-Lys-Lys(ac)-AMC | 134 ± 11 | 148 ± 6 | 221 ± 74 |
| Ac-Arg-His-Lys(ac)-Lys(ac)-AMC | 45 ± 127 | 68 ± 48 | 177 ± 68 |
| Ac-Glu-Thr-Asp-Lys(ac)-AMC | 207 ± 57 | 101 ± 1158 | 67 ± 24 |
| Ac-Glu-Thr-Asp-Lys(*i*-but)-AMC | 56 ± 32 | -22 ± 108 | 51 ± 8 |
| Ac-Glu-Thr-Asp-Lys(cro)-AMC | 104 ± 88 | 58 ± 65 | 91 ± 13 |
| Ac-Glu-Thr-Asp-Lys(dec)-AMC | 175 ± 24 | 167 ± 94 | 262 ± 137 |
| Ac-Glu-Thr-Asp-Lys(glu)-AMC | 92 ± 38 | 135 ± 65 | 68 ± 7 |
| Ac-Thr-Ala-Arg-Lys(ac)-AMC | 185 ± 52 | 490 ± 30 | 242 ± 82 |
| Ac-Thr-Ala-Arg-Lys(cro)-AMC | 342 ± 53 | 972 ± 57 | 800 ± 66 |
| Ac-Thr-Ala-Arg-Lys(hex)-AMC | 418 ± 79 | 1125 ± 83 | 562 ± 56 |
| Ac-Thr-Ala-Arg-Lys(oct)-AMC | 770 ± 40 | 1461 ± 50 | 1198 ± 196 |
| Ac-Thr-Ala-Arg-Lys(lau)-AMC | 534 ± 69 | 580 ± 144 | 616 ± 201 |
| Ac-Thr-Ala-Arg-Lys(myr)-AMC | 112 ± 7 | 328 ± 47 | 328 ± 47 |
| Ac-Thr-Ala-Arg-Lys(lip)-AMC | 562 ± 90 | 758 ± 13 | 703 ± 41 |
| Ac-Thr-Ala-Arg-Lys(hmg)-AMC | 243 ± 35 | 289 ± 6 | 214 ± 22 |
| Ac-Thr-Ala-Arg-Lys(bio)-AMC | 325 ± 42 | 635 ± 7 | 446 ± 77 |
| Ac-Gln-Pro-Lys-Lys(ac)-AMC | 122 ± 72 | 424 ± 29 | 323 ± 26 |
| Ac-Gln-Pro-Lys-Lys(pro)-AMC | 209 ± 14 | 367 ± 122 | 150 ± 31 |
| Ac-Gln-Pro-Lys-Lys(but)-AMC | 216 ± 15 | 484 ± 14 | 335 ± 115 |
| Ac-Gln-Pro-Lys-Lys(*i*-but)-AMC | 330 ± 112 | 445 ± 22 | 280 ± 84 |
| Ac-Gln-Pro-Lys-Lys(*i*-val)-AMC | 271 ± 95 | 827 ± 115 | 457 ± 186 |
| Ac-Gln-Pro-Lys-Lys(cro)-AMC | 276 ± 79 | 399 ± 68 | 250 ± 35 |
| Ac-Gln-Pro-Lys-Lys(hex)-AMC | 280 ± 70 | 1151 ± 47 | 295 ± 87 |
| Ac-Gln-Pro-Lys-Lys(oct)-AMC | 458 ± 74 | 1007 ± 273 | 1066 ± 63 |
| Ac-Gln-Pro-Lys-Lys(dec)-AMC | 1074 ± 80 | 1667 ± 155 | 1686 ± 118 |
| Ac-Gln-Pro-Lys-Lys(lau)-AMC | 366 ± 70 | 1454 ± 124 | 2021 ± 197 |
| Ac-Gln-Pro-Lys-Lys(myr)-AMC | 273 ± 81 | 304 ± 87 | 551 ± 40 |
| Ac-Gln-Pro-Lys-Lys(palm)-AMC | 130 ± 31 | 239 ± 22 | 331 ± 34 |
| Ac-Gln-Pro-Lys-Lys(for)-AMC | 255 ± 30 | 257 ± 112 | 276 ± 106 |
| Ac-Gln-Pro-Lys-Lys(glut)-AMC | -20 ± 17 | 237 ± 54 | 230 ± 64 |

Data depicts the conversion of fluorogenic substrate (in nM ± SD) from two independent experiments performed in duplicate.

#### Experimental methods

##### SIRT7 Expression

*E. coli* codon optimized pET100/D-TOPO vector encoding SIRT7 (aa 1–400) and an N-terminal cleavage site for Tobacco Etch Virus (TEV) protease was designed in GeneArt and purchased from ThermoFisher. SIRT7 was expressed in *E. Coli* BL21(DE3). The cells were grown at 37 °C at 150 rpm until OD = 0.6–0.8 followed by addition of 1 mM isopropyl β-D-1-thiogalactopyranoside and incubation at 18 °C at 150 rpm for 48 h. The cells were harvested by centrifugation at 5000 g for 30 min at 4 °C and disrupted by sonication for 6×1 min in lysis buffer (50 mM Tris, 400 mM NaCl, 20 mM β-mercaptoethanol (BME), 10% glycerol, pH 8) followed by centrifugation at 20,000 g at 4 °C for 30 min. The supernatant was filtered through a 0.45 μM filter and subjected to affinity chromatography using a 5 mL HisTrap HP His-tag purification column operated by a peristaltic pump at 4 °C. The column was prepared by washing with 25 mL water followed by 25 mL lysis buffer. Supernatant was run onto the column followed by washing with 25 mL lysis buffer before eluting the target protein with 25 mL elution buffer [50 mM Tris, 400 mM NaCl, 20 mM β-ME, 10% glycerol, 500 mM imidazole, pH 8.0]. The eluted protein was analyzed by SDS-PAGE and Coomassie staining and dialyzed against lysis buffer [50 mM Tris, 400 mM NaCl, 20 mM BME, 10% glycerol, pH 8.0] at 4 °C for 2 h twice using Spectra/Por 4 dialysis tubing (12–14 kDa MWCO) followed by treatment with TEV protease (0.2 mg/mL) overnight at 4 °C. The sample was then subjected to reverse affinity chromatography using a 5 mL HisTrap HP His-tag purification column at 4 °C to remove the hexa-histidine tag and the TEV protease from the sample. The column was prepared by washing with 25 mL water followed by 25 mL lysis buffer before running the digested sample onto the column. The column was washed with 25 mL lysis buffer before eluting the affinity tag and non-digested protein with 25 mL elution buffer [50 mM Tris, 400 mM NaCl, 20 mM β-ME, 10% glycerol, 500 mM imidazole, pH 8.0). The flow through and following wash were collected and concentrated using an Ultra-15 centrifugal filter unit (cut off value 50 kDa). The concentrated sample was stored at -80 °C. The purified protein exhibited deacetylase activity and we found that it retained activity upon storage at –80 °C for up to 6 months. However, multiple freeze–thaw cycles caused a rapid decrease in activity and the protein should therefore be aliquoted before initial storage.

##### Expression and purification of Avi-tagged full-length human SIRT7

The gene encoding human SIRT7 (aa 1–400, UniProt Q9NRC8) was codon optimized for *E.coli* (GeneArt), cloned into a pET28 plasmid with an N-terminal His10 tag and SUMO tag, and an C-terminal Avi-tag (full coding sequence below). SIRT7 was expressed in E. coli BL21(DE3) cells, co-transfected with a BirA expressing vector. An overnight cell culture was grown in LB medium at 37 °C under shaking (250 rpm). The overnight culture was then transferred to autoinduction medium (OvernightExpress Instant TB Medium, Novagen) and the cells were grown for 5 h at 37 °C under shaking (150 rpm). Then, biotin was added (40 μM final concentration) and the cells were allowed to grow at 16 °C for 16–24 h under shaking (250 rpm). The cells were harvested by centrifugation (6,000 g) for 20 min at 4° C. Cell lysis was performed with a French press cell disruptor (constant systems Ltd) by first suspending the cell pellet in lysis buffer I (50 mM Tris, 250 mM NaCl, 10% glycerol, 1 mM β-mercaptoethanol, 5 mM imidazole, pH 7.6) and adding ½ a tablet of cOmplete™ protease inhibitor cocktail (Roche) as well as lyzosyme (86 μg/mL) and DNase (2 μg/mL). The lysis solution was then run through the disruptor twice at 18 kpsi followed by centrifugation (18,000 g) at 4 °C for 1 h. The supernatant was then subjected to affinity chromatography, using a 5 mL HisTrap HP His-Tag purification column (Cytiva) operated by an ÄKTA fast protein liquid chromatography (FPLC) system, followed by washing with 5 column volumes of washing buffer I (50 mM Tris, 250 mM NaCl, 10% glycerol, 1 mM β-mercaptoethanol, pH 7.6). The protein was then eluted with a gradient of imidazole from 0 to 500 mM in washing buffer over 50 min at a flow rate of 2 mL/min. The eluted protein-containing fractions were analyzed by SDS-PAGE gel electrophoresis and Coomassie staining. The fractions containing pure SIRT7 construct were pooled and treated with His10-tagged

ULP-1 protease (0.1 μM) over night on ice. The cleavage byproducts were then removed by separating the protein on a size exclusion column (HiLoad 16/600 Superdex 200 prep grade, Cytiva), using an ÄKTA FPLC system, using washing buffer I. The protein fractions were analyzed by SDS-PAGE and Coomassie staining. Finally, the fractions were pooled, aliquoted, and stored at -80℃.
Coding region for His10 SUMO full length SIRT7 Avi:

ATGGGTAGCAGTCATCATCACCACCATCACCATCATCATCACAGCAGCGGTCTGGTTCCGCG
TGGTAGCCATATGAGCGATAGCGAAGTTAATCAAGAAGCCAAACCGGAAGTTAAGCCGGAAG
TGAAACCTGAAACACATATTAACCTGAAAGTGAGTGATGGCAGCAGCGAAATCTTCTTCAAAA
TCAAAAAAACAACACCGCTGCGTCGTCTGATGGAAGCATTTGCAAAACGTCAGGGTAAAGAAA
TGGATAGCCTGCGTTTTCTGTATGATGGTATTCGTATTCAGGCAGATCAGACACCGGAAGATC
TGGATATGGAAGATAACGATATTATCGAAGCACATCGTGAGCAGATTGGTGGTATGGCAGCC
GGTGGTCTGAGCCGTAGCGAACGTAAAGCAGCAGAACGTGTTCGTCGTCTGCGTGAAGAACA
GCAGCGTGAACGTCTGCGTCAGGTTAGCCGTATTCTGCGTAAAGCCGCAGCCGAACGTAGC
GCAGAAGAAGGTCGTCTGCTGGCAGAAAGCGCAGATCTGGTTACCGAACTGCAGGGTCGTA
GCCGTCGTCGTGAAGGTCTGAAACGTCGTCAAGAAGAAGTTTGCGACGATCCGGAAGAACTG
CGTGGTAAAGTTCGTGAACTGGCAAGCGCAGTTCGTAATGCAAAATATCTGGTTGTTTATACC
GGTGCAGGTATTAGCACCGCAGCAAGCATTCCGGATTATCGTGGTCCGAATGGTGTTTGGAC
CCTGCTGCAAAAAGGTCGTAGCGTTAGCGCAGCAGATCTGAGCGAAGCAGAACCGACACTGA
CCCATATGAGCATTACCCGTCTGCATGAACAGAAACTGGTTCAGCATGTTGTTAGCCAGAATT
GTGATGGTCTGCATCTGCGTAGCGGTCTGCCTCGTACCGCAATTAGCGAACTGCATGGCAAT
ATGTATATTGAAGTTTGTACCAGCTGTGTGCCGAATCGTGAATATGTTCGTGTTTTTGATGTGA
CCGAACGCACCGCACTGCATCGTCATCAGACCGGTCGTACCTGTCATAAATGTGGCACCCAG
CTGCGTGATACCATTGTTCATTTTGGTGAACGTGGCACCCTGGGTCAGCCGCTGAATTGGGA
AGCAGCCACCGAAGCAGCAAGCCGTGCAGATACCATTCTGTGTCTGGGTAGCAGCCTGAAAG
TTCTGAAAAAGTATCCGCGTCTGTGGTGTATGACCAAACCTCCGAGTCGTCGTCCGAAACTGT
ATATTGTTAATCTGCAGTGGACCCCGAAAGATGATTGGGCAGCACTGAAACTGCACGGTAAAT
GTGATGATGTTATGCGTCTGCTGATGGCAGAACTGGGTTTAGAAATTCCGGCATATAGCCGTT
GGCAGGATCCGATTTTTAGCCTGGCAACACCGCTGCGTGCCGGTGAAGAAGGTAGCCATAGC
CGTAAAAGCCTGTGCCGTAGCCGTGAAGAAGCACCGCCTGGCGATCGTGGTGCACCGCTGA
GCAGCGCACCGATTTTAGGTGGTTGGTTTGGTCGTGGTTGTACCAAACGCACCAAACGTAAA
AAAGTGACCGGCAGTAGCGGCCTGAACGATATTTTTGAAGCCCAGAAAATTGAATGGCATGAA
GGTAGCTAATGA

**Expression and purification of Avi-tagged core domain of human SIRT7**
As for full-length SIRT7, a gene encoding human SIRT7 (aa 74–363) was cloned into a pET28 plasmid with a N-terminal His10-SUMO tag and a C-terminal Avi-tag was (coding sequence below). Expression and cell harvesting were performed as described for Avi-tagged full-length human SUMO SIRT7. The purification by affinity chromatography and size exclusion chromatography were performed as above but with different buffers (lysis buffer II: 50 mM MOPS, 250 mM NaCl, 10% glycerol, 1 mM β-mercaptoethanol, 5 mM imidazole, pH 6.5; and washing buffer II, 50 mM MOPS, 250 mM NaCl, 10% glycerol 1 mM β-mercaptoethanol, pH 6.5). To remove His10-UPL-1 cleavage byproducts, a reverse affinity column chromatography step was performed, using 2 mL His-Select® Nickel Affinity Gel (Sigma-Aldrich) slurry for a final bed volume of 1 mL. The resin was prepared by first washing with 10 column volumes of MilliQ water, followed by 20 column volumes of washing buffer II. The protein sample was then loaded onto the column followed by 20 column volumes of washing buffer II. The protein was then eluted with 8 column volumes elution buffer. The protein was then analyzed by SDS-PAGE and Coomassie staining, aliquoted, and stored at –80℃.
Coding region for His10 SUMO core SIRT7 Avi:

ATGGGTAGCAGTCATCATCACCACCATCACCATCATCATCACAGCAGCGGTCTGGTTCCGCG
TGGTAGCCATATGAGCGATAGCGAAGTTAATCAAGAAGCCAAACCGGAAGTTAAGCCGGAAG
TGAAACCTGAAACACATATTAACCTGAAAGTGAGTGATGGCAGCAGCGAAATCTTCTTCAAAA

TCAAAAAAACAACACCGCTGCGTCGTCTGATGGAAGCATTTGCAAAACGTCAGGGTAAAGAAA
TGGATAGCCTGCGTTTTCTGTATGATGGTATTCGTATTCAGGCAGATCAGACACCGGAAGATC
TGGATATGGAAGATAACGATATTATCGAAGCACATCGTGAGCAGATTGGTGGTCGTCAAGAAG
AAGTTTGCGACGATCCGGAAGAACTGCGTGGTAAAGTTCGTGAACTGGCAAGCGCAGTTCGT
AATGCAAAATATCTGGTTGTTTATACCGGTGCAGGTATTAGCACCGCAGCAAGCATTCCGGAT
TATCGTGGTCCGAATGGTGTTTGGACCCTGCTGCAAAAAGGTCGTAGCGTTAGCGCAGCAGA
TCTGAGCGAAGCAGAACCGACACTGACCCATATGAGCATTACCCGTCTGCATGAACAGAAAC
TGGTTCAGCATGTTGTTAGCCAGAATTGTGATGGTCTGCATCTGCGTAGCGGTCTGCCTCGTA
CCGCAATTAGCGAACTGCATGGCAATATGTATATTGAAGTTTGTACCAGCTGTGTGCCGAATC
GTGAATATGTTCGTGTTTTTGATGTGACCGAACGCACCGCACTGCATCGTCATCAGACCGGTC
GTACCTGTCATAAATGTGGCACCCAGCTGCGTGATACCATTGTTCATTTTGGTGAACGTGGCA
CCCTGGGTCAGCCGCTGAATTGGGAAGCAGCCACCGAAGCAGCAAGCCGTGCAGATACCAT
TCTGTGTCTGGGTAGCAGCCTGAAAGTTCTGAAAAAGTATCCGCGTCTGTGGTGTATGACCAA
ACCTCCGAGTCGTCGTCCGAAACTGTATATTGTTAATCTGCAGTGGACCCCGAAAGATGATTG
GGCAGCACTGAAACTGCACGGTAAATGTGATGATGTTATGCGTCTGCTGATGGCAGAACTGG
GTTTAGAAATTCCGGCATATAGCCGTTGGCAGGATCCGATTTTTAGCCTGGCAACACCGCTGC
GTGCCGGTGAAGAAGGTAGCCATAGCCGTAAAAGCCTGTGCCGTAGCGGCAGTAGCGGCCT
GAACGATATTTTTGAAGCCCAGAAAATTGAATGGCATGAAGGTAGCTAATGA

#### In Vitro Enzymatic Characterization

All fluorescence-based sirtuin deacylase activity assays were performed in black low binding 96-well microtiter plates (Corning half-area wells), with duplicate series in each assay and each assay performed at least twice. Control wells without enzyme were included in each plate. All reactions were performed in assay buffer (see above) with appropriate concentrations of substrates and inhibitors obtained by dilution from 2–200 mM stock solutions in milliQ water or DMSO, and appropriate concentration of enzyme obtained by dilution of the stock provided by the supplier. Previously published sirtuin/HDAC substrates and inhibitors were synthesized in previous reports as highlighted in the Key resources table. Plates were analyzed using a BMG Labtech FLUOstar Omega plate reader with excitation at 360 nm and detecting emission at 460 nm. Fluorescence measurements (RFU) were converted to [AMC] concentrations based on an [AMC]-fluorescence standard curve, and all data analysis was performed using Microsoft Excel and GraphPad Prism.

*Buffers:*

- Tris buffer was prepared as described in Biomol International product sheets BML-KI-143 [50 mM Tris/Cl, 137 mM NaCl, 2.7 mM KCl, $MgCl_2$ 1 mM, 1.0 mg/mL BSA, pH 8.0].
- HEPES buffer was prepared as previously described [2] [50 mM HEPES/Na, 100 mM KCl, 0.001% (v/v) Tween-20, 200 μM tris(2-carboxyethyl)phosphine (TCEP), 1.0 mg/mL BSA, pH 7.4].
- 20 mM phosphate buffer, 0.5 mg/mL BSA, pH 7

#### Endpoint SIRT7 Substrate Screen

Substrate, $NAD^+$ and yeast tRNA were added to each well and the experiment was initiated by addition of a freshly prepared solution of recombinant SIRT7, for a final volume of 25 μL per well in HEPES buffer. The following final concentrations were used: SIRT7 enzyme (500 nM), substrate (50 μM), $NAD^+$ (500 μM), and/or DNA (0.97 mg/mL) or tRNA (4 μM). The plate was incubated at 37 °C for 120 min, then a solution of trypsin and NAM (25 μL, 5 mg/mL and 4 mM, respectively; final concentration 2.5 mg/mL and 2 mM, respectively) was added and the assay development was allowed to proceed for 90 min at RT, before fluorescence measurement and calculation of residual activity.

**Endpoint Inhibition Assay**

Endpoint inhibition assays were performed as previously described.[3] In brief, substrate, $NAD^+$, and inhibitor were added to each well and the experiment was initiated by addition of a freshly prepared solution of relevant sirtuin for a final volume of 25 μL per well. For SIRT7, assays were run with or without DNA (0.75 μg/mL) or tRNA (37 μg/mL) in HEPES buffer or nucleosomes (200 nM) in phosphate buffer. For SIRT5, assays were performed in HEPES buffer, and for SIRT1–3 and 6, assays were performed in Tris buffer. The following final concentrations were used: SIRT enzyme (100–200 nM for SIRT1; 100–200 nM for SIRT2; 300–400 nM for SIRT3; 500–600 nM for SIRT6; 500 nM for SIRT7), substrate (50 μM), $NAD^+$ (500 μM), and inhibitor (10, 33 or 100 μM or fold dilution series for dose–response assays). The plate was incubated at 37 °C for 60 min (120 min for SIRT7), then a solution of trypsin and nicotinamide (NAM; 25 μL, 5 $mg{\cdot}mL^{-1}$ and 4 mM, respectively; final concentrations 2.5 $mg{\cdot}mL^{-1}$ and 2 mM, respectively) was added, and the assay development was allowed to proceed for 90 min at RT before fluorescence measurement and calculation of residual activity. For concentration-response assays, $IC_{50}$ values were obtained by fitting the resulting data to the concentration–response equation using GraphPad Prism.

**HDAC Endpoint Inhibition Assay**

Endpoint inhibition assays against HDACs were performed in 96-well plates (Fischer Scientific, cat. #: 3686) in HEPES buffer containing BSA (0.5 mg/mL). Final volume in each well was 25 µL and a final DMSO concentration of 2% was used. Compound **41** (50 µM, 10 µM, and 2 µM final concentrations) was incubated with enzyme [HDAC1 (BPS cat #50051, 5 nM), HDAC2 (BPS Bioscience cat #50052, 2 nM), HDAC3 (BPS Bioscience cat #50003, 1 nM), HDAC8 (BPS Bioscience cat #50008, 0.25 nM)] and substrate [AC-LGK(AC)-AMC (20 µM) for HDAC1–3; Ac-LGK(TFA)-AMC (100 µM) for HDAC8] at 37 °C for 30 minutes before addition of a solution of trypsin (25 µL, 0.2 $mg{\cdot}mL^{-1}$) in assay buffer. The plate was then incubated at 25 °C for 15 minutes followed by measurement of fluorescence (excitation: 355 nm; emission: 450 nm).

**Michaelis-Menten Assays**

Rate experiments for determination of kinetic parameters were performed in HEPES buffer (no BSA added) in a final volume of 50 μL per well, where Ac-QPKKdec-AMC or Ac-APRKdec-AMC (1.5-fold dilutions) was incubated with SIRT7 (500 nM) and tRNA (4.0 μM) in HEPES buffer or nucleosomes (200 nM) in phosphate buffer. In situ fluorophore release was monitored by fluorescence readings recorded at time points 10, 20, 30 and 40 min at 37 °C to obtain initial rates $v_0$ ($nM{\cdot}s^{-1}$) for each concentration. The data were fitted to the Michaelis-Menten equation to afford $K_M$ (μM) and $k_{cat}$ ($s^{-1}$) values.

**Cellular thermal shift assays (CETSA)**

**Cell culture**

HEK293T (ATCC, CRL-1573) cells were cultured at 37 °C under a humidified 5% CO2 atmosphere in Dulbecco's modified Eagle's medium (DMEM, Thermo Scientific, 11965118) supplemented with 10% (v/v) fetal bovine serum (FBS, Thermo Fisher Scientific, 26140079), 1% penicillin-streptomycin (Sigma-Aldrich, P4333). Cells were sub-cultured every 2-4 days.

**Treatment and Western blotting**

Cellular thermal shifts were measured as previously described.[4] In brief HEK293T cells were plated in 6-well plates and incubated to 80-90% confluency, where medium was replaced with culture medium containing **39** (10 µM in 2% DMSO), **41** (1 or 10 µM in 2% DMSO) or vehicle (2% DMSO); 3 wells/treatment. The cells were treated for 2 hours before media was removed by aspiration and cells were collected in PBS (2 mL/treatment) by scraping, and pelleted by centrifugation (300 g, 5 min). Pellets were resuspended in PBS (2 mL) and pelleted by centrifugation before the washed pellets were suspended in PBS supplemented with cOmplete EDTA-free protease inhibitor cocktail (Sigma-Aldrich, COEDTAF-RO, 330 µL/cell treatment). Cell suspensions were aliquoted into PCR tubes (30 µL) and heated at temperatures ranging from 37.0 °C to 65.0 °C for 3 minutes followed by 3 minutes at 25 °C in a thermal cycler (Eppendorf Mastercycler Nexus Thermal Cycler). The cellular suspensions were then lysed by three freeze/thaw cycles, snap-freezing in a dry-ice/acetone bath followed by thawing at 25 °C in the thermal cycle. The suspensions were subjected to centrifugation (20,000g, 20 min) at 4 °C and the supernatants were collected as whole-cell lysate and mixed with NuPAGE LDS sample buffer (ThermoFisher, NP0007) and sample reducing agent (ThermoFisher, NP0004) followed by heating to 95 °C for 10 min. Samples were then resolved by gel electrophoresis (SDS-PAGE) in NuPAGE gels (12 wells, 4–12% Bis-Tris, 1.0 mm, ThermoFisher, NP0322BOX) with MES running buffer (ThermoFisher, NP000202). Proteins were transferred to PVDF membranes (ThermoFisher, IB24001) using the iBlot2 system and blocked with 5% skim milk in tris-buffered saline with 0.1% tween-20 (TBST) at 25 °C for 1 h. Membranes were then washed with TBST (3 × 5 min), incubated with primary anitbody (Anit-SIRT7, Cell Signaling Technology, CST-5360S, 1:500) in TBST with 5% BSA (4 °C, overnight), washed with TBST (3 × 5 min), incubated with HRP-conjugated secondary antibody (Anti-rabbit IgG HRP-linked antibody, Cell Signaling Technology, CST-7074S, 1:10,000) in TBST with 2% skim milk (25 °C, 1 h), and washed with TBST (3 × 5 min) and TBS (1 × 5 min). Membranes were visualized using enhanced chemiluminescent reagents (Pierce ECL Western blotting substrate, ThermoFisher, 32106) on a syngene PXi4 image analysis system. Protein marker used: Precision Plus Protein All Blue Standard (BioRAD, 161-0373).

**Histone acetylation in HEK293T cells**

Histone acetylation levels were determined as previously described.[5] HEK293T cells were plated in 6-well plates and grown to 70-80% confluency as described above before medium was replaced with culture medium containing 41 (1 or 10 µM in 2% DMSO), SAHA (1 or 10 µM in 2% DMSO) or vehicle (2% DMSO). The cells were treated for 2 hours at 37 °C after which media was aspirated and the cells were collected by scraping in PBS (1 mL), pelleted by centrifugation (300 g, 5 min), aspirated PBS, resuspended in PBS (1 mL), pelleted by centrifugation, resuspended in PBS (1 mL), and treated with RIPA lysis buffer (50 mM Tris, 150 mM NaCl, 1% Triton X-100, 0.5% sodium deoxycholate, 0.1% SDS) supplemented cOmplete EDTA-free protease inhibitor cocktail (Sigma-Aldrich, COEDTAF-RO, 150 µL/cell treatment). Lysis of the cell lysates were done by sonification with a Bandelin Sonopuls mini20 (2 s on, 2 s off, 80% amplitude, 1 min), centrifuged (14000 g, 10 min, 4 °C), and protein concentrations of the supernatants were determined by the bicinchoninic acid assay (BCA1, Sigma-Aldrich). The concentration of the lysates were adjusted with PBS to the lowest concentration, and 20 µg of lysate was visualized by SDS-PAGE as described above. Primary antibodies used: Vinculin (E1E9V) XP (Cell Signaling Technology, #13901, 1:1000), acetyl-histone H3 (Lys18) (D8Z5H) (Cell Signaling Technology, #13998, 1:500), acetyl-histone H3 (K36) (D9T5Q) (Cell Signaling Technology, #27683, 1:500), acetyl-histone H3 (K27) (Cell Signaling Technology, #4353, 1:500), histone H3 (Cell Signaling Technology, #3638, 1:500). Add Ac-H3K18 and H3. Secondary antibodies used: Anti-rabbit IgG HRP-linked antibody (Cell Signaling Technology, #7074S, 1:10000), Anti-mouse IgG HRP-linked antibody (Cell Signaling Technology, #7076S, 1:10000). Protein marker used: Precision Plus Protein All Blue Standard (BioRAD, 161-0373).

**Small angle X-ray scattering**

Recombinant SIRT7 was expressed and purified as previously described including purification by size exclusion chromatography. The sample was buffer exchanged into 50 mM Tris-HCl, pH 8.0 and SAXS measurements were carried out in a concentration series ranging from 1.0-3.9 mg/mL on the EMBL P12 BioSAXS beamline [6] at PETRA III, DESY. All SAXS data analysis and modelling was carried out using programs from the ATSAS software suite v. 2.8.4 [7]. PrimusQT [8] was used for general SAXS data analysis, extraction of molecular parameters and generating a merged SAXS curve from the SAXS curves recorded at low and high concentration. All modelling was based on the merged experimental merged curve. Dammif [9] and Dammin [10] were used for generating an *ab initio* model. 10 models were generated with Dammif in interactive mode and averaged using Damaver [11]. The averaged model was further refined using Dammin in slow mode to give the final *ab initio* model. Crysol [12] was used for fitting atomic structures to experimental data using 500 points in the theoretical scattering curve, constant background subtraction and a maximum scattering angle of q=0.6 $Å^{-1}$. Combined *ab initio* and rigid body modelling was carried out using Coral [13] with default settings. Five models were generated using the Alphafold2 structure (PDB Q9NRC8) as a starting point. The model with the best fit to the experimental merged curve was chosen as the final Coral model.

**Identification of peptide binders via the RaPID screening platform**

**Preparation of flexizyme (eFx)**

Oligonucleotide sequences:

Fx5_F36 = GTAATACGACTCACTATAGGATCGAAAGATTTCCGC
eFx_R45 = ACCTAACGCTAATCCCCTTTCGGGGCCGCGGAAATCTTTCGATCC
T7ex5_F22 = GGCGTAATACGACTCACTATAG
eFx_R18 = ACCTAACGCTAATCCCCT

All oligonucleotides used for preparation of eFx were purchased from IDT. First Fx5_F36 was annealed with eFx_R45, followed by primer extension with Klenow fragment (NEB). The DNA product was then amplified by PCR, using Taq polymerase (VWR) with T7ex5_F22 as the forward primer and eFx_R18 as the reverse primer. The amplified DNA product was then transcribed using T7 RNA polymerase (Thermo Scientific) and purified with an 12% acrylamide, 6 M Urea denaturing PAGE.
DNA template for eFx:
GGCGTAATACGACTCACTATAGGATCGAAAGATTTCCGCGGCCCCGAAAGGGGATTAGCGTT AGGT
RNA sequence of eFx:
GGAUCGAAAGAUUUCCGCGGCCCCGAAAGGGGAUUAGCGUUAGGU

**Preparation of tRNAiniCAU**

Oligonucleotide sequences:
Ini_1_F49 = GTAATACGACTCACTATAGGCGGGGTGGAGCAGCCTGGTAGCTCGTCGG
Ini_1_R44 = GAACCGACGATCTTCGGGTTATGAGCCCGACGAGCTACCAGGCT
T7ex5_F22 = GGCGTAATACGACTCACTATAG
Ini_2_R38 = TGGTTGCGGGGGCCGGATTTGAACCGACGATCTTCGGG
Ini_3_R20_OMe = TG*GTTGCGGGGGCCGGATTT
(G* 2'-OMe)

All oligonucleotides used for preparation of $tRNA^{ini}_{CAU}$ were purchased from Intelligent DNA Technologies. Ini_1_F49 and Ini_1_R44 were annealed followed by a primer extension reaction with Klenow fragment (NEB). The DNA product was then amplified by two sequential PCR reactions using Taq polymerase (VWR). Both PCR reactions used T7ex5_F22 as the forward primer. In PCR1, ini_2_R38 was used as the reverse primer while in PCR2 ini_3_R20_OMe was the reverse primer. The amplified DNA product was then transcribed using T7 RNA polymerase (Thermo Scientific) and thermostable inorganic pyrophosphatase (NEB) The RNA product was subsequently purified with an 8% acrylamide 6 M Urea denaturing PAGE before lastly, it was dissolved in RNase-free water for a final concentration of 250 μM and stored at -80 °C.
DNA template for $tRNA^{ini}_{CAU}$:

GGCGTAATACGACTCACTATAGGCGGGGTGGAGCAGCCTGGTAGCTCGTCGGGCTCATAAC CCGAAGATCGTCGGTTCAAATCCGGCCCCCGCAACCA

RNA sequence of $tRNA^{ini}_{CAU}$:

GGCGGGGUGGAGCAGCCUGGUAGCUCGUCGGGCUCAUAACCCGAAGAUCGUCGGUUCAA AUCCGGCCCCCGCAACCA

**Aminoacylation of $tRNA^{ini}_{CAU}$ with *N*-chloroacetyl-D-Phe-cyanomethyl ester**

The prepared initiator tRNA was aminoacylated with *N*-chloroacetyl-D-Phe. The aminoacylation reaction was carried out by mixing heat-refolded $tRNA^{ini}_{CAU}$ (25 μM), eFx (25 μM), and *N*-chloroacetyl-D-Phe-cyanomethyl ester (5 mM) in HEPES-KOH pH 7.6 (50 mM), $MgCl_2$ (600 mM), 20% DMSO. The reaction mixture was then left for 2 h at 0 °C. To terminate the reaction, 4 reaction volumes of NaOAc (0.3 M) and the product was precipitated with 10 reaction volumes of EtOH. After centrifugation, the pellet was washed twice with NaOAc (0.1 M, pH 5.2) in 70% EtOH and once with 70% EtOH. The pellet was then dried and stored at -80 °C.

**mRNA library preparation and puromycin ligation**

mRNA coding random peptides was prepared by in vitro transcription of custom synthesized DNA templates with degenerate NNK codons (IDT). The mRNA from libraries with NNK lengths of 4, 5, 6, 7, 8, 9, 10, 11, and 12 was mixed. The ratio for each NNK size were respectively: $1.4\times10^{-8}$: $4.5\times10^{-7}$: $1.4\times10^{-5}$: $4.6\times10^{-4}$: 0.015:1:1:1:1. The overall aim is to include as many unique sequences as possible in the final library. The ratios for smaller NNK libraries are therefore lower to avoid overrepresentation of these sequences. The mRNA library was subsequently ligated to puromycin via a nucleic acid linker (5'-phosphate-CTCCCGCCCCCCGTCC-$(iSp18)_5$-CC-3'-puromycin) through the action of T4 RNA ligase 1 (NEB). The ligated library was purified with a phenol chloroform extraction and ethanol precipitation. The library was stored at -80 °C until needed.

N = C, G, A, or T

K = G or T

General sequence for NNK DNA:

GAATTTAATACGACTCACTATAGGGTTAACTTTAAGAAGGAGATATACATATG$(NNK)_n$TGCGGC AGCGGCAGCGGCAGCTAGGACGGGGGGCGGAAA

**Reprogrammed in vitro translation system**

A customized *in vitro* translation system was assembled based on the PUREFrex 2.0 translation system (Cosmo Bio Ltd). The system contained: a custom ordered PUREFrex 2.0 solution II (proteins) lacking release factor 1, PUREFrex 2.0 solution III (ribosomes). PUREFrex 2.0 was supplemented with 50 mM HEPES-KOH pH 7.6, 100 mM KOAc, 2 mM ATP, 2 mM GTP, 1 mM CTP, 1 mM UTP, 20 mM creatine phosphate, 2 mM spermidine, 1 mM DTT, $Mg(OAc)_2$ 5 mM, 1.2 mg/mL total *E.coli* tRNA (Roche). Note a formyl donor was not added to the mixture to prevent initiation with formyl methionine. To this translation solution was added 19 of the 20 canonical amino acids (500 μM), excluding methionine, and the appropriate puromycin-ligated mRNA. For genetic code reprogramming this was supplemented with *in vitro* transcribed initiator $tRNA_{CAU}$ loaded with N-ClAc-D-Phe-OH (50 μM).

**Binding of biotin-SIRT7 to streptavidin magnetic beads**

To determine the binding capacity of streptavidin magnetic beads for biotinylated SIRT7 and biotinylated core SIRT7 protein various volumes of DynaBeads™M-280 Streptavidin (Invitrogen) were washed with selection buffer (50 mM HEPES-KOH, pH 7.6, 250 mM NaCl, 0.05% Tween-20, 1 mM DTT). The beads were then treated with 10 μL of biotinylated protein (1 μM). The beads were incubated with protein for 15 min at 4 °C. After 15 min biotin (25 μM) was added, and the beads were incubated for another 15 min at 4 °C. Supernatant and beads were separated and run on a 4–20%

Mini-PROTEAN TGX gel and was stained using Coomassie stain. Binding capacities were determined to 2.25 pmol/μL bead slurry for full length SIRT7 and 2.37 pmol/μL bead slurry for the core SIRT7 domain.

**First round of selection**

The initial mRNA library ligated to puromycin was added to a 25 μL *in vitro* reprogrammed translation reaction. The reaction was incubated at 37 °C for 40 min. The ribosomes are then dissociated from the mRNA strands by adding 5 μL of 100 mM EDTA and leaving the mixture at 37 °C for 30 min. The peptide ligated mRNA strands were then reverse transcribed using Superscript IV (Thermo Scientific) reverse transcriptase and 2× blocking buffer added (46 μL, 100 mM HEPES, 500 mM NaCl, 0.1% Tween-20, 0.2% acetylated BSA, 2 mM DTT, 0.2% yeast tRNA, 0.2% salmon sperm DNA, pH 7.6) was added. Magnetic beads were prepared corresponding to a final concentration of 200 nM for full length SIRT7 or core SIRT7. The beads were washed in ice cold selection buffer (50 mM HEPES, pH 7.6, 250 mM NaCl, 0,05% Tween-20, 1 mM DTT) before adding 36.8 pmol of the biotinylated target protein. The beads were incubated with the protein at 4 °C for 15 min. Biotin was then added for a final concentration of 25 mM. The beads were then incubated another 15 min at 4 °C. The reverse transcribed peptide library was then added to the beads loaded with biotin-SIRT7. The beads were then rapidly washed three times with ice cold selection buffer before being resuspended in PCR buffer and heat eluted at 95 °C for 5 min. The DNA containing supernatant was then collected and a qPCR analysis was used to quantify the amount of DNA recovered from the first selection round. The eluted DNA sample was then amplified through a PCR reaction using Phusion polymerase (NEB) and was then transcribed with T7 polymerase (Thermo Scientific) overnight at 37 °C. The resulting mRNA was purified by phenol/chloroform extraction and ethanol precipitation followed by puromycin ligation using T4 RNA ligase 1 (NEB) and another round of phenol/chloroform extraction and ethanol precipitation.

**Subsequent selection rounds including negative selection**

The puromycin ligated mRNA library from the last round (5 pmol) was added to an *in vitro* translation reaction (5 μL scale). The translation reaction was incubated at 37 °C for 40 min before addition of 1 μL of 100 mM EDTA (pH 8.0). The reaction was then incubated another 30 min at 37 °C. The peptide attached mRNA was then reverse transcribed for 1 h at 45 °C using Superscript IV (Thermo Scientific) reverse transcriptase after which the reverse transcription solution was blocked with 2x blocking buffer (100 mM HEPES, 500 mM NaCl, 0.1% Tween-20, 0.2% acetylated BSA, 2 mM DTT, 0.2% yeast tRNA. 0.2% salmon sperm DNA, pH 7.6). The next step was to perform a negative selection. Magnetic beads (9X the amounts used for SIRT7) were washed with selection buffer (50 mM HEPES, 250 mM NaCl, 0.05% Tween-20, 1 mM DTT, pH 7.6). Half of the beads were then incubated with biotin (final concentration 25 mM) for 15 min at 4 °C. The two bead samples were then recombined and washed with ice cold selection buffer. The reverse transcribed peptide library was then added to the 1/3 of the beads and incubated for 30 min at 4 °C. This constitutes one round of negative selection, and the supernatant was then used for two additional rounds of negative selection. Next, beads corresponding to a final concentration of 200 nM protein were washed with selection buffer and were subsequently incubated with 7.32 pmol biotinylated target protein for 15 min at 4 °C, followed by addition of biotin at a final concentration of 25 mM. The bead suspension was then incubated for another 15 min at 4 °C before being washed with selection buffer. The supernatant from the third negative selection was then added to the SIRT7 loaded beads and the bead suspension was incubated at 4 °C for 30 min. The beads were rapidly washed three times with selection buffer before they were resuspended in PCR reaction mixture and heat eluted at 95 °C for 5 min. The supernatant was then collected, and the amount of recovered DNA was quantified using qPCR and the eluted DNA sample was then amplified using Phusion PCR. Finally, the DNA was transcribed overnight at 37 °C using T7 polymerase. The resulting mRNA library was then extracted using a phenol/chloroform extraction followed by an ethanol precipitation. The mRNA was then resuspended and ligated to puromycin as previously described. The last step was another phenol/chloroform extraction and ethanol precipitation. The mRNA library was then ready for use in the next selection round.

#### Chemical Synthesis

#### Materials and methods

All reagents and solvents were of analytical grade and used without further purification as obtained from commercial suppliers. Anh. solvents were obtained from a PureSolv-system. Reactions were conducted under an atmosphere of nitrogen whenever anh. solvents were used. Reactions were monitored by thin-layer chromatography (TLC) using silica gel coated plates (analytical $SiO_2$-60, F-254) and by LC-MS. TLC plates were visualized under UV light or by dipping into a solution of potassium permanganate (10 g/L), potassium carbonate (67 g/L) and sodium hydroxide (0.83 g/L) in water. Evaporation of solvents was carried out under reduced pressure at a temperature below 40 °C. LC−MS analyses were performed on a Phenomenex Kinetex column (1.7 μm, 50×2.10 mm) using a Waters Acquity ultra high-performance liquid chromatography (UPLC) system. Gradient A with eluent I (0.1% HCOOH in $H_2O$) and eluent II (0.1% HCOOH in MeCN) rising linearly from 0% to 95% of II during *t* = 0.00−5.20 min was applied at a flow rate of 0.6 mL/min. Preparative reverse-phase HPLC purification was performed on a C8 or C18 Phenomenex Luna column (5 μm, 100 Å, 250×21.2 mm) using an Agilent 1260 LC system equipped with a diode array UV detector and an evaporative light scattering detector (ELSD). Gradient B with eluent III ($H_2O$/MeCN/TFA, 95:5:0.1, v:v) and eluent IV (0.1% TFA in MeCN) rising linearly from 0−30% to 95% of IV during *t* = 5−45 min at a flow rate of 20 mL/min was applied. Analytical HPLC was either performed on a C8 phenomenex Luna column (5 μM, 100 Å, 250×4.60 mm) using an Agilent 1100 series system equipped with a diode array UV detector (method A) or on a C18 Infinity Poroshell 120 column (2.7 μm, 100 × 3.0 mm) using an Agilent 1260 Infinity II series system equipped with a diode array UV detector (method B). Method A with gradient C using eluent III and eluent IV, rising linearly from 0% to 95% of IV during *t* = 5−20 min or *t* = 5−35 min was applied at a flow rate of 1.2 mL/min. Method B with gradient C using gradient C during *t* = 1–11 min was applied at a flow rate of 1.2 mL/min. High-resolution mass spectrometry (HRMS) measurements were recorded either on a maXis G3 quadrupole time-of-flight (TOF) mass spectrometer (Bruker Daltonics, Bremen, Germany) equipped with an electrospray ionization (ESI) source or on an Agilent 1290 UHPLC equipped with a diode array detector and coupled to Agilent 6550 QTOF mass spectrometer operated in positive electrospray or on a Bruker Solarix WR by either matrix assisted laser desorption/ionization, or ESI. Nuclear magnetic resonance (NMR) spectra were recorded either on a Bruker Avance III HD equipped with a cryogenically cooled probe ($^1$H NMR and $^{13}$C NMR recorded at 600 and 151 MHz, respectively) or a Bruker Avance III ($^1$H NMR and $^{13}$C NMR recorded at 400, 101 and 377 MHz, respectively). All spectra were recorded at 298 K. Chemical shifts are reported in ppm relative to deuterated solvent as internal standard ($\delta_H$ DMSO-$d_6$ 2.50 ppm; $\delta_C$ DMSO 39.52 ppm; $\delta_H$ $CDCl_3$ 7.26 ppm; $\delta_C$ $CDCl_3$ 77.16 ppm). Assignments of NMR spectra are based on 2D correlation spectroscopy (COSY, HSQC, TOCSY and HMBC spectra).

##### General solid phase peptide synthesis procedure (SPPS)

Peptides were synthesized on 2-chlorotrityl (2-CTC) polystyrene resin, TentaGel® or ChemMatrix® resin using a Rink amide (RAM) linker by standard SPPS. Resin loading was determined spectrophotometrically, quantifying the amount of released fluorene upon cleavage of the Fmoc group from a small sample.(Gude et al., 2002) Each elongation step was performed by applying the relevant amino acid (3 equiv.), HATU (2.9 equiv.) and *i*$Pr_2$NEt (6 equiv.). Fmoc-deprotection was performed by treatment with DMF/piperidine (4:1, v/v, 4.0 mL/0.1 mmol; 2 min then 15 min), followed by washing with DMF (3×4.0 mL/0.1 mmol resin). The reaction progress was monitored by Kaiser's tests(Kaiser et al., 1970) or by test cleavage and subsequent LC-MS analysis.

**General on-resin Alloc deprotection procedure**
A solution of borane dimethylamine complex (6 equiv.) and $Pd(Ph_3)_4$ (0.1 equiv.) in $CH_2Cl_2$ (2.0 mL/0.1 mmol resin) was added to the fritted syringe containing the resin-bound peptide and the reaction mixture was agitated for 15 min at ambient temperature. The procedure was repeated, and the resin was then washed with $CH_2Cl_2$ (3×4.0 mL), DMF (3×4.0 mL) and $CH_2Cl_2$ (3×4.0 mL).

**General on-resin Teoc deprotection procedure**
A solution of TBAF trihydrate (10 equiv.) in DMF (2.0 mL/0.1 mmol resin) was added to the fritted syringe containing the resin bound peptide and the reaction mixture was agitated for 1 h at 50 °C. The procedure was repeated, and the resin was then washed with DMF (3×4 mL) and $CH_2Cl_2$ (3×4 mL).

**General on-resin thiourea formation procedure**
A solution of the desired amine (2 equiv.) and *i*$Pr_2NEt$ (3 equiv.) in $CH_2Cl_2$ (3.0 mL/0.1 mmol) was added dropwise over 5 min to a solution of Bis(benzotriazol-1-yl)methanethione (2 equiv.) in $CH_2Cl_2$ (1.5 mL/0.1 mmol) at 0 °C. The reaction mixture was stirred for 10 min at 0 °C and concentrated under reduced pressure. The crude residue and *i*$Pr_2NEt$ (2 equiv.) were dissolved in DMF (2.0 mL/0.1 mmol resin) and then added to the fritted syringe containing the resin bound peptide. and the reaction mixture was agitated for 3–16 h. After washing with DMF (3×4.0 mL) and $CH_2Cl_2$ (3×4.0 mL).

**General procedure for global deprotection and cleavage from the resin**
Peptides were cleaved from the resin by TFA/$H_2O$/TIPS (95:2.5:2.5 (v/v/v), 4.0 mL/0.1 mmol; 2 h) and excess TFA removed under a stream of nitrogen unless described otherwise. The resulting crude was triturated with ice-cold diethyl ether and purified by preparative reverse-phase HPLC.

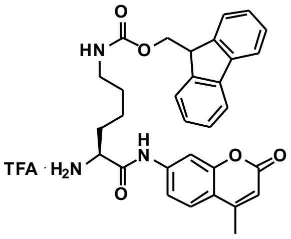

**S1**

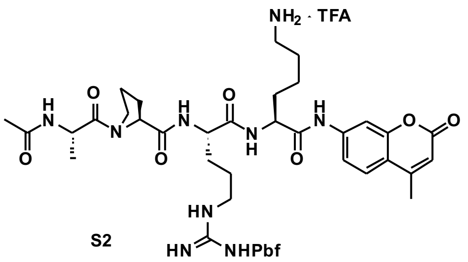

**S2**

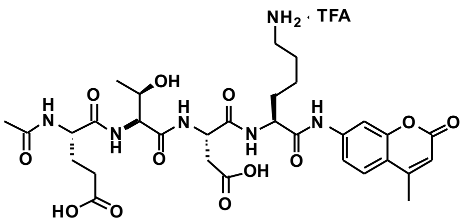

**S3**

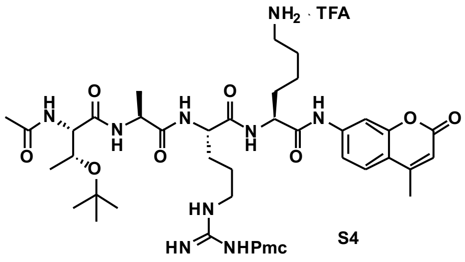

**S4**

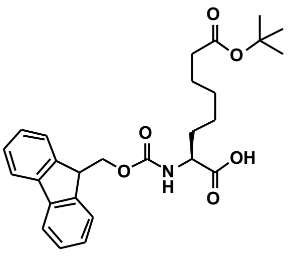

Fmoc-Asu(*t*Bu)-OH (**S5**)

*Compounds **S1** [14], **S2** [15], **S3** [2b], **S4** [16], and **S5** [17] were synthesized in previously.*

#### Synthesis of Fluorogenic Sirtuin Substrates

##### Ac-Gly-Gly-Val-OH (S6)

Ac-Gly-Gly-Val-resin (293 μmol) was synthesized on 2-CTC resin by standard SPPS procedures.

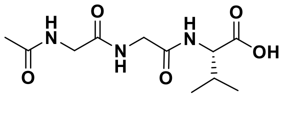

The peptide was cleaved from resin with $CH_2Cl_2$/**TFA** (4:1, v/v, 2×4 mL, 2×30 min) and solvent removed under a stream of nitrogen. Excess TFA was removed by co-evaporation with $CH_2Cl_2$–toluene (1:1, v/v, 15 mL) after which the crude peptide was dissolved in water (10 mL) and frozen to afford the titled compound (76 mg, 95%) as a colorless fluffy TFA-salt after lyophilization. $^1$H NMR (600 MHz, DMSO-$d_6$) δ 12.60 (br s, 1H, COOH), 8.14 (t, $J$ = 5.8 Hz, 1H, $NH_{Gly,1}$), 8.06 (t, $J$ = 5.8 Hz, 1H, $NH_{Gly,2}$), 7.89 (d, $J$ = 8.5 Hz, 1H, $NH_{Val}$), 4.14 (dd, $J$ = 8.5, 5.9 Hz, 1H, $H_{\alpha,Val}$), 3.82–3.73 (m, 2H, $H_{\alpha,Gly,2}$), 3.68 (d, $J$ = 5.8 Hz, 2H, $H_{\alpha,Gly,1}$), 2.10–2.00 (m, 1H, $H_{\beta,Val}$), 1.85 (s, 3H, $CH_{3,acetyl}$), 0.88 (d, $J$ = 6.8 Hz, 3H, $H_{\gamma,Val,1}$), 0.86 (d, $J$ = 6.8 Hz, 3H, $H_{\gamma,Val,2}$). $^{13}$C NMR (151 MHz, DMSO) δ 172.8 (COOH), 169.7 ($CO_{acetyl}$), 169.3 ($CO_{Gly,1}$), 168.9 ($CO_{Gly,2}$), 57.1 ($C_{\alpha,Val}$), 42.1 ($C_{\alpha,Gly,1}$), 41.7 ($C_{\alpha,Gly,2}$), 29.8 ($C_{\beta,Val}$), 22.5 ($CH_{3,acetyl}$), 19.1 ($C_{\gamma,Val,1}$), 17.9 ($C_{\gamma,Val,2}$). ESI-MS $m/z$ calcd for $C_{11}H_{20}N_3O_5^+$ $[M+H]^+$, 274.1; found 274.1. CAS RN: 97530-33-1.

##### Ac-Gly-Gly-Val-Lys-AMC (S7)

Compounds **S1** (52 mg, 0.084 mmol) and **S6** (23 mg, 0.084 mmol) were dissolved in anh. DMF

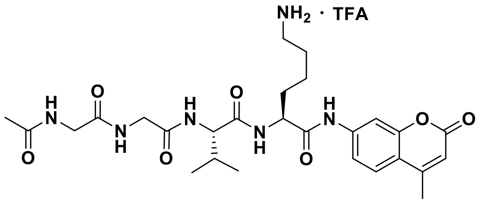

(2.0 mL) and cooled to 0 °C. HOAt (14 mg, 0.101 mmol), $i$Pr$_2$NEt (58 μL, 0.340 mmol) and PyBOP (53 mg, 0.101 mmol) were added to the reaction mixture, which was stirred for 10 min at 0 °C and then for 2 h going towards ambient temperature. The reaction mixture was concentrated under reduced pressure to afford a crude yellow oil, tentatively assigned as Ac-Gly-Gly-Val-Lys(Fmoc)-AMC (ESI-MS $m/z$ calcd for $C_{42}H_{49}N_6O_9^+$ $[M+H]^+$, 781.9; found 781.9). The crude was redissolved in DMF/**piperidine** (4:1, v/v, 2.0 mL) and stirred for 1 h at ambient temperature. Solvent was concentrated under reduced pressure, and preparative reverse-phase HPLC purification afforded the titled compound (40 mg, 72%) as a colorless fluffy material after lyophilization. $^1$H NMR (600 MHz, DMSO-$d_6$) δ 10.38 (s, 1H, $NH_{AMC}$), 8.24 (d, $J$ = 7.4 Hz, 1H, $NH_{\alpha,Lys}$), 8.17 (t, $J$ = 5.8 Hz, 1H, $NH_{Gly,1}$), 8.12 (t, $J$ = 5.8 Hz, 1H, $NH_{Gly,2}$), 7.84 (d, $J$ = 8.4 Hz, 1H, $NH_{Val}$), ) 7.78 (m, 1H, $H8_{AMC}$), 7.73 (d, $J$ = 8.7 Hz, 1H, $H5_{AMC}$), 7.69 (s, 3H, $NH_{\varepsilon,Lys}$), 7.50 (dd, $J$ = 8.7, 2.1 Hz, 1H, $H6_{AMC}$), 6.27 (d, $J$ = 1.2 Hz, 1H, $H3_{AMC}$), 4.40–4.33 (m, 1H, $H_{\alpha,Lys}$), 4.20 (dd, $J$ = 8.4, 6.6 Hz, 1H, $H_{\alpha,Val}$), 3.84–3.72 (m, 2H, $H_{\alpha,Gly,2}$), 3.68 (d, $J$ = 5.8 Hz, 2H, $H_{\alpha,Gly,1}$), 2.84–2.72 (m, 2H, $H_{\varepsilon,Lys}$), 2.40 (d, $J$ = 1.3 Hz, 3H, $CH_{3,AMC}$), 2.06–1.96 (m, 1H, $H_{\beta,Val}$), 1.86 (s, 3H, $CH_{3,acetyl}$), 1.80–1.72 (m, 1H, $H_{\beta,Lys,A}$), 1.71–1.62 (m, 1H, $H_{\beta,Lys,B}$), 1.59–1.49 (m, 2H, $H_{\delta,Lys}$), 1.46–1.29 (m, 2H, $H_{\gamma,Lys}$), 0.88 (d, $J$ = 6.8 Hz, 3H, $H_{\gamma,Val,1}$), 0.84 (d, $J$ = 6.8 Hz, 3H, $H_{\gamma,Val,2}$). $^{13}$C NMR (151 MHz, DMSO) δ 172.2 ($CO_{Val}$), 171.1 ($CO_{\alpha,Lys}$), 169.9 ($CO_{acetyl}$), 169.5 ($CO_{Gly,1}$), 169.0 ($CO_{Gly,2}$), 160.0 ($C2_{AMC}$), 158.0 (q, $J$ = 32.4 Hz, residual $CO_{TFA}$), 153.6 ($C8a_{AMC}$), 153.1 ($C4_{AMC}$), 142.1 ($C7_{AMC}$), 126.0 ($C5_{AMC}$), 115.2 ($C6_{AMC}$), 115.1 ($C4a_{AMC}$), 112.2 ($C3_{AMC}$), 105.7 ($C8_{AMC}$), 57.7 ($C_{\alpha,Val}$), 53.6 ($C_{\alpha,Lys}$), 42.2 ($C_{\alpha,Gly,1}$), 41.9 ($C_{\alpha,Gly,2}$), 38.9 ($C_{\varepsilon,Lys}$), 30.9 ($C_{\beta,Lys}$), 30.4 ($C_{\beta,Val}$), 26.6 ($C_{\delta,Lys}$), 22.5 ($CH_{3,acetyl}$), 22.4 ($C_{\gamma,Lys}$), 19.1 ($C_{\gamma,Val,1}$), 18.1 ($C_{\gamma,Val,2}$), 18.0 ($CH_{3,AMC}$). ESI-MS $m/z$ calcd for $C_{27}H_{39}N_6O_7^+$ $[M+H]^+$, 559.2; found 559.2. AMC=7-amino-4-methylcoumarin.

##### Ac-Gly-Gly-Val-Lys(dec)-AMC (GGVKac)

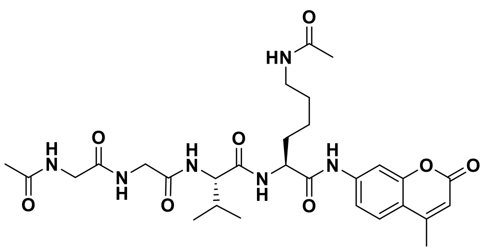

Compound **S7** (9 mg, 0.013 mmol) was dissolved in anh. $CH_2Cl_2$ (2.0 mL) and cooled to 0 °C. $i$Pr$_2$NEt (15 μL, 0.084 mmol) and acetic anhydride (4 μL, 0.042 mmol) were added to the reaction mixture and stirred for 10 min at 0 °C and then for 3 h going towards ambient temperature. Solvent was removed under reduced pressure, and preparative reverse-phase HPLC purification afforded the titled compound (4 mg, 50%) as a colorless fluffy material after lyophilization. $^{1}$H NMR (600 MHz, DMSO-$d_6$) δ 10.32 (s, 1H, $NH_{AMC}$), 8.21–8.09 (m, 3H, $NH_{\alpha,Lys}$, $NH_{Gly,1}$, $NH_{Gly,2}$), 7.83–7.78 (m, 2H, $NH_{Val}$, $NH_{\varepsilon,Lys}$), 7.77 (d, $J$ = 2.1 Hz, 1H, $H8_{AMC}$), 7.72 (d, $J$ = 8.7 Hz, 1H, $H5_{AMC}$), 7.50 (dd, $J$ = 8.7, 2.0 Hz, 1H, $H6_{AMC}$), 6.27 (d, $J$ = 1.3 Hz, 1H, $H3_{AMC}$), 4.35–4.30 (m, 1H, $H_{\alpha,Lys}$), 4.22 (dd, $J$ = 8.6, 6.5 Hz, 1H, $H_{\alpha,Val}$), 3.83–3.73 (m, 2H, $H_{\alpha,Gly,2}$), 3.68 (d, $J$ = 5.8 Hz, 2H, $H_{\alpha,Gly,1}$), 3.01 (q, $J$ = 6.5 Hz, 2H, $H_{\varepsilon,Lys}$), 2.40 (d, $J$ = 1.3 Hz, 3H, $CH_{3,AMC}$), 2.04–1.95 (m, 1H, $H_{\beta,Val}$), 1.85 (s, 3H, $CH_{3,acetyl}$), 1.77 (s, 3H, $COCH_{3,Lys}$), 1.75–1.70 (m, 1H, $H_{\beta,Lys,A}$), 1.69–1.62 (m, 1H, $H_{\beta,Lys,B}$), 1.46–1.18 (m, 4H, $H_{\gamma,Lys}$, $H_{\delta,Lys}$), 0.88 (d, $J$ = 6.8 Hz, 3H, $H_{\gamma,Val,1}$), 0.83 (d, $J$ = 6.8 Hz, 3H, $H_{\gamma,Val,2}$). $^{13}$C NMR (151 MHz, DMSO) δ 171.4 ($CO_{\alpha,Lys}$), 171.1 ($CO_{Val}$), 169.8 ($CO_{acetyl}$), 169.5 ($CO_{Gly,1}$), 168.98 ($CO_{Gly,2}$/$\underline{C}OCH_{3,Lys}$), 168.96 ($CO_{Gly,2}$/$CO_{acetyl,2}$), 160.0 ($C2_{AMC}$), 153.6 ($C8a_{AMC}$), 153.1 ($C4_{AMC}$), 142.2 ($C7_{AMC}$), 125.9 ($C5_{AMC}$), 115.2 ($C6_{AMC}$), 115.1 ($C4a_{AMC}$), 112.3 ($C3_{AMC}$), 105.6 ($C8_{AMC}$), 57.5 ($C_{\alpha,Val}$), 53.8 ($C_{\alpha,Lys}$), 42.1 ($C_{\alpha,Gly,1}$), 42.0 ($C_{\alpha,Gly,2}$), 38.2 ($C_{\varepsilon,Lys}$), 31.1 ($C_{\beta,Lys}$), 30.5 ($C_{\beta,Val}$), 28.8 ($C_{\delta,Lys}$), 23.0 ($C_{\gamma,Lys}$), 22.6 ($CH_{3,acetyl,2}$), 22.4 ($CH_{3,acetyl,1}$), 19.1 ($C_{\gamma,Val,1}$), 17.99 ($C_{\gamma,Val,2}$/$CH_{3,AMC}$), 17.96 ($C_{\gamma,Val,2}$/$CH_{3,AMC}$). Analytical HPLC (Method A) gradient 0–95% eluent IV in eluent III (35 min total runtime), $t_R$ 16.2 min (>98%, $UV_{230}$). HRMS calcd for $C_{29}H_{41}N_6O_8^+$ [M+H]$^+$, 601.2980; found 601.2989. AMC=7-amino-4-methylcoumarin.

**Ac-Gly-Gly-Val-Lys(dec)-AMC (GGVKdec)**

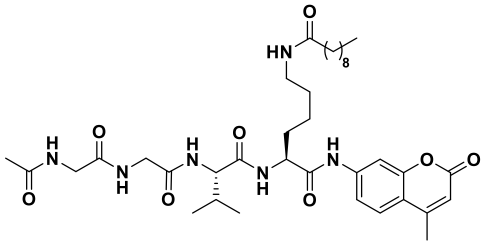

Compound **S7** (9 mg, 0.013 mmol) and capric acid (7 mg, 0.038 mmol) were dissolved in anh. DMF (2.0 mL) and cooled to 0 °C. $i$Pr$_2$NEt (13 µL, 0.076 mmol) and HATU (4 µL, 0.042 mmol) were added to the reaction mixture and was stirred for 10 min at 0 °C and then for 3 h going towards ambient temperature. Solvent was removed under reduced pressure, and preparative reverse-phase HPLC purification afforded the titled compound (4 mg, 92%) as a colorless fluffy material after lyophilization. $^{1}$H NMR (600 MHz, DMSO-$d_6$) δ 10.31 (s, 1H, $NH_{AMC}$), 8.22–8.08 (m, 3H, $NH_{\alpha,Lys}$, $NH_{Gly,1}$, $NH_{Gly,2}$), 7.81 (d, $J$ = 8.6 Hz, 1H, $NH_{Val}$), 7.77 (d, $J$ = 2.0 Hz, 1H, $H8_{AMC}$), 7.75–7.68 (m, 2H, $NH_{\varepsilon,Lys}$, $H5_{AMC}$), 7.50 (dd, $J$ = 8.7, 2.1 Hz, 1H, $H6_{AMC}$), 6.26 (d, $J$ = 1.3 Hz, 1H, $H3_{AMC}$), 4.37–4.29 (m, 1H, $H_{\alpha,Lys}$), 4.22 (dd, $J$ = 8.5, 6.5 Hz, 1H, $H_{\alpha,Val}$), 3.84–3.72 (m, 2H, $H_{\alpha,Gly,2}$), 3.67 (d, $J$ = 5.8 Hz, 2H, $H_{\alpha,Gly,1}$), 3.06–2.94 (m, 2H, $H_{\varepsilon,Lys}$), 2.40 (d, $J$ = 1.3 Hz, 3H, $CH_{3,AMC}$), 2.06–1.96 (m, 3H, $H_{\beta,Val}$, $COCH_2$), 1.85 (s, 3H, $CH_{3,acetyl}$), 1.78–1.61 (m, 2H, $H_{\beta,Lys}$), 1.46–1.06 (m, 18H, $H_{\gamma,Lys}$, $H_{\delta,Lys}$, $(C\underline{H}_2)_7CH_3$), 0.89–0.80 (m, 9H, $H_{\gamma,Val}$, $CH_2C\underline{H}_3$). $^{13}$C NMR (151 MHz, DMSO) δ 171.9 ($CONH_{\varepsilon,Lys}$), 171.3 ($CO_{\alpha,Lys}$), 171.0 ($CO_{Val}$), 169.8 ($CO_{acetyl,1}$), 169.5 ($CO_{Gly,1}$), 169.0 ($CO_{Gly,2}$), 160.0 ($C2_{AMC}$), 153.6 ($C8a_{AMC}$), 153.0 ($C4_{AMC}$), 142.2 ($C7_{AMC}$), 125.9 ($C5_{AMC}$), 115.2 ($C6_{AMC}$), 115.0 ($C4a_{AMC}$), 112.3 ($C3_{AMC}$), 105.6 ($C8_{AMC}$), 57.5 ($C_{\alpha,Val}$), 53.7 ($C_{\alpha,Lys}$), 42.1 ($C_{\alpha,Gly,1}$), 42.0 ($C_{\alpha,Gly,2}$), 38.1 ($C_{\varepsilon,Lys}$), 35.4 ($CO\underline{C}H_2$), 31.2 ($\underline{C}H_2CH_2CH_3$), 31.1 ($C_{\beta,Lys}$), 30.5 ($C_{\beta,Val}$), 28.9–28.6 (6C, $(\underline{C}H_2)_7CH_3$, $C_{\delta,Lys}$), 25.3 ($(\underline{C}H_2)_7CH_3$), 22.9 ($C_{\gamma,Lys}$), 22.4 ($CH_{3,acetyl,1}$), 22.1 ($\underline{C}H_2CH_3$), 19.1 ($C_{\gamma,Val,1}$), 18.0 ($C_{\gamma,Val,2}$/$CH_{3,AMC}$), 17.9 ($C_{\gamma,Val,2}$/$CH_{3,AMC}$), 13.9 ($CH_2\underline{C}H_3$). Analytical HPLC (Method A) gradient 0–95% eluent IV in eluent III (35 min total runtime), $t_R$ 22.8 min (>98%, $UV_{230}$). HRMS calcd for $C_{37}H_{57}N_6O_8^+$ [M+H]$^+$, 713.4232; found 713.4219. AMC=7-amino-4-methylcoumarin.

**Ac-Gly-Gly-Val-Lys(myr)-AMC (GGVKmyr)**

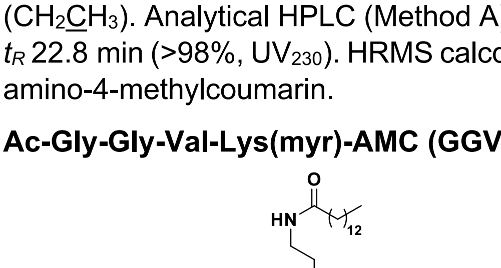

Compound **S7** (8 mg, 0.012 mmol) was dissolved in anh. $CH_2Cl_2$ (2.0 mL) and cooled to 0 °C. $i$Pr$_2$NEt (12 µL, 0.72 mmol) and myristoyl chloride (5 µL, 0.018 mmol) were added to the reaction mixture, which was stirred for 3 h going towards ambient temperature. Solvent was removed under reduced pressure, and preparative reverse-phase HPLC purification to afford the desired substrate (2 mg, 16%) as a colorless fluffy material after lyophilization.$^{1}$H NMR (600 MHz, DMSO-$d_6$) δ 10.32 (s, 1H, $NH_{AMC}$), 8.25–8.07 (m, 3H, $NH_{\alpha,Lys}$, $NH_{Gly,1}$, $NH_{Gly,2}$), 7.81 (d, $J$ = 8.5 Hz, 1H, $NH_{Val}$), 7.77 (d, $J$ = 2.0 Hz, 1H, $H8_{AMC}$), 7.75–7.64 (m, 2H, $NH_{\varepsilon,Lys}$, $H5_{AMC}$), 7.50 (dd, $J$ = 8.7, 2.1 Hz, 1H, $H6_{AMC}$), 6.26 (d, $J$ = 1.3 Hz,

1H, $H3_{AMC}$), 4.36–4.29 (m, 1H, $H_{\alpha,Lys}$), 4.22 (dd, $J$ = 8.5, 6.5 Hz, 1H, $H_{\alpha,Val}$), 3.84–3.73 (m, 2H, $H_{\alpha,Gly,2}$), 3.67 (d, $J$ = 5.8 Hz, 2H, $H_{\alpha,Gly,1}$), 3.06–2.97 (m, 2H, $H_{\varepsilon,Lys}$), 2.40 (d, $J$ = 1.3 Hz, 3H, $CH_{3,AMC}$), 2.06–1.94 (m, 3H, $H_{\beta,Val}$, $COCH_2$), 1.85 (s, 3H, $CH_{3,acetyl}$), 1.79–1.61 (m, 2H, $H_{\beta,Lys}$), 1.47–1.09 (m, 26H, $H_{\gamma,Lys}$, $H_{\delta,Lys}$, $(C\underline{H}_2)_{11}CH_3$), 0.91–0.79 (m, 9H, $H_{\gamma,Val}$, $CH_2C\underline{H}_3$). $^{13}$C NMR (151 MHz, DMSO) δ 171.9 ($CONH_{\varepsilon,Lys}$), 171.3 ($CO_{\alpha,Lys}$), 171.0 ($CO_{Val}$), 169.8 ($CO_{acetyl,1}$), 169.5 ($CO_{Gly,1}$), 169.0 ($CO_{Gly,2}$), 160.0 ($C2_{AMC}$), 153.6 ($C8a_{AMC}$), 153.0 ($C4_{AMC}$), 142.2 ($C7_{AMC}$), 125.9 ($C5_{AMC}$), 115.2 ($C6_{AMC}$), 115.0 ($C4a_{AMC}$), 112.3 ($C3_{AMC}$), 105.6 ($C8_{AMC}$), 57.6 ($C_{\alpha,Val}$), 53.8 ($C_{\alpha,Lys}$), 42.0 ($C_{\alpha,Gly,1}$), 42.0 ($C_{\alpha,Gly,2}$), 38.1 ($C_{\varepsilon,Lys}$), 35.4 ($CO\underline{C}H_2$), 31.3 ($\underline{C}H_2CH_2CH_3$), 31.1 ($C_{\beta,Lys}$), 30.5 ($C_{\beta,Val}$), 29.0–28.6 (10C, $C_{\delta,Lys}$, $(\underline{C}H_2)_{11}CH_3$,), 25.3 ($(\underline{C}H_2)_{11}CH_3$), 22.9 ($C_{\gamma,Lys}$), 22.4 ($CH_{3,acetyl,1}$), 22.1 ($\underline{C}H_2CH_3$), 19.1 ($C_{\gamma,Val,1}$), 18.0 ($C_{\gamma,Val,2}/CH_{3,AMC}$), 17.9 ($C_{\gamma,Val,2}/CH_{3,AMC}$), 13.9 ($CH_2\underline{C}H_3$). Analytical HPLC (Method A) gradient 0–95% eluent IV in eluent III (35 min total runtime), $t_R$ 26.6 min (>98%, $UV_{230}$). HRMS calcd for $C_{41}H_{65}N_6O_8^+$ $[M+H]^+$, 769.4858; found 769.4861 AMC=7-amino-4-methylcoumarin.

**Ac-Ala-Pro-Arg-Lys(dec)-AMC (APRKdec)**

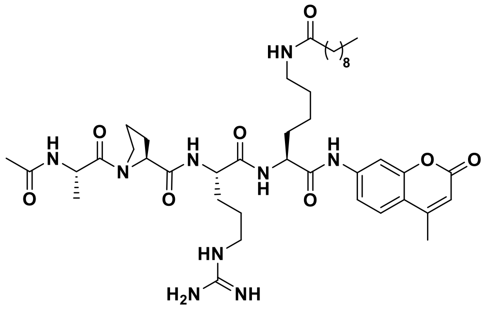

Capric acid (5 mg, 0.029 mmol) and **S2** (10 mg, 0.010 mmol) were dissolved in anh. DMF (2.0 mL) and cooled to 0 °C. *i*$Pr_2NEt$ (10 μL, 0.059 mmol) and HATU (11 mg, 0.029 mmol) were added to the reaction mixture and stirred for 10 min at 0 °C and then for 3 h going towards ambient temperature. Solvent was removed under reduced pressure to afford a yellow oil, tentatively assigned as Ac-Ala-Pro-Arg(Pbf)-Lys(dec)-AMC (ESI-MS *m/z* calcd for $C_{55}H_{82}N_9O_{11}S^+$ $[M+H]^+$, 1076.6; found 1076.6), which was used without further purification. The intermediate was resuspended in TFA–$H_2O$ (98:2, v/v, 1.0 mL) and stirred for 1 h at ambient temperature. Solvent was removed under a stream of nitrogen, and preparative reverse-phase HPLC purification afforded the titled compound (5 mg, 62%) as a colorless fluffy material after lyophilization. $^1$H NMR (600 MHz, DMSO-$d_6$) δ 10.33 (s, 1H, $NH_{AMC}$), 8.11 (d, $J$ = 7.3 Hz, 1H, $NH_{Ala}$), 8.04 (d, $J$ = 7.3 Hz, 1H, $NH_{\alpha,Lys}$), 8.02 (d, $J$ = 7.6 Hz, 1H, $NH_{\alpha,Arg}$), 7.80 (d, $J$ = 2.0 Hz, 1H, $H8_{AMC}$), 7.76–7.70 (m, 2H, $NH_{\varepsilon,Lys}$, $H5_{AMC}$), 7.56 (t, $J$ = 5.7 Hz, 1H, $NH_{\delta,Arg}$), 7.51 (dd, J = 8.7, 2.1 Hz, 1H, $H6_{AMC}$), 6.27 (d, $J$ = 1.4 Hz, 1H, $H3_{AMC}$), 4.52 (p, $J$ = 7.0 Hz, 1H, $H_{\alpha,Ala}$), 4.41–4.29 (m, 2H, $H_{\alpha,Pro}$, $H_{\alpha,Lys}$), 4.29–4.21 (m, 1H, $H_{\alpha,Arg}$), 3.67–3.60 (m, 1H, $H_{\delta,Pro,A}$), 3.59–3.52 (m, 2H, $H_{\delta,Pro,B}$), 3.11 (q, $J$ = 6.8 Hz, 2H, $H_{\delta,Arg}$), 3.07–2.95 (m, 2H, $H_{\varepsilon,Lys}$), 2.40 (d, $J$ = 1.3 Hz, 3H, $CH_{3,AMC}$), 2.19–1.06 (m, 36H, $H_{\beta,Pro}$, $H_{\gamma,Pro}$, $H_{\beta,Lys}$, $H_{\gamma,Lys}$, $H_{\delta,Lys}$, $H_{\beta,Arg}$, $H_{\gamma,Arg}$, $H_{\beta,Ala}$, $COCH_{3,acetyl}$, $(C\underline{H}_2)_8CH_3$), 0.84 (t, $J$ = 7.0 Hz, 3H, $CH_2C\underline{H}_3$). $^{13}$C NMR (151 MHz, DMSO) δ 172.0 ($\underline{C}ONH_{\varepsilon,Lys}$), 171.8 ($CO_{\alpha,Pro}$), 171.4 ($CO_{Arg}$), 171.3 ($CO_{\alpha,Lys}$), 171.2 ($CO_{Ala}$), 168.8 ($\underline{C}OCH_{3,acetyl}$), 160.0 ($C2_{AMC}$), 158.2 (q, $J$ = 31.2 Hz, residual $CO_{TFA}$), 156.7 (NHC(=NH)$NH_2$), 153.6 ($C8a_{AMC}$), 153.1 ($C4_{AMC}$), 142.1 ($C7_{AMC}$), 125.9 ($C5_{AMC}$), 117.1 (q, $J$ = 299.7 Hz, residual $CF_{3,TFA}$), 115.2 ($C6_{AMC}$), 115.1 ($C4a_{AMC}$), 112.3 ($C3_{AMC}$), 105.7 ($C8_{AMC}$), 59.6 ($C_{\alpha,Pro}$), 53.7 ($C_{\alpha,Lys}$), 52.2 ($C_{\alpha,Arg}$), 46.8 ($C_{\delta,Pro}$), 46.2 ($C_{\alpha,Ala}$), 40.4 ($C_{\delta,Arg}$), 38.1 ($C_{\varepsilon,Lys}$), 35.4 ($CO\underline{C}H_2$), 31.29 ($C_{\beta,Lys}$), 31.26 ($\underline{C}H_2CH_2CH_3$), 29.0–28.7 (8C, $C_{\beta,Pro}$, $C_{\beta,Arg}$, $C_{\delta,Lys}$, $(\underline{C}H_2)_7CH_3$), 25.3 ($(\underline{C}H_2)_7CH_3$), 24.9 ($C_{\gamma,Arg}$), 24.5 ($C_{\gamma,Pro}$), 22.8 ($C_{\gamma,Lys}$), 22.1 ($\underline{C}H_2CH_3$), 22.1 ($CO\underline{C}H_{3,acetyl}$), 18.0 ($CH_{3,AMC}$), 16.9 ($C_{\beta,Ala}$), 13.9 ($CH_2\underline{C}H_3$). Two sets of signals (approximately 6:1) were detectable due to rotamers. Only peaks for the major rotamer is given. Analytical HPLC (Method A) gradient 0–95% eluent IV in eluent III (35 min total runtime), $t_R$ 21.5 min (>98%, $UV_{230}$). HRMS calcd for $C_{42}H_{66}N_9O_8^+$ $[M+H]^+$, 824.5028; found 824.5010. AMC=7-amino-4-methylcoumarin.

**Ac-Ala-Pro-Arg-Lys(myr)-AMC (APRKmyr)**

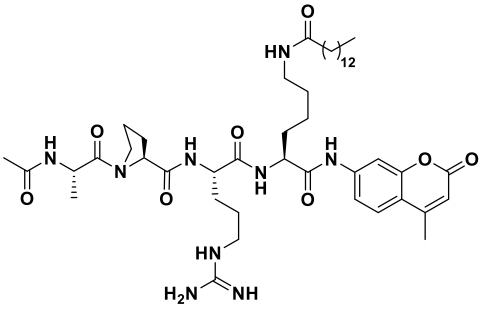

Compound **S2** (12 mg, 0.011 mmol) was dissolved in anh. $CH_2Cl_2$ (2.0 mL) and cooled to 0 °C. *i*$Pr_2NEt$ (12 μL, 0.067 mmol) and myristoyl chloride (9 μL, 0.034 mmol) were added to the reaction mixture and stirred for 10 min at 0°C and then for 3 h going towards ambient temperature. Solvent was removed under reduced pressure to afford a yellow oil, tentatively assigned as Ac-Ala-Pro-Arg(Pbf)-Lys(myr)-AMC (ESI-MS *m/z* calcd for $C_{59}H_{90}N_9O_{11}S^+$ $[M+H]^+$, 1132.6; found 1132.8), which was used without further purification. The intermediate was resuspended in TFA:$H_2O$ (98:2, v/v, 1.0 mL) and stirred for 1 h at ambient temperature. Solvent was removed under a stream of nitrogen, and preparative reverse-phase HPLC purification afforded the titled compound (2 mg, 25%) as a colorless fluffy material after lyophilization. $^1H$ NMR (600 MHz, DMSO-$d_6$) δ 10.33 (s, 1H, $NH_{AMC}$), 8.10 (d, *J* = 7.2 Hz, 1H, $NH_{Ala}$), 8.05 (d, *J* = 7.3 Hz, 1H, $NH_{\alpha,Lys}$), 8.02 (d, *J* = 7.5 Hz, 1H, $NH_{\alpha,Arg}$), 7.80 (d, *J* = 2.0 Hz, 1H, $H8_{AMC}$), 7.77–7.69 (m, 2H, $NH_{\varepsilon,Lys}$, $H5_{AMC}$), 7.55 (t, *J* = 5.7 Hz, 1H, $NH_{\delta,Arg}$), 7.50 (dd, J = 8.7, 2.1 Hz, 1H, $H6_{AMC}$), 6.27 (d, *J* = 1.4 Hz, 1H, $H3_{AMC}$), 4.52 (p, *J* = 7.0 Hz, 1H, $H_{\alpha,Ala}$), 4.40–4.29 (m, 2H, $H_{\alpha,Pro}$, $H_{\alpha,Lys}$), 4.28–4.21 (m, 1H, $H_{\alpha,Arg}$), 3.68–3.60 (m, 1H, $H_{\delta,Pro,A}$), 3.59–3.53 (m, 2H, $H_{\delta,Pro,B}$), 3.11 (q, *J* = 6.7 Hz, 2H, $H_{\delta,Arg}$), 3.06–2.95 (m, 2H, $H_{\varepsilon,Lys}$), 2.40 (d, *J* = 1.3 Hz, 3H, $CH_{3,AMC}$), 2.19–1.04 (m, 44H, $H_{\beta,Pro}$, $H_{\gamma,Pro}$, $H_{\beta,Lys}$, $H_{\gamma,Lys}$, $H_{\delta,Lys}$, $H_{\beta,Arg}$, $H_{\gamma,Arg}$, $H_{\beta,Ala}$, $COCH_{3,acetyl}$, $(C\underline{H_2})_{12}CH_3$), 0.84 (t, *J* = 6.9 Hz, 3H, $CH_2C\underline{H_3}$). $^{13}C$ NMR (151 MHz, DMSO) δ 172.0 ($\underline{C}ONH_{\varepsilon,Lys}$), 171.8 ($CO_{\alpha,Pro}$), 171.4 ($CO_{Arg}$), 171.3 ($CO_{\alpha,Lys}$), 171.2 ($CO_{Ala}$), 168.8 ($\underline{C}OCH_{3,acetyl}$), 160.0 ($C2_{AMC}$), 156.7 (NHC(=NH)$NH_2$), 153.6 ($C8a_{AMC}$), 153.1 ($C4_{AMC}$), 142.1 ($C7_{AMC}$), 125.9 ($C5_{AMC}$), 115.2 ($C6_{AMC}$), 115.1 ($C4a_{AMC}$), 112.3 ($C3_{AMC}$), 105.7 ($C8_{AMC}$), 59.6 ($C_{\alpha,Pro}$), 53.7 ($C_{\alpha,Lys}$), 52.2 ($C_{\alpha,Arg}$), 46.8 ($C_{\delta,Pro}$), 46.2 ($C_{\alpha,Ala}$), 40.4 ($C_{\delta,Arg}$), 38.1 ($C_{\varepsilon,Lys}$), 35.4 ($CO\underline{C}H_2$), 31.3 ($C_{\beta,Lys}$, $\underline{C}H_2CH_2CH_3$), 29.0–28.6 (12C, $C_{\beta,Pro}$, $C_{\beta,Arg}$, $C_{\delta,Lys}$, $(\underline{C}H_2)_{11}CH_3$), 25.3 ($(\underline{C}H_2)_{11}CH_3$), 24.9 ($C_{\gamma,Arg}$), 24.5 ($C_{\gamma,Pro}$), 22.8 ($C_{\gamma,Lys}$), 22.1 ($\underline{C}H_2CH_3$), 22.1 ($CO\underline{C}H_{3,acetyl}$), 18.0 ($CH_{3,AMC}$), 16.9 ($C_{\beta,Ala}$), 13.9 ($CH_2\underline{C}H_3$). Two sets of signals (approximately 5:1) were detectable due to rotamers. Only peaks for the major rotamer is given. Analytical HPLC (Method A) gradient 0–95% eluent IV in eluent III (35 min total runtime), $t_R$ 25.0 min (>98%, $UV_{230}$). HRMS calcd for $C_{46}H_{74}N_9O_8^+$ $[M+H]^+$, 880.5654; found 880.5640. AMC=7-amino-4-methylcoumarin.

**Ac-Glu-Thr-Asp-Lys(glu)-AMC (ETDKglu)**

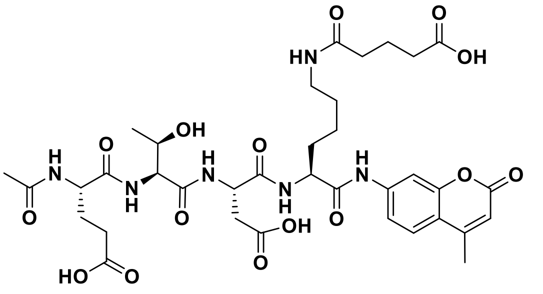

Compound **S3** (35 mg, 0.041 mmol) was suspended in $CH_2Cl_2$ (2.0 mL) with *i*$Pr_2NEt$ (22 μL, 0.122 mmol) and stirred at 0 °C prior to addition of glutaric anhydride (7 mg, 0.061 mmol). The reaction mixture was stirred for 5 min at 0 °C and then for 1 h at ambient temperature. The reaction mixture was diluted with $CH_2Cl_2$ (6 mL) and washed with aq. HCl (0.5 M, 2×5 mL), brine (10 mL), dried with $MgSO_4$ and concentrated under reduced pressure to afford a white solid, tentatively assigned as Ac-Glu(*t*Bu)-Thr(*t*Bu)-Asp(*t*Bu)-Lys(glu)-AMC (39 mg) which was dissolved in TFA:$CH_2Cl_2$:$H_2O$ (50:48:2, v/v/v, 2.0 mL) and stirred at ambient temperature for 1 h. The solution was evaporated to dryness, triturated in ice-cold ether and purified by preparative reverse-phase HPLC to afford the titled compound (21 mg, 62%) as a white fluffy material after lyophilization. $^1H$ NMR (600 MHz, DMSO-$d_6$) δ 12.16 (s, 3H, COOH), 10.21 (s, 1H, $NH_{AMC}$), 8.22 (d, *J* = 7.7 Hz, 1H, $NH_{Asp}$), 8.11 (d, *J* = 7.7 Hz, 1H, $NH_{Glu}$), 7.99 (d, *J* = 7.5 Hz, 1H, $NH_{\alpha,Lys}$), 7.79–7.74 (m, 2H, $H8_{AMC}$, $NH_{\varepsilon,Lys}$), 7.70 (d, *J* = 8.7 Hz, 1H, $H5_{AMC}$), 7.69 (d, *J* = 8.0 Hz, 1H, $NH_{Thr}$) 7.52 (dd, *J* = 8.7, 2.0 Hz, 1H, $H6_{AMC}$), 6.26 (d, *J* = 1.2 Hz, 1H, $H3_{AMC}$), 5.04 (s, 1H, $OH_{Thr}$), 4.61 (m, 1H, $H_{\alpha,Asp}$), 4.35–4.28 (m, 2H, $H_{\alpha,Lys}$, $H_{\alpha,Glu}$), 4.25 (dd, *J* = 8.0, 4.4 Hz, 1H, $H_{\alpha,Thr}$), 4.01 (td, *J* = 11.0, 6.0 Hz, 1H, $H_{\beta,Thr}$), 3.01 (m, 2H, $H_{\varepsilon Lys}$), 2.75 (dd, *J* = 16.7, 5.7 Hz, 1H, $H_{\beta,Asp,A}$), 2.59 (dd, *J* = 16.7, 7.4 Hz, 1H, $H_{\beta,Asp,B}$), 2.40 (d, *J* = 1.1 Hz, 3H, $CH_{3,AMC}$),

2.30–2.23 (m, 2H, $H_{\gamma,Glu}$), 2.17 (t, *J* = 7.4 Hz, 2H, C$\underline{H}_2$COOH), 2.06 (t, *J* = 7.5 Hz, 2H, NHCOC$\underline{H}_2$), 1.95–1.88 (m, 1H, $H_{\beta,Glu,A}$), 1.86 (s, 3H, $COCH_{3,acetyl}$), 1.78–1.59 (m, 5H, $H_{\beta,Glu,B}$, $H_{\beta,Lys}$, NHCOCH$_2$C$\underline{H}_2$), 1.43–1.22 (m, 4H, $H_{\gamma,Lys}$, $H_{\delta,Lys}$), 1.05 (d, *J* = 6.3 Hz, 3H, $H_{\gamma,Thr}$). $^{13}$C NMR (151 MHz, DMSO) δ 174.2 (COOH), 174.0 ($CO_{\delta,Glu}$), 171.9 ($CO_{\gamma,Asp}$), 171.6 (NH$\underline{C}$OCH$_2$/$CO_{\alpha,Glu}$), 171.4 ($CONH_{\varepsilon,Lys}$/$CO_{\alpha,Glu}$), 171.1 ($CO_{\alpha,Lys}$), 170.7 ($CO_{\alpha,Asp}$), 169.9 ($CO_{\alpha,Thr}$), 169.6 ($\underline{C}OCH_{3,acetyl}$), 160.0 ($C2_{AMC}$), 153.6 ($C8a_{AMC}$), 153.1 ($C4_{AMC}$), 142.0 ($C7_{AMC}$), 125.9 ($C5_{AMC}$), 115.4 ($C4a_{AMC}$), 115.2 ($C6_{AMC}$), 112.4 ($C3_{AMC}$), 105.8 ($C8_{AMC}$), 66.7 ($C_{\beta,Thr}$), 57.9 ($C_{\alpha,Thr}$), 53.8 ($C_{\alpha,Lys}$), 52.1 ($C_{\alpha,Glu}$), 49.6 ($C_{\alpha,Asp}$), 38.3 ($C_{\delta,Lys}$), 35.8 ($C_{\beta,Asp}$), 34.5 (NHCO$\underline{C}$H$_2$), 33.0 ($\underline{C}$H$_2$COOH), 31.3 ($C_{\beta,Lys}$), 30.2 ($C_{\gamma,Glu}$), 28.8 ($C_{\delta,Lys}$), 27.0 ($C_{\beta,Glu}$), 22.8 ($C_{\gamma,Lys}$), 22.4 (CO$\underline{C}H_{3,acetyl}$), 20.7 (COCH$_2\underline{C}$H$_2$), 19.2 ($C_{\gamma,Thr}$), 18.0 ($CH_{3,AMC}$). Analytical HPLC (Method A) gradient 0–95% eluent IV in eluent III (35 min total runtime), $t_R$ 15.1 min (>98%, UV$_{230}$). HRMS calcd for $C_{36}H_{49}N_6O_{15}^+$ [M+H]$^+$, 805.3250; found 805.3548. AMC=7-amino-4-methylcoumarin.

**Ac-Thr-Ala-Arg-Lys(hmg)-AMC (TARKhmg)**

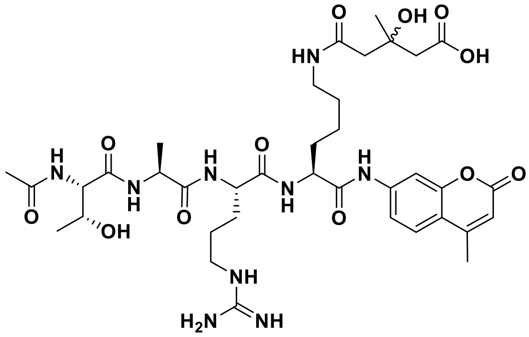

3-Hydroxy-3-methyl-glutaric acid (22 mg, 0.138 mmol) and HOBt (16 mg, 0.118 mmol) were dissolved in DMF (1.5 mL) and DIC (16 μL, 0.105 mmol) was added. The reaction was stirred for 20 min at ambient temperature, where crude **S6** (82 mg, 0.82 mmol) was added and the reaction mixture overnight. A solution of 3-hydroxy-3-methyl-glutaric acid (18 mg, 0.113 mmol) and DIC (17 mg, 0.132 mmol) in $CH_2Cl_2$ (0.5 mL) was added after pre-stirring for 2 h. After stirring for additional 45 min, *i*Pr$_2$NEt (43 μL, 0.248 mmol) was added and after 10 min, the reaction mixture was partitioned between EtOAc (20 mL) and water (10 mL). The aqueous phase was extracted with EtOAc (3×10 mL), and the combined organic layers evaporated to afford a white solid, tentatively assigned as Ac-Thr(*t*Bu)-Ala-Arg(Pmc)-Lys(hmg)-AMC (93 mg), which was dissolved in TFA:$CH_2Cl_2$:TIPS:H2O (95:2:2:1 (v/v/v/v), 3.26 mL) and stirred at ambient temperature for 90 min. Toluene (5 mL) was added, and the solution evaporated to dryness. The resulting residue was then co-evaporated with MeCN–toluene (50:50 (v/v), 3×3.0 mL) and purifed by preparative reverse-phase HPLC to afford the titled compound (41 mg, 62%) as a white fluffy material after lyophilization. $^1$H NMR (DMSO-*d*$_6$, 600 MHz) *δ* 10.41 (s, 1H, $NH_{AMC}$), 8.08 (d, *J* = 7.2 Hz, 1H, $NH_{\alpha,Lys}$), 8.01–7.94 (m, 3H, $NH_{\alpha,Arg}$,$NH_{Ala}$,$NH_{\varepsilon,Lys}$), 7.82 (d, *J* = 8.1 Hz, 1H, $NH_{Thr}$), 7.79 (d, *J* = 2.0 Hz, 1H, $H8_{AMC}$), 7.71 (d, *J* = 8.7 Hz, 1H, $H5_{AMC}$), 7.55 (t, *J* = 5.3 Hz, 1H, $NH_{\delta,Arg}$), 7.47 (dd, *J* = 8.7, 2.1 Hz, 1H, $H6_{AMC}$), 6.26 (d, *J* = 1.4 Hz, 1H, $H3_{AMC}$), 4.38–4.24 (m, 3H, $H_{\alpha,Ala}$,$H_{\alpha,Arg}$,$H_{\alpha,Lys}$), 4.18 (dd, *J* = 8.2, 4.4 Hz, 1H, $H_{\alpha,Thr}$), 4.01–3.92 (m, 1H, $H_{\beta,Thr}$), 3.10 (q, *J* = 6.6 Hz, 2H, $H_{\delta,Arg}$), 3.05 (q, *J* = 6.8 Hz, 2H, $H_{\varepsilon,Lys}$), 2.44–2.40 (m, 2H, C$\underline{H}_2$COOH), 2.39 (d, *J* = 1.2 Hz, 3H, $CH_{3,AMC}$), 2.37–2.33 (br s, 2H, NHCOC$\underline{H}_2$), 1.91 (s, 3H, $COCH_{3,acetyl}$), 1.80–1.61 (m, 3H, $H_{\beta,Arg,A}$,$H_{\beta,Lys}$), 1.61–1.47 (m, 3H, $H_{\beta,Arg,B}$,$H_{\gamma,Arg}$), 1.47–1.27 (m, 4H, $H_{\gamma,Lys}$,$H_{\delta,Lys}$), 1.24 (d, *J* = 7.1 Hz, 3H, $H_{\beta,Ala}$), 1.18 (s, 3H, C$\underline{H}_3$COH), 1.05 (d, *J* = 6.3 Hz, 3H, $H_{\gamma,Thr}$). $^{13}$C NMR (DMSO, 151 MHz) *δ* 172.4 (COOH/$CO_{Ala}$), 172.2 (COOH/$CO_{Ala}$), 171.4 ($CO_{Arg}$), 171.3 ($CO_{Lys}$), 171.0 ($CONH_{\varepsilon}$), 170.2 ($CO_{Thr}$), 169.9 ($CONH_{Thr}$), 160.0 ($C2_{AMC}$), 156.7 (NHC(=NH)NH$_2$), 153.6 ($C8a_{AMC}$), 153.1 ($C4_{AMC}$), 142.1 ($C7_{AMC}$), 125.9 ($C5_{AMC}$), 115.3 ($C6_{AMC}$), 115.1 ($C4a_{AMC}$), 112.3 ($C3_{AMC}$), 105.7 ($C8_{AMC}$), 69.5 (COH), 66.5 ($C_{\beta,Thr}$), 58.5 ($C_{\alpha,Thr}$), 53.7 ($C_{\alpha,Lys}$), 52.2 ($C_{\alpha,Arg}$), 48.4 ($C_{\alpha,Ala}$), 46.1 ($\underline{C}$H$_2$COOH), 45.8 (NHCO$\underline{C}$H$_2$), 40.5 ($C_{\delta,Arg}$), 38.3 ($C_{\varepsilon,Lys}$), 31.4 ($C_{\beta,Lys}$), 29.0 ($C_{\beta,Arg}$), 28.7 ($C_{\delta,Lys}$), 27.3 ($\underline{C}$H$_3$COH), 25.0 ($C_{\gamma,Arg}$), 22.8 ($C_{\gamma,Lys}$), 22.6 (*C*H$_3$CONH$_{thr}$), 19.7 ($C_{\gamma,thr}$), 18.0 ($C_{\beta,Ala}$/$CH_{3,AMC}$), 17.9 ($C_{\beta,Ala}$/$CH_{3,AMC}$). Analytical HPLC (Method A) gradient 0–95% eluent IV in eluent III (20 min total runtime), $t_R$ 10.7 min (>98%, UV$_{230}$). HRMS calcd for $C_{37}H_{56}N_9O_{12}^+$ [M+H]$^+$, 818.4043; found 818.4047. AMC=7-amino-4-methylcoumarin.

#### Synthesis of SIRT7 Inhibitor Series

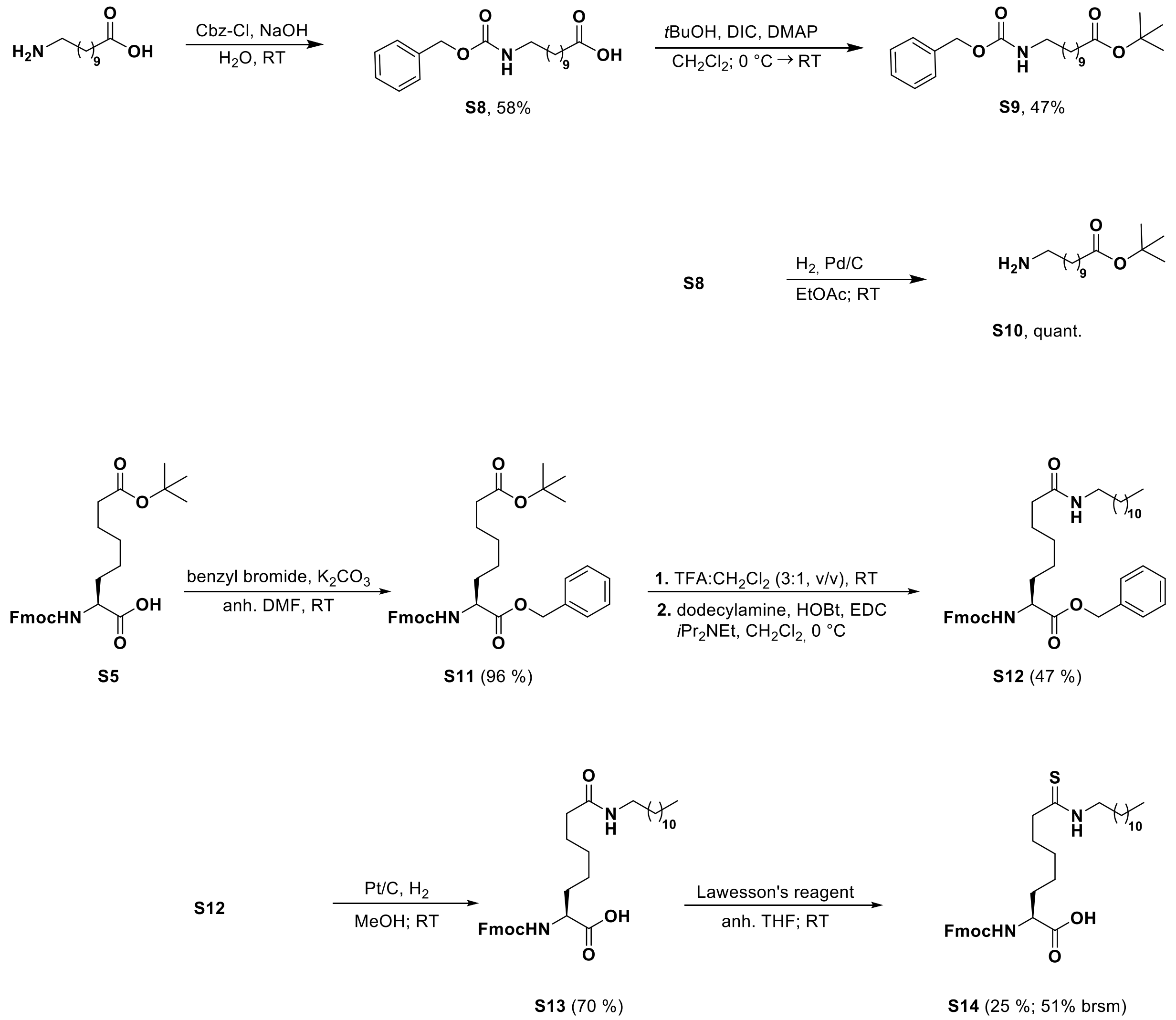

##### 11-(((benzyloxy)carbonyl)amino)undecanoic acid (S8)

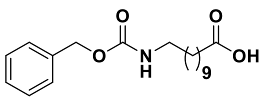

The synthetic procedure was adapted from a previous report.(D'Aléo et al., 2007) In short, benzyloxycarbonyl chloride (933 mg, 5.47 mmol) was added dropwise to a suspension of 11-aminoundecanoic acid (1.00 g, 4.97 mmol) in aq. NaOH (0.45 M, 15 mL) at 0 °C and the reaction mixture was stirred for 2 h at 0 °C. The suspension was acidified with aq. HCl (2 M) until pH < 2 and a white precipitate formed. The precipitate was filtered and washed with ice-cold aq. HCl (0.1 M, 25 mL) and purified by column chromatography (0→25% EtOAc and 0.25% AcOH in heptane) to afford the titled compound (962 mg, 58%) as a white solid. TLC (50% EtOAc and 0.25% AcOH in heptane): $R_f$ = 0.4. $^1$H NMR (400 MHz, DMSO-$d_6$) δ 11.93 (s, COOH), 7.41–7.27 (m, 5H, $H_{Ar}$), 7.19 (t, $J$ = 5.7 Hz, 1H, NH), 5.00 (s, 2H, $CH_{2,Cbz}$), 2.97 (q, $J$ = 6.6 Hz, 2H, C$\underline{H}_2$NHCbz), 2.18 (t, $J$ = 7.3 Hz, 2H, C$\underline{H}_2$COOH), 1.48 (p, $J$ = 7.0 Hz, 2H, C$\underline{H}_2$CH$_2$COOH), 1.38 (p, $J$ = 7.1 Hz, 2H, C$\underline{H}_2$CH$_2$NH), 1.31–1.17 (m, 12H, CH$_2$). $^{13}$C NMR (101 MHz, DMSO) δ 174.5 (COOH), 156.1 ($CO_{Cbz}$), 137.3 ($C1_{Ar,Cbz}$), 128.3 ($C3_{Ar,Cbz}$, $C5_{Ar,Cbz}$), 127.68 ($C4_{Ar,Cbz}$), 127.66 ($C2_{Ar,Cbz}$, $C6_{Ar,Cbz}$), 65.0 ($CH_{2,Cbz}$), 40.1 (NH$\underline{C}$H$_2$, overlap with solvent peak), 33.6 ($\underline{C}$H$_2$COOH), 29.4 ($\underline{C}$H$_2$CH$_2$NH), 28.9–28.5 (5C, CH$_2$), 26.2 ($\underline{C}$H$_2$), 24.5 ($\underline{C}$H$_2$CH$_2$COOH). ESI-MS $m/z$ calcd for $C_{19}H_{28}NO_4^+$ [M-H]$^-$, 334.2; found 334.4. CAS RN: 3422-91-1.

***tert*-butyl 11-(((benzyloxy)carbonyl)amino)undecanoate (S9)**

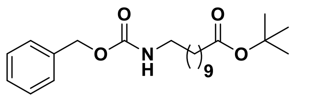

A solution of DIC (91 mg, 0.72 mmol) in anh. $CH_2Cl_2$ (0.5 mL) was added dropwise to a stirring solution of **S8** (200 mg, 0.60 mmol), *tert*-butanol (89 mg, 1.20 mmol) and 4-dimethylamino pyridine (DMAP; 11 mg, 0.10 mmol) in 10 mL anh. $CH_2Cl_2$ at 0 ˚C. The reaction mixture was stirred for 10 min at 0 °C and then overnight at ambient temperature. Additional DIC (91 mg, 0.72 mmol) was added and the reaction mixture was stirred additional 24 h. The reaction mixture was filtered, and the white precipitate washed with $CH_2Cl_2$ (10 mL). The filtrate was concentrated under reduced pressure and purified by column chromatography (0→10% EtOAc heptane) to afford the titled compound (111 mg, 47%) as a colorless oil. TLC (50% EtOAc in heptane): $R_f$ = 0.8. $^1$H NMR (600 MHz, $CDCl_3$) δ 7.38–7.28 (m, 5H, $H_{Ar}$), 5.09 (s, 2H, $CH_{2,Cbz}$), 4.71 (NH), 3.18 (q, *J* = 6.8 Hz, 2H, $NHC\underline{H}_2$), 2.19 (t, *J* = 7.5 Hz, 2H, $C\underline{H}_2CO$), 1.62–1.45 (m, 4H, $CH_2$, $C\underline{H}_2CH_2CO$), 1.44 (s, 9H, $C(CH_3)_3$), 1.32–1.22 (m, 12H, $CH_2$). $^{13}$C NMR (151 MHz, $CDCl_3$) δ 174.5 ($\underline{C}OOt$Bu), 156.5 ($CO_{Cbz}$), 136.8 ($C1_{Ar,Cbz}$), 128.7 ($C3_{Ar,Cbz}$, $C5_{Ar,Cbz}$), 128.3 ($C2_{Ar,Cbz}$, $C6_{Ar,Cbz}$), 128.2 ($C4_{Ar,Cbz}$), 80.0 ($\underline{C}(CH_3)_3$), 66.7 ($CH_{2,Cbz}$), 41.3 ($NH\underline{C}H_2$), 30.1 ($\underline{C}H_2CO$), 29.6 ($NHCH_2\underline{C}H_2$), 29.5–29.2 (5C, $CH_2$), 28.3 ($C(\underline{C}H_3)_3$), 26.9 ($CH_2$), 25.3 ($\underline{C}H_2CH_2COOH$). ESI-MS *m/z* calcd for $C_{18}H_{30}NO_2^+$ $[M\text{-}Boc]^+$, 292.2; found 292.2. CAS RN: 959006-04-3.

***tert*-butyl 11-aminoundecanoate (S10)**

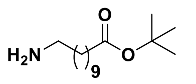

Compound **S9** (101 mg, 0.26 mmol) was added to a solution of Pd/C (10 mg, 10%, w/w) in EtOAc (5 mL) and stirred for 1 h under hydrogen atmosphere. Pd/C was filtered off and solvent removed under reduced pressure to afford the desired amine (62 mg, quant.) as a light-brown oil. $^1$H NMR (400 MHz, $CDCl_3$) δ 2.68 (t, *J* = 7.0 Hz, 2H, $NHC\underline{H}_2$), 2.19 (t, *J* = 7.5 Hz, 2H, $C\underline{H}_2CO$), 1.78 ($NH_2$), 1.62–1.12 (m, 25H, $(C\underline{H}_2)_8CH_2CO$, $C(CH_3)_3$). $^{13}$C NMR (101 MHz, $CDCl_3$) δ 173.5 ($\underline{C}OOt$Bu), 80.0 ($\underline{C}(CH_3)_3$). 42.3 ($NHCH_2$), 35.8 ($\underline{C}H_2CO$), 33.7 ($\underline{C}H_2CH_2NH$), 29.7–29.2 (XC, $CH_2$), 28.3 ($C(\underline{C}H_3)_3$), 27.0 ($CH_2$), 25.2 ($\underline{C}H_2CH_2COOH$). ESI-MS *m/z* calcd for $C_{15}H_{32}NO_2^+$ $[M+1]^+$, 258.2; found 258.3. CAS RN: 220851-29-6. Data is in agreement with literature.(Liu et al., 2002).

**Fmoc-Asu(*t*Bu)-OBn (S11)**

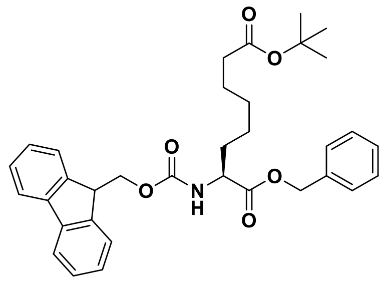

To a solution Fmoc-Asu(*t*Bu)-OH (**S5**; 524 mg, 1.12 mmol) in anh. DMF (8 mL) was added $K_2CO_3$ (309 mg, 2.24 mmol) and benzyl bromide (192 mg, 1.12 mmol) and the reaction mixture was stirred overnight at ambient temperature. Water (50 mL) was added and the slurry solution extracted with EtOAc:heptane (1:3, v/v, 3×30 mL). The combined organic layers were washed with sat. $NaHCO_3$ (2×25 mL) and brine (2×25 mL), dried with $MgSO_4$ and concentrated under reduced pressure to afford the crude product (600 mg, 96%) as a yellow-tainted oil. TLC (30% EtOAc in heptane): $R_f$ = 0.4. $^1$H NMR (600 MHz, $CDCl_3$) δ 7.76 (d, *J* = 7.6 Hz, 2H, $H4_{Fmoc}$, $H5_{Fmoc}$), 7.60 (d, *J* = 7.5 Hz, 2H, $H1_{Fmoc}$, $H8_{Fmoc}$), 7.43–7.27 (m, 9H, $H2_{Fmoc}$, $H3_{Fmoc}$, $H6_{Fmoc}$, $H7_{Fmoc}$, $H_{Bn,Ar}$), 5.24–5.13 (m, 2H, $C\underline{H}_{2,Bn}$), 4.47–4.35 (m, 3H, $H_{\alpha,Asu}$, $H10_{Fmoc}$), 4.22 (t, *J* = 7.2 Hz, 1H, $H9_{Fmoc}$), 2.17 (t, *J* = 7.5 Hz, 2H, $H_{\zeta,Asu}$), 1.89–1.81 (m, 1H, $H_{\beta,Asu,A}$), 1.89–1.81 (m, 1H, $H_{\beta,Asu,A}$), 1.72–1.64 (m, 1H, $H_{\beta,Asu,B}$), 1.57–1.48 (m, 2H, $H_{\varepsilon,Asu}$), 1.44 (s, 9H, $H_{tBu}$), 1.37–1.17 (m, 4H, $H_{\gamma,Asu}$, $H_{\delta,Asu}$). $^{13}$C NMR (151 MHz, $CDCl_3$) δ 173.2 ($CO_{\zeta,Asu}$), 172.5 ($CO_{Asu}$), 156.0 ($CO_{Fmoc}$), 144.1 ($C8a_{Fmoc}/C9a_{Fmoc}$), 143.9 ($C8a_{Fmoc}/C9a_{Fmoc}$), 141.5 ($C4a_{Fmoc}$, $C4b_{Fmoc}$), 135.5 ($C1_{Bn}$), 128.8 ($C3_{Bn}$, $C5_{Bn}$), 128.7 ($C4_{Bn}$), 128.5 ($C2_{Bn}$, $C6_{Bn}$), 127.9 ($C3_{Fmoc}$, $C6_{Fmoc}$), 127.2 ($C2_{Fmoc}$, $C7_{Fmoc}$), 125.2 ($C1_{Fmoc}$, $C8_{Fmoc}$), 120.13 ($C4_{Fmoc}/C5_{Fmoc}$), 120.12 ($C4_{Fmoc}/C5_{Fmoc}$), 80.2 ($C_{tBu}$), 67.3 ($\underline{C}H_{2,Bn}$), 67.2 ($C10_{Fmoc}$), 54.0 ($C_{\alpha,Asu}$), 47.3 ($C9_{Fmoc}$), 35.5 ($C_{\zeta,Asu}$), 32.7 ($C_{\beta,Asu}$), 28.8 ($C_{\delta,Asy}$), 28.3 ($\underline{C}H_{3,tBu}$), 25.0 ($C_{\gamma,Asu}/C_{\varepsilon,Asu}$), 24.9 ($C_{\gamma,Asu}/C_{\varepsilon,Asu}$). ESI-MS *m/z* calcd for $C_{34}H_{40}NO_6^+$ $[M+1]^+$, 558.3; found 558.3. CAS RN: 2231421-06-8.

**Benzyl (*S*)-2-((((9*H*-fluoren-9-yl)methoxy)carbonyl)amino)-8-(dodecylamino)-8-oxooctanoate (S12)**

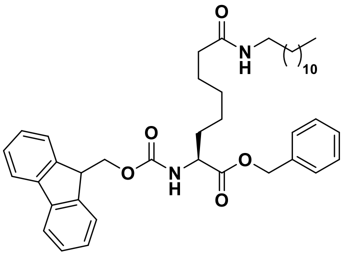

Compound **S11** (583 mg, 1.04 mmol) was dissolved in TFA:$CH_2Cl_2$ (3:1, v/v, 20 mL) and stirred at ambient temperature for 1 h. Solvent was removed under a stream of air and excess TFA was removed by co-evaporation with toluene:MeOH (1:1, v/v, 2×100 mL), providing crude Fmoc-Asu-Bn as a yellow oil (692 mg,) which was used without further purification. The crude oil was redissolved in $CH_2Cl_2$ (15 mL) at stirred at 0 °C. Dodecylamine (185 mg, 1.00 mmol), HOBt (135 mg, 1.00 mmol), *i*$Pr_2$NEt (523 μL, 3.00 mmol) and then EDC•HCl (288 mg, 1.50 mmol) were added to the reaction mixture and stirred at 0 °C for 10 min and then overnight at ambient temperature. The reaction mixture was concentrated under reduced pressure and redissolved in EtOAc (50 mL) and washed with aq. $K_2SO_4$ (5% v/w, 3×50 mL), sat. $NaHCO_3$ (3×50 mL) and brine (2×50 mL). The organic phase was dried over $Na_2SO_4$ and concentrated under pressure to afford the crude product, which was purified by vacuum liquid chromatography (0→40% EtOAC in heptane) to furnish the titled compound (326 mg, 47%) as an off-white solid. TLC (50% EtOAc in heptane): $R_f$ = 0.3. $^1$H NMR (600 MHz, DMSO-$d_6$) δ 7.89 (d, *J* = 7.6 Hz, 2H, $H4_{Fmoc}$, $H5_{Fmoc}$), 7.80 (d, *J* = 7.8 Hz, 1H, $NH_{\alpha,Asu}$), 7.73–7.66 (m, 3H, $H1_{Fmoc}$, $H8_{Fmoc}$, <u>NH</u>$CH_2$), 7.41 (t, *J* = 7.5 Hz, 2H, $H3_{Fmoc}$, $H6_{Fmoc}$), 7.36–7.26 (m, 7H, $H2_{Fmoc}$, $H7_{Fmoc}$, $H_{Bn,Ar}$), 5.16–5.09 (m, 2H, $CH_{2,Bn}$), 4.32–4.26 (m, 2H, $H10_{Fmoc}$), 4.21 (t, *J* = 7.1 Hz, 1H, $H9_{Fmoc}$), 4.05 (td, *J* = 9.3, 7.7m, 5.1 Hz, 1H, $H_{\alpha,Asu}$), 3.00 (q, *J* = 6.6 Hz, 2H, NHC<u>H</u>$_2$), 2.01 (t, *J* = 7.4 Hz, 2H, $H_{\zeta,Asu}$), 1.72–1.58 (m, 2H, $H_{\beta,Asy}$), 1.49–1.40 (m, 2H, $H_{\varepsilon,Asu}$), 1.39–1.08 (m, 24H, $H_{\gamma,Asu}$, $H_{\delta,Asu,}$ (C<u>H</u>$_2$)$_{10}$$CH_3$), 0.84 (t, *J* = 7.0 Hz, 3H, $CH_2$C<u>H</u>$_3$). $^{13}$C NMR (151 MHz, DMSO) δ 172.3 ($CO_{\alpha,Asu}$), 171.7 ($CO_{\zeta,Asu}$), 156.1 ($CO_{Fmoc}$), 143.8 ($C8a_{Fmoc}$/$C9a_{Fmoc}$), 143.7 ($C8a_{Fmoc}$/$C9a_{Fmoc}$), 140.7 ($C4a_{Fmoc}$, $C4b_{Fmoc}$), 136.0 ($C1_{Bn}$), 128.4 ($C3_{Bn}$, $C5_{Bn}$), 128.0 ($C4_{Bn}$), 127.7 ($C2_{Bn}$, $C6_{Bn}$), 127.6 ($C2_{Fmoc}$, $C7_{Fmoc}$), 127.0 ($C3_{Fmoc}$, $C6_{Fmoc}$), 125.2 ($C1_{Fmoc}$, $C8_{Fmoc}$), 120.1 ($C4_{Fmoc}$, $C5_{Fmoc}$), 65.8 (<u>C</u>$H_{2,Bn}$), 65.7 ($C10_{Fmoc}$), 53.9 ($C_{\alpha,Asu}$), 46.6 ($C9_{Fmoc}$), 38.3 (NH<u>C</u>$H_2$), 35.3 ($C_{\zeta,Asu}$), 31.3 (<u>C</u>$H_2CH_2CH_3$), 30.5 ($C_{\beta,Asu}$), 29.1–28.1 (8C, (<u>C</u>$H_2$)$_{10}$$CH_3$), 26.4 ($C_{\delta,Asu}$), 25.2 ($C_{\varepsilon,Asu}$), 25.1 ($C_{\gamma,Asu}$), 22.1 (<u>C</u>$H_2CH_3$), 13.9 ($CH_3$). ESI-MS *m/z* calcd for $C_{42}H_{57}N_2O_5^+$ $[M+1]^+$, 669.4; found 669.4.

**(*S*)-2-((((9*H*-fluoren-9-yl)methoxy)carbonyl)amino)-8-(dodecylamino)-8-oxooctanoic acid (Fmoc-Asu(myr)-OH, S13)**

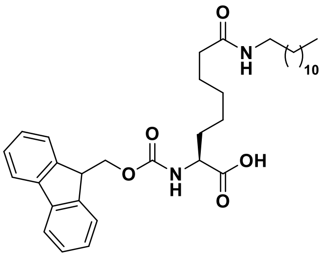

Compound **S12** (79 mg, 0.136 mmol) in MeOH (10 mL) was added to Pt/C (20 mg, 10% w/w) and stirred for 1 h under a hydrogen atmosphere. The reaction mixture was filtered through a pad of celite, concentrated under reduced pressure and purified by column chromatography (0→2.5% MeOH and 0.25% AcOH in $CH_2Cl_2$) to afford the titled compound (55 mg, 70%*) as a colorless solid. TLC (5% MeOH in 0.25% AcOH in $CH_2Cl_2$): $R_f$ = 0.4. $^1$H NMR (600 MHz, DMSO-$d_6$) δ 12.5 (br s, 1H, COOH), 7.89 (d, *J* = 7.6 Hz, 2H, $H4_{Fmoc}$, $H5_{Fmoc}$), 7.72 (dd, *J* = 7.8, 3.0 Hz, 2H, $H1_{Fmoc}$, $H8_{Fmoc}$), 7.69 (t, *J* = 5.8 Hz, 1H, <u>NH</u>$CH_2$), 7.61 (d, *J* = 8.1 Hz, 1H, $NH_{Asu}$), 7.41 (t, J = 7.3 Hz, 2H, $H3_{Fmoc}$, $H6_{Fmoc}$), 7.32 (tt, *J* = 7.4, 1.3 Hz, 2H, $H2_{Fmoc}$, $H7_{Fmoc}$), 4.35–4.18 (m, 3H, $H9_{Fmoc}$, $H10_{Fmoc}$), 3.97–3.85 (m, 1H, $H_{\alpha,Asu}$), 3.00 (q, *J* = 6.8 Hz, 2H, NHC<u>H</u>$_2$), 2.03 (t, *J* = 7.4 Hz, 2H, $H_{\zeta,Asu}$), 1.72–1.64 (m, 1H, $H_{\beta,Asu,A}$), 1.63–1.56 (m, 1H, $H_{\beta,Asu,B}$), 1.55–1.42 (m, 2H, $H_{\varepsilon,Asu}$), 1.39–1.17 (m, 24H, $H_{\gamma,Asu}$, $H_{\delta,Asu}$, (C<u>H</u>$_2$)$_{10}$$CH_3$), 0.84 (t, *J* = 6.9 Hz, 3H, $CH_3$). $^{13}$C NMR (151 MHz, DMSO) δ 174.0 (COOH), 171.8 ($CO_{\zeta,Asu}$), 156.1 ($CO_{Fmoc}$), 143.85 ($C8a_{Fmoc}$/$C9a_{Fmoc}$), 143.77 ($C8a_{Fmoc}$/$C9a_{Fmoc}$), 140.7 ($C4a_{Fmoc}$, $C4b_{Fmoc}$), 127.6 ($C2_{Fmoc}$, $C7_{Fmoc}$), 127.0 ($C3_{Fmoc}$, $C6_{Fmoc}$), 125.2 ($C1_{Fmoc}$, $C8_{Fmoc}$), 120.1 ($C4_{Fmoc}$, $C5_{Fmoc}$), 65.6 ($C10_{Fmoc}$), 53.7 ($C_{\alpha,Asu}$), 46.7 ($C9_{Fmoc}$), 38.3 ($C_{\zeta,Asu}$), 35.4 ($NHCH_2$), 31.3 (<u>C</u>$H_2CH_2CH_3$), 30.6 ($C_{\beta,Asu}$), 29.0–28.2 (8C, $C_{\delta,Asu}$, (<u>C</u>$H_2$)$_9$$CH_3$), 26.4 ($NHCH_2$<u>C</u>$H_2$), 25.3 ($C_{\gamma,Asu}$/$C_{\varepsilon,Asu}$), 25.2 ($C_{\gamma,Asu}$/$C_{\varepsilon,Asu}$), 22.1 (<u>C</u>$H_2CH_3$), 13.9 ($CH_3$). ESI-MS *m/z* calcd for $C_{35}H_{51}N_2O_5^+$ $[M+1]^+$, 579.4; found 579.4. *Partial Fmoc-deprotection observed upon hydrogenation.

**(*S*)-2-((((9*H*-fluoren-9-yl)methoxy)carbonyl)amino)-8-(dodecylamino)-8-thioxooctanoic acid (Fmoc-Asu(thiomyr)-OH, S14)**

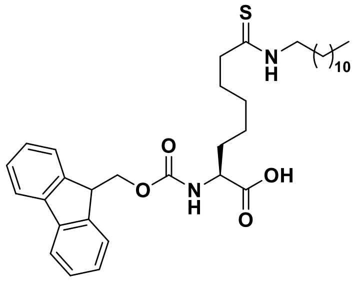

Lawesson's reagent(Lecher et al., 1956; Thomsen et al., 1984) (33 mg, 0.081 mmol) was added to **S13** (78 mg, 0.135 mmol) in anh. THF (1.0 mL) and stirred overnight at ambient temperature The reaction mixture was concentrated and purified by column chromatography (0→2.5% MeOH and 0.25% AcOH in $CH_2Cl_2$) to afford the titled compound (20 mg, 25% (51% brsm)) as a colorless solid. $^{1}$H NMR (600 MHz, DMSO-$d_6$) δ 9.86 (t, *J* = 5.3 Hz, 1H, NHCH$_2$), 7.89 (d, *J* = 7.5 Hz, 2H, H4$_{Fmoc}$, H5$_{Fmoc}$), 7.75–7.69 (m, 2H, H1$_{Fmoc}$, H8$_{Fmoc}$), 7.61 (d, *J* = 8.1 Hz, 1H, NH$_{Asu}$), 7.41 (t, *J* = 7.5 Hz, 2H, H3$_{Fmoc}$, H6$_{Fmoc}$), 7.33 (t, *J* = 7.4 Hz, 2H, H2$_{Fmoc}$, H7$_{Fmoc}$), 4.31–4.18 (m, 3H, H9$_{Fmoc}$, H10$_{Fmoc}$), 3.95–3.88 (m, 1H, H$_{\alpha,Asu}$), 3.50–3.43 (m, 2H, NHCH$_2$), 2.51–2.48 (m, 2H, H$_{\zeta,Asu}$, overlap with solvent peak), 1.73–1.43 (m, 6H, H$_{\beta,Asu}$, H$_{\varepsilon,Asu}$, (CH$_2$)$_{10}$CH$_3$), 1.38–1.19 (m, 22H, H$_{\gamma,Asu}$, H$_{\delta,Asu}$, (CH$_2$)$_{10}$CH$_3$), 0.84 (t, *J* = 7.0 Hz, 3H, CH$_3$). $^{13}$C NMR (151 MHz, DMSO) δ 203.4 (C=S), 174.0 (COOH), 156.1 (CO$_{Fmoc}$), 143.85 (C8a$_{Fmoc}$/C9a$_{Fmoc}$), 143.85 (C8a$_{Fmoc}$/C9a$_{Fmoc}$), 143.77 (C4a$_{Fmoc}$, C4b$_{Fmoc}$), 140.71 (C4a$_{Fmoc}$/C4b$_{Fmoc}$), 140.69 (C4a$_{Fmoc}$/C4b$_{Fmoc}$), 127.6 (C2$_{Fmoc}$, C7$_{Fmoc}$), 127.0 (C3$_{Fmoc}$, C6$_{Fmoc}$), 125.2 (C1$_{Fmoc}$, C8$_{Fmoc}$), 120.08 (C4$_{Fmoc}$/C5$_{Fmoc}$), 120.07 (C4$_{Fmoc}$/C5$_{Fmoc}$), 65.6 (C10$_{Fmoc}$), 53.7 (C$_{\alpha,Asu}$), 46.7 (C9$_{Fmoc}$), 45.04 (C$_{\zeta,Asu}$/NHCH$_2$), 44.96 (C$_{\zeta,Asu}$/NHCH$_2$), 31.3 (CH$_2$CH$_2$CH$_3$), 30.6 (C$_{\beta,Asu}$), 29.0–27.7 (9C, C$_{\delta,Asu}$, C$_{\varepsilon,Asu}$, (CH$_2$)$_9$CH$_3$), 26.4 (NHCH$_2$CH$_2$), 25.3 (C$_{\gamma,Asu}$), 22.1 (CH$_2$CH$_3$), 13.9 (CH$_3$). ESI-MS *m/z* calcd for $C_{35}H_{50}N_2O_4S^+$ [M+1]$^+$, 595.3; found 595.4.

**Benzyl ((6*S*,9*S*,12*S*)-1-amino-6-carbamoyl-1-imino-9-methyl-8,11-dioxo-18-thioxo-2,7,10,17,19-pentaazaheptacosan-12-yl)carbamate (4)**

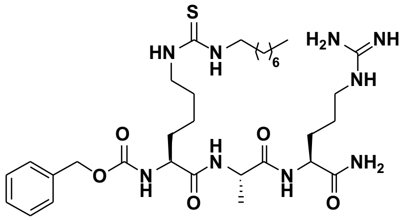

Starting from Cbz-Lys-Ala-Arg(Pbf)-resin (72 mg, estimated loading: 0.56 mmol/g) synthesized from Fmoc-Arg(Pbf)-OH, Fmoc-Ala-OH and Cbz-Lys(Fmoc)-OH by SPPS, the title compound was synthesized by on-resin thiourea formation as described in the general procedures using octylamine as the amine source. Global deprotection and cleavage from the resin, followed by preparative reverse-phase HPLC purification afforded the thiourea (7 mg, 25% based on resin loading) as a colorless fluffy material after lyophilization. $^{1}$H NMR (600 MHz, DMSO-$d_6$) δ 8.05 (d, *J* = 7.0 Hz, 1H, NH$_{Ala}$), 7.83 (d, *J* = 8.0 Hz, 1H, NH$_{\alpha,Arg}$), 7.55 (t, *J* = 5.8 Hz, 1H, NH$_{\delta,Arg}$), 7.42 (d, *J* = 8.0 Hz, 1H, NH$_{\alpha,Lys}$), 7.39–7.22 (m, 8H, NH$_{\varepsilon,Lys}$, NHCH$_2$, H$_{Ar,Cbz}$, CONH$_{2,A}$), 7.08 (s, 1H, CONH$_{2,B}$), 5.06–4.96 (m, 2H, CH$_{2,Cbz}$), 4.25 (p, *J* = 7.1 Hz, 1H, H$_{\alpha,Ala}$), 4.18 (td, *J* = 8.0, 5.5 Hz, 1H, H$_{\alpha,Arg}$), 4.01–3.92 (m, 1H, H$_{\alpha,Lys}$), 3.31 (br s, 2H, H$_{\varepsilon,Lys}$, overlap with residual water), 3.08 (q, *J* = 6.7 Hz, 2H, H$_{\delta,Arg}$), 1.74–1.59 (m, 2H, H$_{\beta,Arg,A}$, H$_{\beta,Lys,A,}$), 1.56–1.38 (m, 8H, H$_{\beta,Arg,B}$, H$_{\gamma,Arg}$, H$_{\beta,Lys,B}$, (CH$_2$)$_7$CH$_3$), 1.36–1.11 (m, 17H, H$_{\gamma,Lys,}$ H$_{\delta,Lys}$, H$_{\beta,Ala}$, (CH$_2$)$_7$CH$_3$), 0.85 (t, *J* = 6.9 Hz, 3H, CH$_2$CH$_3$). $^{13}$C NMR (151 MHz, DMSO) δ 173.1 (CO$_{Arg}$), 172.0 (CO$_{Ala}$), 171.9 (CO$_{Lys}$), 156.6 (NHC(=NH)NH$_2$), 156.0 (CO$_{Cbz}$), 137.0 (C1$_{Ar,Cbz}$), 128.3 (C3$_{Ar,Cbz,}$ C5$_{Ar,Cbz}$), 127.8 (C4$_{Ar,Cbz}$), 127.6 (C2$_{Ar,Cbz,}$ C6$_{Ar,Cbz}$), 65.4 (CH$_{2,Cbz}$), 54.6 (C$_{\alpha,Lys}$), 51.8 (C$_{\alpha,Arg}$), 48.3 (C$_{\alpha,Ala}$), 43.4 (C$_{\varepsilon,Lys}$), 40.4 (C$_{\delta,Arg}$), 31.6 (C$_{\beta,Lys}$), 31.2 (CH$_2$CH$_2$CH$_3$), 29.2 (C$_{\beta,Arg}$), 28.7–28.6 (5C, (CH$_2$)$_7$CH$_3$), 26.4 (C$_{\delta,Lys}$), 25.0 (C$_{\gamma,Arg}$), 23.0 (C$_{\gamma,Lys}$), 22.1 (CH$_2$CH$_3$), 18.0 (C$_{\beta,Ala}$), 13.9 (CH$_2$CH$_3$). The peak for C$_{\varepsilon,Lys}$ was broad and of low intensity and the peak for C=S was not visible in $^{13}$C NMR due to fast quadrupolar relaxation via the nearby $^{14}$N-nuclei. Analytical HPLC (Method A) gradient 0–95% eluent IV in eluent III (35 min total runtime), $t_R$ 21.5 min (>98%, UV$_{230}$). HRMS calcd for $C_{32}H_{56}N_9O_5S^+$ [M+H]$^+$, 678.4120; found 678.4126.

**(6*S*,9*S*,12*S*)-1-amino-12-(((benzyloxy)carbonyl)amino)-6-carbamoyl-1-imino-9-methyl-8,11-dioxo-18-thioxo-2,7,10,17,19-pentaazatriacontan-30-oic acid (5)**

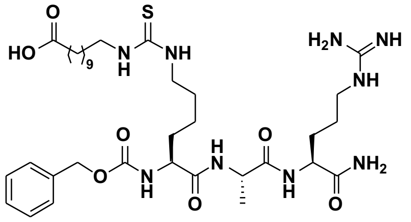

Starting from Cbz-Lys-Ala-Arg(Pbf)-resin (72 mg, estimated loading: 0.56 mmol/g) synthesized from Fmoc-Arg(Pbf)-OH, Fmoc-Ala-OH and Cbz-Lys(Fmoc)-OH by SPPS, the title compound was synthesized by on-resin thiourea formation as described in the general procedures using **S10** as the amine source. Global deprotection and cleavage from the resin, followed by preparative reverse-phase HPLC purification afforded the desired thiourea (5 mg, 15% based on resin loading) as a colorless fluffy material after lyophilization. $^{1}$H NMR (600 MHz, DMSO-$d_6$) δ 8.05 (d, $J$ = 7.0 Hz, 1H, $NH_{Ala}$), 7.83 (d, $J$ = 8.1 Hz, 1H, $NH_{\alpha,Arg}$), 7.55 (t, $J$ = 5.8 Hz, 1H, $NH_{\delta,Arg}$), 7.42 (d, $J$ = 8.0 Hz, 1H, $NH_{\alpha,Lys}$), 7.50–7.25 (m, 8H, $NH_{\varepsilon,Lys}$, $\underline{NH}CH_2$, $H_{Ar,Cbz}$, $CONH_{2,A}$), 7.08 (s, 1H, $CONH_{2,B}$), 5.10–4.95 (m, 2H, $CH_{2,Cbz}$), 4.25 (p, $J$ = 7.1 Hz, 1H, $H_{\alpha,Ala}$), 4.18 (td, $J$ = 8.0, 5.5 Hz, 1H, $H_{\alpha,Arg}$), 4.01–3.94 (m, 1H, $H_{\alpha,Lys}$), 3.31 (br s, 2H, $H_{\varepsilon,Lys}$, overlap with residual water), 3.13–3.03 (m, 2H, $H_{\delta,Arg}$), 2.18 (t, $J$ = 7.4 Hz, 2H, $C\underline{H_2}COOH$), 1.74–1.58 (m, 2H, $H_{\beta,Arg,A}$, $H_{\beta,Lys,A,}$), 1.55–1.05 (m, 29H, $H_{\beta,Arg,B}$, $H_{\gamma,Arg}$, $H_{\beta,Lys,B}$, $H_{\gamma,Lys,}$ $H_{\delta,Lys}$, $H_{\beta,Ala}$, $(C\underline{H_2})_9CH_2COOH$). $^{13}$C NMR (151 MHz, DMSO) δ 174.5 (COOH), 173.1 ($CO_{Arg}$), 172.0 ($CO_{Ala}$), 171.9 ($CO_{Lys}$), 158.3 (q, $J$ = 32.8 Hz, residual $CO_{TFA}$), 156.7 ($NHC(=NH)NH_2$), 156.0 ($CO_{Cbz}$), 137.0 ($C1_{Ar,Cbz}$), 128.3 ($C3_{Ar,Cbz,}$ $C5_{Ar,Cbz}$), 127.8 ($C4_{Ar,Cbz}$), 127.6 ($C2_{Ar,Cbz,}$ $C6_{Ar,Cbz}$), 116.7 (q, $J$ = 297.1 Hz, residual $CF_{3,TFA}$), 65.4 ($CH_{2,Cbz}$), 54.6 ($C_{\alpha,Lys}$), 51.8 ($C_{\alpha,Arg}$), 48.4 ($C_{\alpha,Ala}$), 43.4 ($C_{\varepsilon,Lys}$), 40.4 ($C_{\delta,Arg}$), 33.7 ($\underline{C}H_2COOH$), 31.6 ($C_{\beta,Lys}$), 29.2 ($C_{\beta,Arg}$), 29.0–28.5 (8C, $(\underline{C}H_2)_9CH_2COOH$), 26.4 ($C_{\delta,Lys}$), 25.0 ($C_{\gamma,Arg}$), 24.5 ($\underline{C}H_2CH_2COOH$), 23.0 ($C_{\gamma,Lys}$), 18.0 ($C_{\beta,Ala}$). The peak for $C_{\varepsilon,Lys}$ was broad and of low intensity and the peak for C=S was not visible in $^{13}$C NMR due to fast quadrupolar relaxation via the nearby $^{14}$N-nuclei. Analytical HPLC (Method A) gradient 0–95% eluent IV in eluent III (35 min total runtime), $t_R$ 19.2 min (>95%, $UV_{230}$). HRMS calcd for $C_{35}H_{60}N_9O_7S^+$ [M+H]$^+$, 750.4331; found 750.4321.

**Benzyl ((6*S*,9*S*,12*S*)-1-amino-6-carbamoyl-1-imino-9-methyl-8,11,18-trioxo-2,7,10,19-tetraazahentriacontan-12-yl)carbamate (6)**

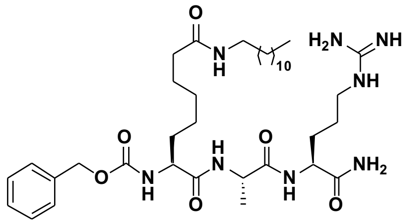

Starting from H-Asu(myr)-Ala-Arg(Pbf)-resin (115 mg, estimated loading: 0.35 mmol/g) synthesized from Fmoc-Arg(Pbf)-OH, Fmoc-D-Ala-OH and Fmoc-Asu(myr)-OH (**S13**) by SPPS, the title compound was synthesized by addition of benzyl chloroformate (17 μL, 0.12 mmol) and *i*$Pr_2NEt$ (42 μL, 0.24 mmol) in $CH_2Cl_2$ (2.0 mL) the resin followed by overnight agitation at ambient temperature. The resin was drained and washed with $CH_2Cl_2$ (3×4.0 mL) and DMF (3×4.0 mL) followed by global deprotection and cleavage from the resin. Preparative reverse-phase HPLC purification afforded the desired inverse amide (3 mg, 11% based on resin loading) as a colorless fluffy material after lyophilization. $^{1}$H NMR (600 MHz, DMSO-$d_6$) δ 8.04 (d, $J$ = 7.0 Hz, 1H, $NH_{Ala}$), 7.83 (d, $J$ = 8.1 Hz, 1H, $NH_{\alpha,Arg}$), 7.72 (t, $J$ = 5.6 Hz, 1H, $N\underline{H}CH_2$), 7.54 (t, $J$ = 5.8 Hz, 1H, $NH_{\delta,Arg}$), 7.41–7.27 (m, 7H, $NH_{Asu}$, $H_{Ar,Cbz}$, $CONH_{2,A}$), 7.08 (s, 1H, $CONH_{2,B}$), 5.08–4.96 (m, 2H, $CH_{2,Cbz}$), 4.24 (p, $J$ = 7.0 Hz, 1H, $H_{\alpha,Ala}$), 4.17 (td, $J$ = 8.1, 5.5 Hz, 1H, $H_{\alpha,Arg}$), 3.96 (td, $J$ = 8.6, 4.9 Hz, 1H, $H_{\alpha,Asu}$), 3.08 (q, $J$ = 6.7 Hz, 2H, $H_{\delta,Arg}$), 2.99 (q, $J$ = 6.6 Hz, 2H,$NHC\underline{H_2}$), 2.02 (t, $J$ = 7.5 Hz, 2H, $H_{\zeta,Asu}$), 1.71–1.64 (m, 1H, $H_{\beta,Arg,A}$), 1.64–1.15 (m, 34H, $H_{\beta,Arg,B}$, $H_{\gamma,Arg}$, $H_{\beta,Asu}$, $H_{\gamma,Asu}$, $H_{\delta,Asu}$, $H_{\varepsilon,Asu}$, $H_{\beta,Ala}$, $(C\underline{H_2})_{10}CH_3$), 0.85 (t, $J$ = 6.9 Hz, 3H, $CH_2C\underline{H_3}$). $^{13}$C NMR (151 MHz, DMSO) δ 173.2 ($CO_{Arg}$), 172.1–172.0 (3C, $CO_{Ala}$, $CO_{\alpha,Asu}$, $CO_{\zeta,Asu}$), 156.7 ($NHC(=NH)NH_2$), 156.1 ($CO_{Cbz}$), 137.0 ($C1_{Ar,Cbz}$), 128.4 ($C3_{Ar,Cbz,}$ $C5_{Ar,Cbz}$), 127.8 ($C4_{Ar,Cbz}$), 127.7 ($C2_{Ar,Cbz,}$ $C6_{Ar,Cbz}$), 65.4 ($CH_{2,Cbz}$), 54.6 ($C_{\alpha,Asu}$), 51.9 ($C_{\alpha,Arg}$), 48.4 ($C_{\alpha,Ala}$), 40.4 ($C_{\delta,Arg}$), 38.4 ($NHC\underline{H_2}$), 35.4 ($C_{\zeta,Asu}$), 31.7 ($C_{\beta,Asu}$), 31.3 ($\underline{C}H_2CH_2CH_3$), 29.2–28.4 (9C, $C_{\beta,Arg}$, $C_{\delta,Asu}$, $(C\underline{H_2})_9CH_3$), 26.4 ($NHCH_2\underline{C}H_2$), 25.3 ($C_{\gamma,Arg}$), 25.2 ($C_{\varepsilon,Asu}$), 25.0 ($C_{\gamma,Asu}$), 22.1 ($\underline{C}H_2CH_3$), 17.9 ($C_{\beta,Ala}$), 14.0 ($CH_2\underline{C}H_3$). Analytical HPLC (Method A) gradient 0–95% eluent IV in eluent III (35 min total runtime), $t_R$ 25.1 min (>98%, $UV_{210}$). ESI-MS calcd for $C_{37}H_{65}N_8O_6^+$ [M+H]$^+$, 717.5; found 717.5.

**Benzyl ((6*S*,9*S*,12*S*)-1-amino-6-carbamoyl-1-imino-9-methyl-8,11-dioxo-18-thioxo-2,7,10,19-tetraazahentriacontan-12-yl)carbamate (7)**

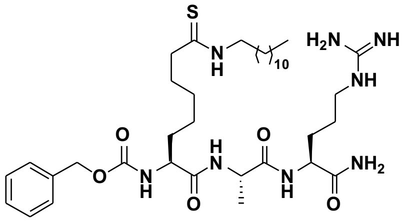

Starting from H-Asu(thiomyr)-Ala-Arg(Pbf)-resin (43 mg, estimated loading: 0.46 mmol/g) synthesized from Fmoc-Arg(Pbf)-OH, Fmoc-Ala-OH and Fmoc-Asu(thiomyr)-OH (**S14**) by SPPS. The title compound was synthesized by addition of benzyl chloroformate (17 µL, 0.12 mmol) and *i*$Pr_2NEt$ (42 µL, 0.24 mmol) in $CH_2Cl_2$ (2.0 mL) the resin followed by overnight agitation at ambient temperature. The resin was drained and washed with $CH_2Cl_2$ (3×4.0 mL) and DMF (3×4.0 mL) followed by global deprotection and cleavage from the resin. Preparative reverse-phase HPLC purification afforded the desired inverse thioamide (1 mg, 5% based on resin loading) as a colorless fluffy material after lyophilization. $^1$H NMR (600 MHz, DMSO-$d_6$) δ 9.85 (t, $J$ = 5.3 Hz, 1H, N$\underline{H}$$CH_2$), 8.01 (d, $J$ = 7.0 Hz, 1H, $NH_{Ala}$), 7.85 (d, $J$ = 8.1 Hz, 1H, $NH_{\alpha,Arg}$), 7.49 (t, $J$ = 5.8 Hz, 1H, $NH_{\delta,Arg}$), 7.41 (d, $J$ = 8.0 Hz, 1H, $NH_{Asu}$), 7.39–7.27 (m, 6H, $H_{Ar,Cbz}$, $CONH_{2,A}$), 7.07 (s, 1H, $CONH_{2,B}$), 5.02 ($m_{ABX}$, $J$ = 12.6 Hz, 2H, $CH_{2,Cbz}$), 4.25 (p, $J$ = 7.1 Hz, 1H, $H_{\alpha,Ala}$), 4.17 (td, $J$ = 8.0, 5.5 Hz, 1H, $H_{\alpha,Arg}$), 3.96 (td, $J$ = 8.9, 4.8 Hz, 1H, $H_{\alpha,Asu}$), 3.49–3.42 (m, 2H, NHC$\underline{H}_2$), 3.08 (q, $J$ = 6.7 Hz, 2H, $H_{\delta,Arg}$), 2.50–2.46 (m, 2H, $H_{\zeta,Asu}$, overlap with solvent peak), 1.72–1.09 (m, 35H, $H_{\beta,Arg}$, $H_{\gamma,Arg}$, $H_{\beta,Asu,}$, $H_{\gamma,Asu}$, $H_{\delta,Asu}$, $H_{\varepsilon,Asu}$, $H_{\beta,Ala}$, (C$\underline{H}_2$)$_{10}$$CH_3$), 0.85 (t, $J$ = 6.9 Hz, 3H, $CH_2$C$\underline{H}_3$). $^{13}$C NMR (151 MHz, DMSO) δ 203.5 (C=S), 173.1 ($CO_{Arg}$), 172.0 ($CO_{Ala}$), 171.8 ($CO_{Asu}$), 156.6 (NHC(=NH)$NH_2$), 156.0 ($CO_{Cbz}$), 137.0 ($C1_{Ar,Cbz}$), 128.3 ($C3_{Ar,Cbz,}$ $C5_{Ar,Cbz}$), 127.8 ($C4_{Ar,Cbz}$), 127.6 ($C2_{Ar,Cbz,}$ $C6_{Ar,Cbz}$), 65.4 ($CH_{2,Cbz}$), 54.6($C_{\alpha,Asu}$), 51.8 ($C_{\alpha,Arg}$), 48.3 ($C_{Ala}$), 45.1 (NHC$\underline{H}_2$), 45.0 ($C_{\zeta,Asu}$), 40.4 ($C_{\delta,Arg}$), 31.7 ($C_{\beta,Asu}$), 31.3 ($\underline{C}$$H_2CH_2CH_3$), 29.1–27.1 (10C, $C_{\varepsilon,Asu}$, $C_{\beta,Arg}$, ($\underline{C}$$H_2$)$_{10}$$CH_3$), 26.4 ($C_{\delta,Asu}$), 25.2 ($C_{\gamma,Arg}$), 25.0 ($C_{\gamma,Asu}$), 22.1 ($\underline{C}$$H_2CH_3$), 18.0 ($C_{\beta,Ala}$), 13.9 ($CH_2$$\underline{C}$$H_3$). Analytical HPLC (Method A) gradient 0–95% eluent IV in eluent III (35 min total runtime), $t_R$ 28.3 min (>98%, $UV_{230}$). ESI-MS calcd for $C_{37}H_{65}N_8O_5S^+$ $[M+H]^+$, 733.5; found 733.5.

**Benzyl ((2*R*,5*S*,8*S*)-1-amino-2,5-dimethyl-1,4,7-trioxo-14-thioxo-3,6,13,15-tetraazaheptacosan-8-yl)carbamate (8)**

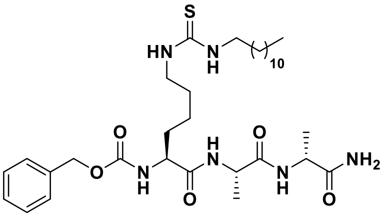

Starting from Cbz-Lys-Ala-D-Ala-resin (81 mg, estimated loading: 0.51 mmol/g) synthesized from Fmoc-D-Ala-OH, Fmoc-Ala-OH and Cbz-Lys(Fmoc)-OH by SPPS, the title compound was synthesized by on-resin thiourea formation as described in the general procedures using dodecylamine as the amine source. Global deprotection and cleavage from the resin, followed by preparative reverse-phase HPLC purification afforded the desired thiourea (1 mg, 4% based on resin loading) as a colorless fluffy material after lyophilization. $^1$H NMR (600 MHz, DMSO-$d_6$) δ 8.12 (d, $J$ = 6.9 Hz, 1H, $NH_{Ala}$), 7.94 (d, $J$ = 7.8 Hz, 1H, $NH_{\alpha,D\text{-}Ala}$), 7.46 (d, $J$ = 7.5 Hz, 1H, $NH_{\alpha,Lys}$), 7.41-7.19 (m, 8H, $NH_{\varepsilon,Lys}$, $\underline{NH}$$CH_2$, $H_{Ar,Cbz}$, $CONH_{2,A}$), 7.05 (d, $J$ = 2.2 Hz, 1H, $CONH_{2,B}$), 5.09–4.94 (m, 2H, $CH_{2,Cbz}$), 4.22 (p, $J$ = 7.0 Hz, 1H, $H_{\alpha,Ala}$), 4.16 (p, $J$ = 7.3 Hz, 1H, $H_{\alpha,D\text{-}Ala}$), 4.00–3.87 (m, 1H, $H_{\alpha,Lys}$*),* 3.31 (br s, 2H, $H_{\varepsilon,Lys}$, overlap with residual water), 1.63 (m, 1H, $H_{\beta,Lys,A}$), 1.55–1.09 (m, 33H, $H_{\beta,D\text{-}Ala}$, $H_{\beta,Ala}$, $H_{\beta,Lys,B}$, $H_{\gamma,Lys}$, $H_{\delta,Lys}$, (C$\underline{H}_2$)$_{11}$$CH_3$), 0.85 (t, $J$ = 6.9 Hz, 3H, $CH_2$C$\underline{H}_3$). $^{13}$C NMR (151 MHz, DMSO) δ 174.2 ($CO_{D\text{-}Ala}$), 172.0 ($CO_{Ala}$), 171.7 ($CO_{Lys}$), 156.2 ($CO_{Cbz}$), 136.9 ($C1_{Ar,Cbz}$), 128.4 ($C3_{Ar,Cbz}$, $C5_{Ar,Cbz}$), 127.8 ($C4_{Ar,Cbz}$), 127.7 ($C2_{Ar,Cbz}$, $C6_{Ar,Cbz}$), 65.5 ($CH_{2,Cbz}$), 54.7 ($C_{\alpha,Lys}$), 48.4 ($C_{\alpha,Ala}$), 48.0 ($C_{\alpha,D\text{-}Ala}$), 43.4 ($C_{\varepsilon,Lys}$), 31.5 ($C_{\beta,Lys}$), 31.3 ($\underline{C}$$H_2CH_2CH_3$), 29.0–28.7 (9C, ($\underline{C}$$H_2$)$_{11}$$CH_3$), 26.4 ($C_{\delta,Lys}$), 22.9 ($C_{\gamma,Lys}$), 22.1 ($\underline{C}$$H_2CH_3$), 18.1 ($C_{\beta,D\text{-}Ala}$) 17.7 ($C_{\beta,Ala}$), 14.0 ($CH_2$$\underline{C}$$H_3$). The peak for $C_{\varepsilon,Lys}$ was broad and of low intensity and the peak for C=S was not visible in $^{13}$C NMR due to fast quadrupolar relaxation via the nearby $^{14}$N-nuclei. Analytical HPLC (Method A) gradient 0–95% eluent IV in eluent III (35 min total runtime), $t_R$ 20.8 min (>98%, $UV_{230}$). HRMS calcd for $C_{33}H_{57}N_6O_5S^+$ $[M+H]^+$, 649.4106; found 649.4100.

**Benzyl ((6*R*,9*S*,12*S*)-1-amino-6-carbamoyl-1-imino-9-methyl-8,11-dioxo-18-thioxo-2,7,10,17,19-pentaazadocosan-12-yl)carbamate (9)**

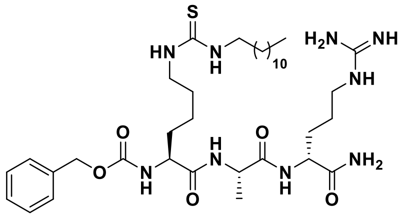

Starting from Cbz-Lys-Ala-D-Arg(Pbf)-resin (78 mg, estimated loading: 0.51 mmol/g) synthesized from Fmoc-D-Arg(Pbf)-OH, Fmoc-Ala-OH and Cbz-Lys(Fmoc)-OH by SPPS, the title compound was synthesized by on-resin thiourea formation as described in the general procedures using dodecylamine as the amine source. Global deprotection and cleavage from the resin, followed by preparative reverse-phase HPLC purification afforded the desired thiourea (3 mg, 9% based on resin loading) as a colorless fluffy material after lyophilization. $^{1}$H NMR (600 MHz, DMSO-$d_6$) δ 8.10 (d, *J* = 6.9 Hz, 1H, $NH_{Ala}$), 7.95 (d, *J* = 8.3 Hz, 1H, $NH_{\alpha,D\text{-}Arg}$), 7.58 (t, *J* = 5.8 Hz, 1H, $NH_{\delta,D\text{-}Arg}$), 7.47 (d, *J* = 7.4 Hz, 1H, $NH_{\alpha,Lys}$), 7.40–7.28 (m, 8H, $NH_{\varepsilon,Lys}$, $\underline{NH}CH_2$, $H_{Ar,Cbz}$, $CONH_{2,A}$), 7.13 (s, 1H, $CONH_{2,B}$), 5.09–4.95 (m, 2H, $CH_{2,Cbz}$), 4.27 (p, *J* = 7.0 Hz, 1H, $H_{\alpha,Ala}$), 4.16 (td, *J* = 8.6, 4.9 Hz, 1H, $H_{\alpha,D\text{-}Arg}$), 3.95 (td, *J* = 8.5, 4.9 Hz, 1H, $H_{\alpha,Lys}$), 3.31 (br s, 2H, $H_{\varepsilon,Lys}$, overlap with residual water), 3.08 (q, *J* = 6.6 Hz, 2H, $H_{\delta,D\text{-}Arg}$), 1.80–1.73 (m, 1H, $H_{\beta,D\text{-}Arg,A}$), 1.67–1.58 (m, 1H, $H_{\beta,Lys,A}$), 1.57–1.08 (m, 33H, $H_{\beta,D\text{-}Arg,B}$, $H_{\gamma,D\text{-}Arg}$, $H_{\beta,Lys,B}$, $H_{\gamma,Lys}$, $H_{\delta,Lys}$, $H_{\beta,Ala}$, $(C\underline{H}_2)_{11}CH_3$), 0.85 (t, *J* = 6.9 Hz, 3H, $CH_2C\underline{H}_3$). $^{13}$C NMR (151 MHz, DMSO) δ 173.3 ($CO_{D\text{-}Arg}$), 172.1 ($CO_{Ala}$), 172.0 ($CO_{Lys}$), 158.1 (q, *J* = 30.1 Hz, residual $CO_{TFA}$), 156.7 ($NHC(=NH)NH_2$), 156.2 ($CO_{Cbz}$), 136.9 ($C1_{Ar,Cbz}$), 128.3 ($C3_{Ar,Cbz}$, $C5_{Ar,Cbz}$), 127.8 ($C4_{Ar,Cbz}$), 127.7 ($C2_{Ar,Cbz}$, $C6_{Ar,Cbz}$), 65.5 ($CH_{2,Cbz}$), 54.8 ($C_{\alpha,Lys}$), 51.8 ($C_{\alpha,D\text{-}Arg}$), 48.4 ($C_{\alpha,Ala}$), 43.4 ($C_{\varepsilon,Lys}$), 40.3 ($C_{\delta,D\text{-}Arg}$), 31.5 ($C_{\beta,Lys}$), 31.3 ($\underline{C}H_2CH_2CH_3$), 29.0–28.7 (10C, $C_{\beta,D\text{-}Arg}$, $(\underline{C}H_2)_{11}CH_3$)), 26.4 ($C_{\delta,Lys}$), 25.1 ($C_{\gamma,D\text{-}Arg}$), 22.9 ($C_{\gamma,Lys}$), 22.1 ($\underline{C}H_2CH_3$), 17.9 ($C_{\beta,Ala}$), 13.9 ($CH_3$). The peak for $C_{\varepsilon,Lys}$ was broad and of low intensity and the peak for C=S was not visible in $^{13}$C NMR due to fast quadrupolar relaxation via the nearby $^{14}$N-nuclei. Analytical HPLC (Method A) gradient 0–95% eluent IV in eluent III (35 min total runtime), $t_R$ 24.6 min (>98%, $UV_{210}$). HRMS calcd for $C_{36}H_{64}N_9O_5S^+$ $[M+H]^+$, 734.4746; found 734.4740.

**(3*S*,6*S*,9*S*)-9-(((benzyloxy)carbonyl)amino)-3-carbamoyl-6-methyl-5,8-dioxo-15-thioxo-4,7,14,16-tetraazaoctacosanoic acid (10)**

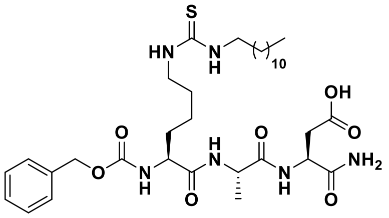

Starting from Cbz-Lys-Ala-Asp(*t*Bu)-resin (94 mg, estimated loading: 0.51 mmol/g) synthesized from Fmoc-Asp(*t*Bu)-OH, Fmoc-Ala-OH and Cbz-Lys(Fmoc)-OH by SPPS, the title compound was synthesized by the on-resin thiourea formation as described in the general procedures using dodecylamine as the amine source. Global deprotection and cleavage from the resin, followed by preparative reverse-phase HPLC purification afforded the desired thiourea (7 mg, 21% based on resin loading) as a colorless fluffy material after lyophilization. $^{1}$H NMR (600 MHz, DMSO-$d_6$) δ 12.28 (br s, 1H, COOH), 8.10 (d, *J* = 6.8 Hz, 1H, $NH_{Ala}$), 7.99 (d, *J* = 8.1 Hz, 1H, $NH_{Asp}$), 7.44 (d, *J* = 7.8 Hz, 1H, $NH_{\alpha,Lys}$), 7.41–7.23 (m, 8H, $NH_{\varepsilon,Lys}$, $\underline{NH}CH_2$, $H_{Ar,Cbz}$), 7.15–7.05 (m, 2H, $CONH_2$), 5.11–4.95 (m, 2H, $CH_{2,Cbz}$), 4.43 (q, *J* = 7.0 Hz, 1H, $H_{\alpha,Asp}$), 4.19 (p, *J* = 7.0 Hz, 1H, $H_{\alpha,Ala}$), 3.97 (td, *J* = 8.6, 4.7 Hz, 1H, $H_{\alpha,Lys}$), 3.32 (br s, 2H, $H_{\varepsilon,Lys}$, overlap with residual water), 2.64 (dd, *J* = 16.4, 5.9 Hz, 1H, $H_{\beta,Asp,A}$), 2.56–2.51 (m, 1H, $H_{\beta,Asp,B}$, overlap with solvent peak), 1.69–1.58 (m, 1H, $H_{\beta,Lys,A}$), 1.56–1.09 (m, 30H, $H_{\beta,Lys,B}$, $H_{\gamma,Lys}$, $H_{\delta,Lys}$, $H_{\beta,Ala}$, $(C\underline{H}_2)_{11}CH_3$), 0.85 (t, *J* = 6.8 Hz, 3H, $CH_2C\underline{H}_3$). $^{13}$C NMR (151 MHz, DMSO) δ 172.3 ($COOH_{,Asp}$), 172.1 ($CO_{Lys}$), 179.1 ($CO_{Ala}$, $CO_{Asp}$), 156.1 ($CO_{Cbz}$), 137.0 ($C1_{Ar,Cbz}$), 128.3 ($C3_{Ar,Cbz,}$ $C5_{Ar,Cbz}$), 127.8 ($C4_{Ar,Cbz}$), 127.7 ($C2_{Ar,Cbz,}$ $C6_{Ar,Cbz}$), 65.4 ($CH_{2,Cbz}$), 54.7 ($C_{\alpha,Lys}$), 49.3 ($C_{\alpha,Asp}$), 48.6 ($C_{\alpha,Ala}$), 43.4 ($C_{\varepsilon,Lys}$), 36.0 ($C_{\beta,Lys}$), 31.4 ($C_{\beta,Lys}$), 31.3 ($\underline{C}H_2CH_2CH_3$), 29.0–28.7 (9C, $(\underline{C}H_2)_{11}CH_3$), 26.4 ($C_{\delta,Lys}$), 23.0 ($C_{\gamma,Lys}$), 22.1 ($\underline{C}H_2CH_3$), 17.8 ($C_{\beta,Ala}$), 13.9 ($CH_2\underline{C}H_3$). The peak for $C_{\varepsilon,Lys}$ was broad and of low intensity and the peak for C=S was not visible in $^{13}$C NMR due to fast quadrupolar relaxation via the nearby $^{14}$N-nuclei. Analytical HPLC (Method A) gradient 0–95% eluent IV in eluent III (35 min total runtime), $t_R$ 24.9 min (>98%, $UV_{230}$). HRMS calcd for $C_{34}H_{56}N_6NaO_7S^+$ $[M+Na]^+$, 715.3823; found 715.3810.

**(4*S*,7*S*,10*S*)-10-(((benzyloxy)carbonyl)amino)-4-carbamoyl-7-methyl-6,9-dioxo-16-thioxo-5,8,15,17-tetraazanonacosanoic acid (11)**

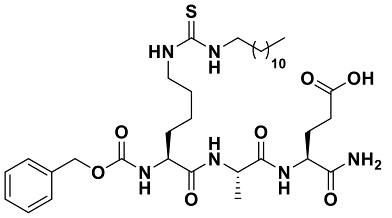

Starting from Cbz-Lys-Ala-Glu(*t*Bu)-resin (78 mg, estimated loading: 0.51 mmol/g) synthesized from Fmoc-Glu(*t*Bu)-OH, Fmoc-Ala-OH and Cbz-Lys(Fmoc)-OH by SPPS, the title compound was synthesized by on-resin thiourea formation as described in the general procedures using dodecylamine as the amine source. Global deprotection and cleavage from the resin, followed by preparative reverse-phase HPLC purification afforded the desired thiourea (1 mg, 5% based on resin loading) as a colorless fluffy material after lyophilization. $^{1}$H NMR (600 MHz, DMSO-$d_6$) δ 12.08 (br s, 1H, COOH), 8.06 (d, *J* = 7.0 Hz, 1H, $NH_{Ala}$), 7.81 (d, *J* = 8.0 Hz, 1H, $NH_{Glu}$), 7.44 (d, *J* = 7.8 Hz, 1H, $NH_{\alpha,Lys}$), 7.39–7.23 (m, 8H, $NH_{\varepsilon,Lys}$, $\underline{NH}CH_2$, $H_{Ar,Cbz}$, $CONH_{2,A}$), 7.07 (s, 1H, $CONH_{2,B}$), 5.09–4.95 (m, 2H, $CH_{2,Cbz}$), 4.24 (p, *J* = 7.1 Hz, 1H, $H_{\alpha,Ala}$), 4.16 (td, *J* = 8.2, 5.1 Hz, 1H, $H_{\alpha,Glu}$), 3.96 (ddd, *J* = 9.5, 7.8, 4.8 Hz, 1H, $H_{\alpha,Lys}$), 3.32 (br s, 2H, $H_{\varepsilon,Lys}$, overlap with residual water), 2.26–2.14 (m, 2H, $H_{\gamma,Glu}$), 1.95–1.85 (m, 1H, $H_{\beta,Glu,A}$), 1.80–1.69 (m, 1H, $H_{\beta,Glu,B}$), 1.68–1.58 (m, 1H, $H_{\beta,Lys,A}$), 1.55–1.09 (m, 30H, $H_{\beta,Lys,B}$, $H_{\gamma,Lys}$, $H_{\delta,Lys}$, $H_{\beta,Ala}$, $(C\underline{H}_2)_{11}CH_3$), 0.85 (t, *J* = 7.0 Hz, 3H, $CH_2C\underline{H}_3$). $^{13}$C NMR (151 MHz, DMSO) δ 174.0 (COOH), 172.9 ($CO_{Glu}$), 172.0 ($CO_{Ala}$), 171.9 ($CO_{Lys}$), 156.1 ($CO_{Cbz}$), 137.0 ($C1_{Ar,Cbz}$), 128.3 ($C3_{Ar,Cbz}$, $C5_{Ar,Cbz}$), 127.7 ($C4_{Ar,Cbz}$), 127.6 ($C2_{Ar,Cbz}$, $C6_{Ar,Cbz}$), 65.4 ($CH_{2,Cbz}$), 54.7 ($C_{\alpha,Lys}$), 51.7 ($C_{\alpha,Glu}$), 48.4 ($C_{\alpha,Ala}$), 43.4 ($C_{\varepsilon,Lys}$), 31.5 ($C_{\beta,Lys}$), 31.3 ($\underline{C}H_2CH_2CH_3$), 30.2 ($C_{\gamma,Glu}$), 29.0–28.7 (9C, $(\underline{C}H_2)_{11}CH_3$), 27.4 ($C_{\beta,Glu}$), 26.4 ($C_{\delta,Lys}$), 23.0 ($C_{\gamma,Lys}$), 22.1 ($\underline{C}H_2CH_3$), 17.8 ($C_{\beta,Ala}$), 13.9 ($CH_2\underline{C}H_3$). The peak for $C_{\varepsilon,Lys}$ was broad and of low intensity and the peak for C=S was not visible in $^{13}$C NMR due to fast quadrupolar relaxation via the nearby $^{14}$N-nuclei. Analytical HPLC (Method A) gradient 0–95% eluent IV in eluent III (35 min total runtime), $t_R$ 25.6 min (>98%, $UV_{230}$). HRMS calcd for $C_{35}H_{59}N_6O_7S^+$ $[M+H]^+$, 707.4160; found 707.4152.

**(4*R*,7*S*,10*S*)-10-(((benzyloxy)carbonyl)amino)-4-carbamoyl-7-methyl-6,9-dioxo-16-thioxo-5,8,15,17-tetraazanonacosanoic acid (12)**

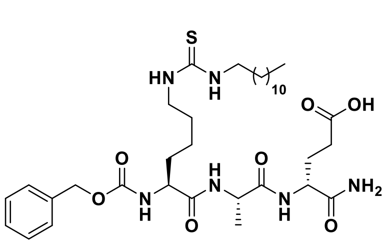

Starting from Cbz-Lys-Ala-D-Glu(*t*Bu)-resin (78 mg, estimated loading: 0.51 mmol/g) synthesized from Fmoc-D-Glu(*t*Bu)-OH, Fmoc-Ala-OH and Cbz-Lys(Fmoc)-OH by SPPS, the title compound was synthesized by the on-resin thiourea formation as described in the general procedures using dodecylamine as the amine source. Global deprotection and cleavage from the resin, followed by preparative reverse-phase HPLC purification afforded the desired thiourea (9 mg, 33% based on resin loading) as a colorless fluffy material after lyophilization. $^{1}$H NMR (600 MHz, DMSO-$d_6$) δ 12.07 (s, 1H, COOH), 8.10 (d, *J* = 6.8 Hz, 1H, $NH_{Ala}$), 7.95 (d, *J* = 8.3 Hz, 1H, $NH_{D\text{-}Glu}$), 7.46 (d, *J* = 7.6 Hz, 1H, $NH_{\alpha,Lys}$), 7.42–7.18 (m, 8H, $NH_{\varepsilon,Lys}$, $\underline{NH}CH_2$, $H_{Ar,Cbz}$, $CONH_{2,A}$), 7.11 (s, 1H, $CONH_{2,B}$), 5.11–4.94 (m, 2H, $CH_{2,Cbz}$), 4.24 (p, *J* = 7.0 Hz, 1H, $H_{\alpha,Ala}$), 4.17 (td, *J* = 8.7, 4.8 Hz, 1H, $H_{\alpha,D\text{-}Glu}$), 3.95 (td, *J* = 8.4, 5.0 Hz, 1H, $H_{\alpha,Lys}$), 3.32 (br s, 2H, $H_{\varepsilon,Lys}$, overlap with residual water), 2.20 (t, *J* = 7.8, 2H, $H_{\gamma,D\text{-}Glu}$), 2.02–1.93 (m, 1H, $H_{\beta,D\text{-}Glu,A}$), 1.76–1.66 (m, 1H, $H_{\beta,D\text{-}Glu,B}$), 1.62 (ddt, *J* = 15.0, 10.5, 5.2 Hz, 1H, $H_{\beta,Lys,A}$), 1.55–1.09 (m, 30H, $H_{\beta,Lys,B}$, $H_{\gamma,Lys}$, $H_{\delta,Lys}$, $H_{\beta,Ala}$, $(C\underline{H}_2)_{11}CH_3$), 0.85 (t, *J* = 7.0 Hz, 3H, $CH_2C\underline{H}_3$). $^{13}$C NMR (151 MHz, DMSO) δ 173.8 (COOH), 173.1 ($CO_{D\text{-}Glu}$), 172.1 ($CO_{Lys}$), 172.0 ($CO_{D\text{-}Glu}$), 156.1 ($CO_{Cbz}$), 136.9 ($C1_{Ar,Cbz}$), 128.3 ($C3_{Ar,Cbz}$,$C5_{Ar,Cbz}$), 127.8 ($C4_{Ar,Cbz}$), 127.7 ($C2_{Ar,Cbz}$, $C6_{Ar,Cbz}$), 65.5 ($CH_{2,Cbz}$), 54.7 ($C_{\alpha,Lys}$), 51.6 ($C_{\alpha,D\text{-}Glu}$), 48.5 ($C_{\alpha,Ala}$), 43.3 ($C_{\varepsilon,Lys}$), 31.5 ($C_{\beta,Lys}$), 31.3 ($\underline{C}H_2CH_2CH_3$), 30.1 ($C_{\gamma,D\text{-}Glu}$), 29.0–28.7 (9C, $(\underline{C}H_2)_{11}CH_3$), 27.1 ($C_{\beta,D\text{-}Glu}$), 26.4 ($C_{\delta,Lys}$), 22.9 ($C_{\gamma,Lys}$), 22.1 ($\underline{C}H_2CH_3$), 17.8 ($C_{\beta,Ala}$), 13.9 ($CH_2\underline{C}H_3$). The peak for $C_{\varepsilon,Lys}$ was broad and of low intensity and the peak for C=S was not visible in $^{13}$C NMR due to fast quadrupolar relaxation via the nearby $^{14}$N-nuclei. Analytical HPLC (Method A) gradient 0–95% eluent IV in eluent III (35 min total runtime), $t_R$ 24.9 min (>98%, $UV_{230}$). HRMS calcd for $C_{35}H_{59}N_6O_7S^+$ $[M+H]^+$, 707.4160; found 707.4152.

**Methyl (4*S*,7*S*,10*S*)-10-(((benzyloxy)carbonyl)amino)-4-carbamoyl-7-methyl-6,9-dioxo-16-thioxo-5,8,15,17-tetraazanonacosanoate (13)**

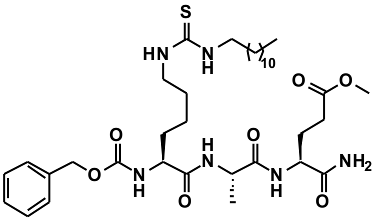

Starting from Cbz-Lys-Ala-Glu(OMe)-resin (87 mg, estimated loading: 0.51 mmol/g) synthesized from Fmoc-Glu(OMe)-OH, Fmoc-Ala-OH and Cbz-Lys(Fmoc)-OH by SPPS, the title compound was synthesized by the on-resin thiourea formation as described in the general procedures using dodecylamine as the amine source. Global deprotection and cleavage from resin, followed by preparative reverse-phase HPLC purification afforded the desired thiourea (9 mg, 27% based on resin loading) as a colorless fluffy material after lyophilization. $^{1}$H NMR (600 MHz, DMSO-$d_6$) δ 8.07 (d, $J$ = 7.1 Hz, 1H, $NH_{Ala}$), 7.78 (d, $J$ = 8.1 Hz, 1H, $NH_{Glu(OMe)}$), 7.43 (d, $J$ = 7.8 Hz, 1H, $NH_{\alpha,Lys}$), 7.41–7.19 (m, 8H, $NH_{\varepsilon,Lys}$, $\underline{NH}CH_2$, $H_{Ar,Cbz}$, $CONH_{2,A}$), 7.09 (s, 1H, $CONH_{2,B}$), 5.08–4.93 (m, 2H, $CH_{2,Cbz}$), 4.24 (p, $J$ = 7.1 Hz, 1H, $H_{\alpha,Ala}$), 4.16 (td, $J$ = 8.3, 5.1 Hz, 1H, $H_{\alpha,Glu(OMe)}$), 4.00–3.93 (m, 1H, $H_{\alpha,Lys}$), 3.57 (s, 3H, $OCH_3$), 3.32 (br s, 2H, $H_{\varepsilon,Lys}$, overlap with residual water), 2.29 (t, $J$ = 8.0 Hz, 2H, $H_{\gamma,Glu(OMe)}$), 2.00–1.89 (m, 1H, $H_{\beta,Glu(OMe),A}$), 1.82–1.68 (m, 1H, $H_{\beta,Glu(OMe),B}$), 1.68–1.58 (m, 1H, $H_{\beta,Lys,A}$), 1.56–1.06 (m, 30H, $H_{\beta,Lys,B}$, $H_{\gamma,Lys}$, $H_{\delta,Lys}$, $H_{\beta,Ala}$, $(C\underline{H_2})_{11}CH_3$), 0.85 (t, $J$ = 6.9 Hz, 3H, $CH_2C\underline{H_3}$). $^{13}$C NMR (151 MHz, DMSO) δ 172.8 ($CO_{\alpha,Glu(OMe)}$, $CO_{\delta,Glu(OMe)}$), 172.0 ($CO_{Ala}$), 171.9 ($CO_{Lys}$), 156.1 ($CO_{Cbz}$), 137.0 ($C1_{Ar,Cbz}$), 128.3 ($C3_{Ar,Cbz,}$ $C5_{Ar,Cbz}$), 127.7 ($C4_{Ar,Cbz}$), 127.6 ($C2_{Ar,Cbz,}$ $C6_{Ar,Cbz}$), 65.4 ($CH_{2,Cbz}$), 54.7 ($C_{\alpha,Lys}$), 51.5 ($C_{\alpha,Glu(OMe)}$), 51.3 ($OCH_3$), 48.3 ($C_{\alpha,Ala}$), 43.3 ($C_{\varepsilon,Lys}$), 31.5 ($C_{\beta,Lys}$), 31.3 ($\underline{C}H_2CH_2CH_3$), 29.7 ($C_{\gamma,Glu(OMe)}$), 29.0–28.7 (9C, $(\underline{C}H_2)_{11}CH_3$), 27.3 ($C_{\beta,Glu(OMe)}$), 26.4 ($C_{\delta,Lys}$), 23.0 ($C_{\gamma,Lys}$), 22.1 ($\underline{C}H_2CH_3$), 17.7 ($C_{\beta,Ala}$), 13.9 ($CH_2\underline{C}H_3$). The peak for $C_{\varepsilon,Lys}$ was broad and of low intensity and the peak for C=S was not visible in $^{13}$C NMR due to fast quadrupolar relaxation via the nearby $^{14}$N-nuclei. Analytical HPLC gradient 0–95% eluent IV in eluent III (35 min total runtime), $t_R$ 26.1 min (>99%, $UV_{230}$). HRMS calcd for $C_{36}H_{60}N_6NaO_7S^+$ $[M+H]^+$, 743.4136; found 743.4127.

**(4*S*,10*S*)-10-(((benzyloxy)carbonyl)amino)-4-carbamoyl-6,9-dioxo-16-thioxo-5,8,15,17-tetraazanonacosanoic acid (14)**

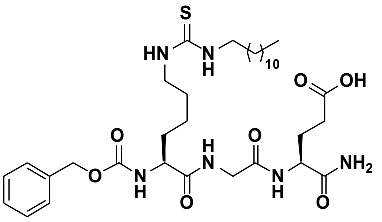

Starting from Cbz-Lys-Gly-Glu(*t*Bu)-resin (79 mg, estimated loading: 0.51 mmol/g) synthesized from Fmoc-Glu(*t*Bu)-OH, Fmoc-Gly-OH and Cbz-Lys(Fmoc)-OH by SPPS, the title compound was synthesized by the on-resin thiourea formation as described in the general procedures using dodecylamine as the amine source. Global deprotection and cleavage from the resin, followed by preparative reverse-phase HPLC purification afforded the desired thiourea (8 mg, 28% based on resin loading) as a colorless fluffy material after lyophilization. $^{1}$H NMR (600 MHz, DMSO-$d_6$) δ 12.08 (s, 1H, COOH), 8.19 (t, $J$ = 5.7 Hz, 1H, $NH_{Gly}$), 7.79 (d, $J$ = 8.1 Hz, 1H, $NH_{Glu}$), 7.51 (d, $J$ = 7.4 Hz, 1H, $NH_{\alpha,Lys}$), 7.39–7.23 (m, 8H, $NH_{\varepsilon,Lys}$, $\underline{NH}CH_2$, $H_{Ar,Cbz}$, $CONH_{2,A}$), 7.09 (s, 1H, $CONH_{2,B}$), 5.07–4.96 (m, 2H, $CH_{2,Cbz}$), 4.19 (td, $J$ = 8.5, 5.1 Hz, 1H, $H_{\alpha,Glu}$), 3.94 (ddd, $J$ = 9.1, 7.4, 5.1 Hz, 1H, $H_{\alpha,Lys}$), 3.79–3.67 (m, 2H, $H_{\alpha,Gly}$), 3.32 (br s, 2H, $H_{\varepsilon,Lys}$, overlap with residual water), 2.25–2.18 (m, 2H, $H_{\gamma,Glu}$), 1.99–1.89 (m, 1H, $H_{\beta,Glu,A}$), 1.78–1.69 (m, 1H, $H_{\beta,Glu,B}$), 1.69–1.60 (m, 1H, $H_{\beta,Lys,A}$), 1.58–1.10 (m, 27H, $H_{\beta,Lys,B}$, $H_{\gamma,Lys}$, $H_{\delta,Lys}$ $(C\underline{H_2})_{11}CH_3$), 0.85 (t, $J$ = 6.9 Hz, 3H, $CH_3$). $^{13}$C NMR (151 MHz, DMSO) δ 173.9 (COOH), 173.0 ($CO_{Glu}$), 172.4 ($CO_{Lys}$), 168.7 ($CO_{Gly}$), 156.1 ($CO_{Cbz}$), 136.9 ($C1_{Ar,Cbz}$), 128.3 ($C3_{Ar,Cbz}$, $C5_{Ar,Cbz}$), 127.8 ($C4_{Ar,Cbz}$), 127.7 ($C2_{Ar,Cbz}$, $C6_{Ar,Cbz}$), 65.5 ($CH_{2,Cbz}$), 54.9 ($C_{\alpha,Lys}$), 51.6 ($C_{\alpha,Glu}$), 43.4 ($C_{\varepsilon,Lys}$), 42.1 ($C_{\alpha,Gly}$), 31.32 ($C_{\beta,Lys}$), 31.28 ($\underline{C}H_2CH_2CH_3$), 30.1 ($C_{\gamma,Glu}$), 29.0–28.5 (9C, $(\underline{C}H_2)_{11}CH_3$), 27.2 ($C_{\beta,Glu}$), 26.4 ($C_{\delta,Lys}$), 23.0 ($C_{\gamma,Lys}$), 22.1 ($\underline{C}H_2CH_3$), 13.9 ($CH_3$). The peak for $C_{\varepsilon,Lys}$ was broad and of low intensity and the peak for C=S was not visible in $^{13}$C NMR due to fast quadrupolar relaxation via the nearby $^{14}$N-nuclei. Analytical HPLC gradient 0–95% eluent IV in eluent III (35 min total runtime), $t_R$ 24.5 min (>98%, $UV_{230}$). ESI-MS calcd for $C_{34}H_{57}N_6O_7S^+$ $[M+H]^+$, 693.4; found 693.4.

**(4*S*,7*R*,10*S*)-10-(((benzyloxy)carbonyl)amino)-4-carbamoyl-7-methyl-6,9-dioxo-16-thioxo-5,8,15,17-tetraazanonacosanoic acid (15)**

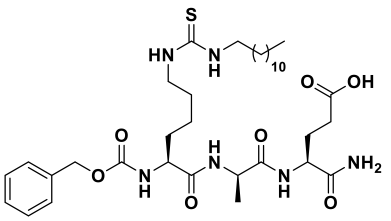

Starting from Cbz-Lys-D-Ala-Glu(*t*Bu)-resin (79 mg, estimated loading: 0.51 mmol/g) synthesized from Fmoc-Glu(*t*Bu)-OH, Fmoc-D-Ala-OH and Cbz-Lys(Fmoc)-OH by SPPS, the title compound was synthesized by the on-resin thiourea formation as described in the general procedures using dodecylamine as the amine source. Global deprotection and cleavage from the resin followed by preparative reverse-phase HPLC purification afforded the desired thiourea (5 mg, 18% based on resin loading) as a colorless fluffy material after lyophilization. $^{1}$H NMR (600 MHz, DMSO-*d*$_6$) δ 12.08 (s, 1H, COOH), 8.16 (d, *J* = 7.2 Hz, 1H, $NH_{D\text{-}Ala}$), 7.88 (d, *J* = 8.3 Hz, 1H, $NH_{Glu}$), 7.46 (d, *J* = 7.4 Hz, 1H, $NH_{\alpha,Lys}$), 7.38–7.27 (m, 8H, $NH_{\varepsilon,Lys}$, $\underline{NH}CH_2$, $H_{Ar,Cbz}$, $CONH_{2,A}$), 7.08 (s, 1H, $CONH_{2,B}$), 5.07–4.95 (m, 2H, $CH_{2,Cbz}$), 4.25 (p, *J* = 7.1 Hz, 1H, $H_{\alpha,D\text{-}Ala}$), 4.18 (td, *J* = 8.8, 5.0 Hz, 1H, $H_{\alpha,Glu}$), 3.99–3.92 (m, 1H, $H_{\alpha,Lys}$), 3.32 (br s, 2H, $H_{\varepsilon,Lys}$, overlap with residual water), 2.23–2.17 (m, 2H, $H_{\gamma,Glu}$), 2.01–1.92 (m, 1H, $H_{\beta,Glu,A}$), 1.78–1.69 (m, 1H, $H_{\beta,Glu,B}$), 1.64–1.55 (m, 1H, $H_{\beta,Lys,A}$), 1.55–1.14 (m, 30H, $H_{\beta,Lys,B}$, $H_{\gamma,Lys}$, $H_{\delta,Lys}$, $H_{\beta,D\text{-}Ala}$, $(C\underline{H}_2)_{11}CH_3$), 0.85 (t, *J* = 7.0 Hz, 3H, $CH_2C\underline{H}_3$). $^{13}$C NMR (151 MHz, DMSO) δ 173.9 (COOH), 173.1 ($CO_{Glu}$), 172.0 ($CO_{Ala}$), 171.9 ($CO_{Lys}$), 156.1 ($CO_{Cbz}$), 136.9 ($C1_{Ar,Cbz}$), 128.3 ($C3_{Ar,Cbz}$, $C5_{Ar,Cbz}$), 127.8 ($C4_{Ar,Cbz}$), 127.7 ($C2_{Ar,Cbz}$, $C6_{Ar,Cbz}$), 65.5 ($CH_{2,Cbz}$), 54.8 ($C_{\alpha,Lys}$), 51.7 ($C_{\alpha,Glu}$), 48.4 ($C_{\alpha,D\text{-}Ala}$), 43.4 ($C_{\varepsilon,Lys}$), 31.4 ($C_{\beta,Lys}$), 31.3 ($\underline{C}H_2CH_2CH_3$), 30.1 ($C_{\gamma,Glu}$), 29.0–28.7 (9C, $(\underline{C}H_2)_{11}CH_3$), 27.1 ($C_{\beta,Glu}$), 26.4 ($C_{\delta,Lys}$), 22.9 ($C_{\gamma,Lys}$), 22.1 ($\underline{C}H_2CH_3$), 18.1 ($C_{\beta,D\text{-}Ala}$), 13.9 ($CH_2\underline{C}H_3$). The peak for $C_{\varepsilon,Lys}$ was broad and of low intensity and the peak for C=S was not visible in $^{13}$C NMR due to fast quadrupolar relaxation via the nearby $^{14}$N-nuclei. Analytical HPLC (Method A) gradient 0–95% eluent IV in eluent III (35 min total runtime), $t_R$ 26.6 min (>98%, $UV_{230}$). ESI-MS calcd for $C_{35}H_{59}N_6O_7S^+$ $[M+H]^+$, 707.4; found 707.5.

**(4*S*,7*S*,10*S*)-10-(((benzyloxy)carbonyl)amino)-4-carbamoyl-7-(hydroxymethyl)-6,9-dioxo-16-thioxo-5,8,15,17-tetraazanonacosanoic acid (16)**

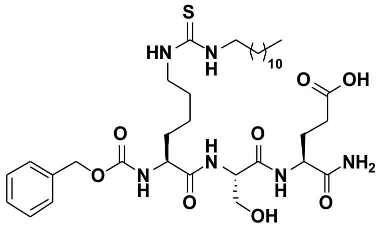

Starting from Cbz-Lys-Ser(*t*Bu)-Glu(*t*Bu)-resin (115 mg, estimated loading: 0.35 mmol/g) synthesized from Fmoc-Glu(*t*Bu)-OH, Fmoc-Ser(*t*Bu)-OH and Cbz-Lys(Fmoc)-OH by SPPS, the title compound was synthesized by the on-resin thiourea formation as described in the general procedures using dodecylamine as the amine source. Global deprotection and cleavage from the resin, followed by preparative reverse-phase HPLC purification afforded the desired thiourea (5 mg, 17% based on resin loading) as a colorless fluffy material after lyophilization. $^{1}$H NMR (600 MHz, DMSO-*d*$_6$) δ 12.07 (s, 1H, COOH), 7.96 (d, *J* = 7.4 Hz, 1H, $NH_{Ser}$), 7.91 (d, *J* = 8.0 Hz, 1H, $NH_{Glu}$), 7.47 (d, *J* = 7.9 Hz, 1H, $NH_{\alpha,Lys}$), 7.38–7.23 (m, 8H, $NH_{\varepsilon,Lys}$, $\underline{NH}CH_2$, $H_{Ar,Cbz}$, $CONH_{2,A}$), 7.15 (s, 1H, $CONH_{2,B}$), 5.11 (s, 1H, $OH_{Ser}$), 5.06–4.98 (m, 2H, $CH_{2,Cbz}$), 4.33–4.24 (m, 1H, $H_{\alpha,Ser}$), 4.18 (td, *J* = 8.5, 4.8 Hz, 1H, $H_{\alpha,Glu}$), 4.04–3.98 (m, 1H, $H_{\alpha,Lys}$), 3.69–3.62 (m, 1H, $H_{\beta,Ser,A}$), 3.56–3.50 (m, 1H, $H_{\beta,Ser,B}$), 3.32 (br s, 2H, $H_{\varepsilon,Lys}$, overlap with residual water), 2.24 (t, *J* = 8.0, 2H, $H_{\gamma,Glu}$), 2.02–1.94 (m, 1H, $H_{\beta,Glu,A}$), 1.79–1.69 (m, 1H, $H_{\beta,Glu,B}$), 1.69–1.61 (m, 1H, $H_{\beta,Lys,A}$), 1.57–1.09 (m, 27H, $H_{\beta,Lys,B}$, $H_{\gamma,Lys}$, $H_{\delta,Lys}$, $(C\underline{H}_2)_{11}CH_3$), 0.85 (t, *J* = 6.9 Hz, 3H, $CH_3$). $^{13}$C NMR (151 MHz, DMSO) δ 173.9 (COOH), 173.1 ($CO_{Glu}$), 172.2 ($CO_{Lys}$), 170.0 ($CO_{Ser}$), 156.1 ($CO_{Cbz}$), 137.0 ($C1_{Ar,Cbz}$), 128.3 ($C3_{Ar,Cbz}$, $C5_{Ar,Cbz}$), 127.7 ($C4_{Ar,Cbz}$), 127.6 ($C2_{Ar,Cbz}$, $C6_{Ar,Cbz}$), 65.4 ($CH_{2,Cbz}$), 61.4 ($C_{\beta,Ser}$) 57.9 ($C_{\alpha,Ser}$), 54.8 ($C_{\alpha,Lys}$), 51.9 ($C_{\alpha,Glu}$), 43.4 ($C_{\varepsilon,Lys}$), 31.5 ($C_{\beta,Lys}$), 31.3 ($\underline{C}H_2CH_2CH_3$), 30.2 ($C_{\gamma,Glu}$), 29.0–28.5 (9C, $(\underline{C}H_2)_{11}$), 27.0 ($C_{\beta,Glu}$), 26.4 ($C_{\delta,Lys}$), 23.0 ($C_{\gamma,Lys}$), 22.1 ($\underline{C}H_2CH_3$), 13.9 ($CH_3$). The peak for $C_{\varepsilon,Lys}$ was broad and of low intensity and the peak for C=S was not visible in $^{13}$C NMR due to fast quadrupolar relaxation via the nearby $^{14}$N-nuclei. Analytical HPLC (Method A) gradient 0–95% eluent IV in eluent III (35 min total runtime), $t_R$ 26.3 min (>97%, $UV_{230}$). ESI-MS calcd for $C_{35}H_{59}N_6O_8S^+$ $[M+H]^+$, 723.4; found 723.5.

**(4*S*,7*S*,10*S*)-10-(((benzyloxy)carbonyl)amino)-4-carbamoyl-7-((R)-1-hydroxyethyl)-6,9-dioxo-16-thioxo-5,8,15,17-tetraazanonacosanoic acid (17)**

Starting from Cbz-Lys-Thr(*t*Bu)-Glu(*t*Bu)-resin (115 mg, estimated loading: 0.35 mmol/g) synthesized from Fmoc-Glu(*t*Bu)-OH, Fmoc-Thr(*t*Bu)-OH and Cbz-Lys(Fmoc)-OH by SPPS, the title compound was synthesized by the on-resin thiourea formation as described in the general procedures using dodecylamine as the amine source. Global deprotection and cleavage from the resin, followed by preparative reverse-phase HPLC purification afforded the desired thiourea (4 mg, 14% based on resin loading) as a colorless fluffy material after lyophilization.

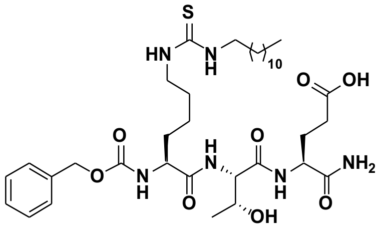

$^{1}$H NMR (600 MHz, DMSO-*d*$_6$) δ 12.08 (br s, 1H, COOH), 7.79 (d, *J* = 8.0 Hz, 1H, $NH_{Glu}$), 7.74 (d, *J* = 8.1 Hz, 1H, $NH_{Thr}$), 7.54 (d, *J* = 7.9 Hz, 1H, $NH_{\alpha,Lys}$), 7.42–7.21 (m, 8H, $NH_{\varepsilon,Lys}$, $\underline{NH}CH_2$, $H_{Ar,Cbz}$, $CONH_{2,A}$), 7.15 (s, 1H, $CONH_{2,B}$), 5.03 (s, 2H, $CH_{2,Cbz}$), 4.25–4.17 (m, 2H, $H_{\alpha,Thr}$, $H_{\alpha,Glu}$), 4.09–4.02 (m, 2H, $H_{\alpha,Lys}$, $H_{\beta,Thr}$), 3.32 (br s, 2H, $H_{\varepsilon,Lys}$, overlap with residual water), 2.23 (t, *J* = 8.0, 2H, $H_{\gamma,Glu}$), 1.99–1.91 (m, 1H, $H_{\beta,Glu,A}$), 1.80–1.71 (m, 1H, $H_{\beta,Glu,B}$), 1.71–1.63 (m, 1H, $H_{\beta,Lys,A}$), 1.58–1.10 (m, 27H, $H_{\beta,Lys,B}$, $H_{\gamma,Lys}$, $H_{\delta,Lys}$, $(C\underline{H_2})_{11}CH_3$), 1.03 (d, *J* =6.3, 3H, $H_{\gamma,Thr,}$), 0.85 (t, *J* = 6.9 Hz, 3H, $CH_2C\underline{H_3}$). $^{13}$C NMR (151 MHz, DMSO) δ 174.0 (COOH), 173.1 ($CO_{Glu}$), 172.3 ($CO_{Lys}$), 169.7 ($CO_{Thr}$), 156.1 ($CO_{Cbz}$), 137.0 ($C1_{Ar,Cbz}$), 128.3 ($C3_{Ar,Cbz}$, $C5_{Ar,Cbz}$), 127.7 ($C4_{Ar,Cbz}$), 127.6 ($C2_{Ar,Cbz,}$ $C6_{Ar,Cbz}$), 66.4 ($C_{\beta,Thr}$), 65.4 ($CH_{2,Cbz}$), 57.9 ($C_{\alpha,Thr}$), 54.8 ($C_{\alpha,Lys}$), 51.8 ($C_{\alpha,Glu}$), 43.5 ($C_{\varepsilon,Lys}$), 31.4 ($C_{\beta,Lys}$), 31.3 ($\underline{C}H_2CH_2CH_3$), 30.2 ($C_{\gamma,Glu}$), 29.0–28.5 (9C, $(\underline{C}H_2)_{11}CH_3$), 27.4 ($C_{\beta,Glu}$), 26.4 ($C_{\delta,Lys}$), 23.0 ($C_{\gamma,Lys}$), 22.1 ($\underline{C}H_2CH_3$), 19.4 ($C_{\gamma,Thr}$), 13.9 ($CH_2\underline{C}H_3$). The peak for $C_{\varepsilon,Lys}$ was broad and of low intensity and the peak for C=S was not visible in $^{13}$C NMR due to fast quadrupolar relaxation via the nearby $^{14}$N-nuclei. Analytical HPLC (Method A) gradient 0–95% eluent IV in eluent III (35 min total runtime), $t_R$ 26.6 min (>98%, $UV_{230}$). ESI-MS calcd for $C_{36}H_{61}N_6O_8S^+$ [M+H]$^+$, 737.4; found 737.6.

**(4*S*,7*S*,10*S*)-7-benzyl-10-(((benzyloxy)carbonyl)amino)-4-carbamoyl-6,9-dioxo-16-thioxo-5,8,15,17-tetraazanonacosanoic acid (18)**

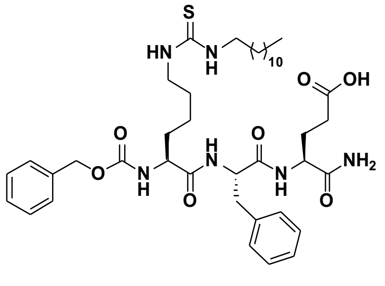

Starting from Cbz-Lys-Phe-Glu(*t*Bu)-resin (115 mg, estimated loading: 0.35 mmol/g) synthesized from Fmoc-Glu(*t*Bu)-OH, Fmoc-Phe-OH and Cbz-Lys(Fmoc)-OH by SPPS, the title compound was synthesized by the on-resin thiourea formation as described in the general procedures using dodecylamine as the amine source. Global deprotection and cleavage from the resin, followed by preparative reverse-phase HPLC purification afforded the desired thiourea (3 mg, 8% based on resin loading) as a colorless fluffy material after lyophilization. $^{1}$H NMR (600 MHz, DMSO-*d*$_6$) δ 12.07 (s, 1H, COOH), 7.98 (d, *J* = 8.1 Hz, 1H, $NH_{Phe}$), 7.92 (d, *J* = 8.0 Hz, 1H, $NH_{Glu}$), 7.42–7.10 (m, 15H, $NH_{\alpha,Lys}$, $NH_{\varepsilon,Lys}$, $\underline{NH}CH_2$, $H_{Ar,Cbz}$, $H_{Ar,Phe}$, $CONH_2$), 5.06–4.96 (m, 2H, $CH_{2,Cbz}$), 4.51 (td, *J* = 8.6, 4.8 Hz, 1H, $H_{\alpha,Phe}$), 4.19 (td, *J* = 8.3, 5.1 Hz, 1H, $H_{\alpha,Glu}$), 3.90 (td, *J* = 8.4, 5.1 Hz, 1H, $H_{\alpha,Lys}$), 3.32 (br s, 2H, $H_{\varepsilon,Lys}$, overlap with residual water), 3.02 (dd, *J* = 13.9, 4.7 Hz, 1H, $H_{\beta,Phe,A}$), 2.83 (dd, *J* = 13.9, 9.3 Hz, 1H, $H_{\beta,Phe,B}$), 2.27–2.15 (m, 2H, $H_{\gamma,Glu}$), 1.96–1.88 (m, 1H, $H_{\beta,Glu,A}$), 1.80–1.71 (m, 1H, $H_{\beta,Glu,B}$), 1.57–1.08 (m, 28H, $H_{\beta,Lys}$, $H_{\gamma,Lys}$, $H_{\delta,Lys}$, $(C\underline{H_2})_{11}CH_3$), 0.85 (t, *J* = 6.9 Hz, 3H, $CH_3$). $^{13}$C NMR (151 MHz, DMSO) δ 173.9 (COOH), 172.8 ($CO_{Glu}$), 171.9 ($CO_{Lys}$), 170.8 ($CO_{Phe}$), 156.0 ($CO_{Cbz}$), 137.6 ($C1_{Ar,Phe}$), 136.9 ($C1_{Ar,Cbz}$), 129.2 ($C3_{Ar,Phe}$, $C5_{Ar,Phe}$), 128.3 ($C3_{Ar,Cbz}$, $C5_{Ar,Cbz}$), 128.0 ($C2_{Ar,Phe}$, $C6_{Ar,Phe}$), 127.8 ($C4_{Ar,Cbz}$), 127.7 ($C2_{Ar,Cbz,}$ $C6_{Ar,Cbz}$), 126.2 ($C4_{Ar,Phe}$), 66.5 ($CH_{2,Cbz}$), 54.9 ($C_{\alpha,Lys}$), 53.8 ($C_{\alpha,Phe}$), 51.8 ($C_{\alpha,Glu}$), 43.3 ($C_{\varepsilon,Lys}$), 37.1 ($C_{\beta,Phe}$), 31.6 ($C_{\beta,Lys}$), 31.3 ($\underline{C}H_2CH_2CH_3$), 30.1 ($C_{\gamma,Glu}$), 29.0–28.5 (9C, $(\underline{C}H_2)_{11}CH_3$), 27.4 ($C_{\beta,Glu}$), 26.4 ($C_{\delta,Lys}$), 22.9 ($C_{\gamma,Lys}$), 22.1 ($\underline{C}H_2CH_3$), 13.9 ($CH_3$). The peak for $C_{\varepsilon,Lys}$ was broad and of low intensity and the peak for C=S was not visible in $^{13}$C NMR due to fast quadrupolar relaxation via the nearby $^{14}$N-nuclei. Analytical HPLC (Method A) gradient 0–95% eluent IV in eluent III (35 min total runtime), $t_R$ 28.3 min (>98%, $UV_{230}$). ESI-MS calcd for $C_{41}H_{63}N_6O_7S^+$ [M+H]$^+$, 783.4; found 784.5.

**(4*S*,7*S*,10*S*)-10-(((benzyloxy)carbonyl)amino)-4-carbamoyl-7-(4-hydroxybenzyl)-6,9-dioxo-16-thioxo-5,8,15,17-tetraazanonacosanoic acid (19)**

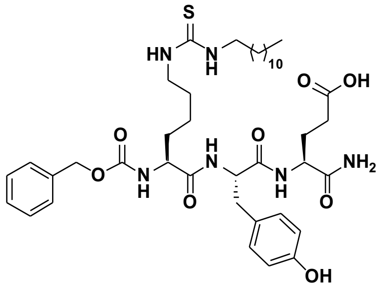

Starting from Cbz-Lys-Tyr(*t*Bu)-Glu(*t*Bu)-resin (115 mg, estimated loading: 0.35 mmol/g) synthesized from Fmoc-Glu(*t*Bu)-OH, Fmoc-Tyr(*t*Bu)-OH and Cbz-Lys(Fmoc)-OH by SPPS, the title compound was synthesized by the on-resin thiourea formation as described in the general procedures using dodecylamine as the amine source. Global deprotection and cleavage from the resin, followed by preparative reverse-phase HPLC purification afforded the desired thiourea (3 mg, 9% based on resin loading) as a colorless fluffy material after lyophilization. $^1$H NMR (600 MHz, DMSO-$d_6$) δ 12.06 (br s, 1H, COOH), 9.14 (s, 1H, $OH_{Tyr}$), 7.92–7.86 (m, 2H, $NH_{Tyr}$, $NH_{Glu}$), 7.42–7.23 (m, 8H, $NH_{\alpha,Lys}$, $NH_{\varepsilon,Lys}$, <u>NH</u>$CH_2$, $H_{Ar,Cbz}$), 7.16 (s, 1H, $CONH_{2,A}$), 7.08 (s, 1H, $CONH_{2,B}$), 7.00 (d, 8.1 Hz, 2H, $H2_{Ar,Tyr}$, $H6_{Ar,Tyr}$), 6.62 (d, 8.4 Hz, 2H, $H3_{Ar,Tyr}$, $H5_{Ar,Tyr}$), 5.06–4.95 (m, 2H, $CH_{2,Cbz}$), 4.41 (td, *J* = 8.4, 4.9 Hz, 1H, $H_{\alpha,Tyr}$), 4.18 (td, *J* = 8.2, 5.1 Hz, 1H, $H_{\alpha,Glu}$), 3.94–3.87 (m, 1H, $H_{\alpha,Lys}$), 3.32 (br s, 2H, $H_{\varepsilon,Lys}$, overlap with residual water), 2.92 (dd, *J* = 14.0, 4.8 Hz, 1H, $H_{\beta,Tyr,A}$), 2.72 (dd, *J* = 14.1, 8.9 Hz, 1H, $H_{\beta,Tyr,B}$), 2.24–2.15 (m, 2H, $H_{\gamma,Glu}$), 1.95–1.88 (m, 1H, $H_{\beta,Glu,A}$), 1.80–1.70 (m, 1H, $H_{\beta,Glu,B}$), 1.57–1.50 (m, 1H, $H_{\beta,Lys,A}$), 1.49–1.10 (m, 27H, $H_{\beta,Lys,B}$, $H_{\gamma,Lys}$, $H_{\delta,Lys}$, (C<u>H</u>$_2$)$_{11}$CH$_3$), 0.85 (t, *J* = 6.9 Hz, 3H, $CH_3$). $^{13}$C NMR (151 MHz, DMSO) δ 173.9 (COOH), 172.9 ($CO_{Glu}$), 171.9 ($CO_{Lys}$), 170.9 ($CO_{Tyr}$), 156.0 ($CO_{Cbz}$), 155.8 ($C4_{Ar,Tyr}$), 136.9 ($C1_{Ar,Cbz}$), 130.1 ($C2_{Ar,Tyr}$, $C6_{Ar,Tyr}$), 128.3 ($C3_{Ar,Cbz}$, $C5_{Ar,Cbz}$), 127.6 ($C1_{Ar,Tyr}$), 127.8 ($C4_{Ar,Cbz}$), 127.7 ($C2_{Ar,Cbz,}$ $C6_{Ar,Cbz}$), 114.8 ($C3_{Ar,Tyr}$, $C5_{Ar,Tyr}$),) 66.5 ($CH_{2,Cbz}$), 54.9 ($C_{\alpha,Lys}$), 54.1 ($C_{\alpha,Tyr}$), 51.7 ($C_{\alpha,Glu}$), 43.4 ($C_{\varepsilon,Lys}$), 36.3 ($C_{\beta,Tyr}$), 31.5 ($C_{\beta,Lys}$), 31.3 (<u>C</u>H$_2$CH$_2$CH$_3$), 30.1 ($C_{\gamma,Glu}$), 29.0–28.5 (9C, (<u>C</u>H$_2$)$_{11}$CH$_3$), 27.4 ($C_{\beta,Glu}$), 26.4 ($C_{\delta,Lys}$), 22.9 ($C_{\gamma,Lys}$), 22.1 (<u>C</u>H$_2$CH$_3$), 13.9 ($CH_3$). The peak for $C_{\varepsilon,Lys}$ was broad and of low intensity and the peak for C=S was not visible in $^{13}$C NMR due to fast quadrupolar relaxation via the nearby $^{14}$N-nuclei. Analytical HPLC (Method A) gradient 0–95% eluent IV in eluent III (35 min total runtime), $t_R$ 27.6 min (>98%, $UV_{254}$). ESI-MS calcd for $C_{41}H_{63}N_6O_8S^+$ $[M+H]^+$, 799.4; found 799.4.

**(4*S*,7*S*,10*S*)-7-(4-aminobutyl)-10-(((benzyloxy)carbonyl)amino)-4-carbamoyl-6,9-dioxo-16-thioxo-5,8,15,17-tetraazanonacosanoic acid (20)**

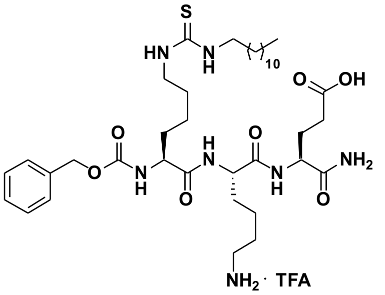

Starting from Cbz-Lys-Lys(Boc)-Glu(*t*Bu)-resin (79 mg, estimated loading: 0.51 mmol/g) synthesized from Fmoc-Glu(*t*Bu)-OH, Fmoc-Lys(Boc)-OH and Cbz-Lys(Fmoc)-OH by SPPS, the title compound was synthesized by the on-resin thiourea formation as described in the general procedures using dodecylamine as the amine source. Global deprotection and cleavage from the resin, followed by preparative reverse-phase HPLC purification afforded the desired thiourea (10 mg, 29% based on resin loading) as a colorless fluffy TFA-salt after lyophilization. $^1$H NMR (600 MHz, DMSO-$d_6$) δ 12.12 (s, 1H, COOH), 8.01 (d, *J* = 7.8 Hz, 1H, $NH_{\alpha,Lys}$), 7.81 (d, *J* = 7.9 Hz, 1H, $NH_{Glu}$), 7.70–7.65 (m, 3H, $NH_3{}^+{}_{\varepsilon,Lys}$), 7.44 (d, *J* = 7.7 Hz, 1H, $NH_{\alpha,Lys(Dtu)}$), 7.39–7.23 (m, 7H, $NH_{\varepsilon,Lys(Dtu)}$, <u>NH</u>$CH_2$, $H_{Ar,Cbz}$, $CONH_{2,A}$), 7.08 (s, 1H, $CONH_{2,B}$), 5.02 ($m_{ABX}$, *J* = 12.7, 2H, $CH_{2,Cbz}$), 4.26–4.14 (m, 2H, $H_{\alpha,Lys(Dtu)}$, $H_{\alpha,Glu}$), 4.00–3.93 (m, 1H, $H_{\alpha,Lys}$), 3.36 (br s, 2H, $H_{\varepsilon,Lys(Dtu)}$, overlap with residual water), 2.79–2.71 (m, 2H, $H_{\varepsilon,Lys}$), 2.21 (m, 2H, $H_{\gamma,Glu}$), 1.94–1.86 (m, 1H, $H_{\beta,Glu,A}$), 1.80–1.16 (m, 35H, $H_{\beta,Glu,B}$ $H_{\beta,Lys}$, $H_{\gamma,Lys}$, $H_{\delta,Lys}$, $H_{\beta,Lys(Dtu)}$, $H_{\gamma,Lys(Dtu)}$, $H_{\delta,Lys(Dtu)}$, (C<u>H</u>$_2$)$_{11}$CH$_3$), 0.85 (t, *J* = 7.0 Hz, 3H, $CH_3$). $^{13}$C NMR (151 MHz, DMSO) δ 173.9 (COOH), 172.9 ($CO_{Glu}$), 172.1 ($CO_{Lys(Dtu)}$), 171.9 ($CO_{Lys}$), 157.9 (q, *J* = 30.8 Hz, $CO_{TFA}$), 156.1 ($CO_{Cbz}$), 137.0 ($C1_{Ar,Cbz}$), 128.3 ($C3_{Ar,Cbz}$, $C5_{Ar,Cbz}$), 127.8 ($C4_{Ar,Cbz}$), 127.6 ($C2_{Ar,Cbz,}$ $C6_{Ar,Cbz}$), 117.3 (q, *J* = 300.6 Hz, $CF_{3,TFA}$), 65.4 ($CH_{2,Cbz}$), 54.8 ($C_{\alpha,Lys(Dtu)}$), 52.3 ($C_{\alpha,Lys}$), 51.6 ($C_{\alpha,Glu}$), 43.4 ($C_{\varepsilon,Lys(Dtu)}$), 38.7 ($C_{\varepsilon,Lys}$), 31.5 ($C_{\beta,Lys(Dtu)}$), 31.3 (<u>C</u>H$_2$CH$_2$CH$_3$), 31.1 ($C_{\beta,Lys}$), 30.1 ($C_{\gamma,Glu}$), 29.0–28.5 (9C, (<u>C</u>H$_2$)$_{11}$CH$_3$), 27.4 ($C_{\beta,Glu}$), 26.6 ($C_{\delta,Lys}$), 26.4 ($C_{\delta,Lys(Dtu)}$), 23.1 ($C_{\gamma,Lys(Dtu)}$), 22.1 (<u>C</u>H$_2$CH$_3$, $C_{\gamma,Lys}$), 13.9 ($CH_3$). The peak for $C_{\varepsilon,Lys(Dtu)}$ was broad and of low intensity and the peak for C=S was not visible in $^{13}$C NMR due to fast quadrupolar relaxation via the nearby $^{14}$N-nuclei. Analytical HPLC (Method A) gradient 0–95% eluent IV in eluent III (35 min total runtime), $t_R$ 23.2 min (>97%, $UV_{230}$). ESI-MS calcd for $C_{38}H_{66}N_7O_7S^+$ $[M+H]^+$, 764.5; found 764.6. Dtu = 1-dodecylthiourea.

**(4*S*,7*R*,10*S*)-7-(4-aminobutyl)-10-(((benzyloxy)carbonyl)amino)-4-carbamoyl-6,9-dioxo-16-thioxo-5,8,15,17-tetraazanonacosanoic acid (21)**

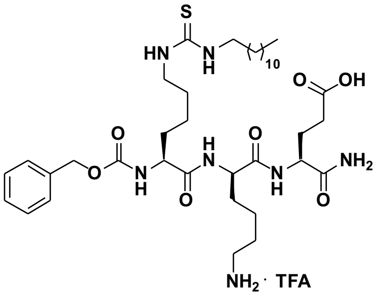

Starting from Cbz-Lys-D-Lys(Boc)-Glu(*t*Bu)-resin (79 mg, estimated loading: 0.51 mmol/g) synthesized from Fmoc-Glu(*t*Bu)-OH, Fmoc-D-Lys(Boc)-OH and Cbz-Lys(Fmoc)-OH by SPPS, the title compound was synthesized by the on-resin thiourea formation as described in the general procedures using dodecylamine as the amine source. Global deprotection and cleavage from the resin, followed by preparative reverse-phase HPLC purification afforded the desired thiourea (9 mg, 29% based on resin loading) as a colorless fluffy material after lyophilization. $^1$H NMR (600 MHz, DMSO-$d_6$) δ 12.11 (br s, 1H, COOH), 8.15 (d, $J$ = 7.8 Hz, 1H, $NH_{\alpha,D\text{-}Lys}$), 7.93 (1d, $J$ = 8.2 Hz, 1H, $NH_{Glu}$), 7.68 (m, 3H, $NH_3{}^+{}_{\varepsilon,Lys}$), 7.48 (d, $J$ = 7.2 Hz, 1H, $NH_{\alpha,Lys(Dtu)}$), 7.40–7.26 (m, 8H, $NH_{\varepsilon,Lys(Dtu)}$, $\underline{NH}CH_2$, $H_{Ar,Cbz}$, $CONH_{2,A}$), 7.09 (s, 1H, $CONH_{2,B}$), 5.09–4.93 (m, 2H, $CH_{2,Cbz}$), 4.24–4.17 (m, 2H, $H_{\alpha,Glu}$, $H_{\alpha,D\text{-}Lys}$), 4.00–3.93 (m, 1H, $H_{\alpha,Lys(Dtu)}$), 3.32 (br s, 2H, $H_{\varepsilon,Lys(Dtu)}$, overlap with residual water), 2.77–2.70 (m, 2H, $H_{\varepsilon,D\text{-}Lys}$), 2.24–2.15 (m, 2H, $H_{\gamma,Glu}$), 2.00–1.92 (m, 1H, $H_{\beta,Glu,A}$), 1.79–1.10 (m, 35H, $H_{\beta,Glu,B}$, $H_{\beta,Lys}$, $H_{\gamma,Lys}$, $H_{\delta,Lys}$, $H_{\beta,Lys(Dtu)}$, $H_{\gamma,Lys(Dtu)}$, $H_{\delta,Lys(Dtu)}$, $(C\underline{H}_2)_{11}CH_3$), 0.85 (t, $J$ = 7.0 Hz, 3H, $CH_3$). $^{13}$C NMR (151 MHz, DMSO) δ 173.9 (COOH), 173.0 ($CO_{Lys(Dtu)}$), 172.3 ($CO_{Lys}$), 171.9 ($CO_{Glu}$), 157.9 (d, $J$ = 30.6 Hz, $CO_{TFA}$), 156.1 ($CO_{Cbz}$), 136.7 ($C1_{Ar,Cbz}$), 128.3 ($C3_{Ar,Cbz}$, $C5_{Ar,Cbz}$), 127.8 ($C4_{Ar,Cbz}$, $C2_{Ar,Cbz}$, $C6_{Ar,Cbz}$), 117.32 (d, $J$ = 300.8 Hz, $CF_{3,TFA}$), 65.5 ($CH_{2,Cbz}$), 54.9 ($C_{\alpha,Lys(Dtu)}$), 52.4 ($C_{\alpha,Lys}$), 51.6 ($C_{\alpha,Glu}$), 43.5 ($C_{\varepsilon,Lys(Dtu)}$), 38.6 ($C_{\varepsilon,Lys}$), 31.4 ($C_{\beta,Lys(Dtu)}$), 31.3 ($\underline{C}H_2CH_2CH_3$), 31.0 ($C_{\beta,Lys}$), 30.1 ($C_{\gamma,Glu}$), 29.0–28.5 (9C, $(\underline{C}H_2)_{11}CH_3$), 27.1 ($C_{\beta,Glu}$), 26.5 ($C_{\delta,Lys}$), 26.4 ($C_{\delta,Lys(Dtu)}$), 23.0 ($C_{\gamma,Lys(Dtu)}$), 22.2 ($C_{\gamma,Lys}$), 22.1 ($\underline{C}H_2CH_3$), 13.9 ($CH_3$). The peak for $C_{\varepsilon,Lys(Dtu)}$ was broad and of low intensity and the peak for C=S was not visible in $^{13}$C NMR due to fast quadrupolar relaxation via the nearby $^{14}$N-nuclei. Analytical HPLC (Method A) gradient 0–95% eluent IV in eluent III (35 min total runtime), $t_R$ 23.2 min (>98%, $UV_{230}$). ESI-MS calcd for $C_{38}H_{66}N_7O_7S^+$ $[M+H]^+$, 764.5; found 764.7. Dtu = 1-dodecylthiourea.

**(4*S*,7*S*,10*S*)-10-(((benzyloxy)carbonyl)amino)-4-carbamoyl-7-(3-guanidinopropyl)-6,9-dioxo-16-thioxo-5,8,15,17-tetraazanonacosanoic acid (22)**

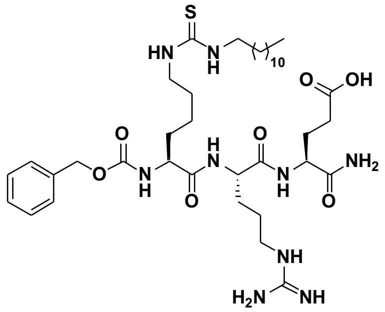

Starting from Cbz-Lys-Arg(Pbf)-Glu(*t*Bu)-resin (79 mg, estimated loading: 0.51 mmol/g) synthesized from Fmoc-Glu(*t*Bu)-OH, Fmoc-Arg(Pbf)-OH and Cbz-Lys(Fmoc)-OH by SPPS, the title compound was synthesized by the on-resin thiourea formation as described in the general procedures using dodecylamine as the amine source. Global deprotection and cleavage from the resin, followed by preparative reverse-phase HPLC purification afforded the desired thiourea (6 mg, 19% based on resin loading) as a colorless fluffy material after lyophilization. $^1$H NMR (600 MHz, DMSO-$d_6$) δ 12.12 (s, COOH), 8.05 (d, $J$ = 7.7 Hz, 1H, $NH_{\alpha,Arg}$), 7.84 (d, $J$ = 7.8 Hz, 1H, $NH_{Glu}$), 7.54 (t, $J$ = 5.7 Hz, 1H, $NH_{\delta,Arg}$), 7.46 (d, $J$ = 7.8 Hz, 1H, $NH_{\alpha,Lys}$), 7.43–7.24 (m, 8H, $NH_{\varepsilon,Lys}$, $\underline{NH}CH_2$, $H_{Ar,Cbz}$, $CONH_{2,A}$), 7.08 (s, 1H, $CONH_{2,B}$), 5.07–4.97 (m, 2H, $CH_{2,Cbz}$), 4.29–4.23 (m, 1H, $H_{\alpha,Arg}$), 4.19 (td, $J$ = 8.1, 5.2 Hz, 1H, $H_{\alpha,Glu}$), 4.04–3.93 (m, 1H, $H_{\alpha,Lys}$), 3.32 (br s, 2H, $H_{\varepsilon,Lys}$, overlap with residual water), 3.09 (q, $J$ = 6.6 Hz, 2H, $H_{\delta,Arg}$), 2.27–2.15 (m, 2H, $H_{\gamma,Glu}$), 1.95–1.86 (m, 1H, $H_{\beta,Glu,A}$), 1.80–1.13 (m, 33H, $H_{\beta,Glu,A}$, $H_{\beta,Lys}$, $H_{\gamma,Lys}$, $H_{\delta,Lys}$, $H_{\beta,Arg}$, $H_{\gamma,Arg}$, $(C\underline{H}_2)_{11}CH_3$), 0.85 (t, $J$ = 6.9 Hz, 3H, $CH_3$). $^{13}$C NMR (151 MHz, DMSO) δ 173.9 (COOH), 172.9 ($CO_{Glu}$), 172.2 ($CO_{Lys}$), 171.1 ($CO_{Arg}$), 156.1 ($CO_{Cbz}$), 137.0 ($C1_{Ar,Cbz}$), 128.3 ($C3_{Ar,Cbz}$, $C5_{Ar,Cbz}$), 127.8 ($C4_{Ar,Cbz}$), 127.6 ($C2_{Ar,Cbz}$, $C6_{Ar,Cbz}$), 65.4 ($CH_{2,Cbz}$), 54.8

($C_{\alpha,Lys}$), 52.2 ($C_{\alpha,Arg}$), 51.7 ($C_{\alpha,Glu}$), 43.3 ($C_{\varepsilon,Lys}$), 40.4 ($C_{\delta,Arg}$), 31.5 ($C_{\beta,Lys}$), 31.3 ($\underline{C}H_2CH_2CH_3$), 30.1 ($C_{\gamma,Glu}$), 29.0–28.5 (10C, $C_{\beta,Arg}$, ($\underline{C}H_2)_{11}CH_3$), 27.5 ($C_{\beta,Glu}$), 26.4 ($C_{\delta,Lys}$), 24.9 ($C_{\gamma,Arg}$), 23.1 ($C_{\gamma,Lys}$), 22.1 ($\underline{C}H_2CH_3$), 13.9 ($CH_3$). The peak for $C_{\varepsilon,Lys}$ was broad and of low intensity and the peak for C=S was not visible in $^{13}C$ NMR due to fast quadrupolar relaxation via the nearby $^{14}N$-nuclei. Analytical HPLC (Method A) gradient 0–95% eluent IV in eluent III (35 min total runtime), $t_R$ 23.2 min (>98%, $UV_{230}$). HRMS calcd for $C_{38}H_{66}N_9O_7S^+$ $[M+H]^+$, 792.5; found 792.7.

**(4*S*,7*S*,10*S*)-10-(((benzyloxy)carbonyl)amino)-4-carbamoyl-7-(2-carboxyethyl)-6,9-dioxo-16-thioxo-5,8,15,17-tetraazanonacosanoic acid (23)**

Starting from Cbz-Lys-Glu(*t*Bu)-Glu(*t*Bu)-resin (115 mg, estimated loading: 0.35 mmol/g) synthesized from Fmoc-Glu(*t*Bu)-OH and Cbz-Lys(Fmoc)-OH by SPPS, the title compound was synthesized by the on-resin thiourea formation as described in the general procedures using dodecylamine as the amine source. Global deprotection and cleavage from the resin, followed by preparative reverse-phase HPLC purification afforded the desired thiourea (6 mg, 20% based on resin loading) as a colorless fluffy material after lyophilization. $^1H$ NMR (600 MHz, DMSO-$d_6$) δ 12.11 (br s, 2H, $COOH_{,1}$, $COOH_{,2}$), 8.05 (d, $J$ = 7.6 Hz, 1H, $NH_{Glu,1}$), 7.83 (d, $J$ = 7.9 Hz, 1H, $NH_{Glu,2}$), 7.46 (d, $J$ = 7.7 Hz, 1H, $NH_{\alpha,Lys}$), 7.41–7.20 (m, 8H, $NH_{\varepsilon,Lys}$, $\underline{NH}CH_2$, $H_{Ar,Cbz}$, $CONH_{2,A}$), 7.07 (d, $J$ = 2.2 Hz, 1H, $CONH_{2,B}$), 5.09–4.97 (m, 2H, $CH_{2,Cbz}$), 4.24 (td, $J$ = 8.1, 5.4 Hz, 1H, $H_{\alpha,Glu,1}$), 4.17 (td, $J$ = 8.2, 5.1 Hz, 1H, $H_{\alpha,Glu,2}$), 4.02–3.93 (m, 1H, $H_{\alpha,Lys}$), 3.32 (br s, 2H, $H_{\varepsilon,Lys}$, overlap with residual water), 2.26–2.16 (m, 4H, $H_{\gamma,Glu,1}$, $H_{\gamma,Glu,2}$), 1.96–1.85 (m, 2H, $H_{\beta,Glu,1,A}$, $H_{\beta,Glu,2,A}$), 1.81–1.69 (m, 2H, $H_{\beta,Glu,1,B}$, $H_{\beta,Glu,2,B}$), 1.69–1.58 (1H, $H_{\beta,Lys,A}$), 1.56–1.18 (m, 27H, $H_{\beta,Lys,B}$, $H_{\gamma,Lys}$, $H_{\delta,Lys}$, ($C\underline{H}_2)_{11}CH_3$), 0.85 (t, $J$ = 7.0 Hz, 3H, $CH_3$). $^{13}C$ NMR (151 MHz, DMSO) δ 174.0 ($COOH_{,1}$), 173.9 ($COOH_{,2}$), 172.9 ($CO_{Glu,2}$), 172.2 ($CO_{Lys}$), 170.9 ($CO_{Glu,1}$), 156.1 ($CO_{Cbz}$), 137.0 ($C1_{Ar,Cbz}$), 128.3 ($C3_{Ar,Cbz}$, $C5_{Ar,Cbz}$), 127.8 ($C4_{Ar,Cbz}$), 127.7 ($C2_{Ar,Cbz}$, $C6_{Ar,Cbz}$), 65.4 ($CH_{2,Cbz}$), 54.8 ($C_{\alpha,Lys}$), 52.0 ($C_{\alpha,Glu,1}$), 51.7 ($C_{\alpha,Glu,2}$), 43.4 ($C_{\varepsilon,Lys}$), 31.4 ($C_{\beta,Lys}$), 31.3 ($\underline{C}H_2CH_2CH_3$), 30.1 ($C_{\gamma,Glu,1}$, $C_{\gamma,Glu,2}$), 29.0–28.5 (9C, ($\underline{C}H_2)_{11}CH_3$), 27.3 ($C_{\beta,Glu,2}$), 27.1 ($C_{\beta,Glu,1}$), 26.4 ($C_{\delta,Lys}$), 23.0 ($C_{\gamma,Lys}$), 22.1 ($\underline{C}H_2CH_3$), 13.9 ($CH_3$). The peak for $C_{\varepsilon,Lys}$ was broad and of low intensity and the peak for C=S was not visible in $^{13}C$ NMR due to fast quadrupolar relaxation via the nearby $^{14}N$-nuclei. Analytical HPLC (Method A) gradient 0–95% eluent IV in eluent III (35 min total runtime), $t_R$ 26.2 min (>98%, $UV_{230}$). ESI-MS calcd for $C_{37}H_{61}N_6O_9S^+$ $[M+H]^+$, 765.4; found 765.5.

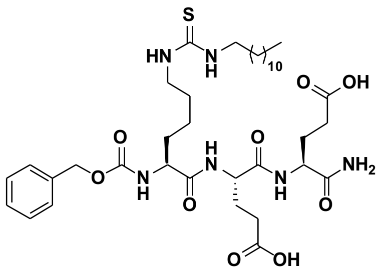

**(4*S*,7*R*,10*S*)-10-(((benzyloxy)carbonyl)amino)-4-carbamoyl-7-(2-carboxyethyl)-6,9-dioxo-16-thioxo-5,8,15,17-tetraazanonacosanoic acid (24)**

Starting from Cbz-Lys-D-Glu(*t*Bu)-Glu(*t*Bu)-resin (115 mg, estimated loading: 0.35 mmol/g) synthesized from Fmoc-Glu(*t*Bu)-OH, Fmoc-D-Glu(*t*Bu)-OH and Cbz-Lys(Fmoc)-OH by SPPS, the title compound was synthesized by the on-resin thiourea formation as described in the general procedures using dodecylamine as the amine source. Global deprotection and cleavage from the resin, followed by preparative reverse-phase HPLC purification afforded the desired thiourea (7 mg, 22% based on resin loading) as a colorless fluffy material after lyophilization. $^1H$ NMR (600 MHz, DMSO-$d_6$) δ 12.10 (s, 2H, $COOH_{,1}$, $COOH_{,2}$), 8.19 (d, $J$ = 7.6 Hz, 1H, $NH_{D\text{-}Glu}$), 7.89 (d, $J$ = 8.2 Hz, 1H, $NH_{Glu}$), 7.47 (d, $J$ = 7.2 Hz, 1H, $NH_{\alpha,Lys}$), 7.38–7.21 (m, 8H, $NH_{\varepsilon,Lys}$, $\underline{NH}CH_2$, $H_{Ar,Cbz}$, $CONH_{2,A}$), 7.08 (s, 1H, $CONH_{2,B}$), 5.07–4.95 (m, 2H, $CH_{2,Cbz}$), 4.24 (td, $J$ = 8.3, 5.3 Hz, 1H, $H_{\alpha,\,D\text{-}Glu}$), 4.18 (td, $J$ = 8.6, 5.1 Hz, 1H, $H_{\alpha,Glu}$), 3.96 (ddd, $J$ = 8.9, 7.2, 5.5 Hz, 1H, $H_{\alpha,Lys}$), 3.32 (br s, 2H, $H_{\varepsilon,Lys}$, overlap with residual water), 2.27–2.14 (m, 4H, $H_{\gamma,Glu}$, $H_{\gamma,D\text{-}Glu}$), 2.02–1.91 (m, 2H, $H_{\beta,Glu,A}$, $H_{\beta,D\text{-}Glu,A}$), 1.80–1.69 (m, 2H, $H_{\beta,Glu,B}$, $H_{\beta,D\text{-}Glu,B}$), 1.64–1.12 (m, 28H, $H_{\beta,Lys}$, $H_{\gamma,Lys}$, $H_{\delta,Lys}$, ($C\underline{H}_2)_{11}CH_3$), 0.85 (t, $J$ = 6.9 Hz, 3H, $CH_3$). $^{13}C$ NMR (151 MHz, DMSO) δ 173.84 (COOH), 173.83 (COOH), 173.0 ($CO_{Glu}$), 172.3 ($CO_{Lys}$), 171.0 ($CO_{D\text{-}Glu}$), 156.1 ($CO_{Cbz}$), 136.9 ($C1_{Ar,Cbz}$), 128.3 ($C3_{Ar,Cbz}$, $C5_{Ar,Cbz}$), 127.8 ($C2_{Ar,Cbz}$, $C4_{Ar,Cbz}$, $C6_{Ar,Cbz}$), 65.5 ($CH_{2,Cbz}$),

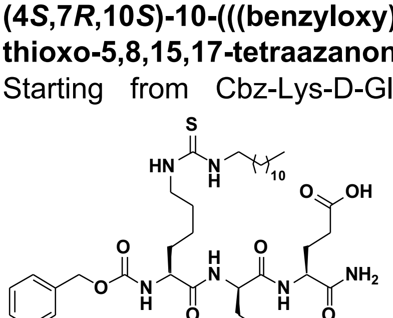

54.9 ($C_{\alpha,Lys}$), 52.1 ($C_{\alpha,D\text{-}Glu}$), 51.7 ($C_{\alpha,Glu}$), 43.4 ($C_{\varepsilon,Lys}$), 31.32 ($C_{\beta,Lys}$), 31.28 ($\underline{C}H_2CH_2CH_3$), 30.1 ($C_{\gamma,Glu}$), 30.0 ($C_{\gamma,D\text{-}Glu}$), 29.0–28.5 (9C, $(\underline{C}H_2)_{11}CH_3$), 27.03 ($C_{\beta,Glu}/C_{\beta,D\text{-}Glu}$), 26.99 ($C_{\beta,Glu}/C_{\beta,D\text{-}Glu}$), 26.4 ($C_{\delta,Lys}$), 22.9 ($C_{\gamma,Lys}$), 22.1 ($\underline{C}H_2CH_3$), 13.9 ($CH_3$).The peak for $C_{\varepsilon,Lys}$ was broad and of low intensity and the peak for C=S was not visible in $^{13}$C NMR due to fast quadrupolar relaxation via the nearby $^{14}$N-nuclei. Analytical HPLC (Method A) gradient 0–95% eluent IV in eluent III (35 min total runtime), $t_R$ 26.2 min (>98%, $UV_{230}$). ESI-MS calcd for $C_{37}H_{61}N_6O_9S^+$ [M+H]$^+$, 765.4; found 765.4.

**(4*S*,7*S*,10*S*)-7-(3-amino-3-oxopropyl)-10-(((benzyloxy)carbonyl)amino)-4-carbamoyl-6,9-dioxo-16-thioxo-5,8,15,17-tetraazanonacosanoic acid (25)**

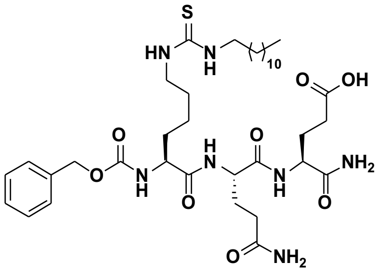

Starting from Cbz-Lys-Gln(Trt)-Glu(*t*Bu)-resin (79 mg, estimated loading: 0.51 mmol/g) synthesized from Fmoc-Glu(*t*Bu)-OH, Fmoc-Gln(Trt)-OH and Cbz-Lys(Fmoc)-OH by SPPS, the title compound was synthesized by the on-resin thiourea formation as described in the general procedures using dodecylamine as the amine source. Global deprotection and cleavage from the resin, followed by preparative reverse-phase HPLC purification afforded the desired thiourea (7 mg, 23% based on resin loading) as a colorless fluffy material after lyophilization. $^1$H NMR (600 MHz, DMSO-$d_6$) δ 12.08 (s, 1H, COOH), 8.09 (d, $J$ = 7.5 Hz, 1H, $NH_{\alpha,Gln}$), 7.83 (d, $J$ = 7.9 Hz, 1H, $NH_{Glu}$), 7.46 (d, $J$ = 7.6 Hz, 1H, $NH_{\alpha,Lys}$), 7.41–7.20 (m, 9H, $NH_{\varepsilon,Lys}$, $\underline{NH}CH_2$, $H_{Ar,Cbz}$, $CONH_{2,A}$, $NH_{\delta,Gln,A}$), 7.08 (s, 1H, $CONH_{2,B}$), 6.76 (s, 1H, $NH_{\delta,Gln,B}$), 5.09–4.96 (m, 2H, $CH_{2,Cbz}$), 4.23–4.14 (m, 2H, $H_{\alpha,Gln}$, $H_{\alpha,Glu}$), 4.00–3.93 (m, 1H, $H_{\alpha,Lys}$), 3.31 (br s, 2H, $H_{\varepsilon,Lys}$, overlap with residual water), 2.24–2.17 (m, 2H, $H_{\gamma,Glu}$), 2.15–2.03 (m, 2H, $H_{\gamma,Gln}$), 1.96–1.85 (m, 2H, $H_{\beta,Glu,A}$, $H_{\beta,Gln,A}$), 1.80–1.70 (m, 2H, $H_{\beta,Glu,B}$, $H_{\beta,Gln,B}$), 1.66–1.59 (m, 1H, $H_{\beta,Lys,A}$), 1.37–1.17 (m, 27H, $H_{\beta,Lys,B}$, $H_{\gamma,Lys}$, $H_{\delta,Lys}$, $(C\underline{H_2})_{11}CH_3$), 0.85 (t, $J$ = 7.0 Hz, 3H, $CH_3$). $^{13}$C NMR (151 MHz, DMSO) δ 173.9 (COOH), 173.8 ($CO_{\delta,Gln}$), 173.0 ($CO_{Glu}$), 172.2 ($CO_{Lys}$), 171.1 ($CO_{\alpha,Gln}$), 156.1 ($CO_{Cbz}$), 137.0 ($C1_{Ar,Cbz}$), 128.3 ($C3_{Ar,Cbz}$, $C5_{Ar,Cbz}$), 127.8 ($C4_{Ar,Cbz}$), 127.7 ($C2_{Ar,Cbz}$, $C6_{Ar,Cbz}$), 65.4 ($CH_{2,Cbz}$), 54.8 ($C_{\alpha,Lys}$), 52.4 ($C_{\alpha,Gln}$) 51.7 ($C_{\alpha,Glu}$), 43.5 ($C_{\varepsilon,Lys}$), 31.5 ($C_{\beta,Lys}$), 31.4 ($C_{\gamma,Gln}$), 31.3 ($\underline{C}H_2CH_2CH_3$), 30.1 ($C_{\gamma,Glu}$), 29.0–28.7 (9C, $(\underline{C}H_2)_{11}CH_3$), 27.6 ($C_{\gamma,Gln}$), 27.3 ($C_{\beta,Glu}$), 26.4 ($C_{\delta,Lys}$), 23.0 ($C_{\gamma,Lys}$), 22.1 ($\underline{C}H_2CH_3$), 13.9 ($CH_3$). The peak for $C_{\varepsilon,Lys}$ was broad and of low intensity and the peak for C=S was not visible in $^{13}$C NMR due to fast quadrupolar relaxation via the nearby $^{14}$N-nuclei. Analytical HPLC (Method A) gradient 0–95% eluent IV in eluent III (35 min total runtime), $t_R$ 25.8 min (>98%, $UV_{230}$). ESI-MS calcd for $C_{37}H_{62}N_7O_8S^+$ [M+H]$^+$, 764.4; found 764.6.

**(4*S*,7*S*,10*S*)-10-(((benzyloxy)carbonyl)amino)-4-carbamoyl-7-isobutyl-6,9-dioxo-16-thioxo-5,8,15,17-tetraazanonacosanoic acid (26)**

Starting from Cbz-Lys-Leu-Glu(*t*Bu)-resin (79 mg, estimated loading: 0.51 mmol/g) synthesized from

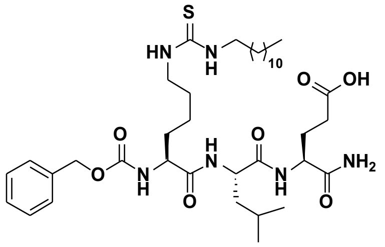

Fmoc-Glu(*t*Bu)-OH, Fmoc-Leu-OH and Cbz-Lys(Fmoc)-OH by SPPS, the title compound was synthesized by the on-resin thiourea formation as described in the general procedures using dodecylamine as the amine source. Global deprotection and cleavage from the resin, followed by preparative reverse-phase HPLC purification afforded the desired thiourea (5 mg, 16% based on resin loading) as a colorless fluffy material after lyophilization. $^1$H NMR (600 MHz, DMSO-$d_6$) δ 12.08 (br s, 1H, COOH), 7.98 (d, $J$ = 7.9 Hz, 1H, $NH_{Leu}$), 7.76 (d, $J$ = 8.0 Hz, 1H, $NH_{Glu}$), 7.43 (d, $J$ = 7.8 Hz, 1H, $NH_{\alpha,Lys}$), 7.39–7.19 (m, 8H, $NH_{\varepsilon,Lys}$, $\underline{NH}CH_2$, $H_{Ar,Cbz}$, $CONH_{2,A}$), 7.07 (s, 1H, $CONH_{2,B}$), 5.06–4.93 (m, 2H, $CH_{2,Cbz}$), 4.26 (td, $J$ = 7.7 Hz, 1H, $H_{\alpha,Leu}$), 4.17 (td, $J$ = 8.3, 5.1 Hz, 1H, $H_{\alpha,Glu}$), 4.00–3.93 (m, 1H, $H_{\alpha,Lys}$), 3.34 (br s, 2H, $H_{\varepsilon,Lys}$, overlap with residual water), 2.24–2.16 (m, 2H, $H_{\gamma,Glu}$), 1.94–1.87 (m, 1H, $H_{\beta,Glu,A}$), 1.79–1.70 (m, 1H, $H_{\beta,Glu,B}$), 1.68–1.15 (m, 32H, $H_{\beta,Lys}$, $H_{\gamma,Lys}$, $H_{\delta,Lys}$, $H_{\beta,Leu}$, $H_{\gamma,Leu}$, $(C\underline{H_2})_{11}CH_3$), 0.90–0.78 (m, 9H, $H_{\delta,Leu,1}$, $H_{\delta,Leu,2}$, $CH_2C\underline{H_3}$). $^{13}$C NMR (151 MHz, DMSO) δ 173.9 (COOH), 172.9 ($CO_{Glu}$), 172.0 ($CO_{Lys}$), 171.8 ($CO_{Leu}$), 156.1 ($CO_{Cbz}$), 137.0 ($C1_{Ar,Cbz}$), 128.3 ($C3_{Ar,Cbz}$, $C5_{Ar,Cbz}$), 127.8 ($C4_{Ar,Cbz}$), 127.7 ($C2_{Ar,Cbz}$, $C6_{Ar,Cbz}$), 65.4 ($CH_{2,Cbz}$), 54.8 ($C_{\alpha,Lys}$), 51.6 ($C_{\alpha,Glu}$), 51.2 ($C_{\alpha,Leu}$), 43.4 ($C_{\varepsilon,Lys}$), 40.4

($C_{\beta,Leu}$), 31.5 ($C_{\beta,Lys}$), 31.3 ($\underline{C}H_2CH_2CH_3$), 30.1 ($C_{\gamma,Glu}$), 29.1–28.5 (9C, ($\underline{C}H_2)_{11}CH_3$), 27.4 ($C_{\beta,Glu}$), 26.4 ($C_{\delta,Lys}$), 24.1($C_{\delta,Leu,1}$), 23.1 ($C_{\gamma,Leu}$), 23.0 ($C_{\gamma,Lys}$), 22.1 ($\underline{C}H_2CH_3$), 21.5 ($C_{\delta,Leu,2}$), 13.9 ($CH_2\underline{C}H_3$). The peak for $C_{\varepsilon,Lys}$ was broad and of low intensity and the peak for C=S was not visible in $^{13}$C NMR due to fast quadrupolar relaxation via the nearby $^{14}$N-nuclei. Analytical HPLC (Method A) gradient 0–95% eluent IV in eluent III (35 min total runtime), $t_R$ 26.1 min (>98%, $UV_{230}$). ESI-MS calcd for $C_{36}H_{65}N_6O_7S^+$ $[M+H]^+$, 749.5; found 749.6.

**(4*S*,7*S*,10*S*)-10-(((benzyloxy)carbonyl)amino)-4-carbamoyl-7-isopropyl-6,9-dioxo-16-thioxo-5,8,15,17-tetraazanonacosanoic acid (27)**

Starting from Cbz-Lys-Val-Glu(*t*Bu)-resin (115 mg, estimated loading: 0.35 mmol/g) synthesized from Fmoc-Glu(*t*Bu)-OH, Fmoc-Val-OH and Cbz-Lys(Fmoc)-OH by SPPS, the title compound was synthesized by the on-resin thiourea formation as described in the general procedures using dodecylamine as the amine source. Global deprotection and cleavage from the resin, followed by preparative reverse-phase HPLC purification afforded the desired thiourea (4 mg, 14% based on resin loading) as a colorless fluffy material after lyophilization. $^1$H NMR (600 MHz, DMSO-$d_6$) δ 12.08 (s, 1H, COOH), 7.88 (d, $J$ = 7.8 Hz, 1H, $NH_{Glu}$), 7.76 (d, $J$ = 8.6 Hz, 1H, $NH_{Val}$), 7.47 (d, $J$ = 8.1 Hz, 1H, $NH_{\alpha,Lys}$), 7.39–7.26 (m, 8H, $NH_{\varepsilon,Lys}$, $\underline{NH}CH_2$, $H_{Ar,Cbz}$, $CONH_{2,A}$), 7.06 (s, 1H, $CONH_{2,B}$), 5.04–4.98 (m, 2H, $CH_{2,Cbz}$), 4.23–4.14 (m, 2H, $H_{\alpha,Val}$, $H_{\alpha,Glu}$), 4.02 (td, $J$ = 8.8, 4.8 Hz, 1H, $H_{\alpha,Lys}$), 3.32 (br s, 2H, $H_{\varepsilon,Lys}$, overlap with residual water), 2.20 (t, $J$ = 9,5, 8.9 Hz, 2H, $H_{\gamma,Glu}$), 2.03–1.95 (m, 1H, $H_{\beta,Val}$), 1.94–1.85 (m, 1H, $H_{\beta,Glu,A}$), 1.79–1.70 (m, 1H, $H_{\beta,Glu,B}$), 1.67–1.59 (m, 1H, $H_{\beta,Lys,A}$), 1.56–1.09 (m, 29H, $H_{\beta,Lys,B}$, $H_{\gamma,Lys}$, $H_{\delta,Lys}$, $H_{\gamma,Leu,}$ ($C\underline{H}_2)_{11}CH_3$), 0.91–0.79 (m, 9H, $H_{\gamma,Val}$, $CH_2C\underline{H}_3$). $^{13}$C NMR (151 MHz, DMSO) δ 173.8 (COOH), 172.8 ($CO_{Glu}$), 172.0 ($CO_{Lys}$), 170.7 ($CO_{Val}$), 156.0 ($CO_{Cbz}$), 137.0 ($C1_{Ar,Cbz}$), 128.3 ($C3_{Ar,Cbz}$, $C5_{Ar,Cbz}$), 127.7 ($C4_{Ar,Cbz}$), 127.6 ($C2_{Ar,Cbz,}$ $C6_{Ar,Cbz}$), 66.3 ($CH_{2,Cbz}$), 57.5 ($C_{\alpha,Val}$), 54.8 ($C_{\alpha,Lys}$), 51.7 ($C_{\alpha,Glu}$), 43.5 ($C_{\varepsilon,Lys}$), 31.5 ($C_{\beta,Lys}$), 31.3 ($\underline{C}H_2CH_2CH_3$), 30.5 ($C_{\beta,Val}$), 30.1 ($C_{\gamma,Glu}$), 29.0–28.5 (9C, ($\underline{C}H_2)_{11}CH_3$), 27.4 ($C_{\beta,Glu}$), 26.4 ($C_{\delta,Lys}$), 23.0 ($C_{\gamma,Lys}$), 22.1 ($\underline{C}H_2CH_3$), 19.2 ($C_{\gamma,Val,1}$), 18.0 ($C_{\gamma,Val,2}$), 13.9 ($CH_2\underline{C}H_3$). The peak for $C_{\varepsilon,Lys}$ was broad and of low intensity and the peak for C=S was not visible in $^{13}$C NMR due to fast quadrupolar relaxation via the nearby $^{14}$N-nuclei. Analytical HPLC (Method A) gradient 0–95% eluent IV in eluent III (35 min total runtime), $t_R$ 27.6 min (>95%, $UV_{230}$). ESI-MS calcd for $C_{37}H_{63}N_6O_7S^+$ $[M+H]^+$, 735.4; found 735.5.

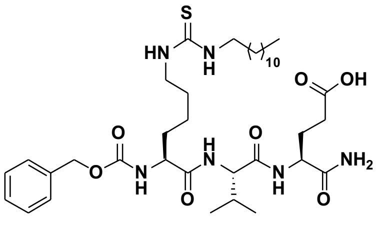

**(4*S*,7*R*,10*S*)-10-(((benzyloxy)carbonyl)amino)-4-carbamoyl-7-isopropyl-6,9-dioxo-16-thioxo-5,8,15,17-tetraazanonacosanoic acid (28)**

Starting from Cbz-Lys-D-Val-Glu(*t*Bu)-resin (115 mg, estimated loading: 0.35 mmol/g) synthesized from Fmoc-Glu(*t*Bu)-OH, Fmoc-D-Val-OH and Cbz-Lys(Fmoc)-OH by SPPS, the title compound was synthesized by the on-resin thiourea formation as described in the general procedures using dodecylamine as the amine source. Global deprotection and cleavage from the resin, followed by preparative reverse-phase HPLC purification afforded the desired thiourea (3 mg, 10% based on resin loading) as a colorless fluffy material after lyophilization. $^1$H NMR (600 MHz, DMSO-$d_6$) δ 12.08 (s, 1H, COOH), 8.05 (d, $J$ = 8.2 Hz, 1H, $NH_{Glu}$), 7.93 (d, $J$ = 8.4 Hz, 1H, $NH_{D-Val}$), 7.45–7.19 (m, 10H, $NH_{\alpha,Lys}$, $NH_{\varepsilon,Lys}$, $\underline{NH}CH_2$, $H_{Ar,Cbz}$, $CONH_{2,A}$), 7.06 (s, 1H, $CONH_{2,B}$), 5.01 ($m_{ABX}$, $J$ = 12.5 Hz, 2H, $CH_{2,Cbz}$), 4.21 (td, $J$ = 8.6, 5.0 Hz, $H_{\alpha,Glu}$), 4.17–4.12 (m, 1H, $H_{\alpha,D-Val}$), 4.07 (td, $J$ = 8.4, 5.2 Hz, 1H, $H_{\alpha,Lys}$), 3.32 (br s, 2H, $H_{\varepsilon,Lys}$, overlap with residual water), 2.27–2.15 (m, 2H, $H_{\gamma,Glu}$), 2.05–1.89 (m, 2H, $H_{\beta,D-Val}$, $H_{\beta,Glu,A}$), 1.78–1.68 (m, 1H, $H_{\beta,Glu,B}$), 1.66–1.15 (m, 28H, $H_{\beta,Lys,}$ $H_{\gamma,Lys}$, $H_{\delta,Lys}$, ($C\underline{H}_2)_{11}CH_3$), 0.88–0.80 (m, 9H, $H_{\gamma,D-Val}$, $CH_2C\underline{H}_3$). $^{13}$C NMR (151 MHz, DMSO) δ 173.8 (COOH), 173.1 ($CO_{Glu}$), 172.4 ($CO_{Lys}$), 170.9 ($CO_{D-Val}$), 156.0 ($CO_{Cbz}$), 137.0 ($C1_{Ar,Cbz}$), 128.3 ($C3_{Ar,Cbz}$, $C5_{Ar,Cbz}$), 127.8 ($C4_{Ar,Cbz}$), 127.7 ($C2_{Ar,Cbz,}$ $C6_{Ar,Cbz}$), 66.4 ($CH_{2,Cbz}$), 57.9 ($C_{\alpha,D-Val}$), 54.8 ($C_{\alpha,Lys}$), 51.6 ($C_{\alpha,Glu}$), 43.4 ($C_{\varepsilon,Lys}$), 31.7 ($C_{\beta,Lys}$), 31.3 ($\underline{C}H_2CH_2CH_3$), 30.1 ($C_{\gamma,Glu}$, $C_{\beta,D-Val}$), 29.0–28.4 (9C,

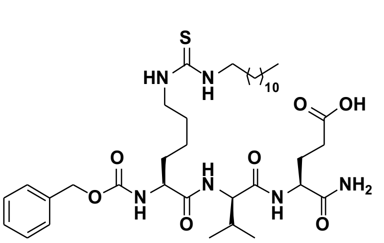

($\underline{C}H_2)_{11}CH_3$), 27.0 ($C_{\beta,Glu}$), 26.4 ($C_{\delta,Lys}$), 22.9 ($C_{\gamma,Lys}$), 22.1 ($\underline{C}H_2CH_3$), 19.2 ($C_{\gamma,D\text{-}Val,1}$), 17.9 ($C_{\gamma,D\text{-}Val,2}$), 13.9 ($CH_2\underline{C}H_3$). The peak for $C_{\varepsilon,Lys}$ was broad and of low intensity and the peak for C=S was not visible in $^{13}C$ NMR due to fast quadrupolar relaxation via the nearby $^{14}N$-nuclei. Analytical HPLC (Method A) gradient 0–95% eluent IV in eluent III (35 min total runtime), $t_R$ 27.5 min (>95%, $UV_{230}$). ESI-MS calcd for $C_{37}H_{63}N_6O_7S^+$ $[M+H]^+$, 735.4; found 735.6.

**(4*S*,7*S*,10*S*)-10-(((benzyloxy)carbonyl)amino)-7-((S)-sec-butyl)-4-carbamoyl-6,9-dioxo-16-thioxo-5,8,15,17-tetraazanonacosanoic acid (29)**

Starting from Cbz-Lys-Ile-Glu(*t*Bu)-resin (115 mg, estimated loading: 0.35 mmol/g) synthesized from Fmoc-Glu(*t*Bu)-OH, Fmoc-Ile-OH and Cbz-Lys(Fmoc)-OH by SPPS, the title compound was synthesized by the on-resin thiourea formation as described in the general procedures using dodecylamine as the amine source. Global deprotection and cleavage from the resin, followed by preparative reverse-phase HPLC purification afforded the desired thiourea (5 mg, 17% based on resin loading) as a colorless fluffy material after lyophilization. $^1H$ NMR (600 MHz, DMSO-$d_6$) δ 12.08 (s, 1H, COOH), 7.87 (d, $J$ = 7.9 Hz, 1H, $NH_{Ile}$), 7.79 (d, $J$ = 8.5 Hz, 1H, $NH_{Glu}$), 7.44 (d, $J$ = 7.8 Hz, 1H, $NH_{\alpha,Lys}$), 7.38–7.22 (m, 8H, $NH_{\varepsilon,Lys}$, $\underline{NH}CH_2$, $H_{Ar,Cbz}$, $CONH_{2,A}$), 7.06 (s, 1H, $CONH_{2,B}$), 5.06–5.01 (m, 2H, $CH_{2,Cbz}$), 4.23–4.14 (m, 2H, $H_{\alpha,Ile}$, $H_{\alpha,Glu}$), 4.00–3.93 (td, $J$ = 8.7, 4.9 Hz, 1H, $H_{\alpha,Lys}$), 3.31 (br s, 2H, $H_{\varepsilon,Lys}$, overlap with residual water), 2.23–2.17 (m, 2H, $H_{\gamma,Glu}$), 1.93–1.85 (m, 1H, $H_{\beta,Glu,A}$), 1.79–1.70 (m, 2H, $H_{\beta,Glu,B}$, $H_{\beta,Ile}$), 1.65–1.58 (m, 1H, $H_{\beta,Lys,A}$), 1.55–1.18 (m, 30H, $H_{\beta,Lys}$ $H_{\gamma,Lys}$, $CH_{\delta,Lys}$, $H_{2,\gamma,Ile}$, $(C\underline{H}_2)_{11}CH_3$), 0.88–0.74 (m, 9H, $H_{3,\gamma,Ile}$, $H_{\delta,Ile}$, $CH_2CH_2C\underline{H}_3$). $^{13}C$ NMR (151 MHz, DMSO) δ 173.8 (COOH), 172.8 ($CO_{Glu}$), 172.0 ($CO_{Lys}$), 170.8 ($CO_{Ile}$), 156.0 ($CO_{Cbz}$), 137.0 ($C1_{Ar,Cbz}$), 128.3 ($C3_{Ar,Cbz}$, $C5_{Ar,Cbz}$), 127.7 ($C4_{Ar,Cbz}$), 127.6 ($C2_{Ar,Cbz,}$ $C6_{Ar,Cbz}$), 66.3 ($CH_{2,Cbz}$), 56.8 ($C_{\alpha,Ile}$), 54.7 ($C_{\alpha,Lys}$), 51.6 ($C_{\alpha,Glu}$), 43.5 ($C_{\varepsilon,Lys}$), 36.5 ($C_{\beta,Ile}$), 31.5 ($C_{\beta,Lys}$), 31.3 ($\underline{C}H_2CH_2CH_3$), 30.1 ($C_{\gamma,Glu}$), 29.0–28.5 (9C, ($\underline{C}H_2)_{11}CH_3$), 27.4 ($C_{\beta,Glu}$), 26.4 ($C_{\delta,Lys}$), 24.2 ($CH_{2,\gamma,Ile}$), 23.0 ($C_{\gamma,Lys}$), 22.1 ($CH_2\underline{C}H_2CH_3$), 15.3 ($CH_{3,\gamma,Ile}$), 13.9 ($CH_2CH_2\underline{C}H_3$), 11.0 ($C_{\delta,Ile}$). The peak for $C_{\varepsilon,Lys}$ was broad and of low intensity and the peak for C=S was not visible in $^{13}C$ NMR due to fast quadrupolar relaxation via the nearby $^{14}N$-nuclei. Analytical HPLC (Method A) gradient 0–95% eluent IV in eluent III (35 min total runtime), $t_R$ 28.0 min (>98%, $UV_{230}$). ESI-MS calcd for $C_{38}H_{65}N_6O_7S^+$ $[M+H]^+$, 749.5; found 749.5.

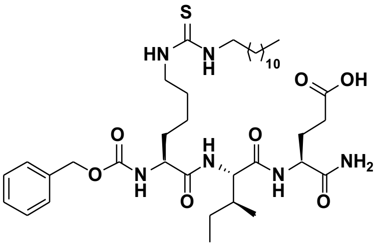

**(4*S*,7*S*,10*S*)-10-acetamido-4-carbamoyl-7-isopropyl-6,9-dioxo-16-thioxo-5,8,15,17-tetraazanonacosanoic acid (30)**

Starting from H-Lys(Alloc)-Val-Glu-resin (115 mg, estimated loading: 0.35 mmol/g) synthesized from Fmoc-Glu(*t*Bu)-OH, Fmoc-Val-OH and Fmoc-Lys(Alloc)-OH by SPPS, the title compound was synthesized by capping the resin with a solution of $Ac_2O$:DMF (1:4, 2.0 mL) for 1 h at ambient temperature. The resin was drained and washed with DMF (3×4.0 mL) and $CH_2Cl_2$ (3×4.0 mL) followed by Alloc-deprotection and on-resin thiourea formation as described in the general procedures using dodecylamine as the amine source. Global deprotection and cleavage from the resin, followed by preparative reverse-phase HPLC purification afforded the desired thiourea (2 mg, 9% based on resin loading) as a colorless fluffy material after lyophilization. $^1H$ NMR (600 MHz, DMSO-$d_6$) δ 12.09 (br s, 1H, COOH), 8.02 (d, $J$ = 7.9 Hz, 1H, $NH_{\alpha,Lys}$), 7.84 (d, $J$ = 7.8 Hz, 1H, $NH_{Glu}$), 7.79 (d, $J$ = 8.5 Hz, 1H, $NH_{Val}$), 7.39–7.22 (m, 3H, $NH_{\varepsilon,Lys}$, $\underline{NH}CH_2$, $CONH_{2,A}$), 7.06 (s, 1H, $CONH_{2,B}$), 4.26 (td, J = 8.5, 5.2 Hz, 1H, $H_{\alpha,Lys}$), 4.19 (td, $J$ = 8.1, 5.2 Hz, 1H, $H_{\alpha,Glu}$), 4.14–4.08 (m, 1H, $H_{\alpha,Val}$), 3.32 (br s, 2H, $H_{\varepsilon,Lys}$, overlap with residual water), 2.20 (t, $J$ = 9.3, 6.8 Hz, 2H, $H_{\gamma,Glu}$), 1.99 (h, $J$ = 6.7 Hz, 1H, $H_{\beta,Val}$), 1.92–1.85 (m, 1H, $H_{\beta,Glu,A}$), 1.84 (s, 3H, $CH_3CO$), 1.79–1.70 (m, 1H, $H_{\beta,Glu,B}$), 1.68–1.59 (m, 1H, $H_{\beta,Lys,A}$), 1.52–1.40 (m, 5H, $H_{\beta,Lys,B}$, $H_{\delta,Lys}$, $NHC\underline{H}_2$), 1.33–1.11 (m, 22H, $H_{\gamma,Lys}$, $(C\underline{H}_2)_{10}CH_3$), 0.90–0.77 (m, 9H, $H_{\gamma,Val}$, $CH_2C\underline{H}_3$). $^{13}C$ NMR (151 MHz, DMSO) δ 173.9 (COOH), 172.9 ($CO_{Glu}$), 172.0 ($CO_{Lys}$), 170.7 ($CO_{Val}$), 169.3 ($CH_3\underline{C}O$), 57.7 ($C_{\alpha,Val}$), 52.6 ($C_{\alpha,Lys}$), 51.7 ($C_{\alpha,Glu}$), 43.4 ($C_{\varepsilon,Lys}$), 31.5 ($C_{\beta,Lys}$), 31.3 ($\underline{C}H_2CH_2CH_3$),

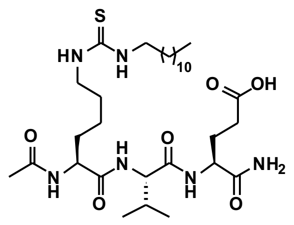

30.2 ($C_{\beta,Val}$), 30.1 ($C_{\gamma,Glu}$), 29.0–28.6 (C9, (<u>C</u>$H_2$)$_{11}$$CH_3$), 27.4 ($C_{\beta,Glu}$), 26.4 ($C_{\delta,Lys}$), 22.9 ($C_{\gamma,Lys}$), 22.4 (<u>C</u>$H_3$CO), 22.1 (<u>C</u>$H_2CH_3$), 19.2 ($C_{\gamma,Val,1}$), 18.0 ($C_{\gamma,Val,2}$), 13.9 ($CH_2$<u>C</u>$H_3$). The peak for $C_{\varepsilon,Lys}$ was broad and of low intensity and the peak for C=S was not visible in $^{13}$C NMR due to fast quadrupolar relaxation via the nearby $^{14}$N-nuclei. Analytical HPLC (Method A) gradient 0–95% eluent IV in eluent III (35 min total runtime), $t_R$ 24.9 min (>97%, $UV_{230}$). ESI-MS calcd for $C_{31}H_{59}N_6O_6S^+$ $[M+H]^+$, 643.4; found 643.6.

**(4*S*,7*S*,10*S*)-10-benzamido-4-carbamoyl-7-isopropyl-6,9-dioxo-16-thioxo-5,8,15,17-tetraazanonacosanoic acid (31)**

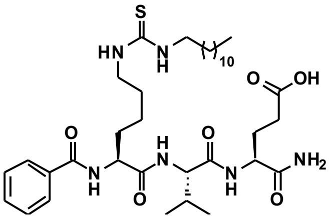

Starting from H-Lys(Teoc)-Val-Glu(*t*Bu)-resin (85 mg, estimated loading: 0.47 mmol/g) synthesized from Fmoc-Glu(*t*Bu)-OH, Fmoc-Val-OH and Fmoc-Lys(Teic)-OH by SPPS, the title compound was synthesized by addition of benzoyl chloride (14 µL, 0.12 mmol) and *i*$Pr_2$NEt (42 µL, 0.24 mmol) in $CH_2Cl_2$ (2.0 mL) to the resin, which was agitated for overnight at ambient temperature. The resin was drained and washed with $CH_2Cl_2$ (3×4.0 mL) and DMF (3×4.0 mL) followed by Teoc deprotection and on-resin thiourea formation as described in the general procedures using dodecylamine as the amine source. Global deprotection and cleavage from the resin, followed by preparative reverse-phase HPLC purification afforded the desired thiourea (7 mg, 25% based on resin loading) as a colorless fluffy material after lyophilization. $^1$H NMR (600 MHz, DMSO-*d*$_6$) δ 12.09 (s, 1H, COOH), 8.48 (d, $J$ = 7.9 Hz, 1H, $NH_{\alpha,Lys}$), 7.93–7.86 (m, 3H, $NH_{Glu}$, $C2_{Ar}$, $C6_{Ar}$), 7.82 (d, $J$ = 8.6 Hz, 1H, $NH_{Val}$), 7.57–7.51 (m, 1H, $H4_{Ar}$), 7.49–7.45 (m, 2H, $H3_{Ar}$, $H5_{Ar}$), 7.37–7.21 (m, 3H, $NH_{\varepsilon,Lys}$, <u>NH</u>$CH_2$, $CONH_{2,A}$), 7.05 (d, $J$ = 2.1 Hz, 1H, $CONH_{2,B}$), 4.50–4.42 (m, 1H, $H_{\alpha,Lys}$), 4.24–4.13 (m, 2H, $H_{\alpha,Val}$, $H_{\alpha,Glu}$), 3.32 (br s, 2H, $H_{\varepsilon,Lys}$, overlap with residual water), 2.27–2.16 (m, 2H, $H_{\gamma,Glu}$), 2.09–1.96 (m, 1H, $H_{\beta,Val}$), 1.95–1.86 (m, 1H, $H_{\beta,Glu,A}$), 1.79–1.67 (m, 3H, $H_{\beta,Glu,B}$, $H_{\beta,Lys}$), 1.56–1.17 (m, 26H, $H_{\gamma,Lys}$, $H_{\delta,Lys}$, (C<u>H</u>$_2$)$_{11}$$CH_3$), 0.88–0.80 (m, 9H, $H_{\gamma,Val}$, $CH_2$C<u>H</u>$_3$). $^{13}$C NMR (151 MHz, DMSO) δ 173.9 (COOH), 172.9 ($CO_{Glu}$), 172.0 ($CO_{Lys}$), 170.7 ($CO_{Val}$), 166.6 ($CO_{Ar}$), 134.2 ($C1_{Ar}$), 131.3 ($C4_{Ar}$), 128.2 ($C3_{Ar}$, $C5_{Ar}$), 127.5 ($C2_{Ar}$, $C6_{Ar}$), 57.6 ($C_{\alpha,Val}$), 53.6 ($C_{\alpha,Lys}$), 51.7 ($C_{\alpha,Glu}$), 43.5 ($C_{\varepsilon,Lys}$), 31.3 ($C_{\beta,Lys}$), 31.1 (<u>C</u>$H_2CH_2CH_3$), 30.4 ($C_{\beta,Val}$), 30.1 ($C_{\gamma,Glu}$), 29.0–28.7 (9C, (<u>C</u>$H_2$)$_{11}$$CH_3$), 27.4 ($C_{\beta,Glu}$), 26.4 ($C_{\delta,Lys}$), 23.3 ($C_{\gamma,Lys}$), 22.1 (<u>C</u>$H_2CH_3$), 19.2 ($C_{\gamma,Val,1}$), 17.9 ($C_{\gamma,Val,2}$), 13.9 ($CH_2$<u>C</u>$H_3$). The peak for $C_{\varepsilon,Lys}$ was broad and of low intensity and the peak for C=S was not visible in $^{13}$C NMR due to fast quadrupolar relaxation via the nearby $^{14}$N-nuclei. Analytical HPLC (Method A) gradient 0–95% eluent IV in eluent III (35 min total runtime), $t_R$ 27.4 min (>98%, $UV_{230}$). ESI-MS calcd for $C_{36}H_{61}N_6O_6S^+$ $[M+H]^+$, 705.4; found 705.4.

**(4*S*,7*S*,10*S*)-4-carbamoyl-10-((3-carboxyphenyl)sulfonamido)-7-isopropyl-6,9-dioxo-16-thioxo-5,8,15,17-tetraazanonacosanoic acid (32)**

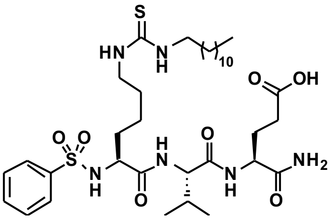

Starting from H-Lys(Teoc)-Val-Glu(*t*Bu)-resin (85 mg, estimated loading: 0.47 mmol/g) synthesized from Fmoc-Glu(*t*Bu)-OH, Fmoc-Val-OH and Fmoc-Lys(Teoc)-OH by SPPS, the title compound was synthesized by addition of benzenesulfonyl chloride (15 µL, 0.12 mmol) and *i*$Pr_2$NEt (42 µL, 0.24 mmol) in $CH_2Cl_2$ (4 mL) to the resin followed by overnight agitation at ambient temperature. The resin was drained and washed with $CH_2Cl_2$ (3×4.0 mL) and DMF (3×4.0 mL) followed by Teoc deprotection and on-resin thiourea formation as described in the general procedures using dodecylamine as the amine source. Global deprotection and cleavage from the resin, followed by preparative reverse-phase HPLC purification afforded the desired thiourea (6 mg, 20% based on resin loading) as a colorless fluffy material after lyophilization. $^1$H NMR (600 MHz, DMSO-*d*$_6$) δ 12.07 (br s, 1H, COOH), 7.94 (d, $J$ = 8.8 Hz, 1H, $NH_{\alpha,Lys}$), 7.88 (d, $J$ = 8.4 Hz, 1H, $NH_{Glu}$), 7.80 (d, $J$ = 7.8 Hz, 1H, $NH_{Val}$), 7.79–7.73 (m, 2H, $C2_{Ar}$, $C6_{Ar}$), 7.61–7.55 (m, 1H, $H4_{Ar}$), 7.55–7.48 (m, 2H, $H3_{Ar}$, $H5_{Ar}$), 7.32–7.18 (m, 3H, $NH_{\varepsilon,Lys}$, <u>NH</u>$CH_2$, $CONH_{2,A}$), 7.04 (d, $J$ = 2.1 Hz, 1H, $CONH_{2,B}$), 4.17 (td, $J$ = 8.1, 5.3 Hz, 1H, $H_{\alpha,Glu}$),

3.93 (dd, *J* = 8.4, 6.5 Hz, 1H, $H_{\alpha,Val}$), 3.85–3.76 (m, 1H, $H_{\alpha,Lys}$), 3.32 (br s, 2H, $H_{\varepsilon,Lys}$, overlap with residual water), 1.91–1.82 (m, 1H, $H_{\beta,Val}$, $H_{\beta,Glu,A}$), 1.78–1.68 (m, 1H, $H_{\beta,Glu,B}$), 1.56–1.17 (m, 26H, $H_{\gamma,Lys}$, $H_{\delta,Lys}$, $(C\underline{H}_2)_{11}CH_3$), 0.85 (t, *J* = 6.9 Hz, 3H, $CH_2C\underline{H}_3$), 0.78–0.68 (m, 6H, $H_{\gamma,Val,}$). $^{13}C$ NMR (151 MHz, DMSO) δ 173.9 (COOH), 172.8 ($CO_{Glu}$), 170.8 ($CO_{Lys}$), 170.5 ($CO_{Val}$), 141.1 ($C1_{Ar}$), 132.1 ($C4_{Ar}$), 128.8 ($C3_{Ar}$, $C5_{Ar}$), 126.4 ($C2_{Ar}$, $C6_{Ar}$), 57.6 ($C_{\alpha,Val}$), 55.9 ($C_{\alpha,Lys}$), 51.6 ($C_{\alpha,Glu}$), 43.4 ($C_{\varepsilon,Lys}$), 32.7 ($C_{\beta,Lys}$), 31.3 ($\underline{C}H_2CH_2CH_3$), 30.2 ($C_{\beta,Val}$), 30.0 ($C_{\gamma,Glu}$), 29.0–28.3 (9C, $(\underline{C}H_2)_{11}CH_3$), 27.3 ($C_{\beta,Glu}$), 26.4 ($C_{\delta,Lys}$), 22.5 ($C_{\gamma,Lys}$), 22.1 ($\underline{C}H_2CH_3$), 19.1 ($C_{\gamma,Val,1}$), 18.0 ($C_{\gamma,Val,2}$), 13.9 ($CH_2\underline{C}H_3$). The peak for $C_{\varepsilon,Lys}$ was broad and of low intensity and the peak for C=S was not visible in $^{13}C$ NMR due to fast quadrupolar relaxation via the nearby $^{14}N$-nuclei. Analytical HPLC (Method A) gradient 0–95% eluent IV in eluent III (35 min total runtime), $t_R$ 27.3 min (>98%, $UV_{230}$). ESI-MS calcd for $C_{35}H_{61}N_6O_7S_2^+$ $[M+H]^+$, 741.4; found 741.4

**(5*S*,8*S*,11*S*)-11-carbamoyl-5-(6-(dodecylamino)-6-oxohexyl)-8-isopropyl-3,6,9-trioxo-1-phenyl-2-oxa-4,7,10-triazatetradecan-14-oic acid (33)**

Starting from H-Asu(myr)-Val-Glu(*t*Bu)-resin (85 mg, estimated loading: 0.47 mmol/g) synthesized from Fmoc-Glu(*t*Bu)-OH, Fmoc- Val-OH and Fmoc-Asu(myr)-OH (**S13**) by SPPS, the title compound was synthesized by addition of benzyl chloroformate (17 µL, 0.12 mmol) and *i*$Pr_2NEt$ (42 µL, 0.24 mmol) in $CH_2Cl_2$ (2.0 mL) the resin followed by overnight agitation at ambient temperature. The resin was drained and washed with $CH_2Cl_2$ (3×4.0 mL) and DMF (3×4.0 mL) followed by followed by global deprotection and cleavage from the resin. Preparative reverse-phase HPLC purification afforded the desired inverse amide (4 mg, 14% based on resin loading) as a colorless fluffy material after lyophilization. $^1H$ NMR (600 MHz, DMSO-$d_6$) δ 12.08 (s, 1H, COOH), 7.88 (d, *J* = 7.8 Hz, 1H, $NH_{Glu}$), 7.73 (d, *J* = 8.7 Hz, 1H, $NH_{Val}$), 7.68 (t, *J* = 5.6 Hz, 1H, $N\underline{H}CH_2$), 7.45 (d, *J* = 8.2 Hz, 1H, $NH_{Asu}$), 7.38–7.22 (m, 6H, $H_{Ar,Cbz}$, $CONH_{2,A}$), 7.04 (br s, 1H, $CONH_{2,B}$), 5.02 (s, 2H, $CH_{2,Cbz}$), 4.23–4.14 (m, 2H, $H_{\alpha,Glu}$, $H_{\alpha,Val}$), 4.01 (td, *J* = 8.7, 5.0 Hz, 1H, $H_{\alpha,Asu}$), 2.99 (q, *J* = 6.6 Hz, 2H,$NHC\underline{H}_2$), 2.25–2.16 (m, 2H, $H_{\gamma,Glu}$), 2.05–1.94 (m, 3H, $H_{\zeta,Asu}$, $H_{\beta,Val}$), 1.93–1.84 (m, 1H, $H_{\beta,Glu,A}$), 1.79–1.70 (m, 1H, $H_{\beta,Glu,B}$), 1.64–1.54 (m, 1H, $H_{\beta,Asu,A}$), 1.53–1.16 (m, 27H, $H_{\beta,Asu,B}$, $H_{\gamma,Asu}$, $H_{\delta,Asu}$, $H_{\varepsilon,Asu}$, $(C\underline{H}_2)_{10}CH_3$), 0.89–0.75 (m, 9H, $H_{\gamma,Val}$, $CH_2C\underline{H}_3$). $^{13}C$ NMR (151 MHz, DMSO) δ 173.8 (COOH), 172.8 ($CO_{Glu}$), 172.0 ($CO_{\alpha,Asu}$), 171.9 ($CO_{\zeta,Asu}$), 170.7 ($CO_{Val}$), 155.9 ($CO_{Cbz}$), 137.0 ($C1_{Ar,Cbz}$), 128.3 ($C3_{Ar,Cbz,}$ $C5_{Ar,Cbz}$), 127.7 ($C4_{Ar,Cbz}$), 127.6 ($C2_{Ar,Cbz,}$ $C6_{Ar,Cbz}$), 65.3 ($CH_{2,Cbz}$), 57.5 ($C_{\alpha,Val}$), 54.8 ($C_{\alpha,Asu}$), 51.7 ($C_{\alpha,Glu}$), 38.4 ($NHC\underline{H}_2$), 35.4 ($C_{\zeta,Asu}$), 31.7 ($C_{\beta,Asu}$), 31.3 ($\underline{C}H_2CH_2CH_3$), 30.5 ($C_{\beta,Val}$), 30.1 ($C_{\gamma,Glu}$), 29.1–28.4 (9C, $C_{\delta,Asu}$, $(C\underline{H}_2)_9CH_3$), 27.4 ($C_{\beta,Glu}$), 26.4 ($NHCH_2\underline{C}H_2$), 25.2 ($C_{\gamma,Asu}$, $C_{\varepsilon,Asu}$), 22.1 ($\underline{C}H_2CH_3$), 19.2 ($C_{\gamma,Val,1}$), 18.0 ($C_{\gamma,Val,2}$), 13.9 ($CH_2\underline{C}H_3$). Analytical HPLC (Method A) gradient 0–95% eluent IV in eluent III (35 min total runtime), $t_R$ 27.8 min (>97%, $UV_{210}$). HRMS calcd for $C_{38}H_{64}N_5O_8^+$ $[M+H]^+$, 718.4749; found 718.4733.

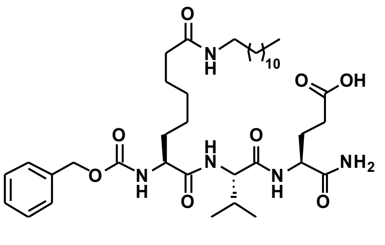

**(5*S*,8*S*,11*S*)-11-carbamoyl-5-(6-(dodecylamino)-6-thioxohexyl)-8-isopropyl-3,6,9-trioxo-1-phenyl-2-oxa-4,7,10-triazatetradecan-14-oic acid (34)**

Starting from H-Asu(thiomyr)-Val-Glu(*t*Bu)-resin (85 mg, estimated loading: 0.47 mmol/g) synthesized from Fmoc-Glu(*t*Bu)-OH, Fmoc- Val-OH and Fmoc-Asu(thiomyr)-OH (**S14**) by SPPS the title compound was synthesized by addition of benzyl chloroformate (17 µL, 0.12 mmol) and *i*$Pr_2NEt$ (42 µL, 0.24 mmol) in $CH_2Cl_2$ (2.0 mL) the resin followed by overnight agitation at ambient temperature. The resin was drained and washed with $CH_2Cl_2$ (3×4.0 mL) and DMF (3×4.0 mL) followed by global deprotection and cleavage from the resin. Preparative reverse-phase HPLC purification afforded the desired inverse thioamide (4 mg, 13% based on resin loading) as a colorless fluffy material after lyophilization. $^1H$ NMR (600 MHz, DMSO-$d_6$) δ 12.09 (s, 1H, COOH), 9.84 (t, *J* = 5.4 Hz, 1H, $N\underline{H}CH_2$), 7.90 (d, *J* = 7.8 Hz, 1H, $NH_{Glu}$), 7.71 (d, *J* = 8.7 Hz, 1H, $NH_{Val}$), 7.47 (d, *J* = 8.1 Hz, 1H, $NH_{Asu}$), 7.40–7.23 (m, 6H, $H_{Ar,Cbz}$, $CONH_{2,A}$), 7.08–7.00 (m, 1H, $CONH_{2,B}$), 5.02

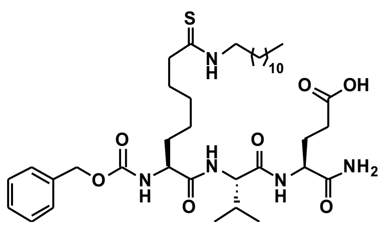

(s, 2H, $CH_{2,Cbz}$), 4.22–4.14 (m, 2H, $H_{\alpha,Glu}$, $H_{\alpha,Val}$), 4.01 (td, *J* = 8.7, 4.9 Hz, 1H, $H_{\alpha,Asu}$), 3.45 (td, *J* = 7.2, 5.3 Hz, 2H, $NHC\underline{H}_2$), 2.49–2.47 (m, 1H, $H_{\zeta,Asu}$, overlap with solvent peak), 2.24–2.17 (m, 2H, $H_{\gamma,Glu}$), 1.98 (hept, *J* = 6.7 Hz, 1H, $H_{\beta,Val}$), 1.93–1.84 (m, 1H, $H_{\beta,Glu,A}$), 1.79–1.71 (m, 1H, $H_{\beta,Glu,B}$), 1.66–1.46 (m, 6H, $C_{\beta,Asu}$, $C_{\varepsilon,Asu}$, $(C\underline{H}_2)_{10}CH_3$), 1.38–1.18 (m, 22H, $H_{\gamma,Asu}$, $H_{\delta,Asu}$, $(C\underline{H}_2)_{10}CH_3$), 0.90–0.71 (m, 9H, $H_{\gamma,Val}$, $CH_2C\underline{H}_3$). $^{13}C$ NMR (151 MHz, DMSO) δ 203.5 (C=S), 173.9 (COOH), 172.8 ($CO_{Glu}$), 172.0 ($CO_{\alpha,Asu}$), 170.7 ($CO_{Val}$), 155.9 ($CO_{Cbz}$), 137.0 ($C1_{Ar,Cbz}$), 128.3 ($C3_{Ar,Cbz,}$ $C5_{Ar,Cbz}$), 127.7 ($C4_{Ar,Cbz}$), 127.6 ($C2_{Ar,Cbz,}$ $C6_{Ar,Cbz}$), 65.3 ($CH_{2,Cbz}$), 57.5 ($C_{\alpha,Val}$), 54.8 ($C_{\alpha,Asu}$), 51.7 ($C_{\alpha,Glu}$), 45.1 ($C_{\zeta,Asu}$/$NHC\underline{H}_2$), 45.0 ($C_{\zeta,Asu}$/$NHC\underline{H}_2$), 31.7 ($C_{\beta,Asu}$), 31.3 ($\underline{C}H_2CH_2CH_3$), 30.5 ($C_{\beta,Val}$), 30.1 ($C_{\gamma,Glu}$), 29.0–28.7 (8C, $(C\underline{H}_2)_9CH_3$), 27.9 ($C_{\delta,Asu}$), 27.4 ($C_{\beta,Glu}$), 27.1 ($C_{\varepsilon,Asu}$), 26.4 ($NHCH_2\underline{C}H_2$), 25.2 ($C_{\gamma,Asu}$), 22.1 ($\underline{C}H_2CH_3$), 19.2 ($C_{\gamma,Val,1}$), 17.9 ($C_{\gamma,Val,2}$), 13.9 ($CH_2\underline{C}H_3$). Analytical HPLC (Method A) gradient 0–95% eluent IV in eluent III (35 min total runtime), $t_R$ 29.3 min (>98%, $UV_{230}$). HRMS calcd for $C_{38}H_{64}N_5O_7S^+$ $[M+H]^+$, 734.4521; found 734.4518.

**Synthesis of RaPID-derived hit peptide macrocycles**

**General procedure for synthesis of hit peptides from RaPID**

The desired peptide was synthesized using standard Fmoc-based SPPS with the Rink amide linker on TentaGel® resin (87 mg, estimated loading: 0.23 mmol/g). The N-terminal amino group was then functionalized with a chloroacetyl group, by reacting the free amine with a solution of chloroacetic acid (6 µL, 0.10 mmol), DIC (16 µL 0.10 mmol), and HOAt (6 µL, 0.04 mmol) in DMF (2.0 mL) for 1 h at ambient temperature with agitation. The resin was drained and washed with DMF (3×2.0 mL), $CH_3OH$ (3×2.0 mL), and $CH_2Cl_2$ (3×2.0 mL) followed by global deprotection and cleavage from the resin. The resulting pellet was dissolved in DMSO (2.0 mL) and *i*$Pr_2NEt$ (8 drops) was added. The reaction was left at ambient temperature overnight with agitation. The solution was then acidified with TFA and diluted with MeCN–$H_2O$ (8 mL, 1:1; v/v) or neat MeCN. Preparative reverse-phase HPLC purification was used to purify the macrocycle and fractions containing pure compound were pooled and lyophilized.

**Characterization data for peptide 35**

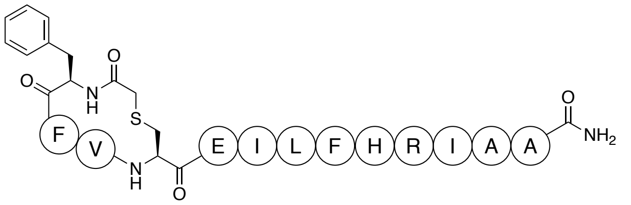

The peptide was synthesized at 20 µmol scale according to the general procedure. Preparative reverse-phase HPLC purification on the C18 column afforded the macrocycle (7 mg, 21% based on resin loading) as a colorless fluffy material after lyophilization. Analytical HPLC (Method B) gradient 0% eluent IV for 1 min followed by 0–95% eluent IV in eluent III over 10 minutes, $t_R$ = 7.22 min (>96%, $UV_{215}$). HRMS calcd for $C_{78}H_{114}N_{19}O_{16}S^+$, $[M+H]^+$, 1604.8405; found, 1604.8408.

**Characterization data for peptide 36**

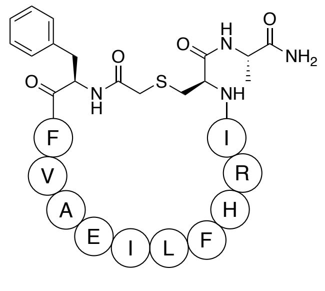

The peptide was synthesized at 40 µmol scale according to the general procedure. Preparative reverse-phase HPLC purification on the C8 column afforded the macrocycle (1.2 mg, 1.8% based on resin loading) as a colorless material after lyophilization. Analytical HPLC (Method B) gradient 0% eluent IV for 1 min followed by 0–95% eluent IV in eluent III over 10 minutes, $t_R$ = 6.88 min (>98%, $UV_{215}$), HRMS calcd for $C_{78}H_{114}N_{19}O_{16}S^+$, $[M+H]^+$, 1604.8406; found, 1604.8434.

**Characterization data for peptide 37**

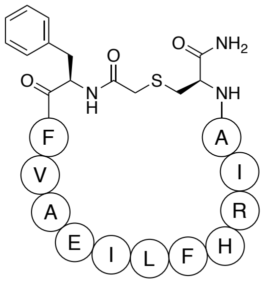

The peptide was synthesized at 40 µmol scale according to the general procedure. Preparative reverse-phase HPLC purification on the C8 column afforded the macrocycle (0.6 mg, 0.9% based on resin loading) as a colorless material after lyophilization. Analytical HPLC (Method B) gradient 0% eluent IV for 1 min followed by 0–95% eluent IV in eluent III over 10 minutes, $t_R$ = 6.96 min (97%, $UV_{215}$), HRMS calcd for $C_{78}H_{114}N_{19}O_{16}S^+$, $[M+H]^+$, 1604.8406; found, 1604.8418.

**Characterization data for peptide 38**

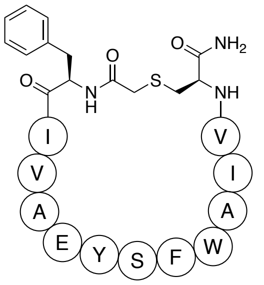

The peptide was synthesized at 40 µmol scale according to the general procedure. Preparative reverse-phase HPLC purification on the C8 column afforded the macrocycle (0.4 mg, 0.6% based on resin loading) as a colorless material after lyophilization. Analytical HPLC (Method B) gradient 0% eluent IV for 1 min followed by 0–95% eluent IV in eluent III over 10 minutes, $t_R$ = 7.15 min (97%, $UV_{215}$), HRMS calcd for $C_{78}H_{114}N_{19}O_{16}S^+$, $[M+H]^+$, 1586.7712; found, 1586.7757.

**Characterization data for peptide 39**

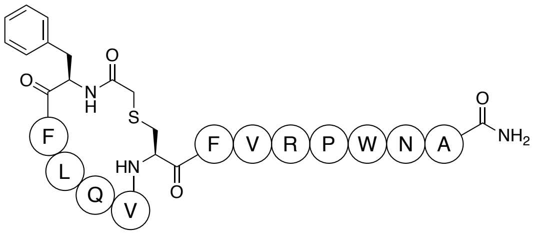

The peptide was synthesized at 20 µmol scale according to the general procedure. Preparative reverse-phase HPLC purification on the C18 column afforded the macrocycle (3 mg, 9% based on resin loading) as a colorless material after lyophilization. Analytical HPLC (Method B) gradient 0% eluent IV for 1 min followed by 0–95% eluent IV in eluent III over 10 minutes, $t_R$ = 6.66 min (>96%, $UV_{215}$), HRMS calcd for $C_{82}H_{113}N_{20}O_{16}S^+$, $[M+H]^+$, 1665.8357; found, 1665.8356.

**Characterization data for peptide 40**

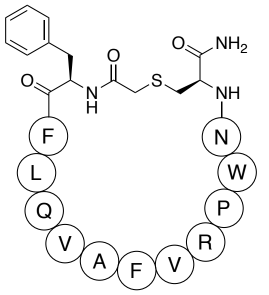

The peptide was synthesized at 40 µmol scale according to the general procedure. Preparative reverse-phase HPLC purification on the C8 column afforded the macrocycle (5 mg, 7.4% based on resin loading) as a colorless material after lyophilization. Analytical HPLC (Method B) gradient 0% eluent IV for 1 min followed by 0–95% eluent IV in eluent III over 10 minutes, $t_R$ = 6.73 min (96%, $UV_{215}$), HRMS calcd for $C_{82}H_{113}N_{20}O_{16}S^+$, $[M+H]^+$, 1665.8359; found, 1665.8371.

**Characterization data for peptide 41**

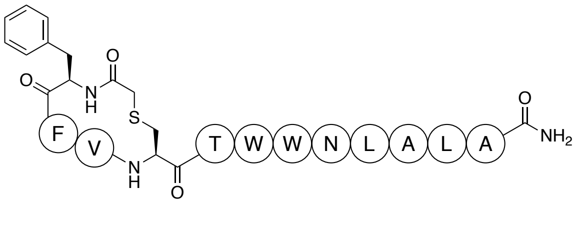

The peptide was synthesized at 40 µmol scale according to the general procedure. Preparative reverse-phase HPLC purification on the C18 column afforded the macrocycle (17.4 mg, 29% based on resin loading) as a colorless material after lyophilization. Analytical HPLC (Method B) gradient

0% eluent IV for 1 min followed by 0–95% eluent IV in eluent III over 10 minutes, $t_R$ = 8.07 min (>98%, $UV_{215}$), HRMS calcd for $C_{76}H_{101}N_{16}O_{15}S^+$, $[M+H]^+$, 1509.7348; found, 1509.7381.

**Characterization data for peptide 42**

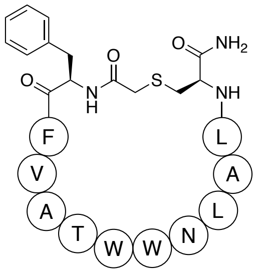

The peptide was synthesized at 40 µmol scale according to the general procedure. Preparative reverse-phase HPLC purification on the C18 column afforded the macrocycle (3.1 mg, 5.1% based on resin loading) as a colorless material after lyophilization. Analytical HPLC (Method B) gradient 0% eluent IV for 1 min followed by 0–95% eluent IV in eluent III over 10 minutes, $t_R$ = 7.34 min (95%, $UV_{215}$), HRMS calcd for $C_{76}H_{101}N_{16}O_{15}S^+$, $[M+H]^+$, 1509.7348; found, 1509.7357.

**Characterization data for peptide 43**

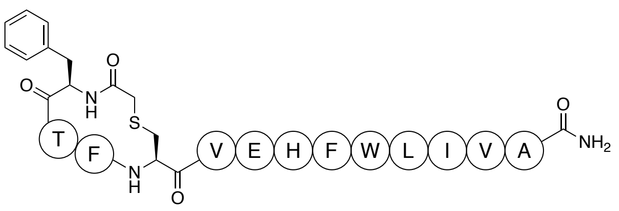

The peptide was synthesized at 40 µmol scale according to the general procedure. Preparative reverse-phase HPLC purification on the C18 column afforded the macrocycle (8.7 mg, 13.2 % based on resin loading) as a colorless material after lyophilization. Analytical HPLC (Method B) gradient 0% eluent IV for 1 min followed by 0–95% eluent IV in eluent III over 10 minutes, $t_R$ = 7.42 min (97%, $UV_{215}$), HRMS calcd for $C_{83}H_{112}N_{17}O_{17}S^+$, $[M+H]^+$, 1650.813; found, 1650.8189.

**Characterization data for peptide 44**

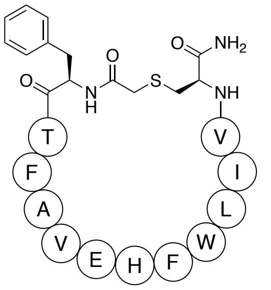

The peptide was synthesized at 40 µmol scale according to the general procedure. Preparative reverse-phase HPLC purification on the C8 column afforded the macrocycle (1.8 mg, 2.7% based on resin loading) as a colorless material after lyophilization. Analytical HPLC (Method B) gradient 0% eluent IV for 1 min followed by 0–95% eluent IV in eluent III over 10 minutes, $t_R$ = 6.95 min (95%, $UV_{215}$), HRMS calcd for $C_{83}H_{112}N_{17}O_{17}S^+$, $[M+H]^+$, 1650.8137; found, 1650.8178.

**Characterization data for peptide 45**

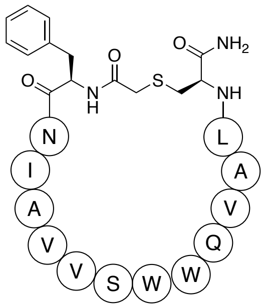

The peptide was synthesized at 40 µmol scale according to the general procedure. Preparative reverse-phase HPLC purification on the C8 column afforded the macrocycle (0.3 mg, 0.5% based on resin loading) as a colorless material after lyophilization. Analytical HPLC (Method B) gradient 0% eluent IV for 1 min followed by 0–95% eluent IV in eluent III over 10 minutes, $t_R$ = 6.62 min (97%, $UV_{215}$), HRMS calcd for $C_{81}H_{116}N_{19}O_{18}S^+$, $[M+H]^+$, 1674.8461; found, 1674.8484.

**Characterization data for peptide 46**

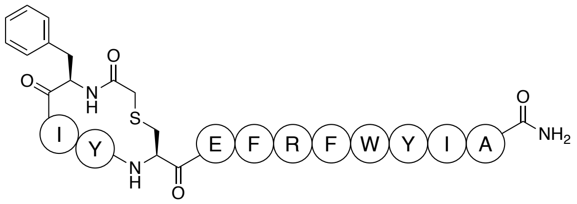

The peptide was synthesized at 40 μmol scale according to the general procedure. Preparative reverse-phase HPLC purification on the C18 column afforded the macrocycle (14.2 mg, 21% based on resin loading) as a colorless material after lyophilization. Analytical HPLC (Method B) gradient 0% eluent IV for 1 min followed by 0–95% eluent IV in eluent III over 10 minutes, $t_R$ = 6.97 min (95%, $UV_{215}$), HRMS calcd for $C_{87}H_{110}N_{17}O_{17}S^+$, $[M+H]^+$, 1696.7981; found, 1696.8116.

**Characterization data for peptide 47**

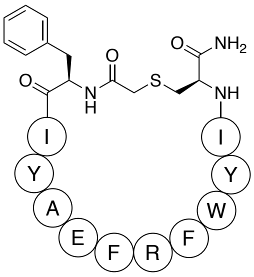

The peptide was synthesized at 40 μmol scale according to the general procedure. Preparative reverse-phase HPLC purification on the C8 column afforded the macrocycle (0.4 mg, 0.5% based on resin loading) as a colorless material after lyophilization. Analytical HPLC (Method B) gradient 0% eluent IV for 1 min followed by 0–95% eluent IV in eluent III over 10 minutes, $t_R$ = 6.75 min (96%, $UV_{215}$), HRMS calcd for $C_{87}H_{110}N_{17}O_{17}S^+$, $[M+H]^+$, 1696.7981; found, 1696.8066.

### Copies of analytical HPLC chromatograms

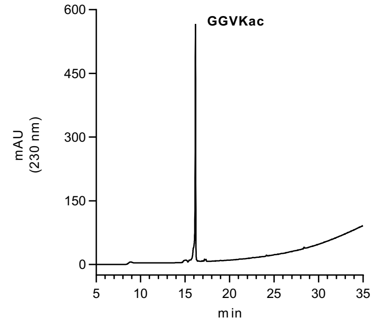

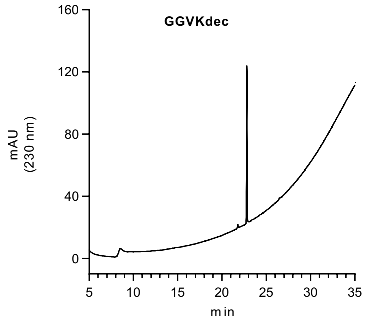

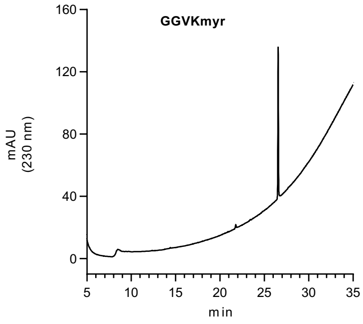

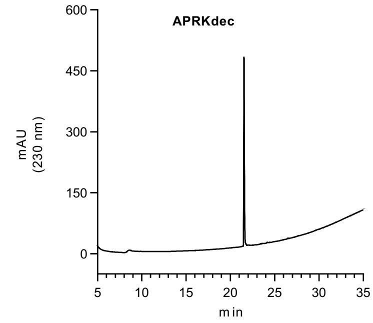

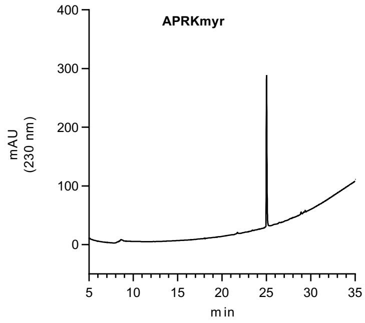

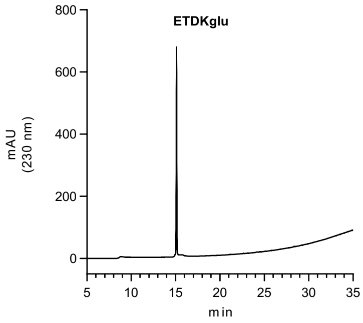

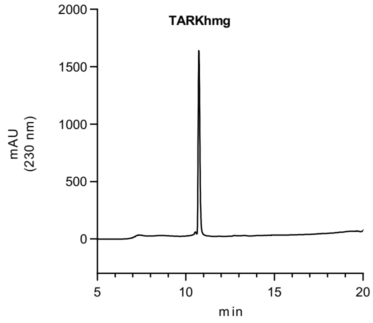

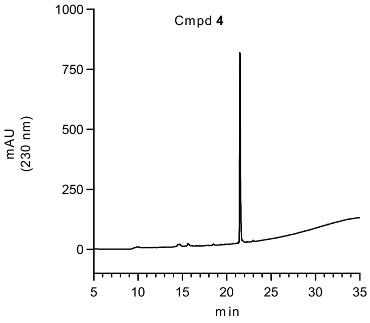

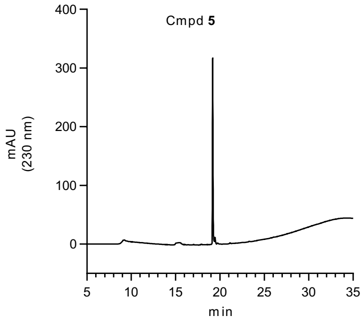

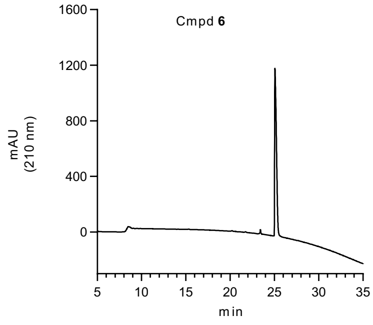

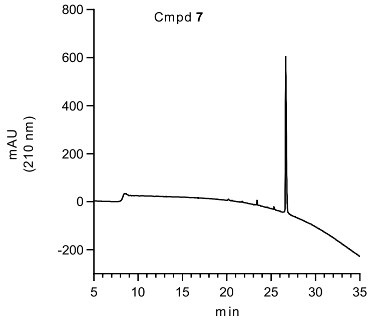

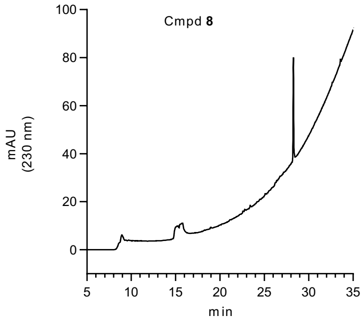

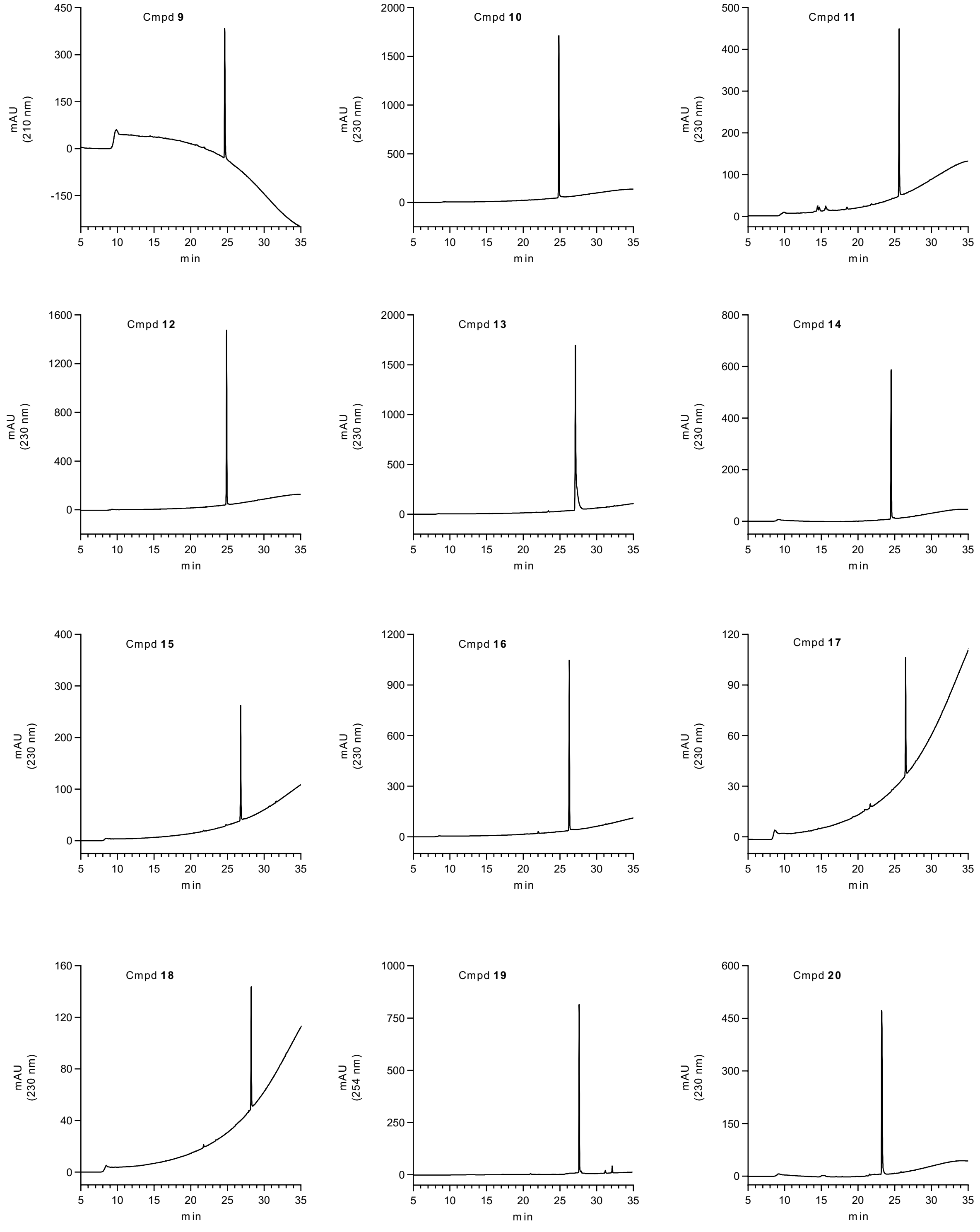
Cmpd **9**
mAU (210 nm)
min
Cmpd **10**
mAU (230 nm)
min
Cmpd **11**
mAU (230 nm)
min
Cmpd **12**
mAU (230 nm)
min
Cmpd **13**
mAU (230 nm)
min
Cmpd **14**
mAU (230 nm)
min
Cmpd **15**
mAU (230 nm)
min
Cmpd **16**
mAU (230 nm)
min
Cmpd **17**
mAU (230 nm)
min
Cmpd **18**
mAU (230 nm)
min
Cmpd **19**
mAU (254 nm)
min
Cmpd **20**
mAU (230 nm)
min

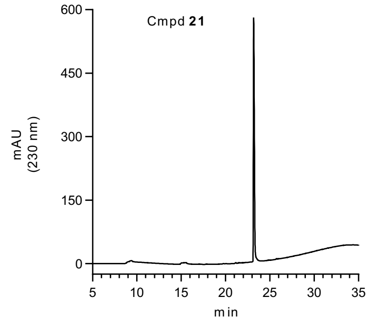
Cmpd 21
mAU (230 nm)
min

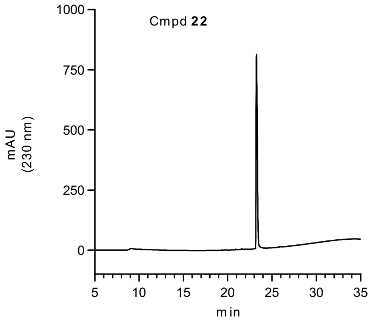
Cmpd 22
mAU (230 nm)
min

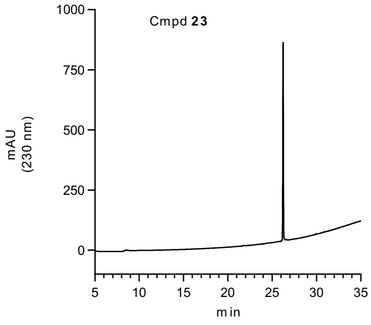
Cmpd 23
mAU (230 nm)
min

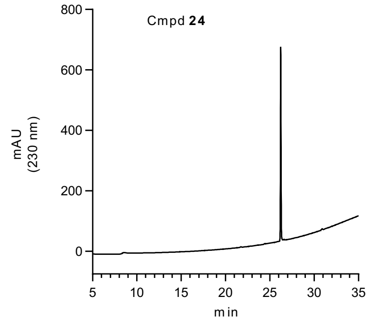
Cmpd 24
mAU (230 nm)
min

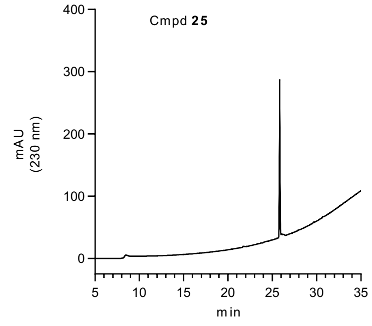
Cmpd 25
mAU (230 nm)
min

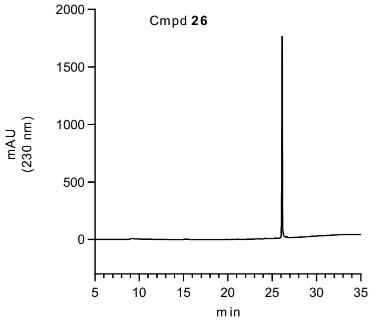
Cmpd 26
mAU (230 nm)
min

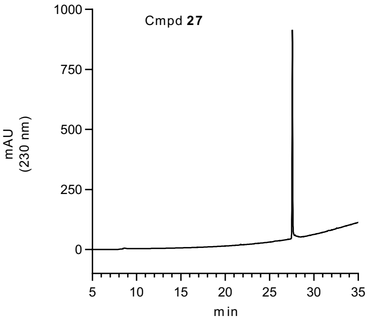
Cmpd 27
mAU (230 nm)
min

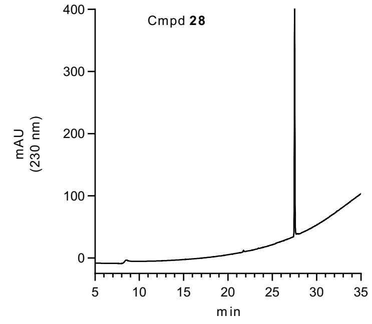
Cmpd 28
mAU (230 nm)
min

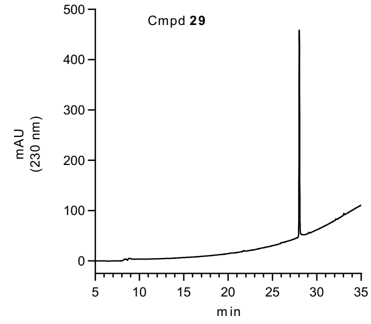
Cmpd 29
mAU (230 nm)
min

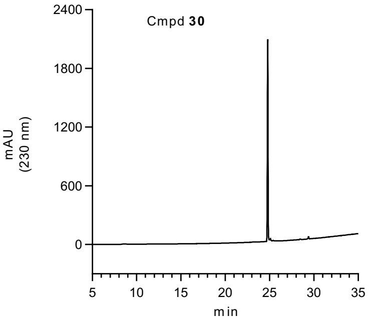
Cmpd 30
mAU (230 nm)
min

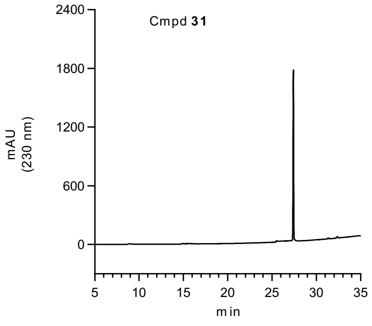
Cmpd 31
mAU (230 nm)
min

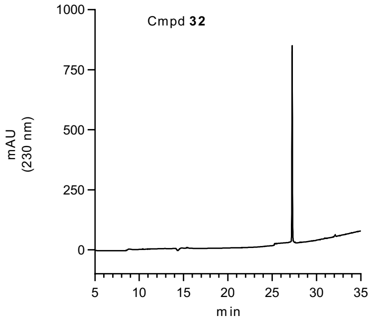
Cmpd 32
mAU (230 nm)
min

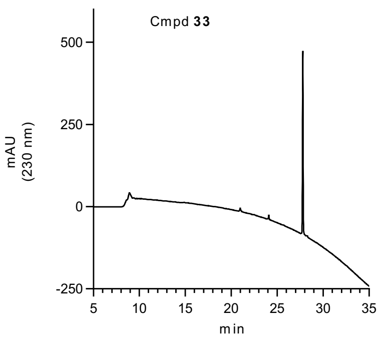

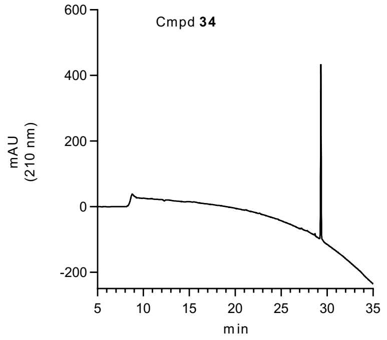

Peptide 35

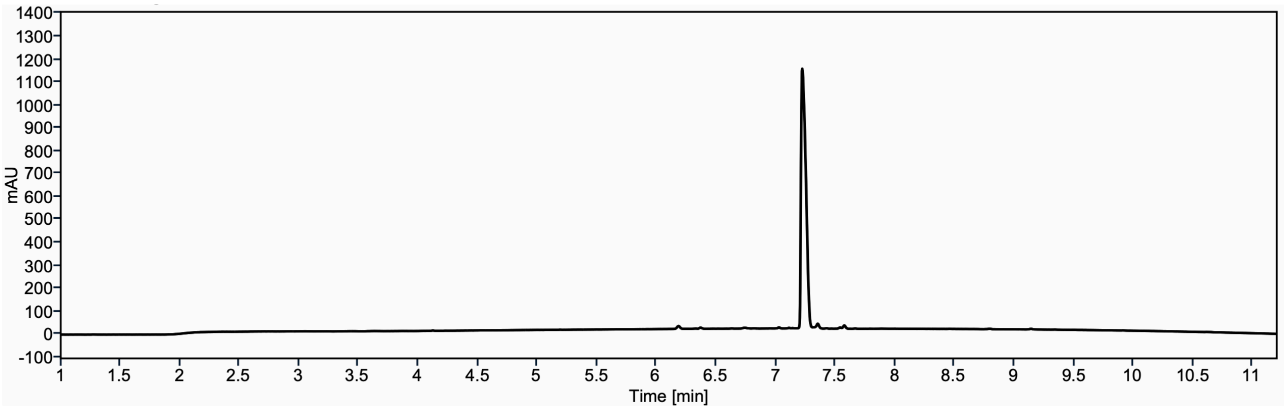

Peptide 36

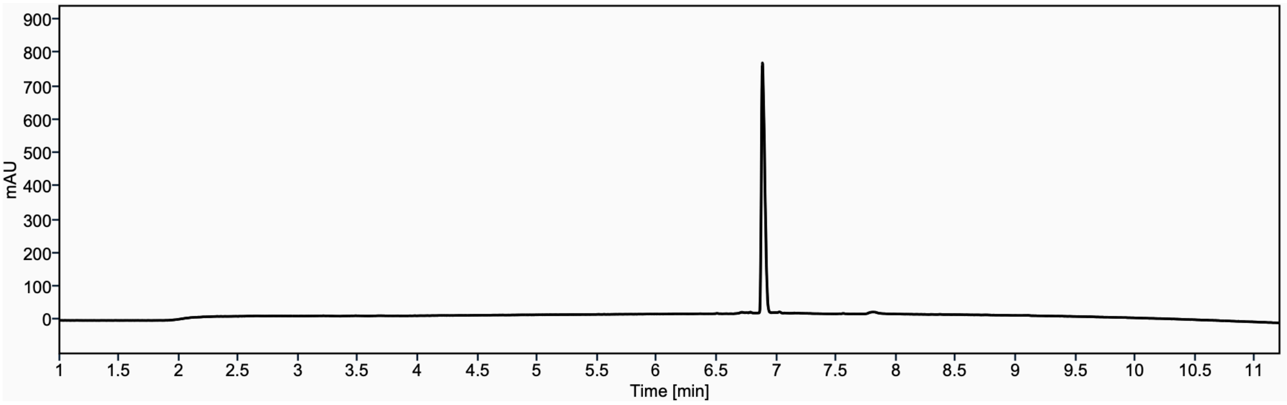

Peptide 37

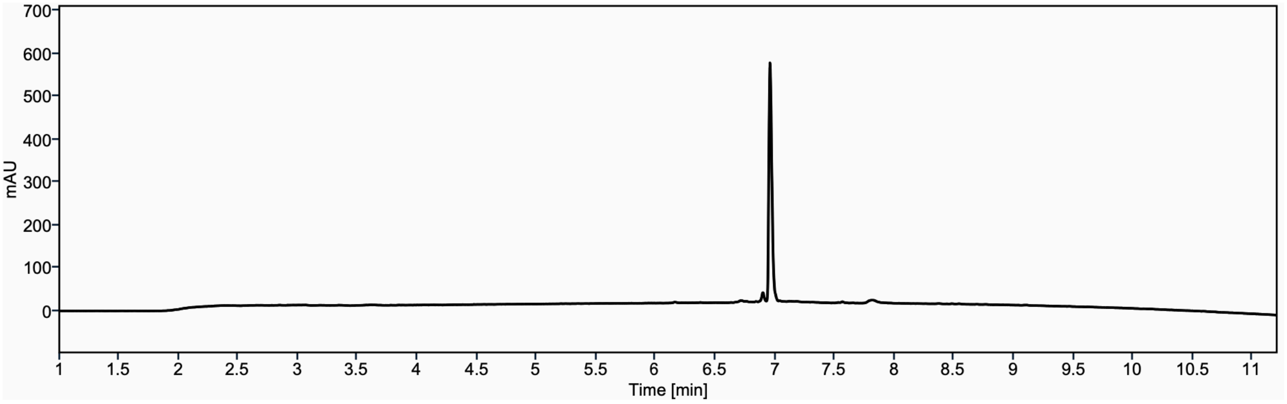

Peptide 38

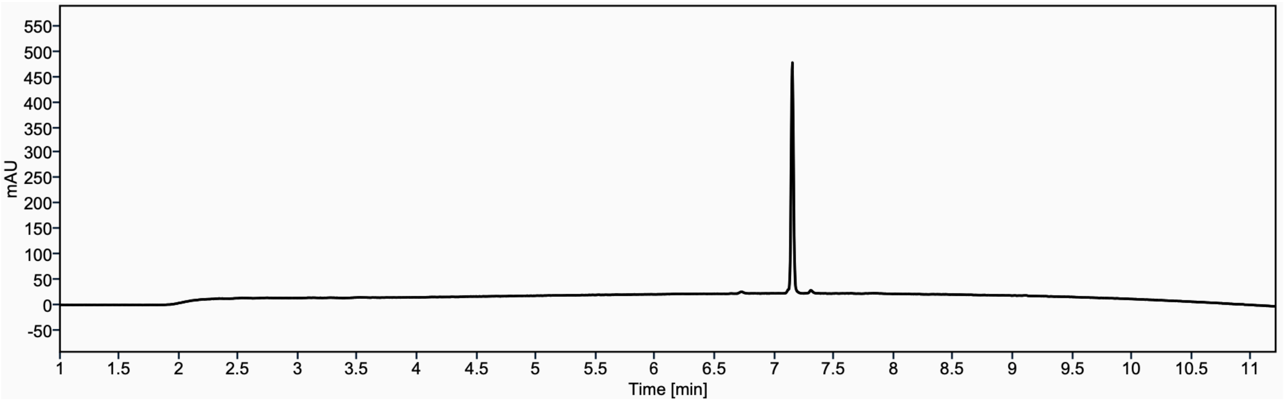

Peptide 39

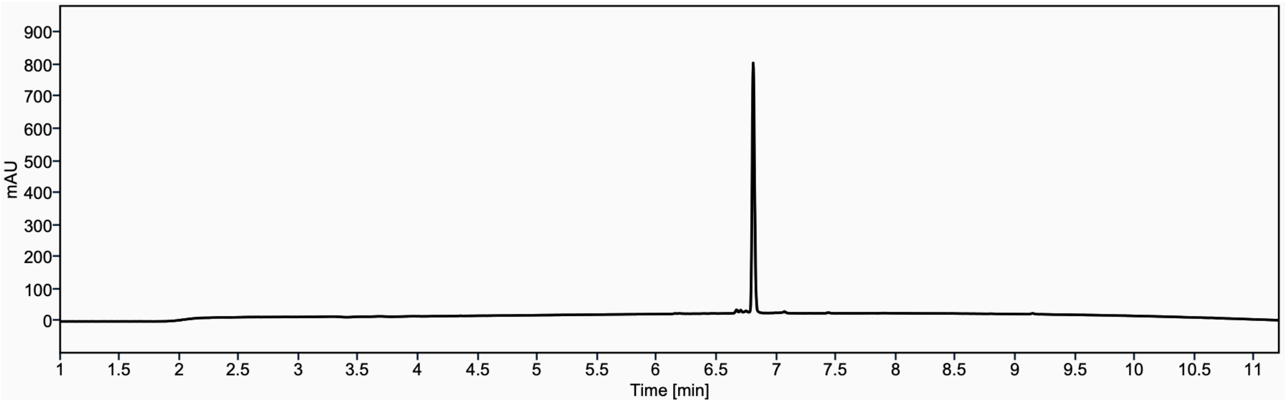

Peptide 40

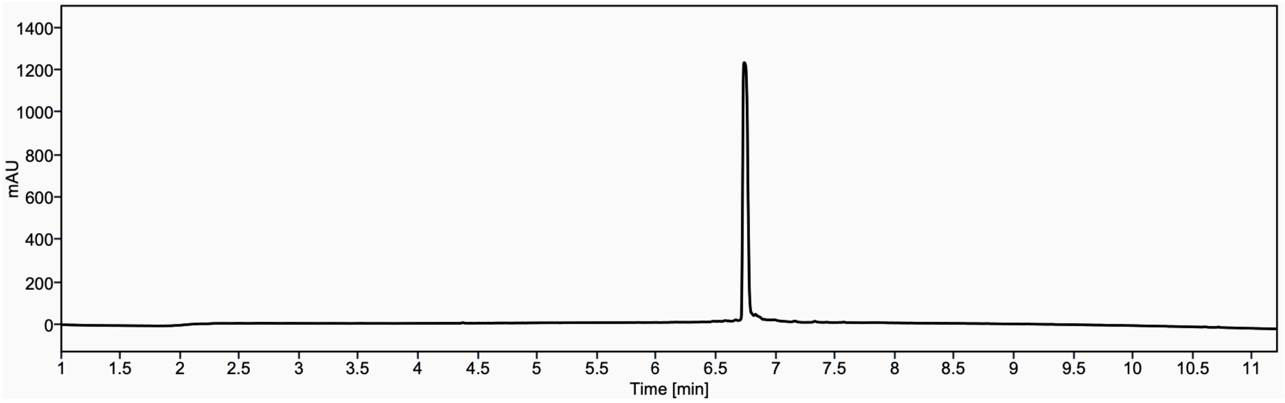

Peptide 41

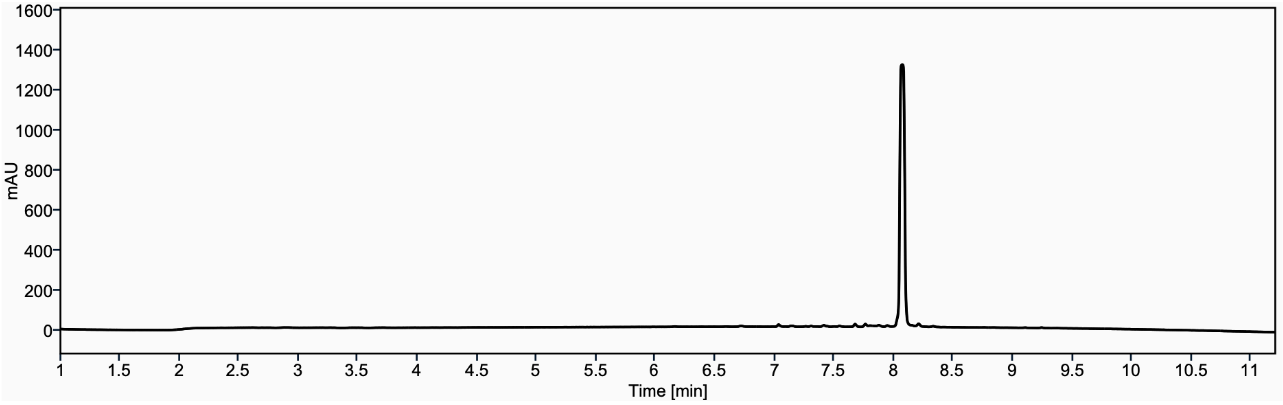

Peptide 42

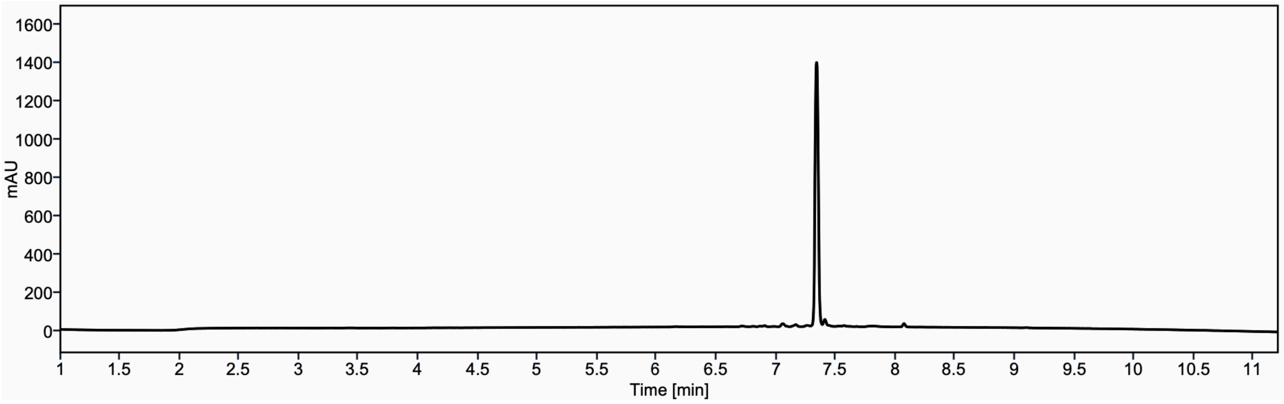

Peptide 43

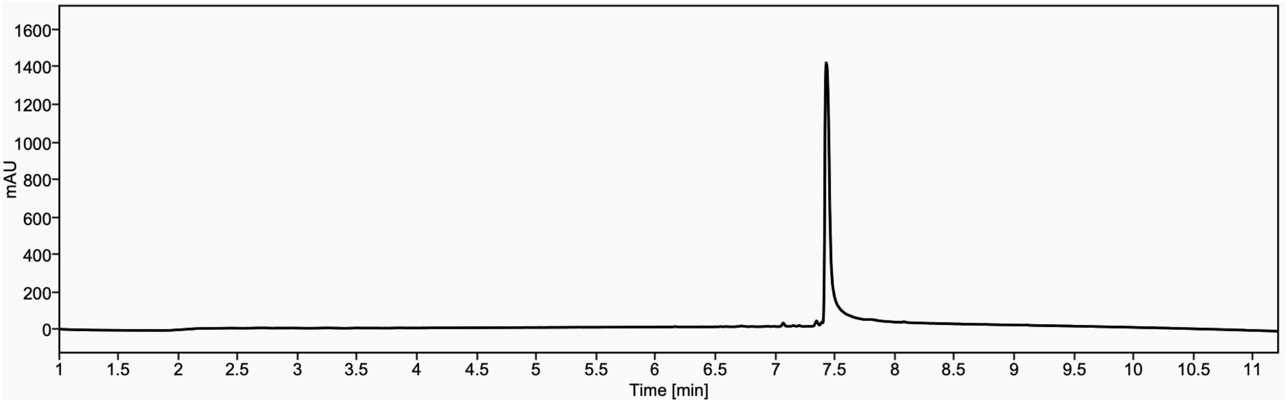

Peptide 44

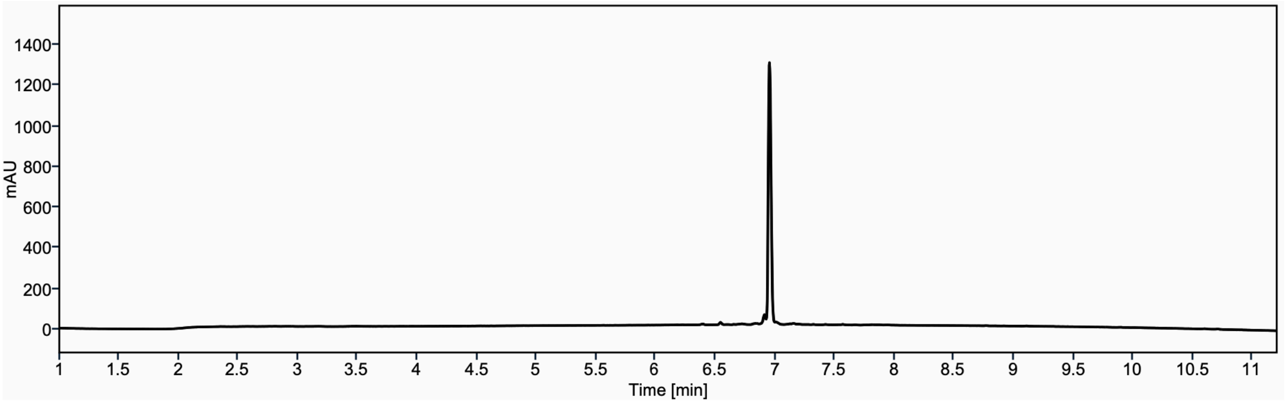

Peptide 45

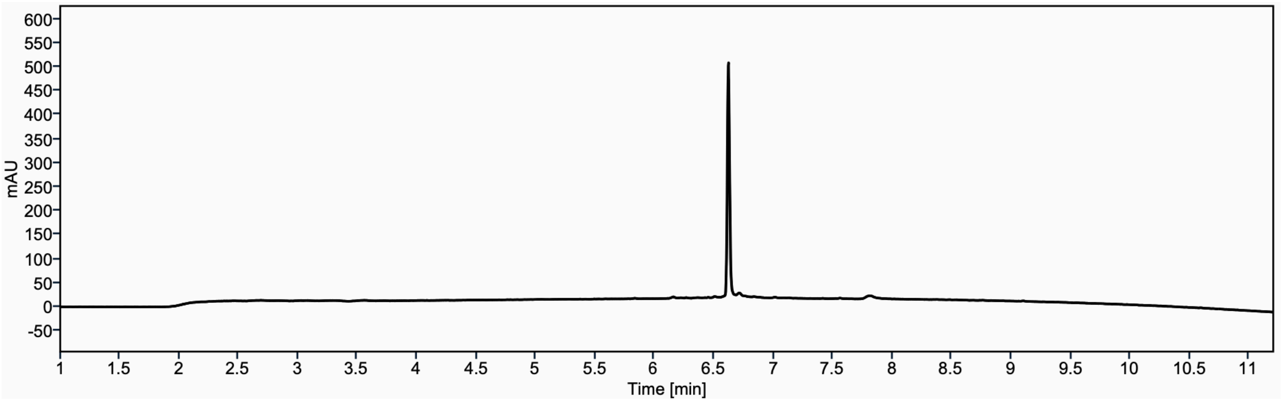

Peptide 46

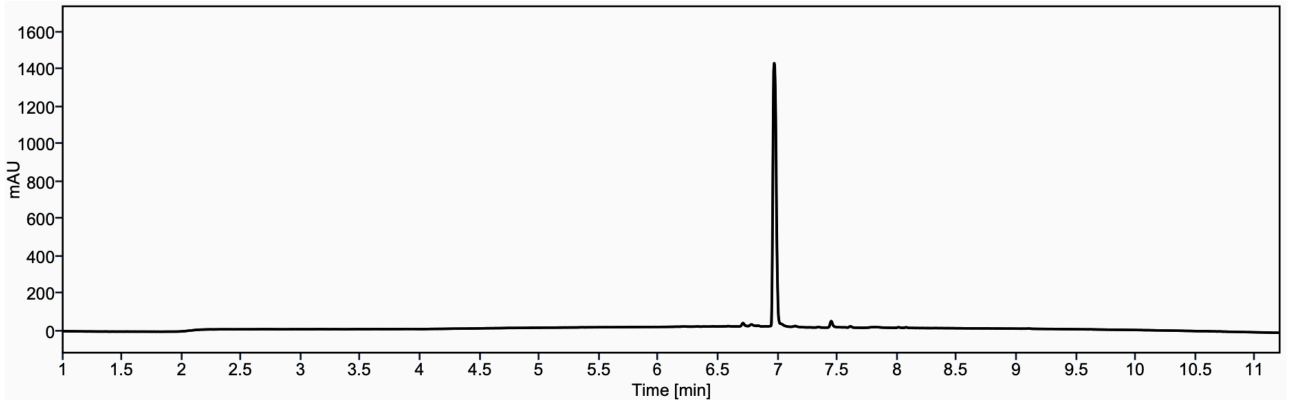

Peptide 47

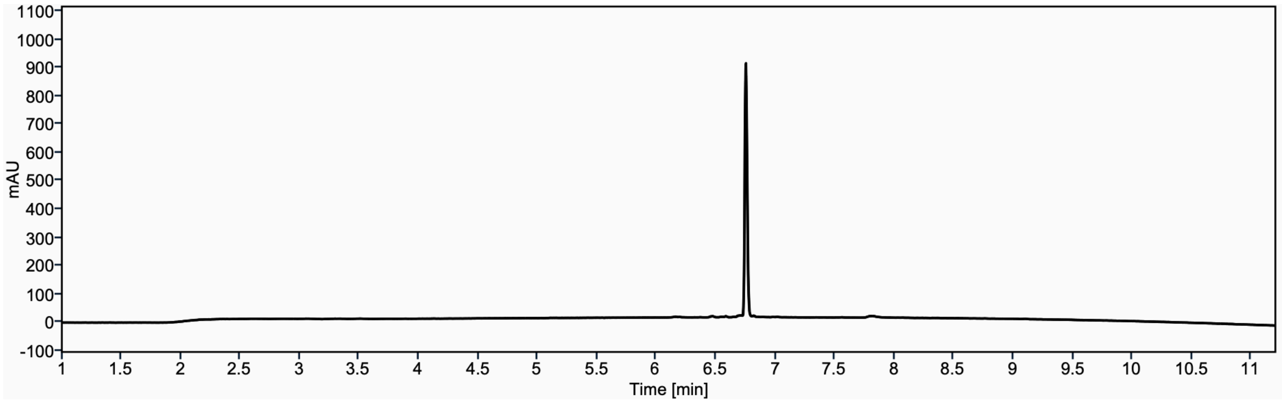

### Copies of NMR spectra

$^{1}H$ NMR (600 MHz, DMSO-$d_6$) of **GGVKac**

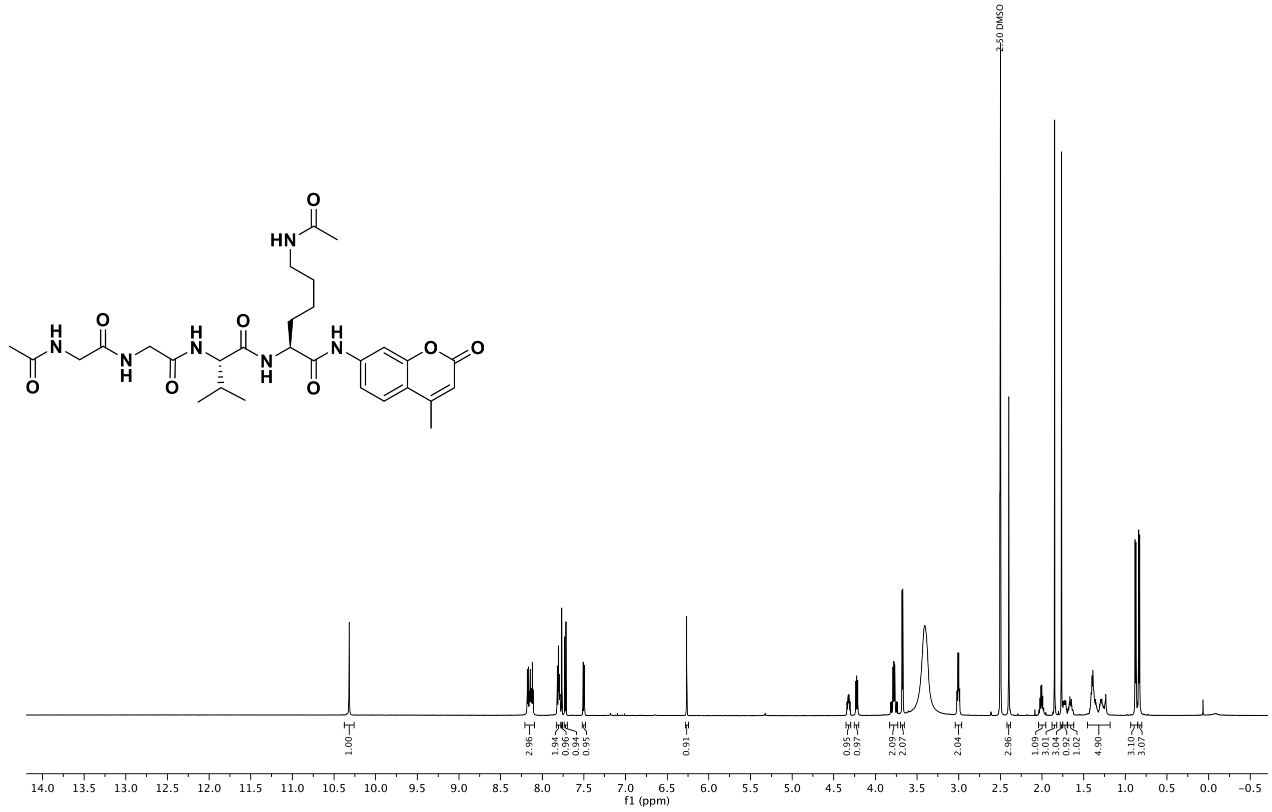

$^{13}C$ NMR (151 MHz, DMSO) of **GGVKac**

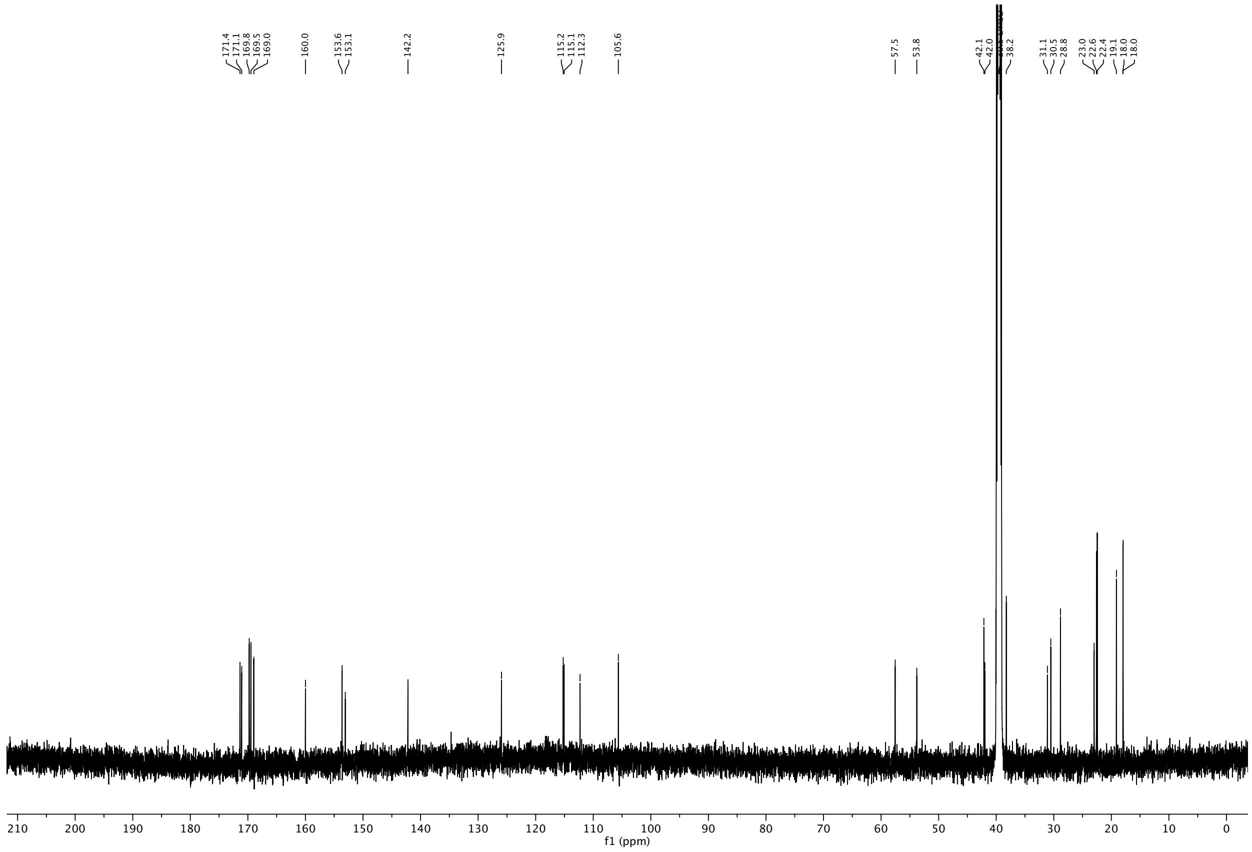

$^{1}$H NMR (600 MHz, DMSO-$d_6$) of **GGVKdec**

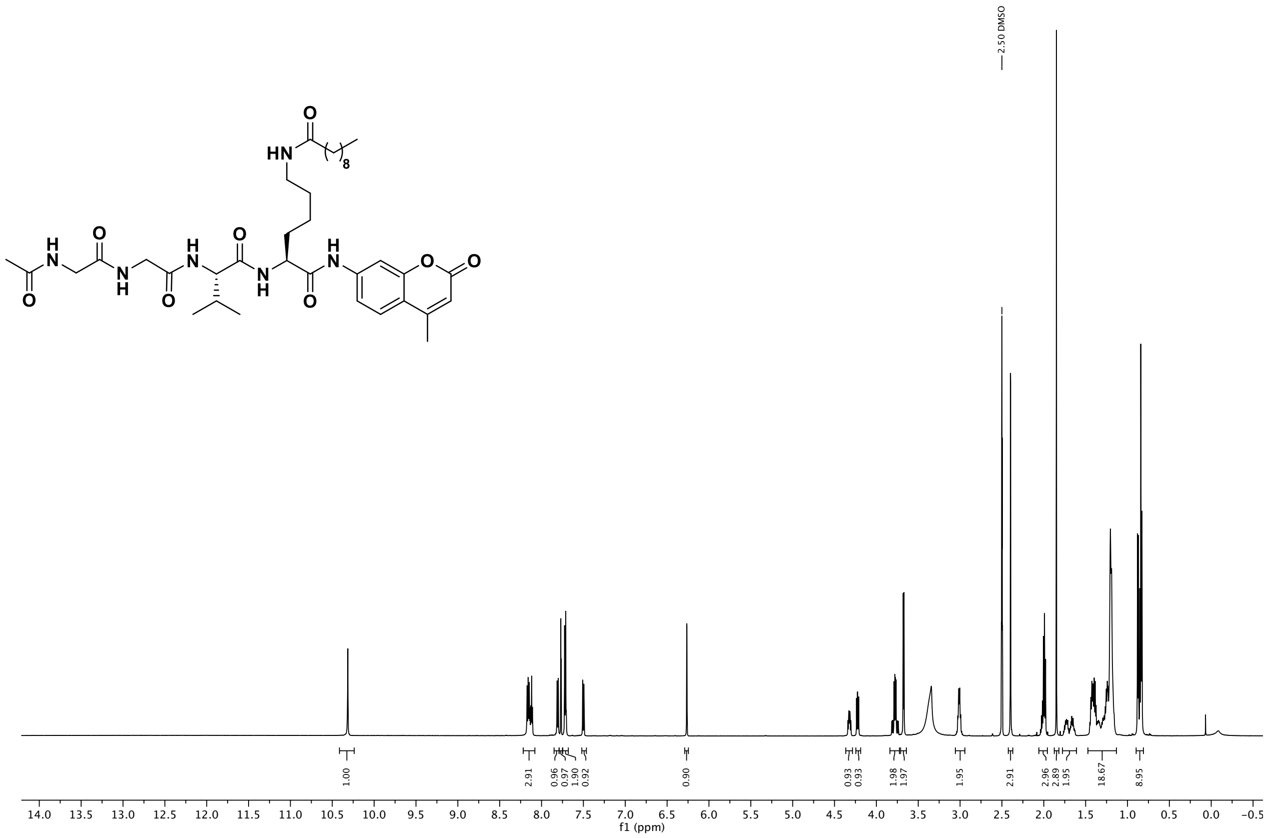

$^{13}$C NMR (151 MHz, DMSO) of **GGVKdec**

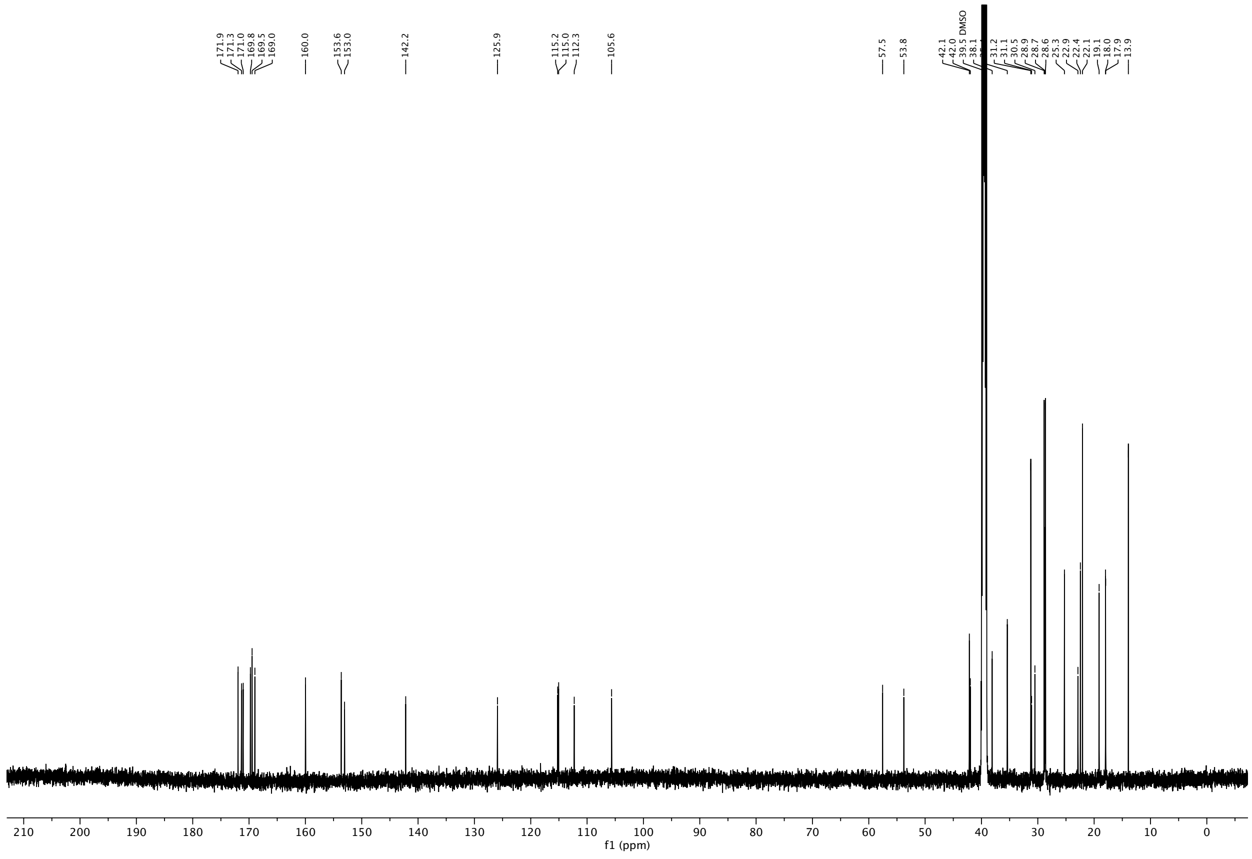

$^{1}$H NMR (600 MHz, DMSO-$d_6$) of **GGVKmyr**

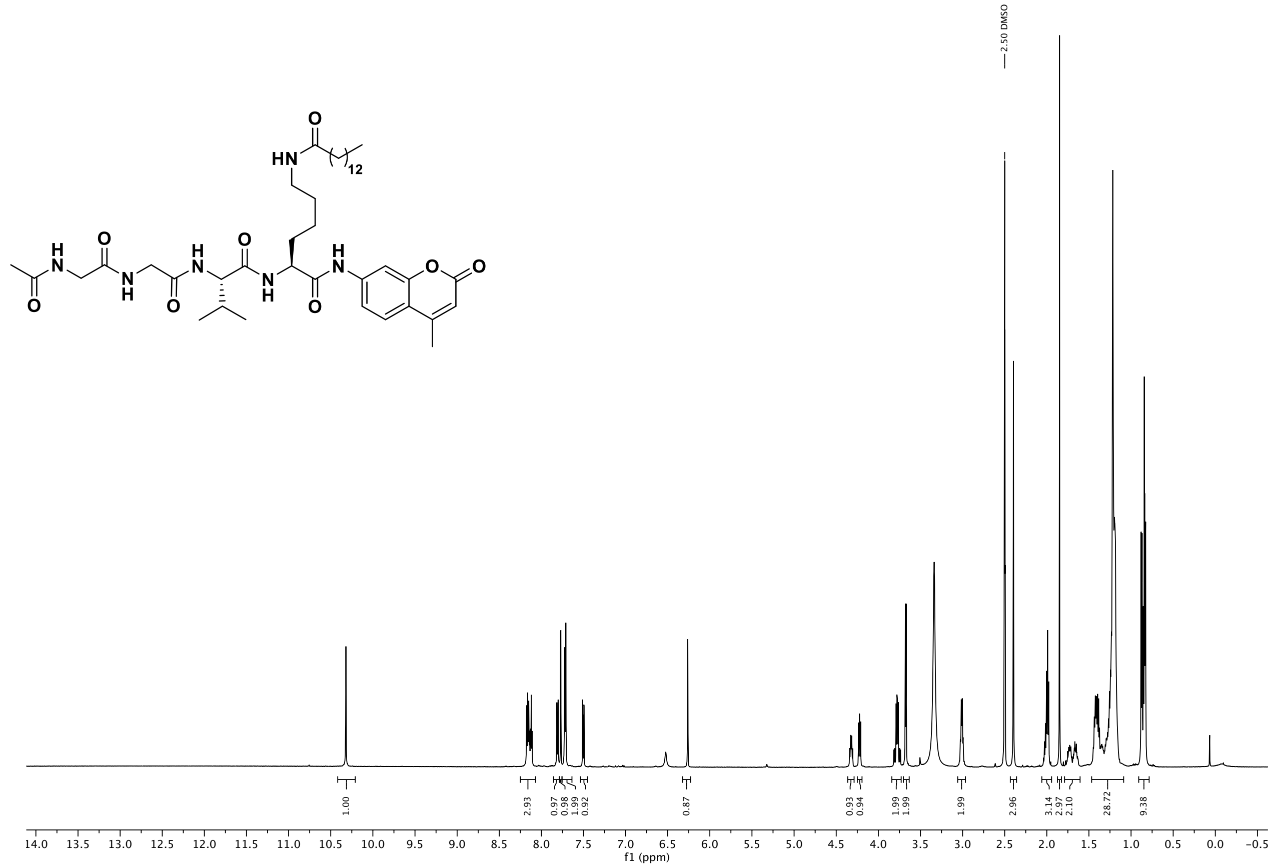

$^{13}$C NMR (151 MHz, DMSO) of **GGVKmyr**

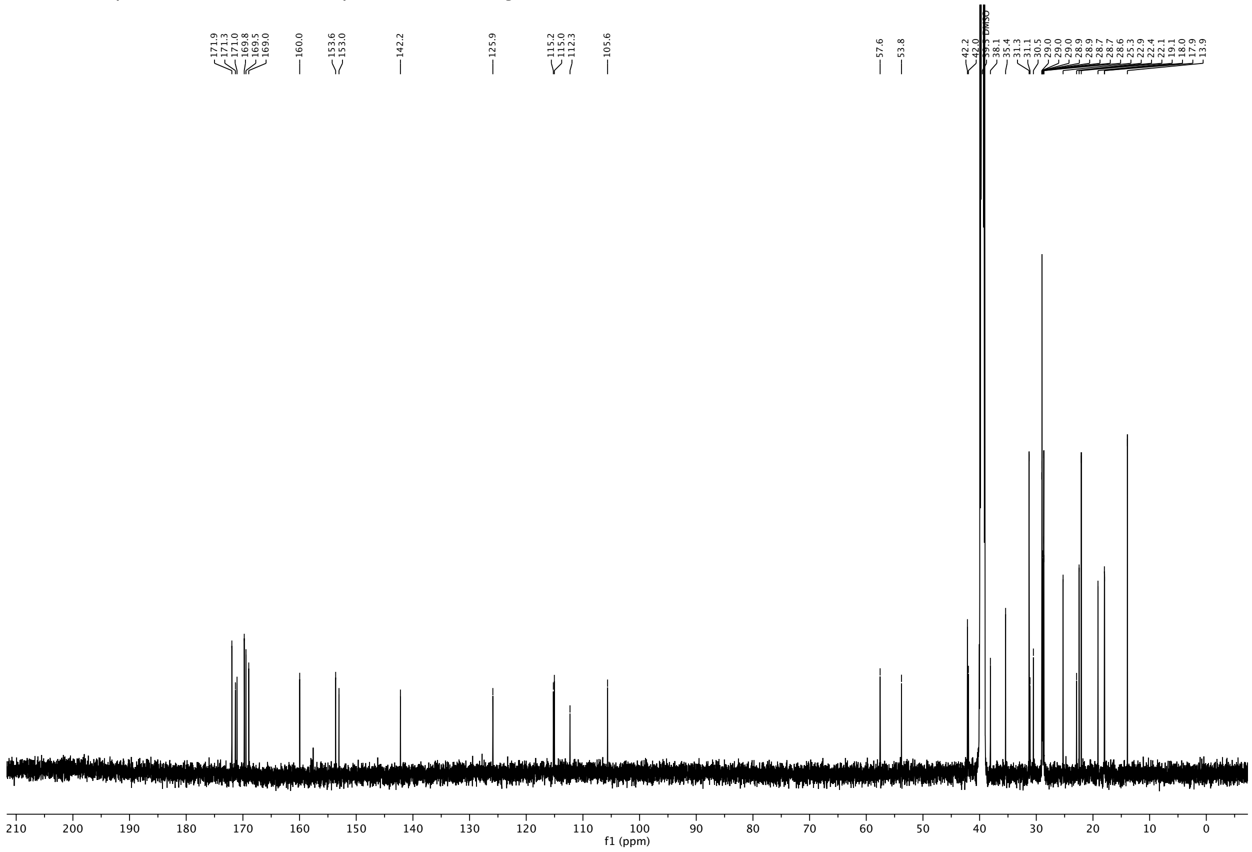

$^{1}H$ NMR (600 MHz, DMSO-$d_6$) of **APRKdec**

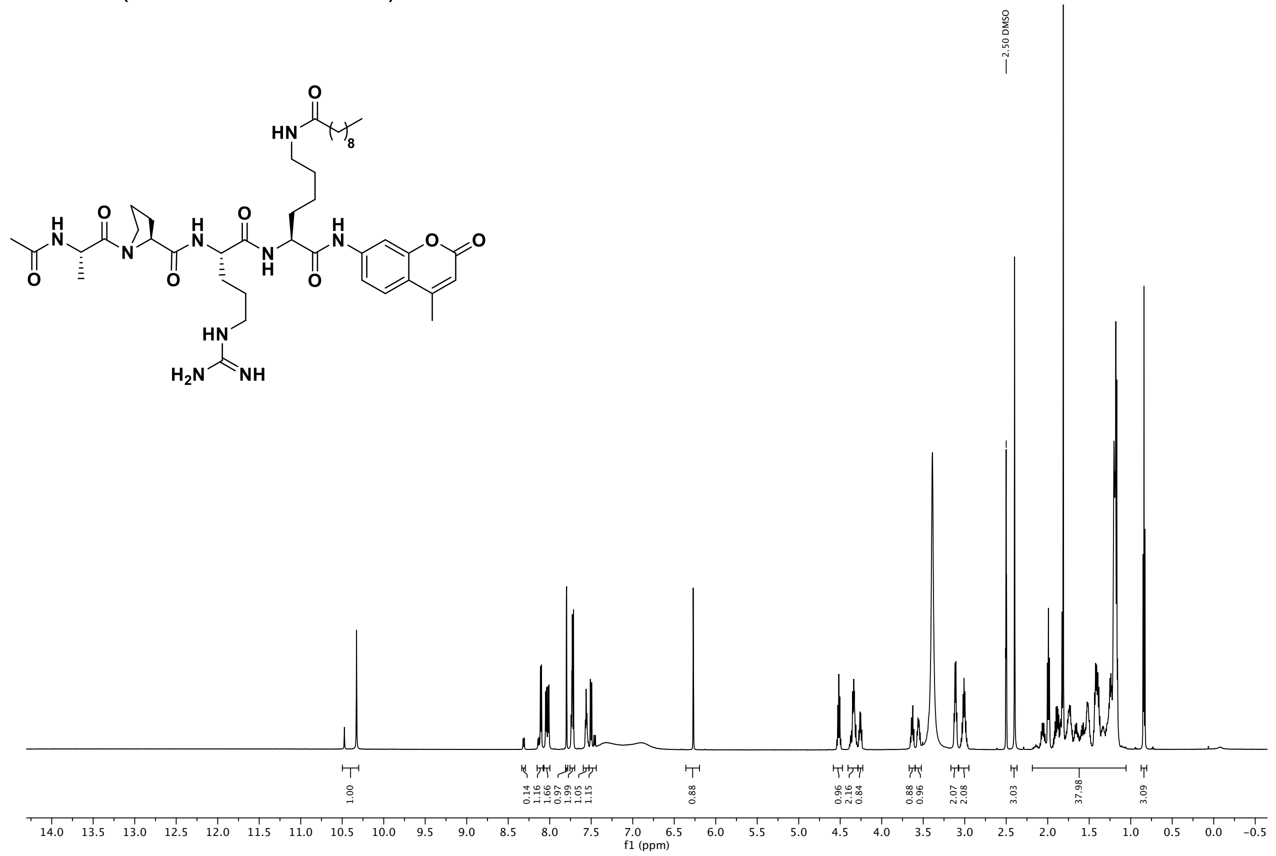

$^{13}C$ NMR (151 MHz, DMSO) of **APRKdec**

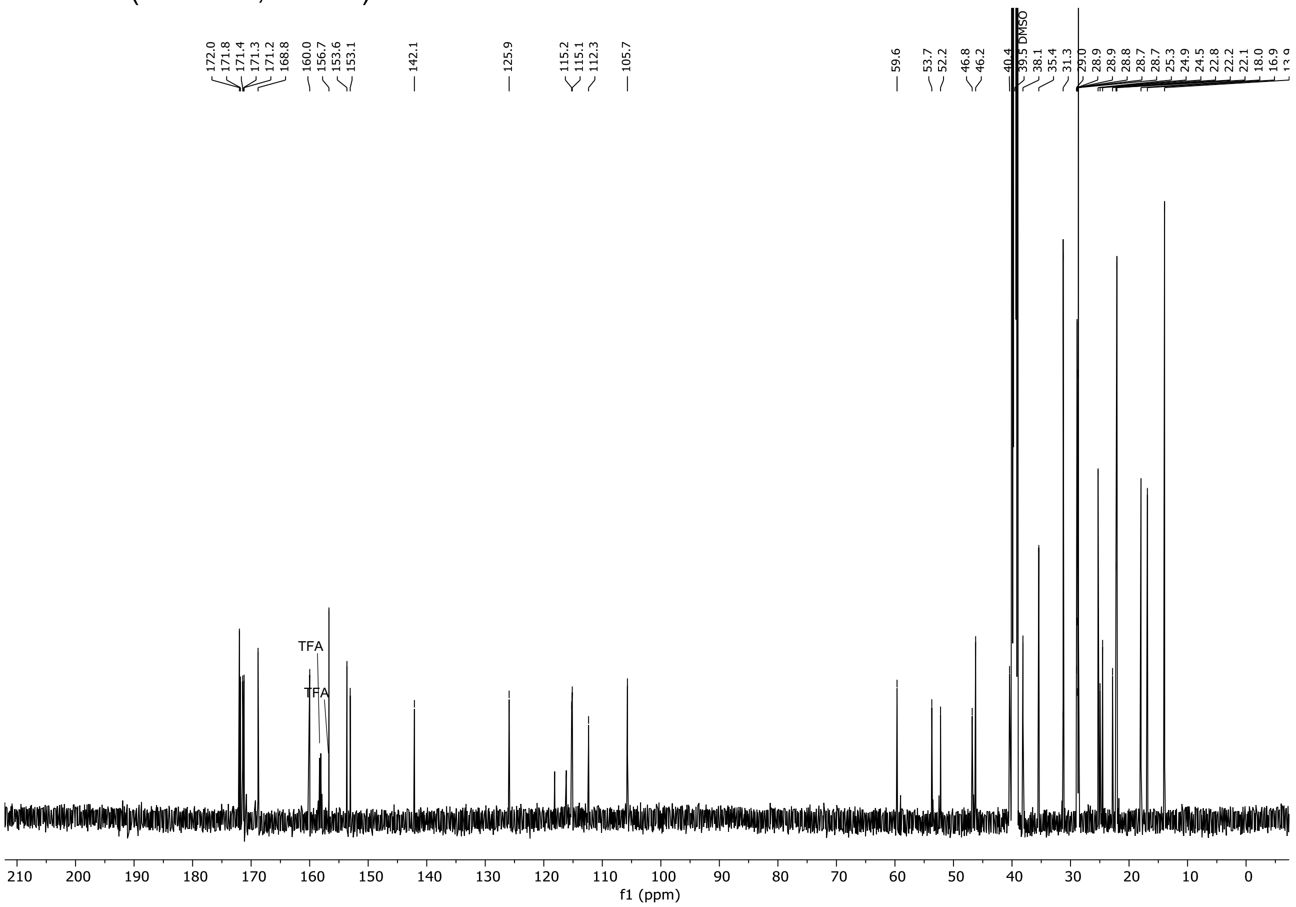

$^{1}H$ NMR (600 MHz, DMSO-$d_6$) of compound **APRKmyr**

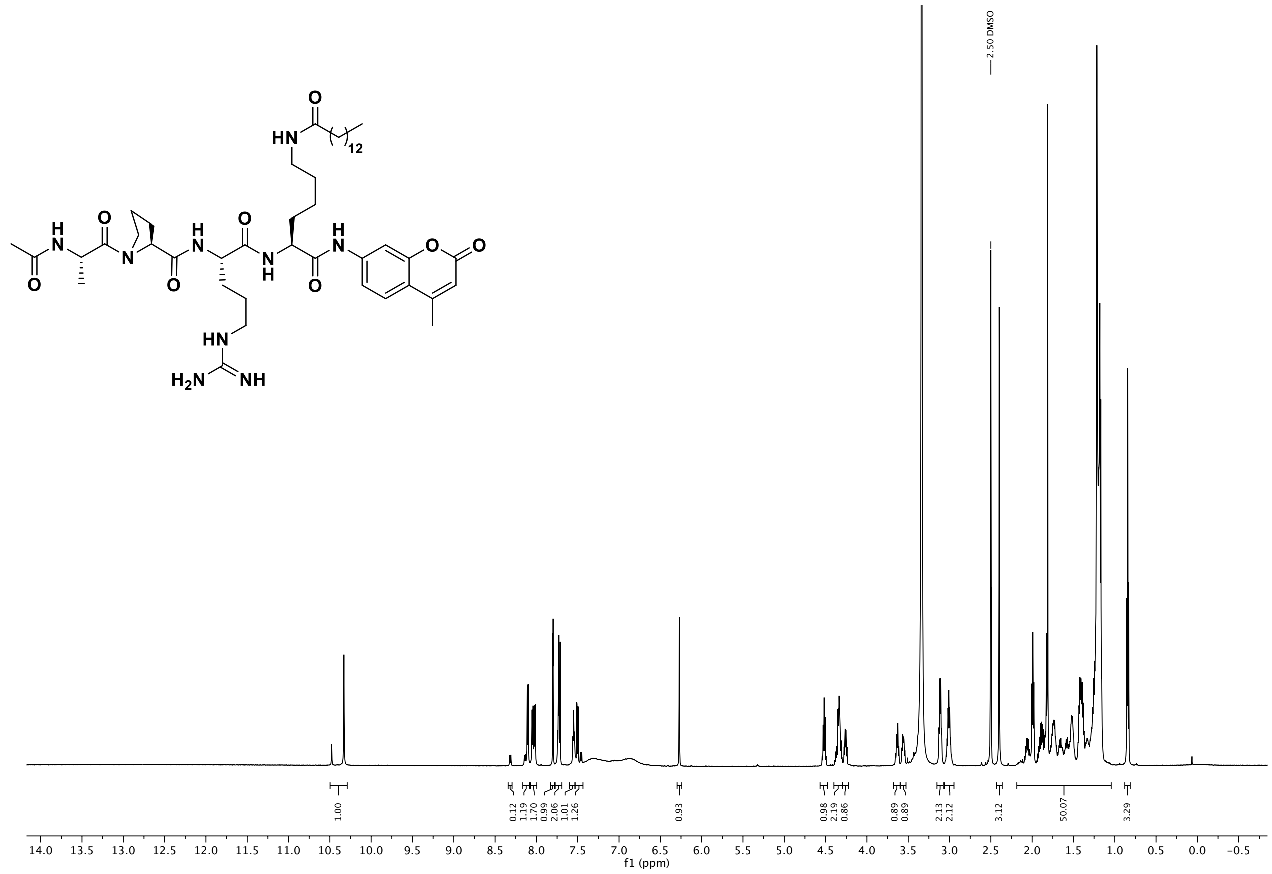

$^{13}C$ NMR (151 MHz, DMSO) of **APRKmyr**

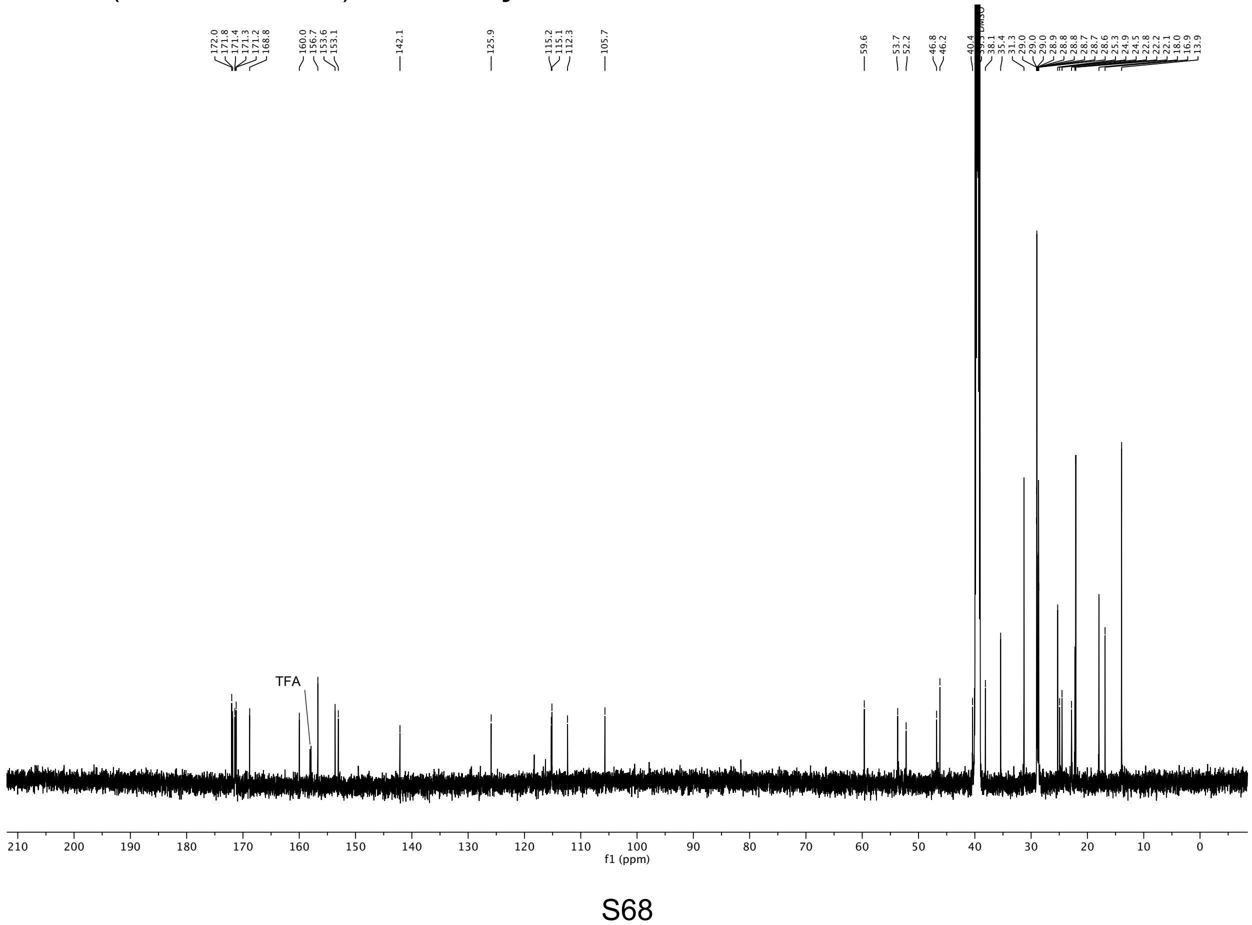

$^{1}H$ NMR (600 MHz, DMSO-$d_6$) of **ETDKglu**

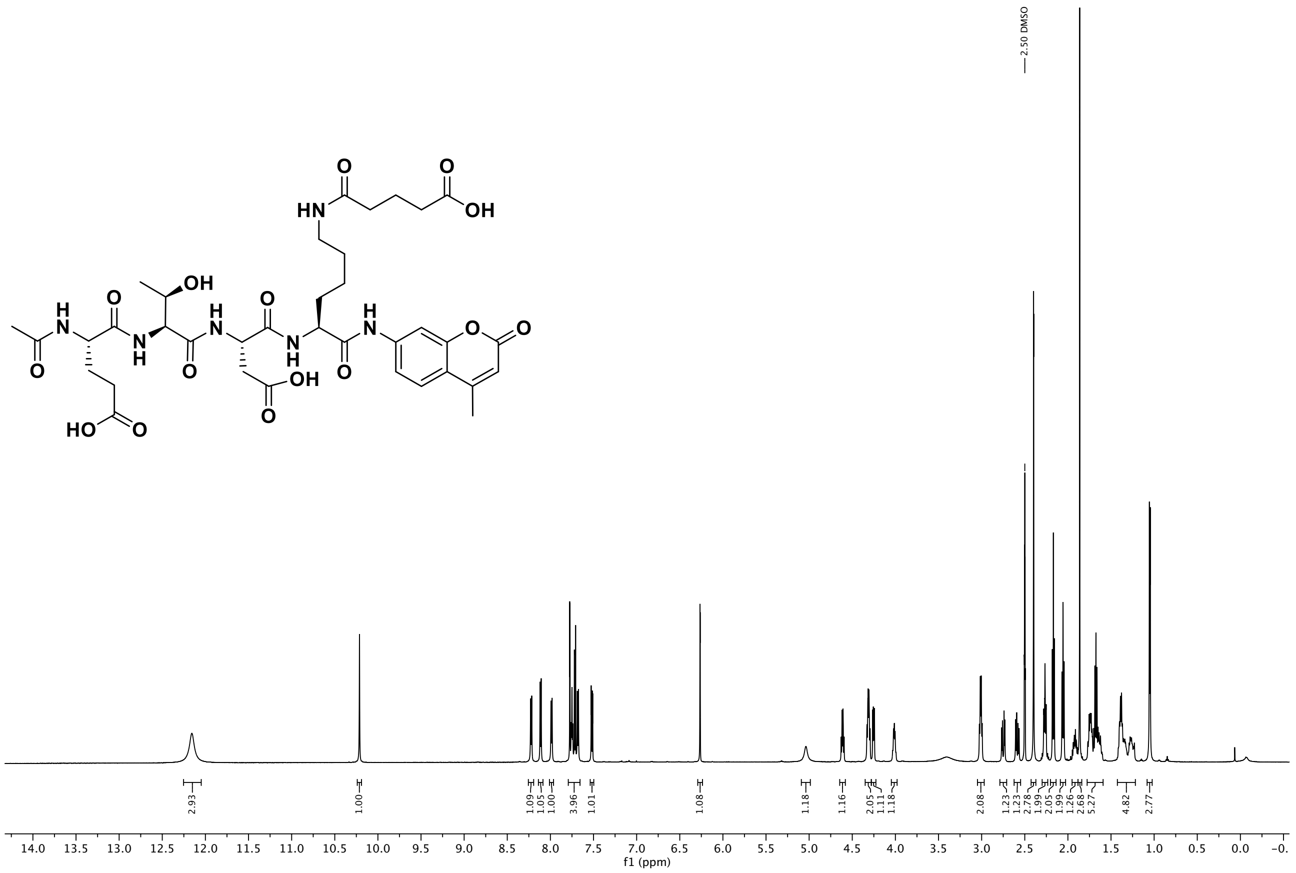

$^{13}C$ NMR (151 MHz, DMSO) of **ETDKglu**

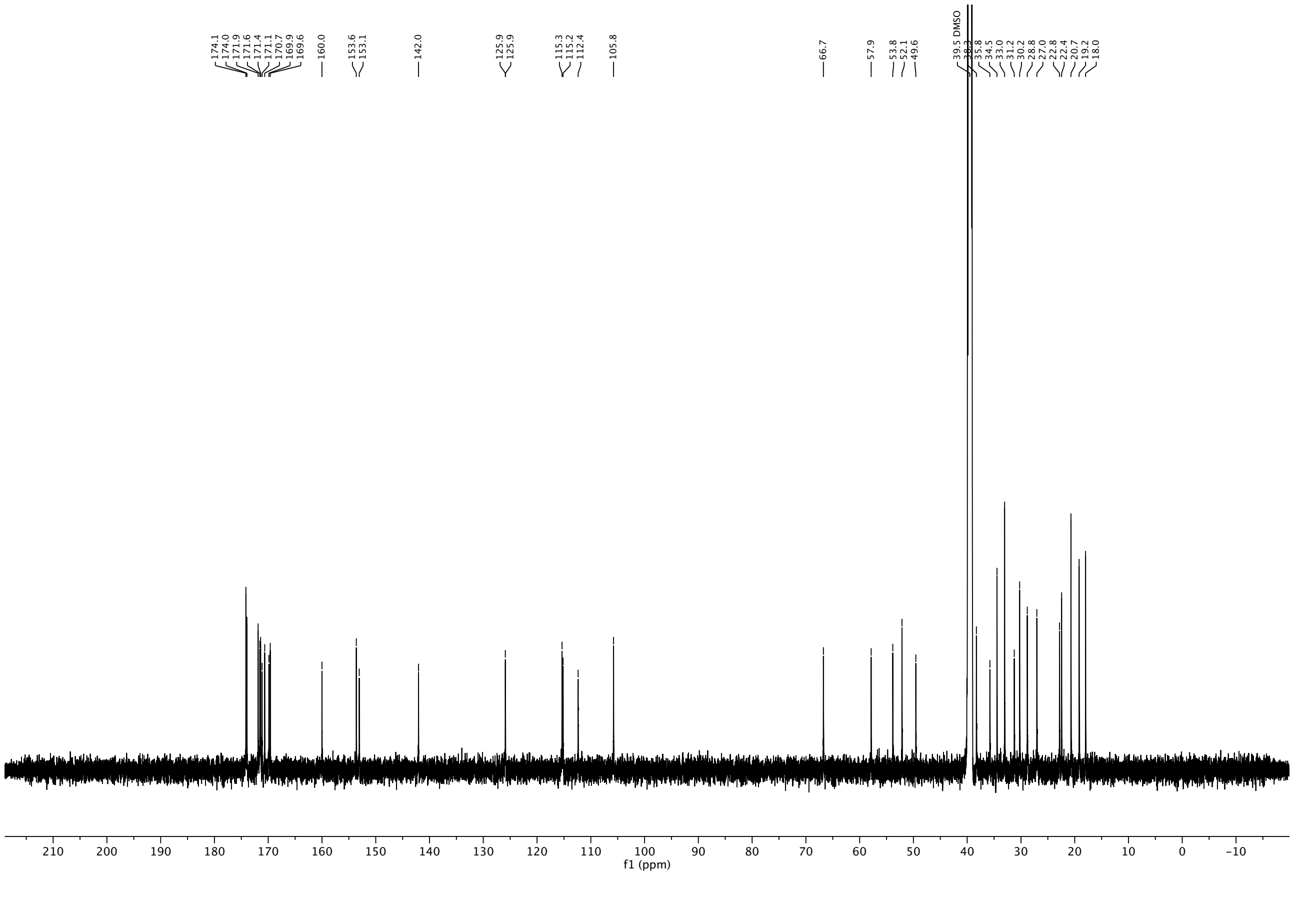

$^{1}$H NMR (600 MHz, DMSO-$d_6$) of **TARKhmg**

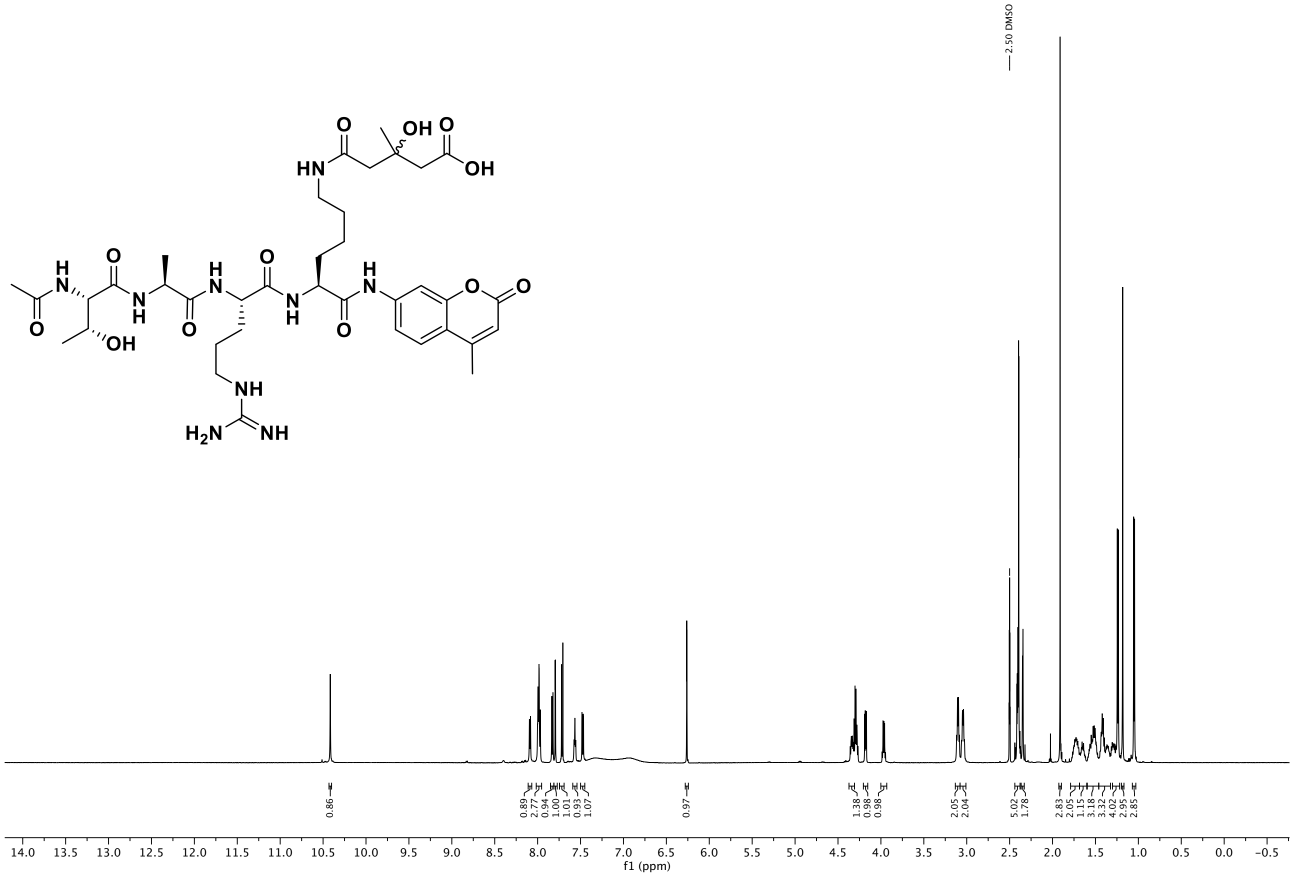

$^{13}$C NMR (151 MHz, DMSO) of **TARKhmg.**

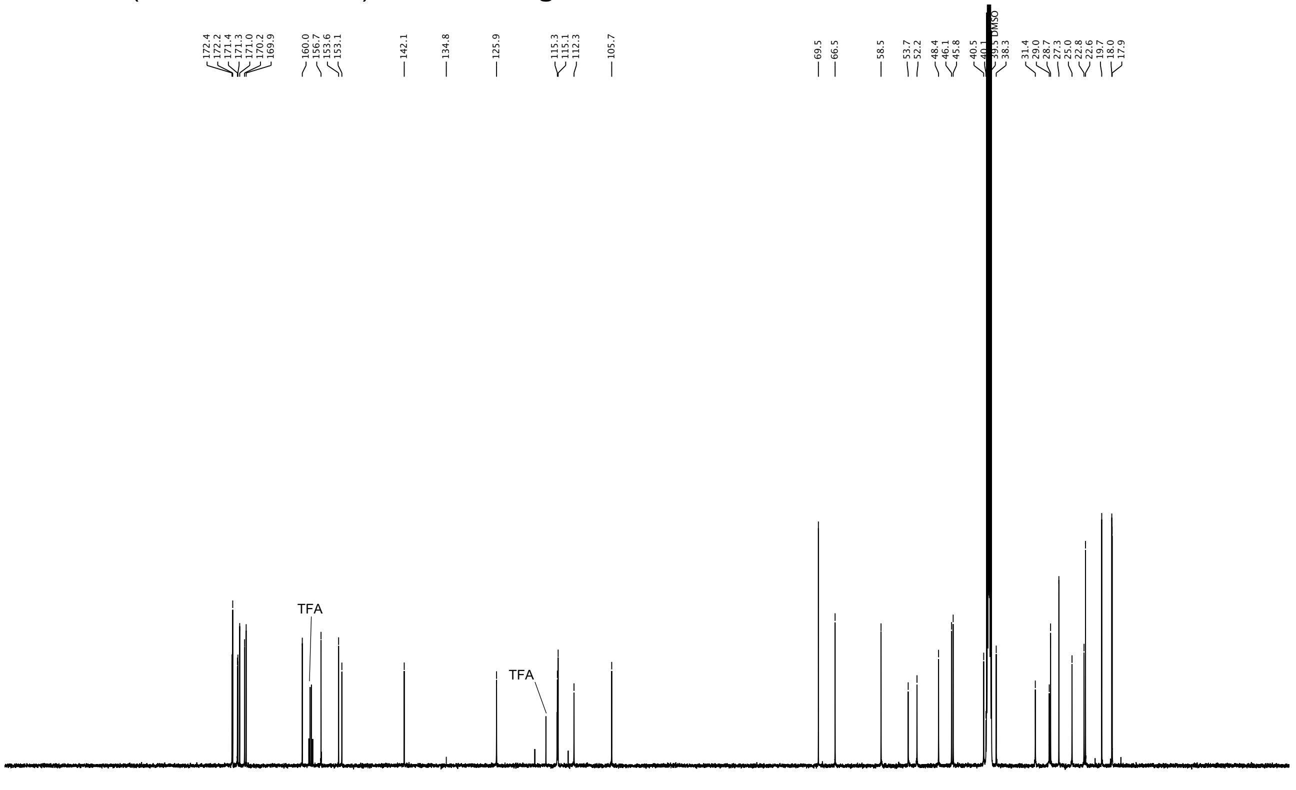

[1]H NMR (600 MHz, DMSO-$d_6$) of compound **4**.

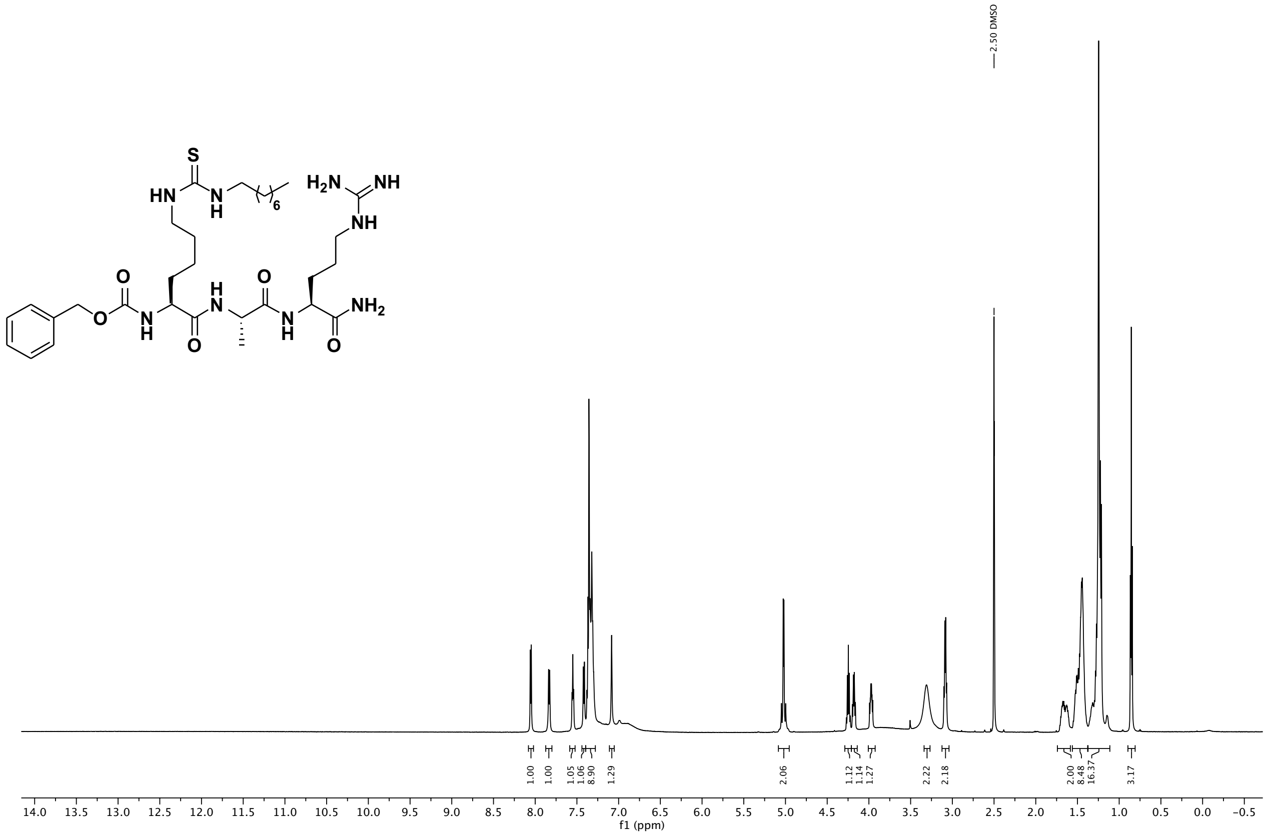

[13]C NMR (151 MHz, DMSO) of compound **4**.

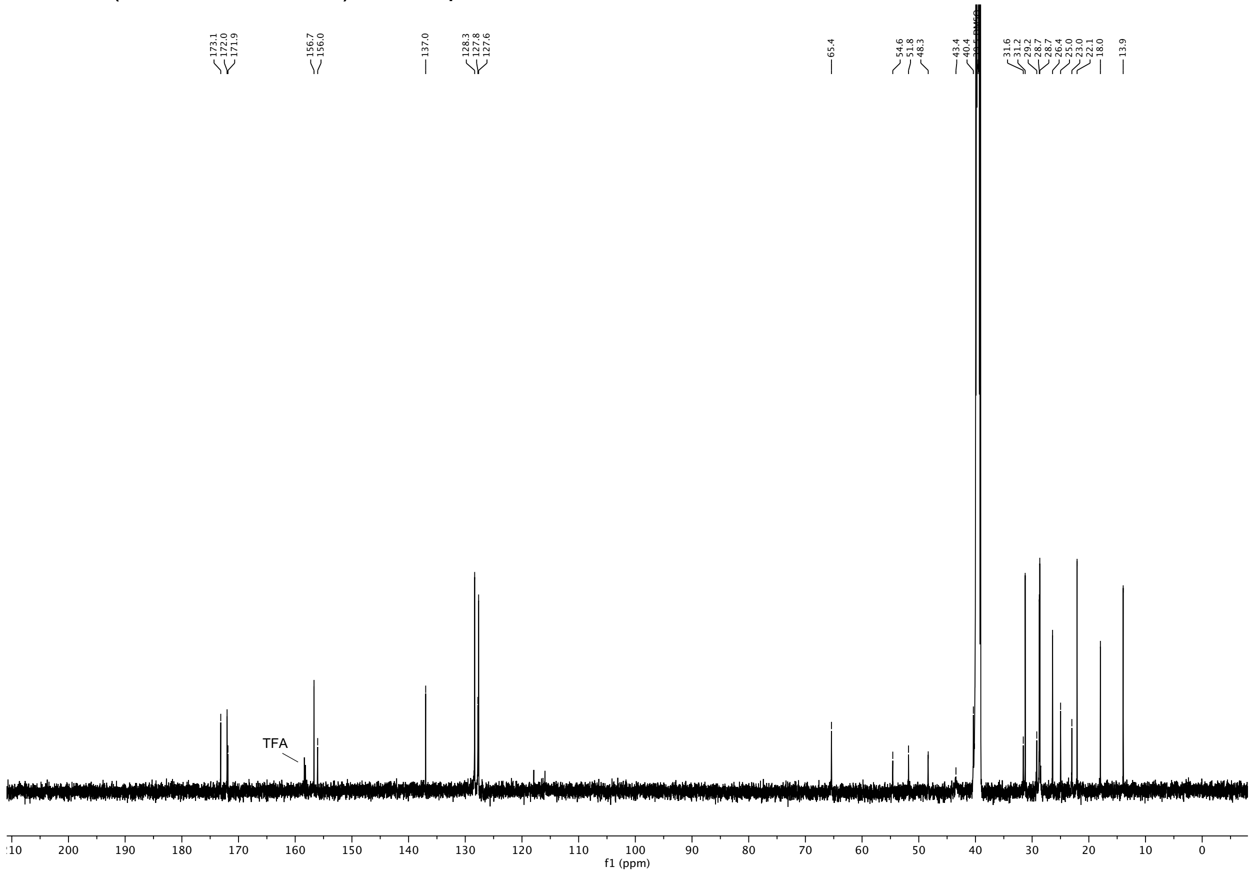

[1]H NMR (600 MHz, DMSO-$d_6$) of compound **5**.

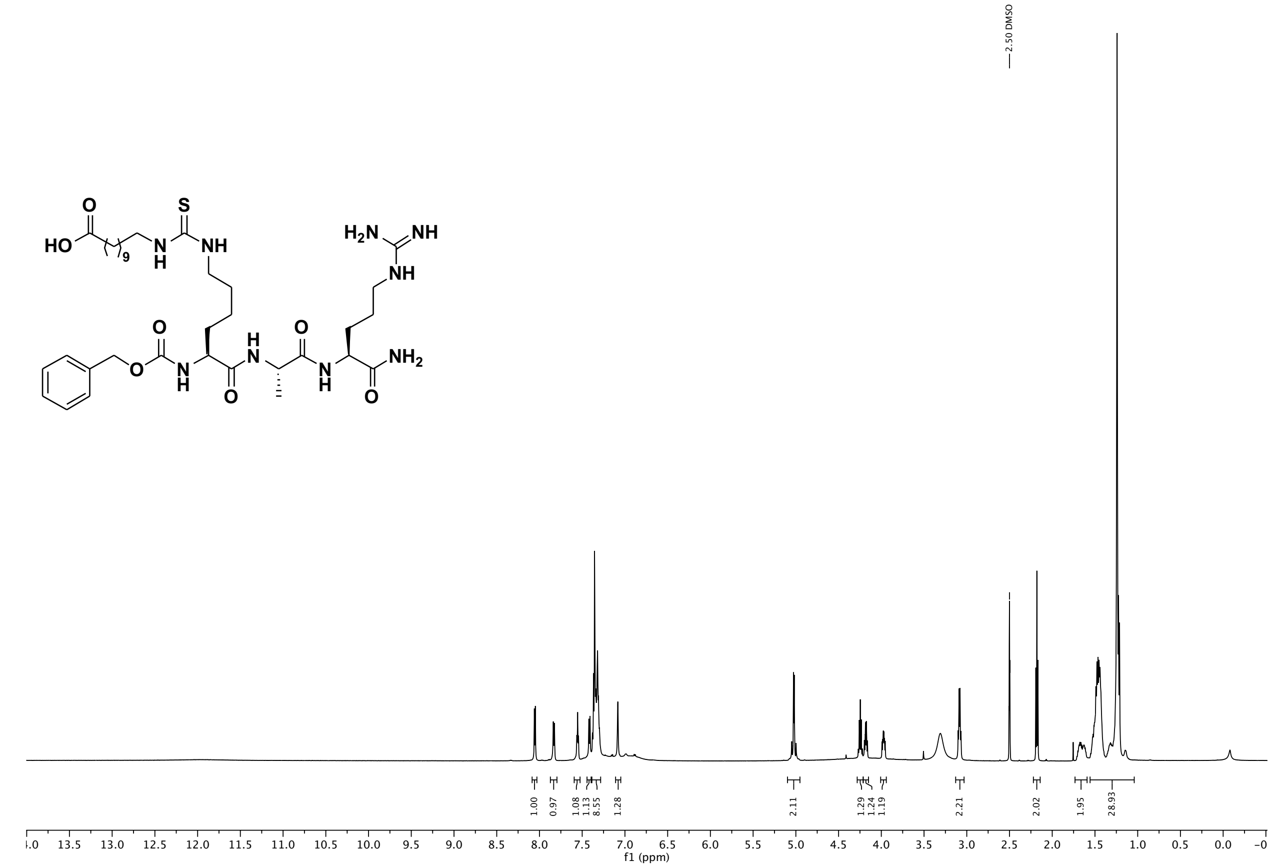

[13]C NMR (151 MHz, DMSO) of compound **5**.

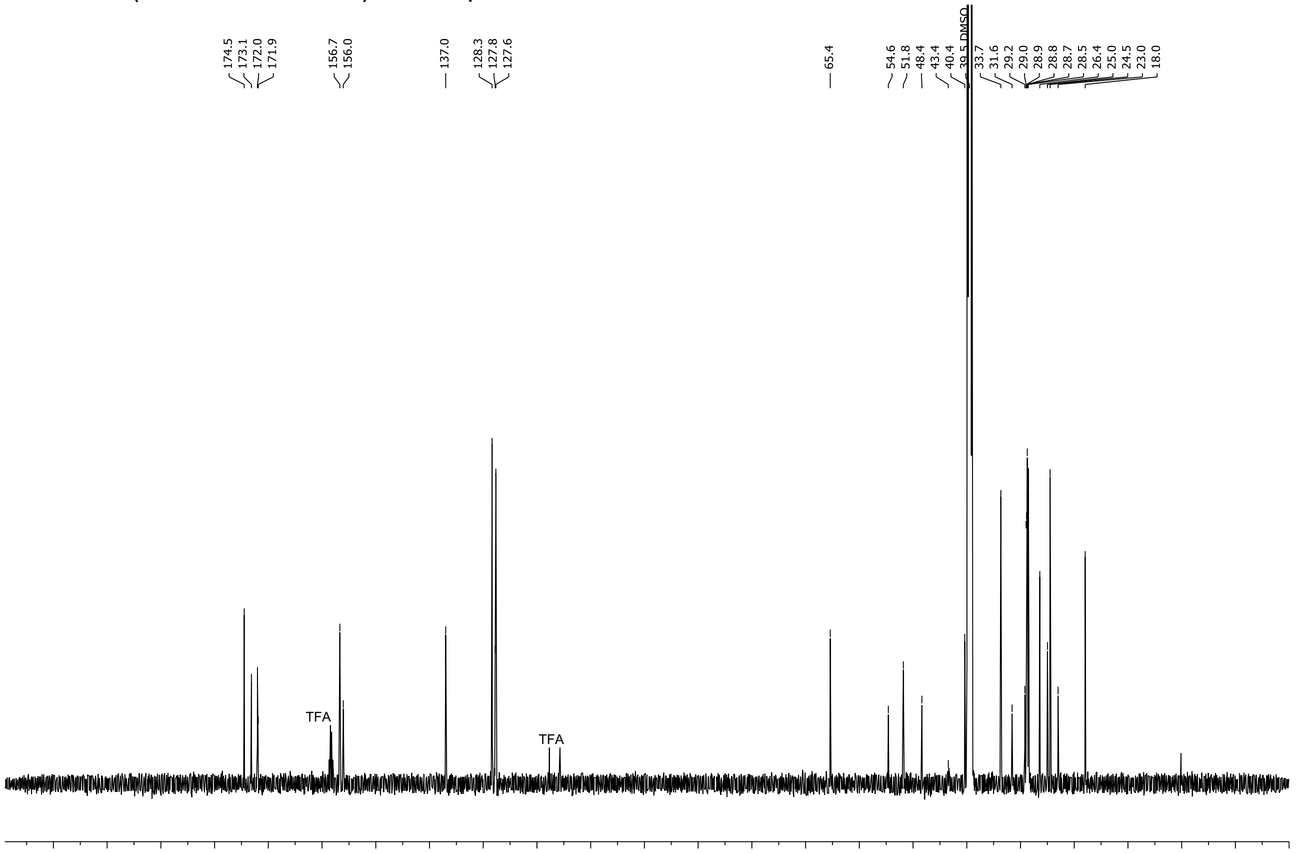

$^{1}H$ NMR (600 MHz, DMSO-$d_6$) of compound **6**.

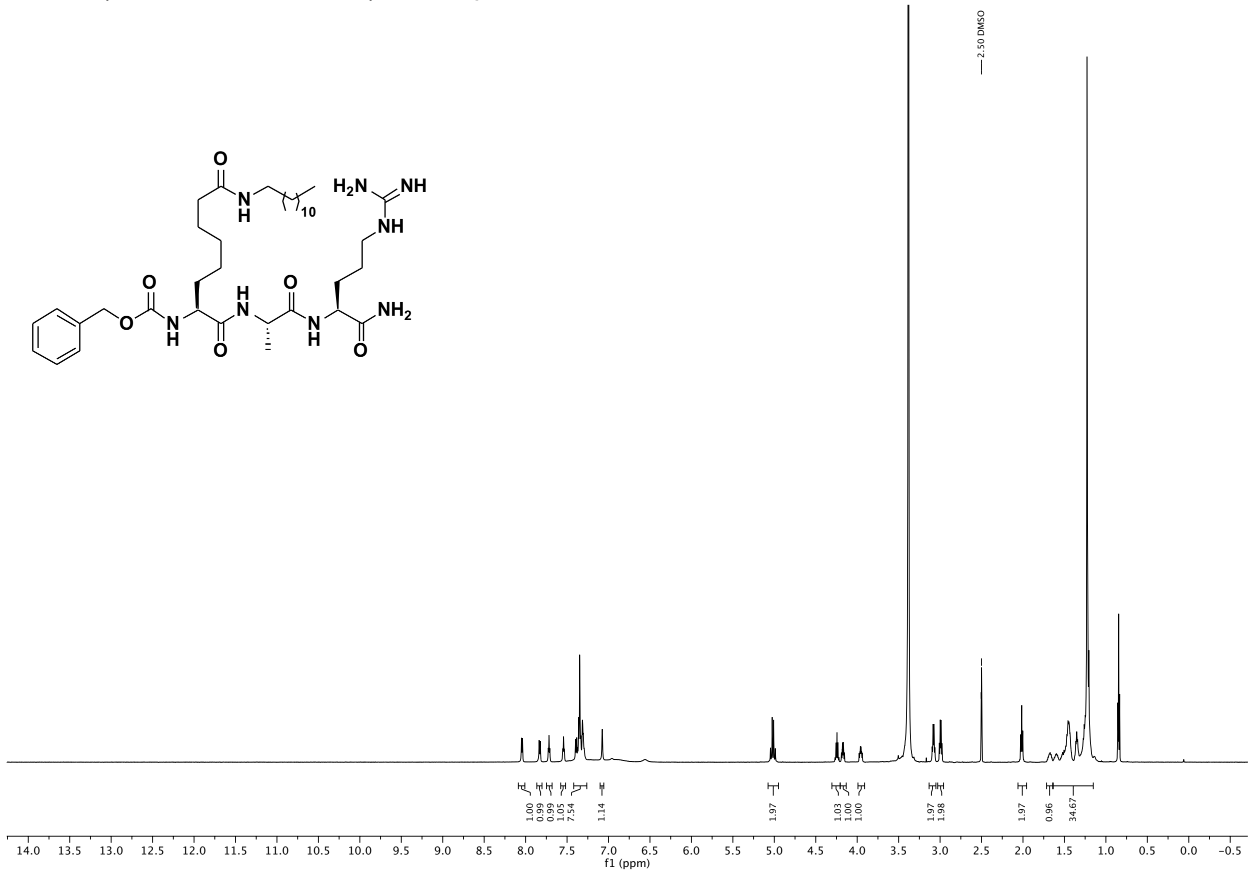

$^{13}C$ NMR (151 MHz, DMSO) of compound **6**.

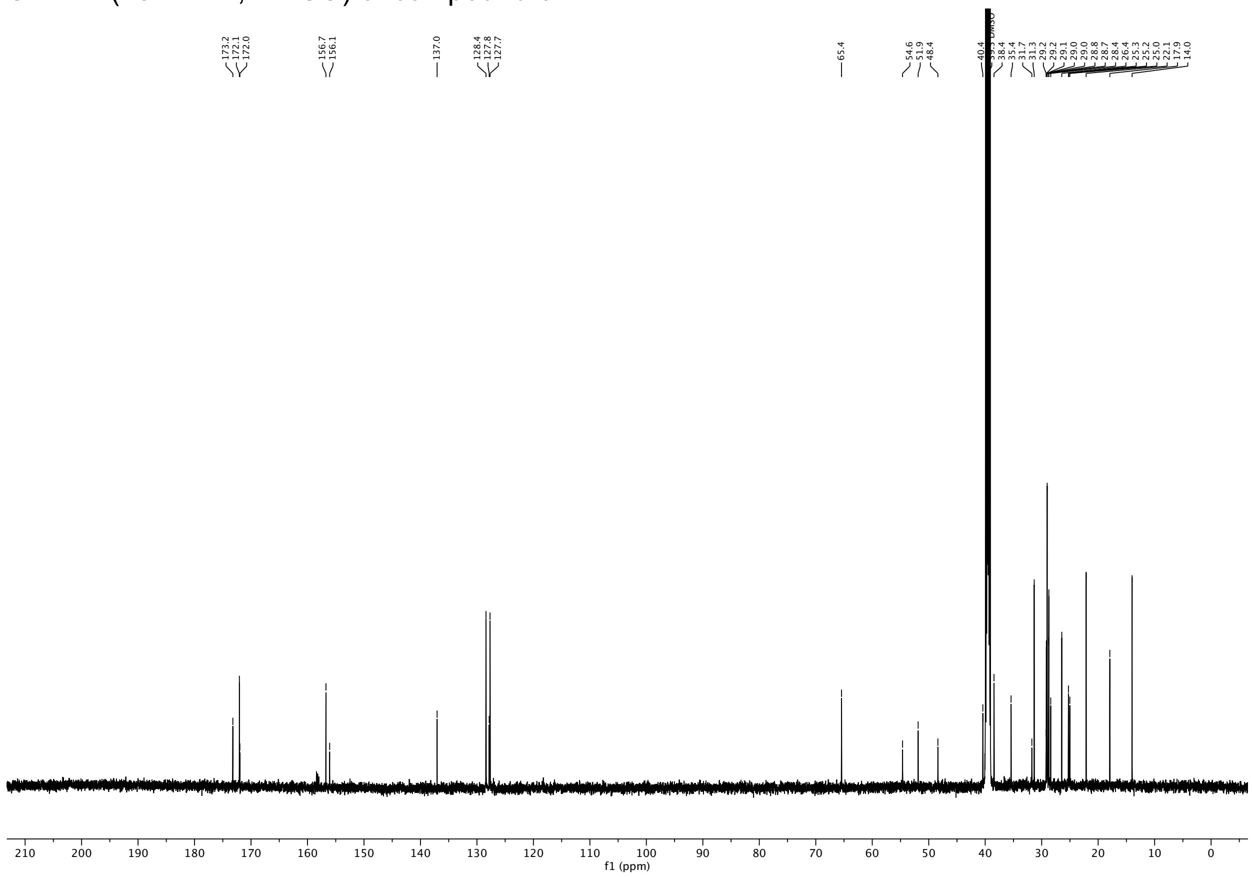

$^{1}$H NMR (600 MHz, DMSO-$d_6$) of compound **7**.

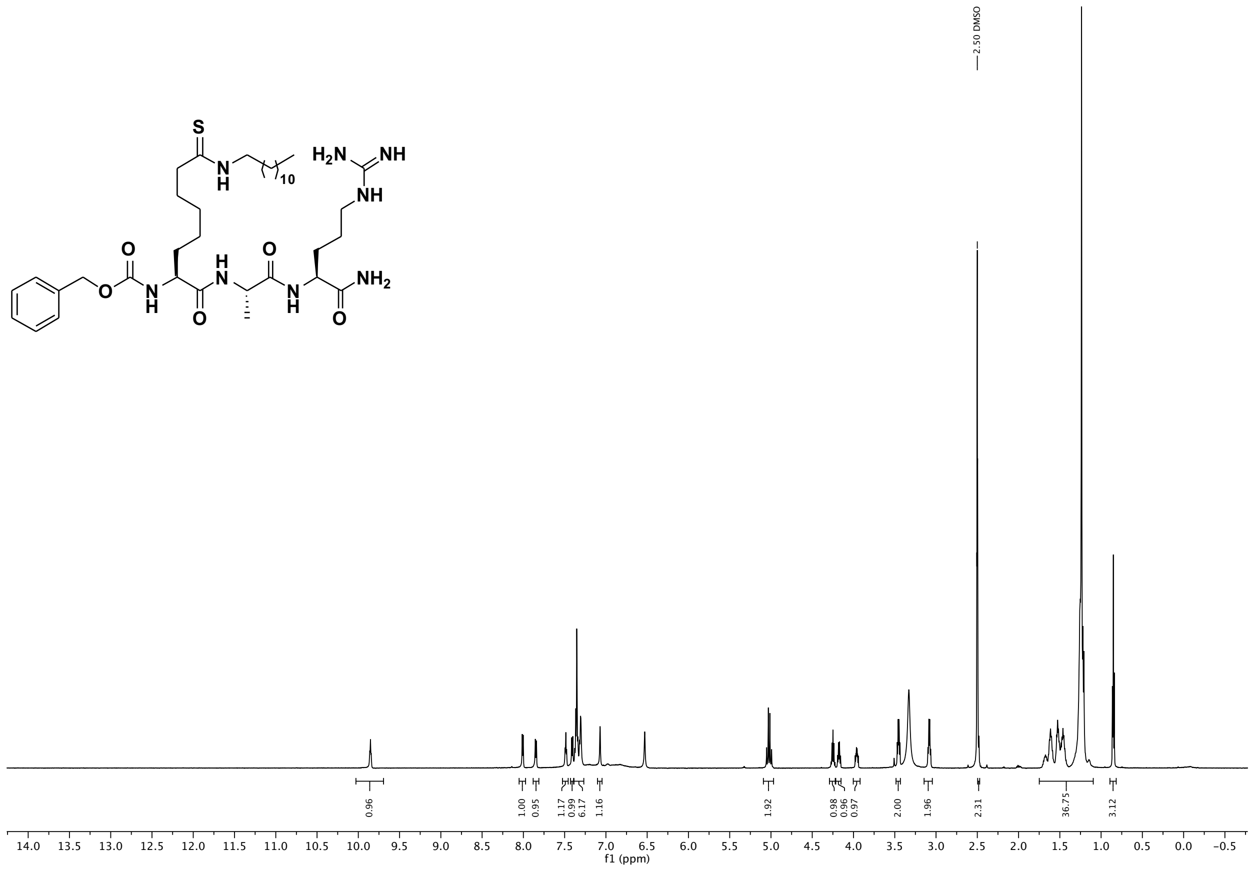

$^{13}$C NMR (151 MHz, DMSO) of compound **7**.

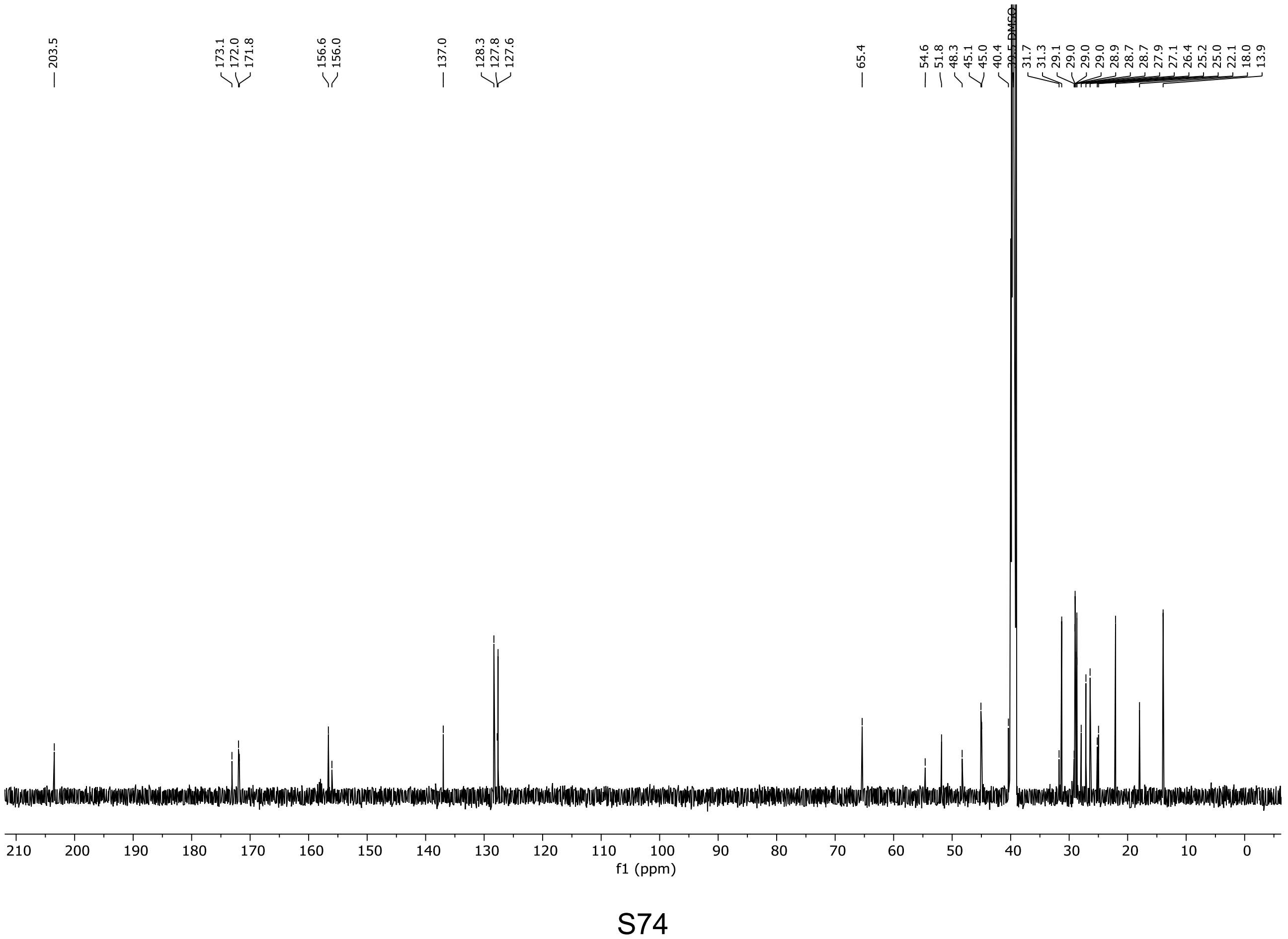

[1]H NMR (600 MHz, DMSO-$d_6$) of compound **8**.

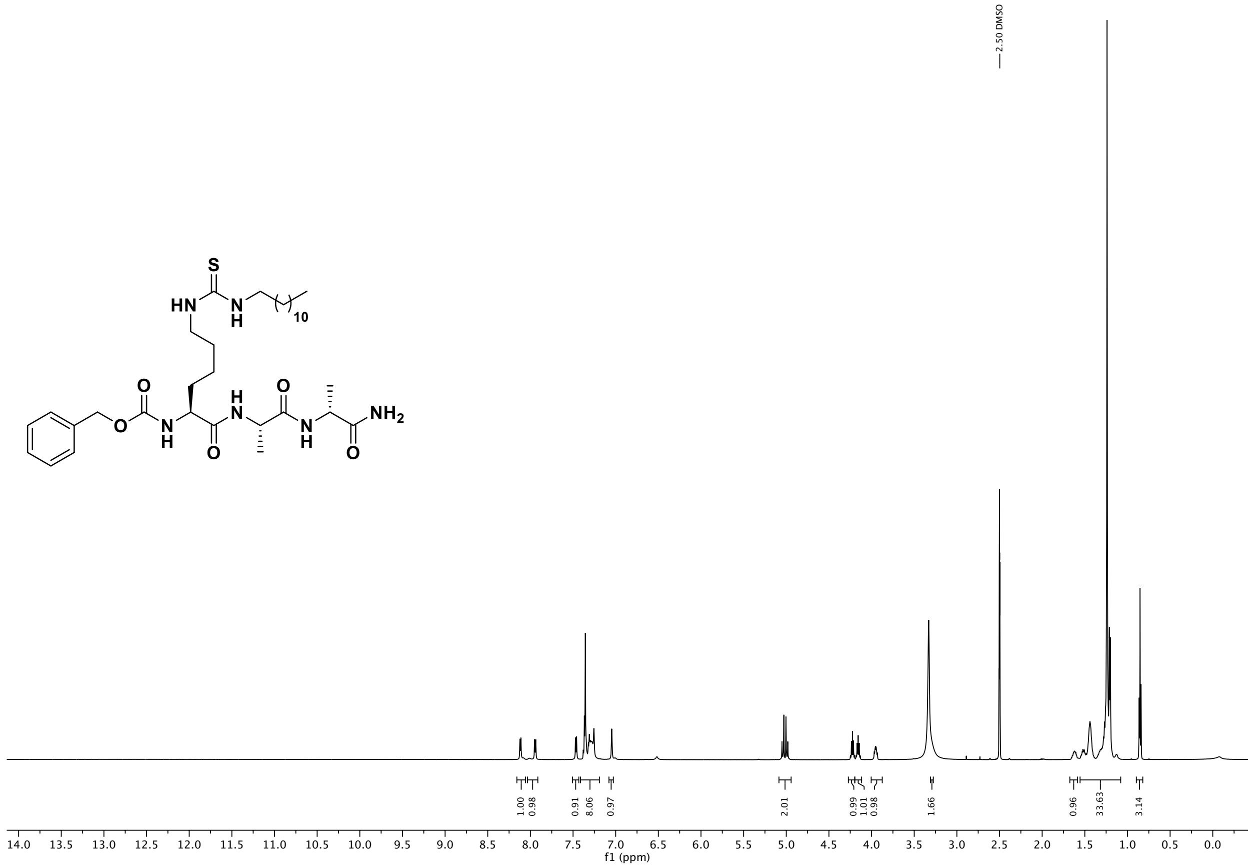

[13]C NMR (151 MHz, DMSO) of compound **8**.

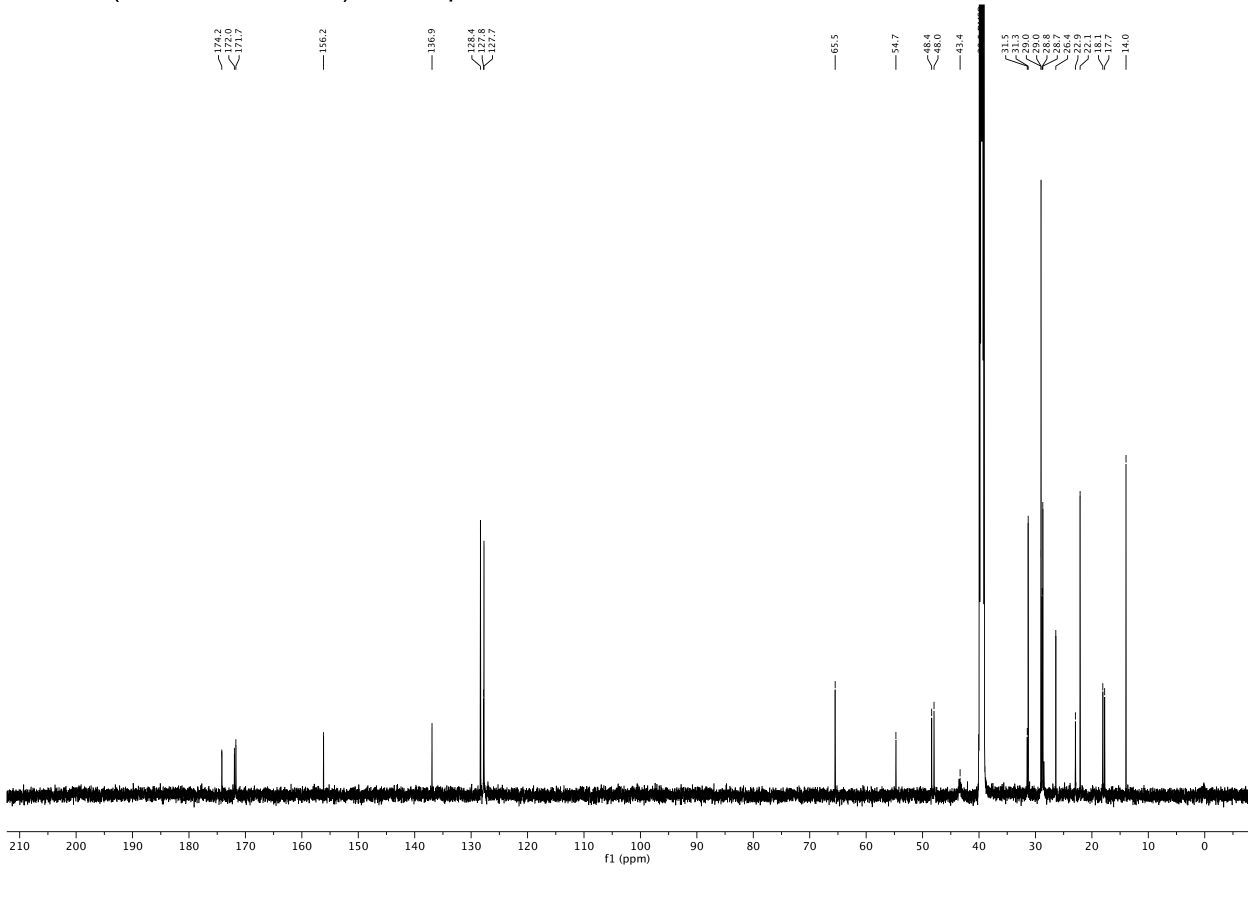

$^{1}H$ NMR (600 MHz, DMSO-$d_6$) of compound **9**.

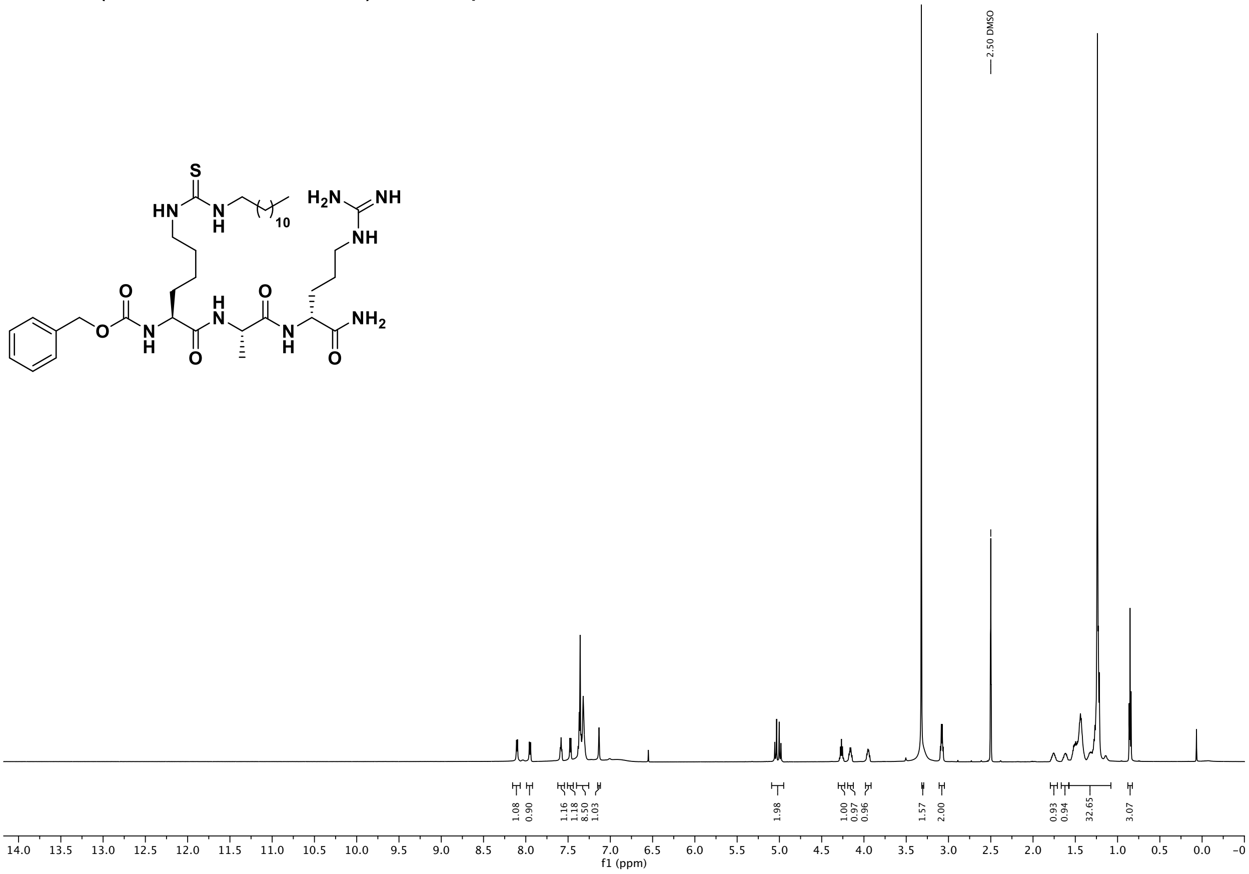

$^{13}C$ NMR (151 MHz, DMSO) of compound **9**.

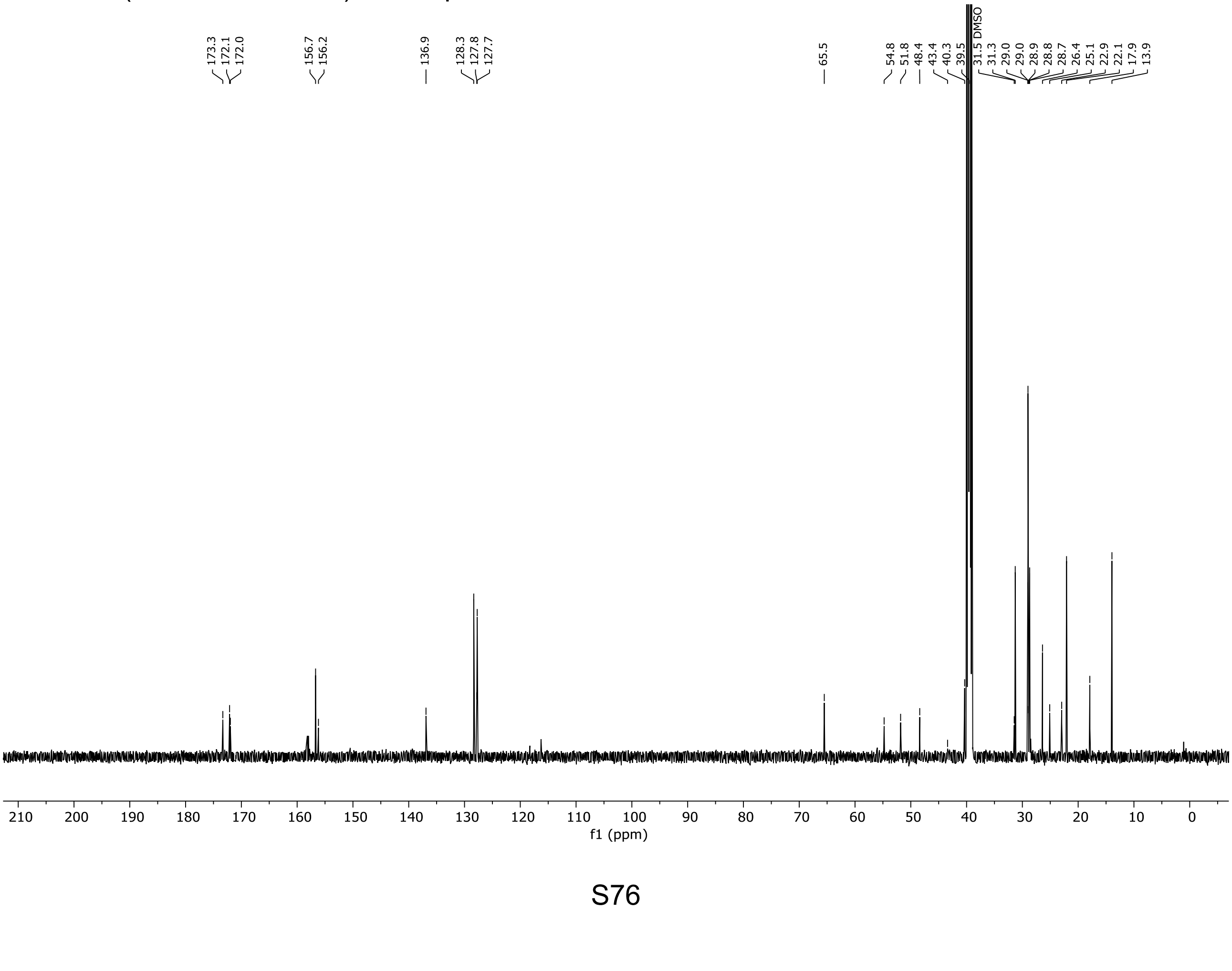

[1] H NMR (600 MHz, DMSO-$d_6$) of compound **10**.

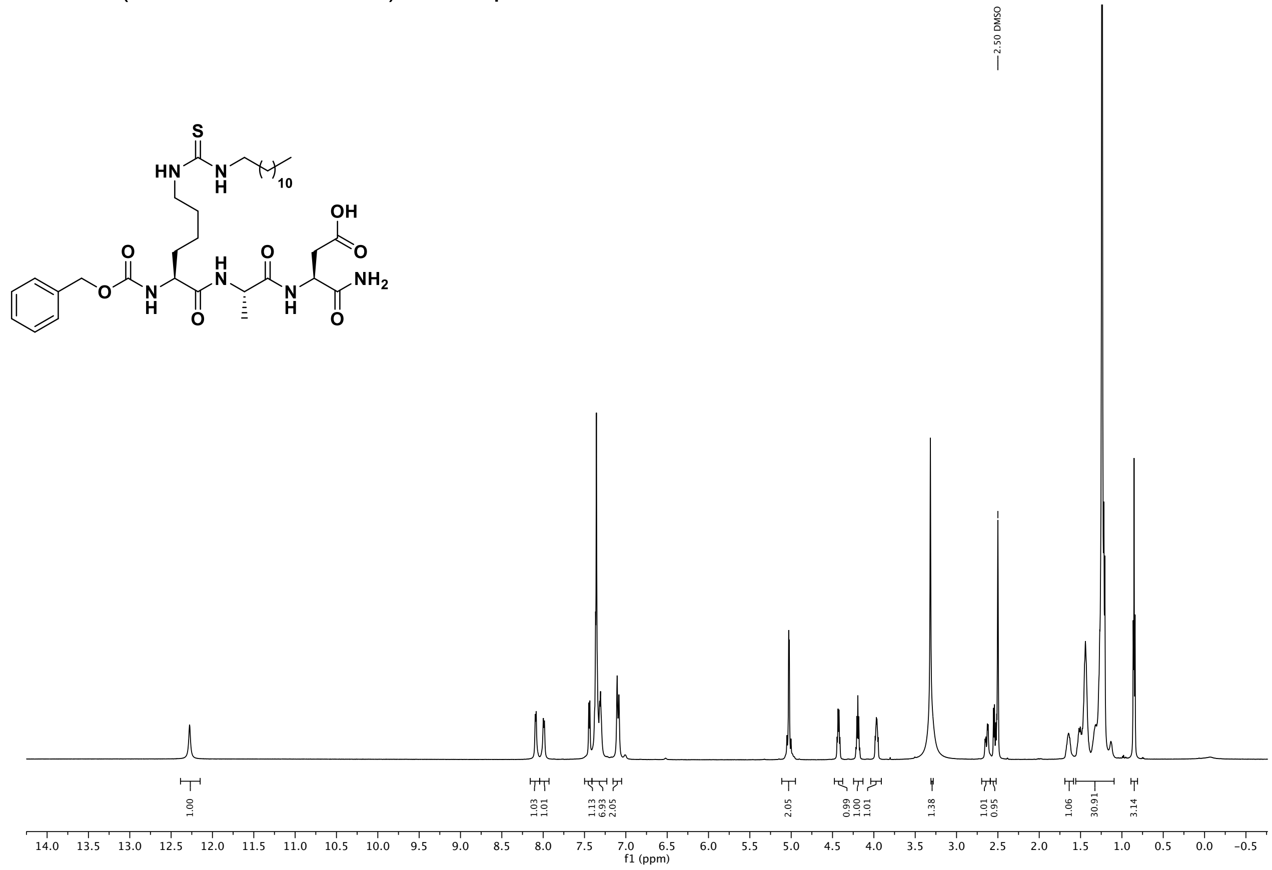

[13] C NMR (151 MHz, DMSO) of compound **10**.

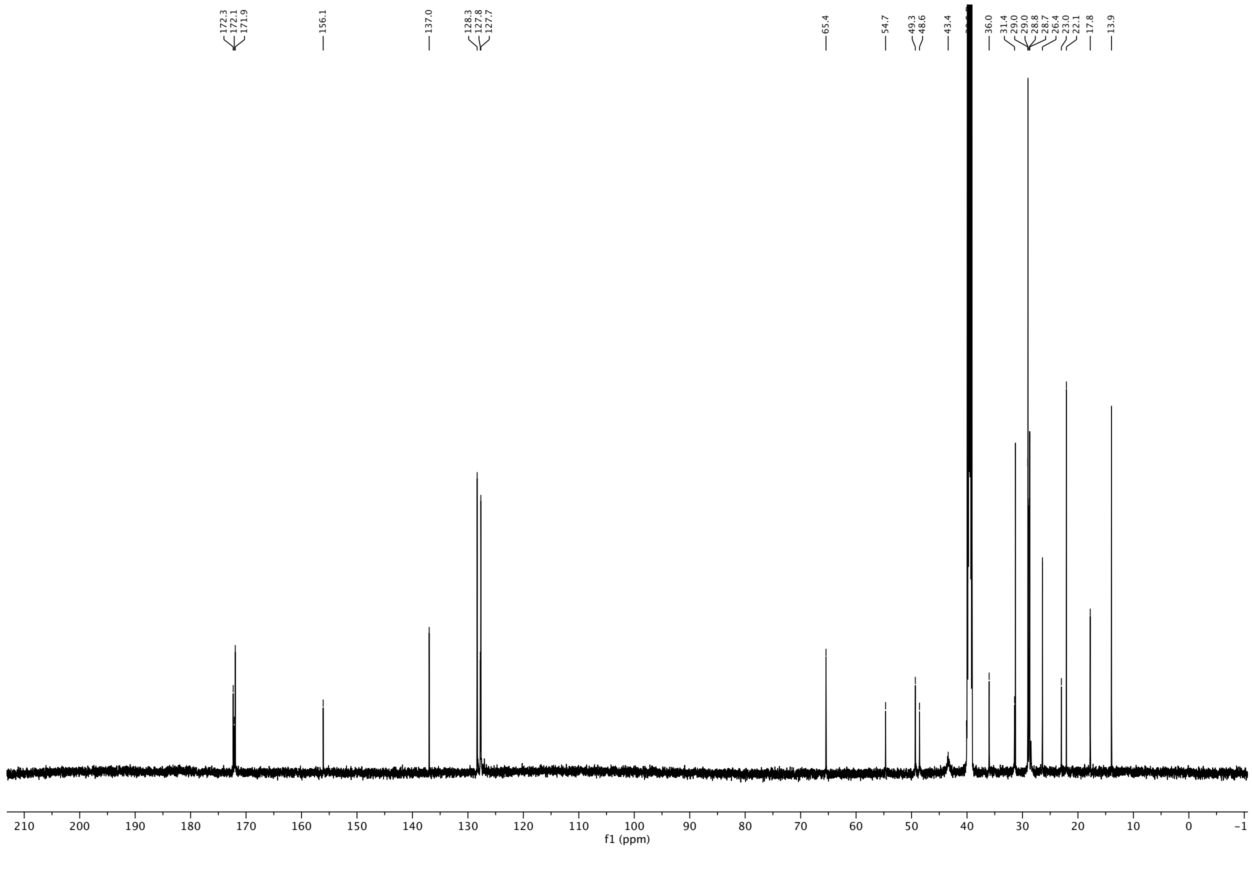

[1]H NMR (600 MHz, DMSO-$d_6$) of compound **11**.

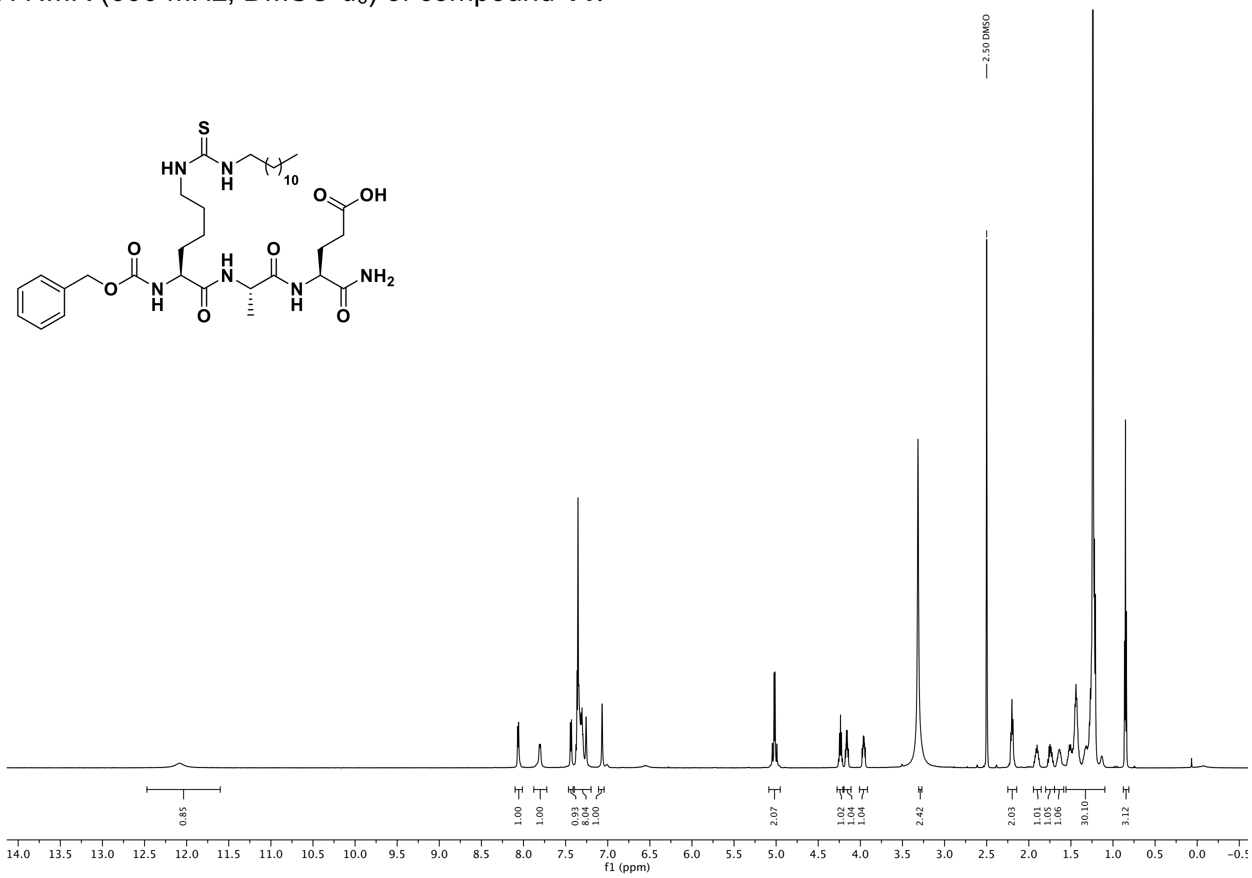

[13]C NMR (151 MHz, DMSO) of compound **11**.

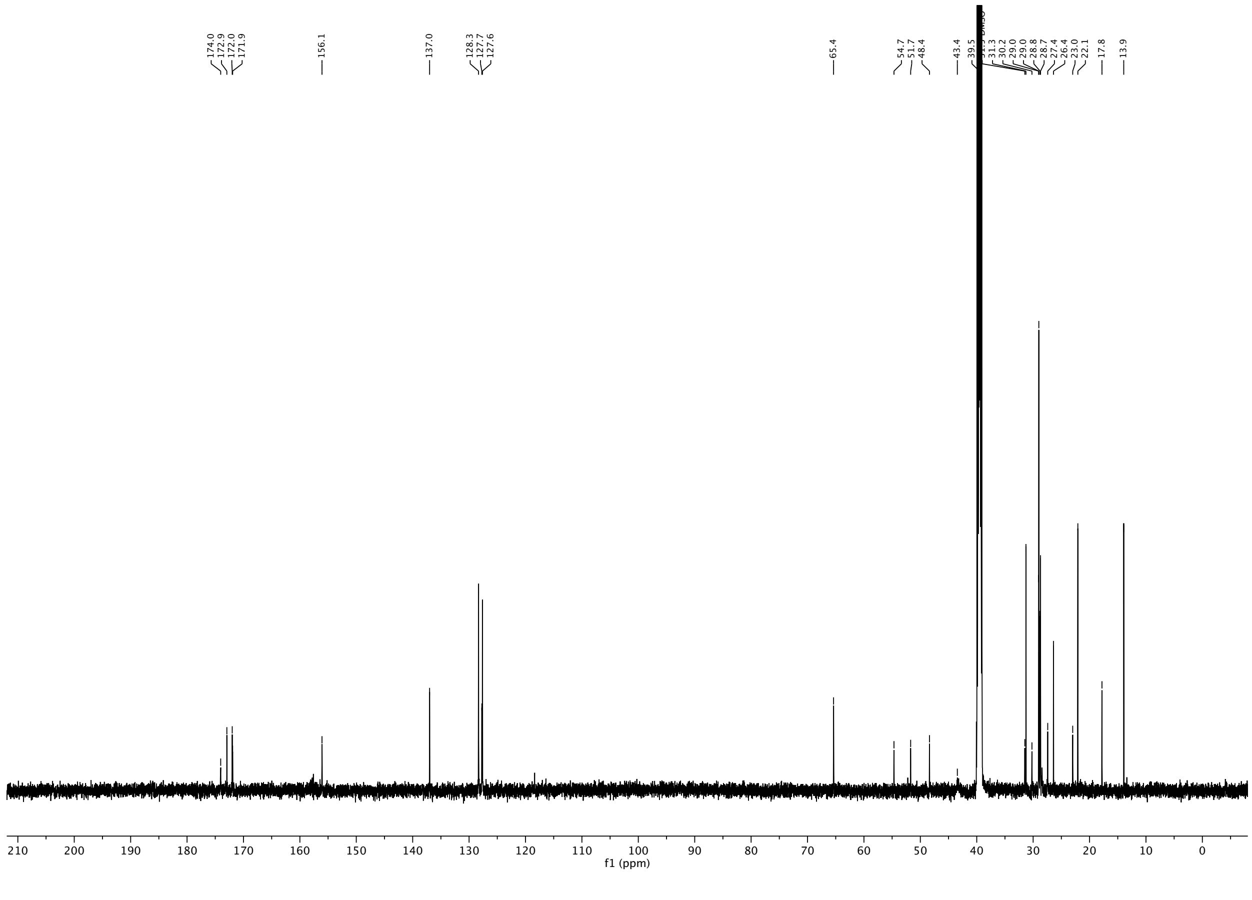

$^{1}H$ NMR (600 MHz, DMSO-$d_6$) of compound **12**.

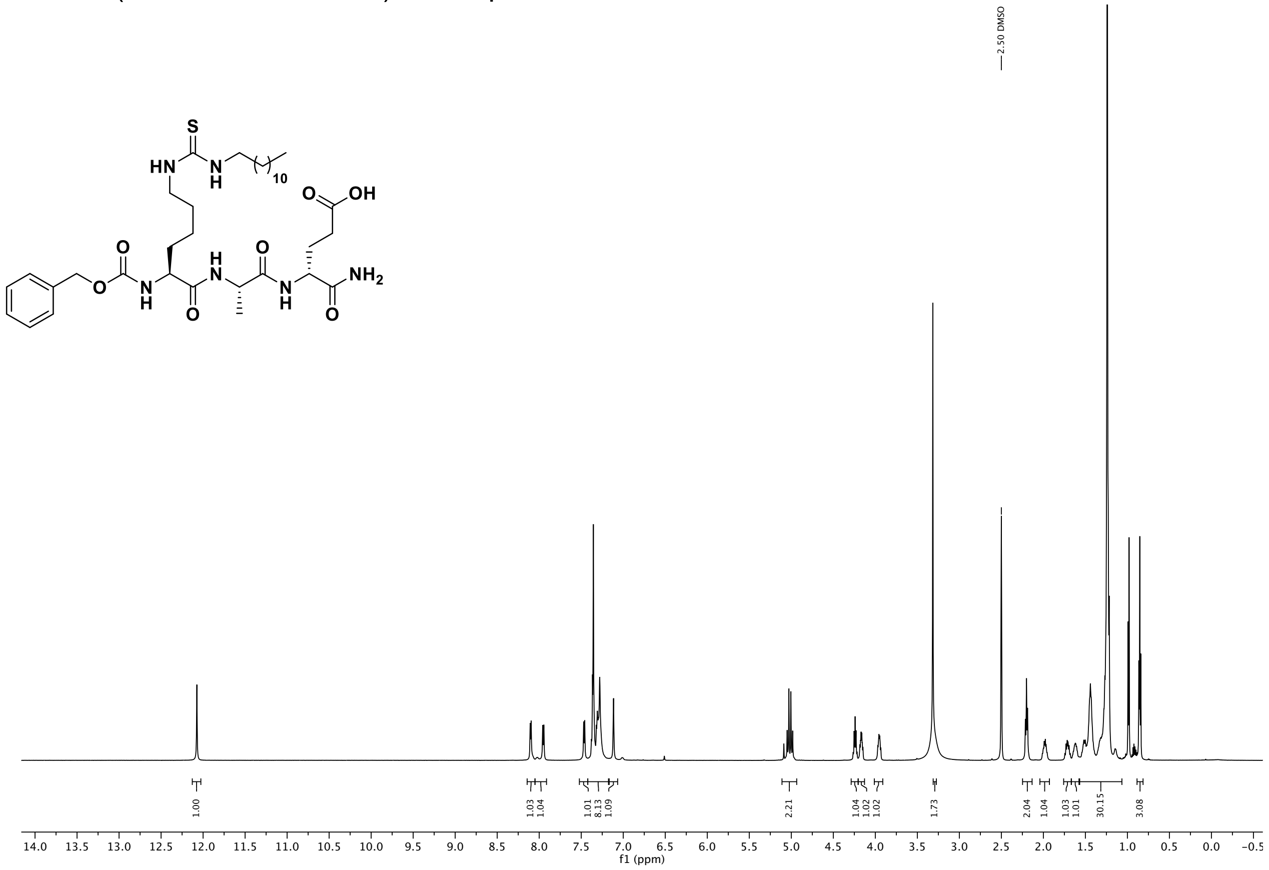

$^{13}C$ NMR (151 MHz, DMSO) of compound **12**.

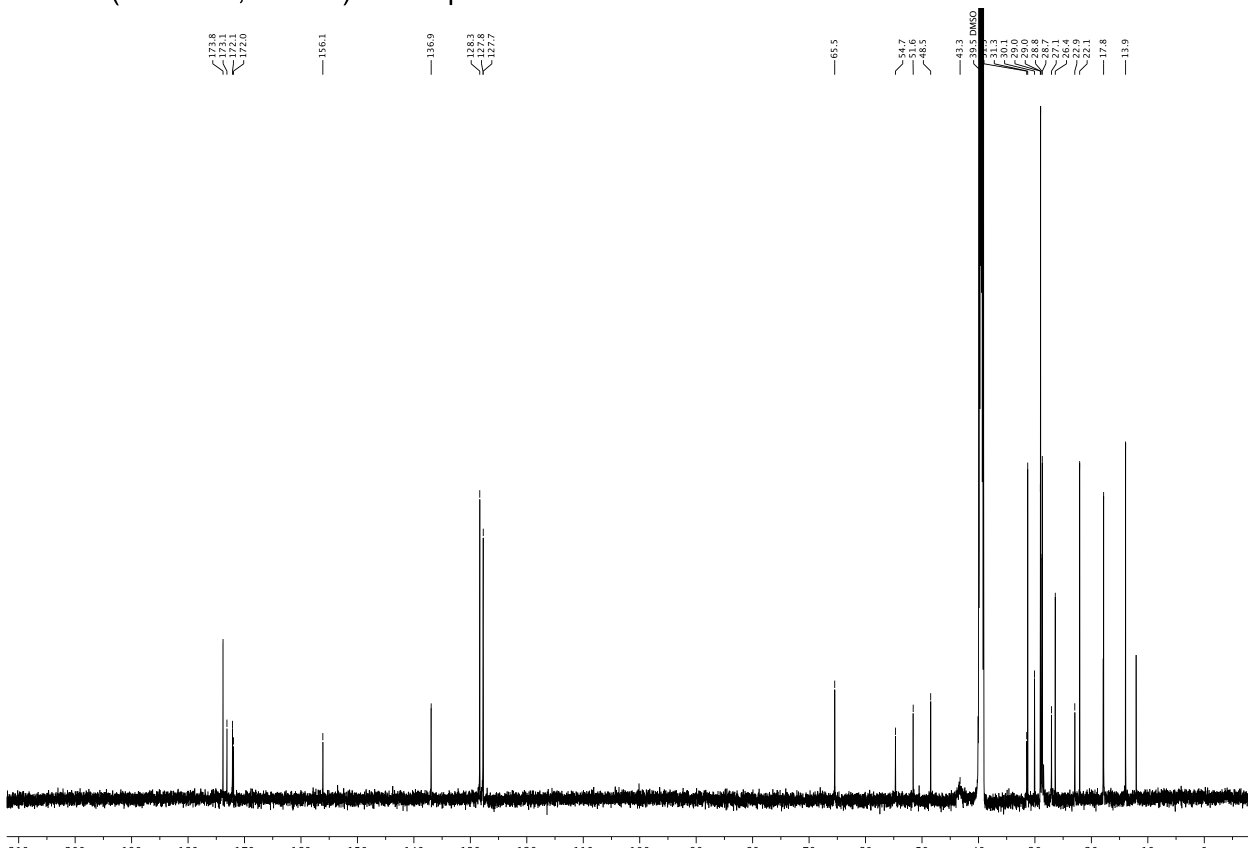

[1]H NMR (600 MHz, DMSO-$d_6$) of compound **13**.

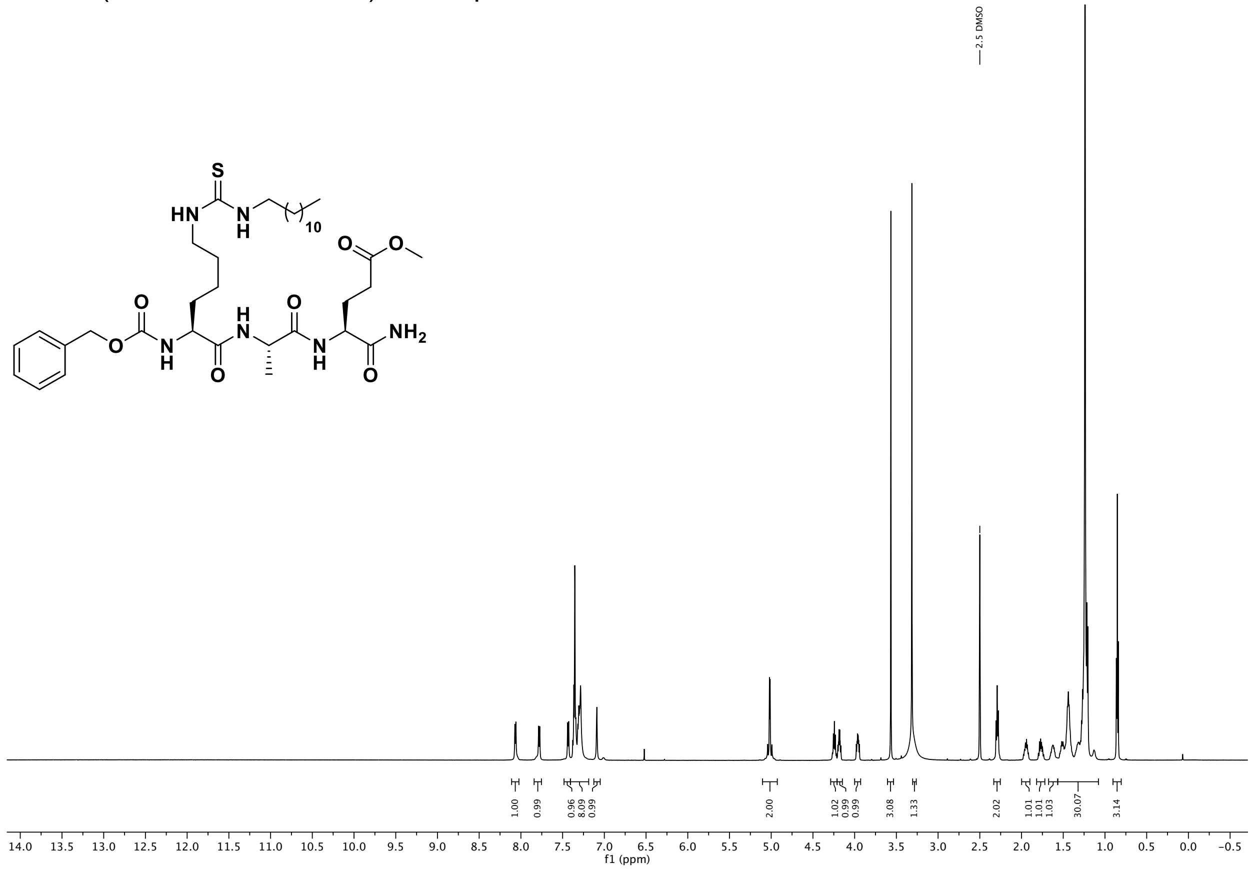

[13]C NMR (151 MHz, DMSO) of compound **13**.

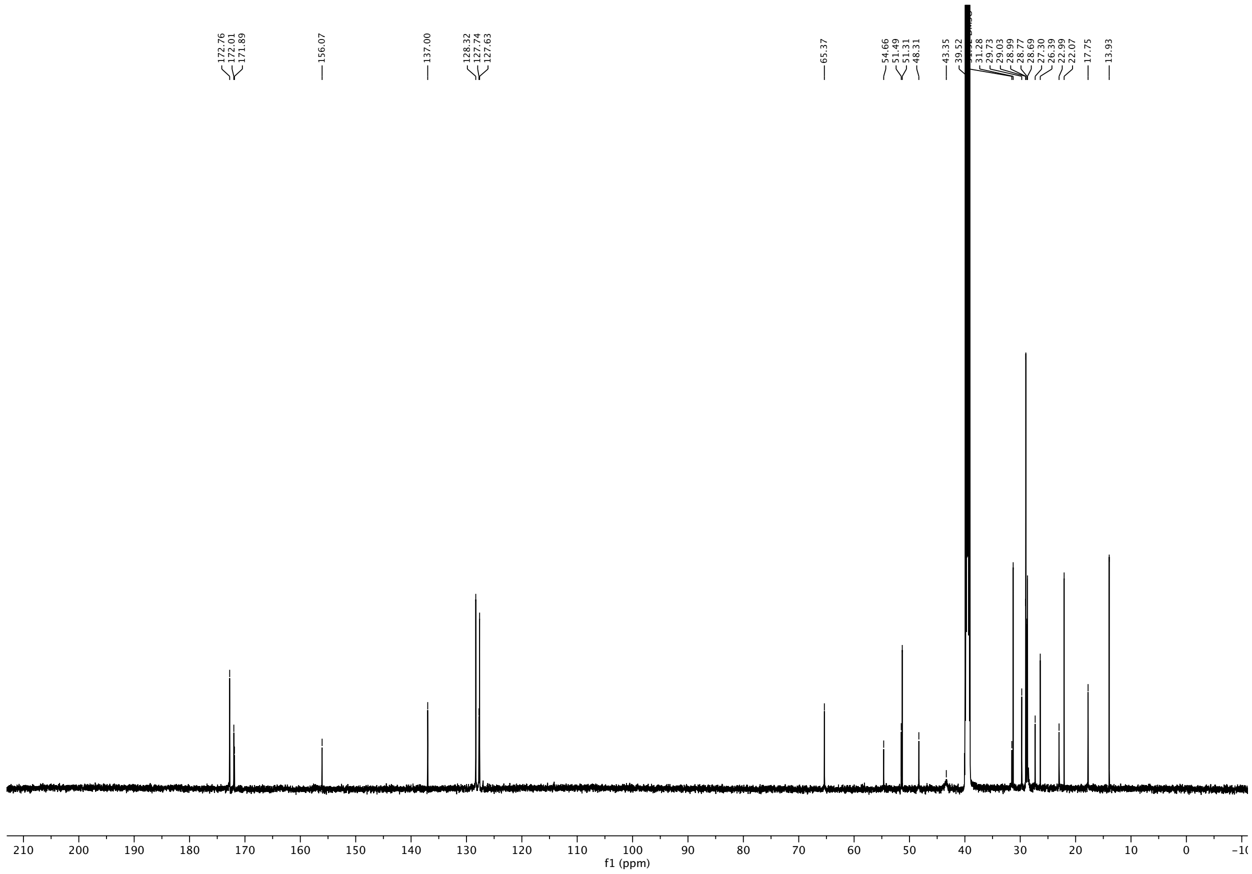

$^{1}$H NMR (600 MHz, DMSO-$d_6$) of compound **14**.

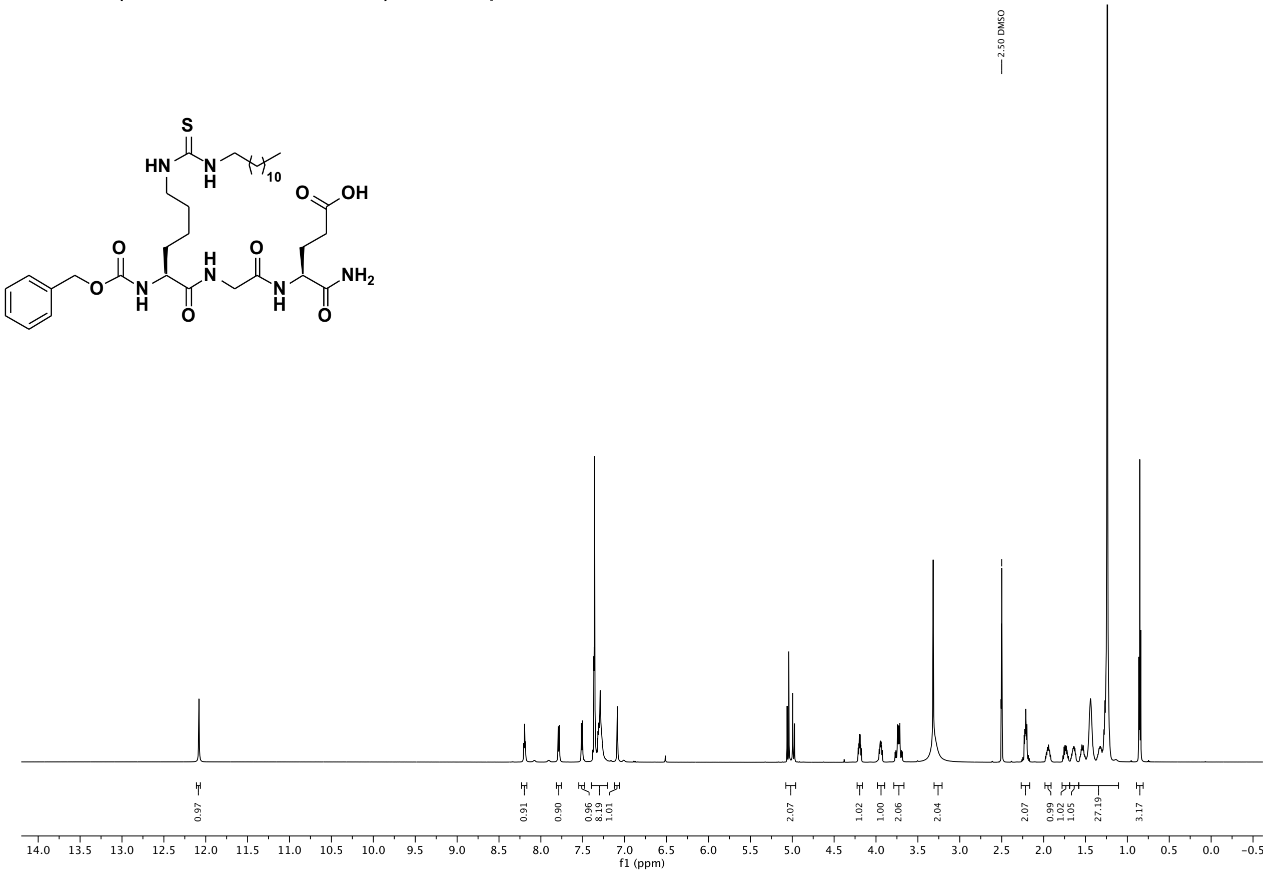

$^{13}$C NMR (151 MHz, DMSO) of compound **14**.

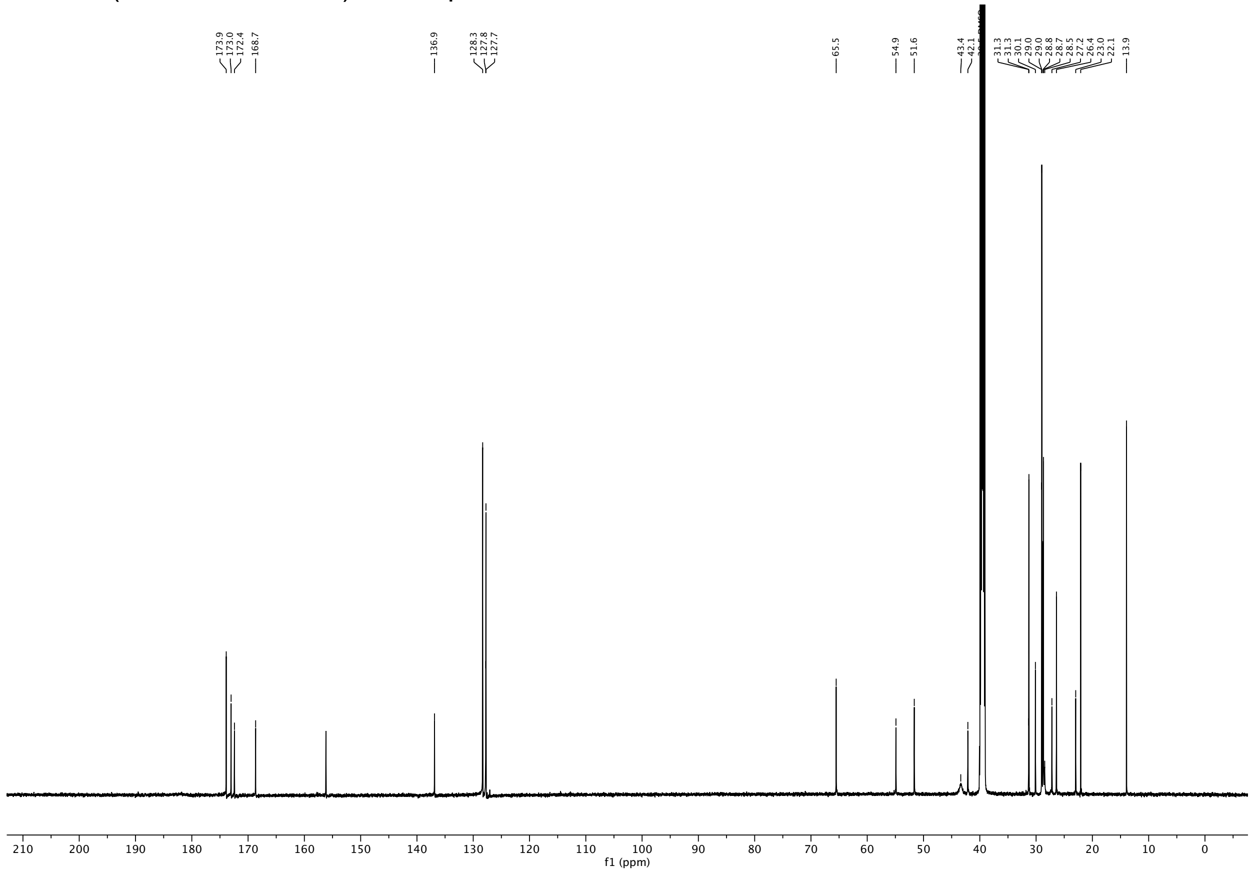

[1]H NMR (600 MHz, DMSO-$d_6$) of compound **15**.

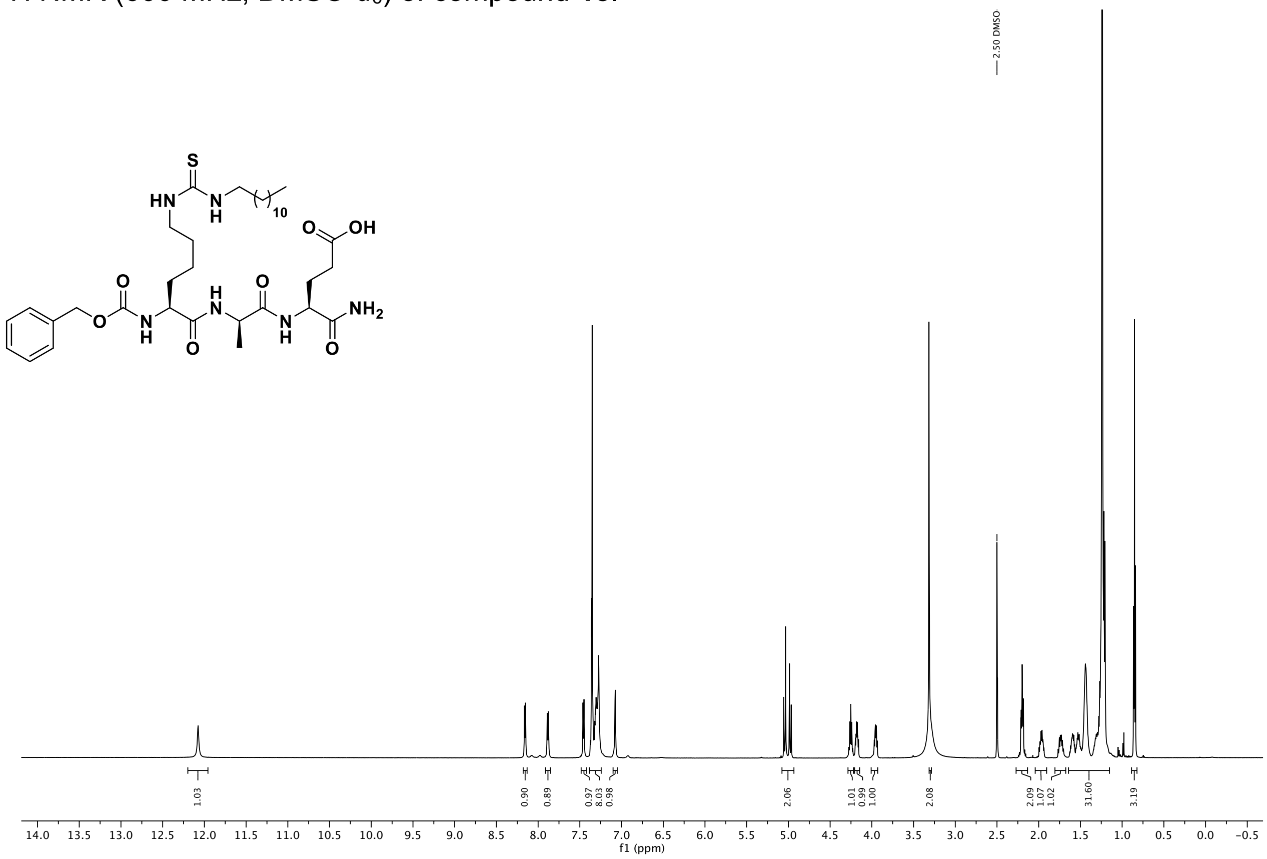

[13]C NMR (151 MHz, DMSO) of compound **15**.

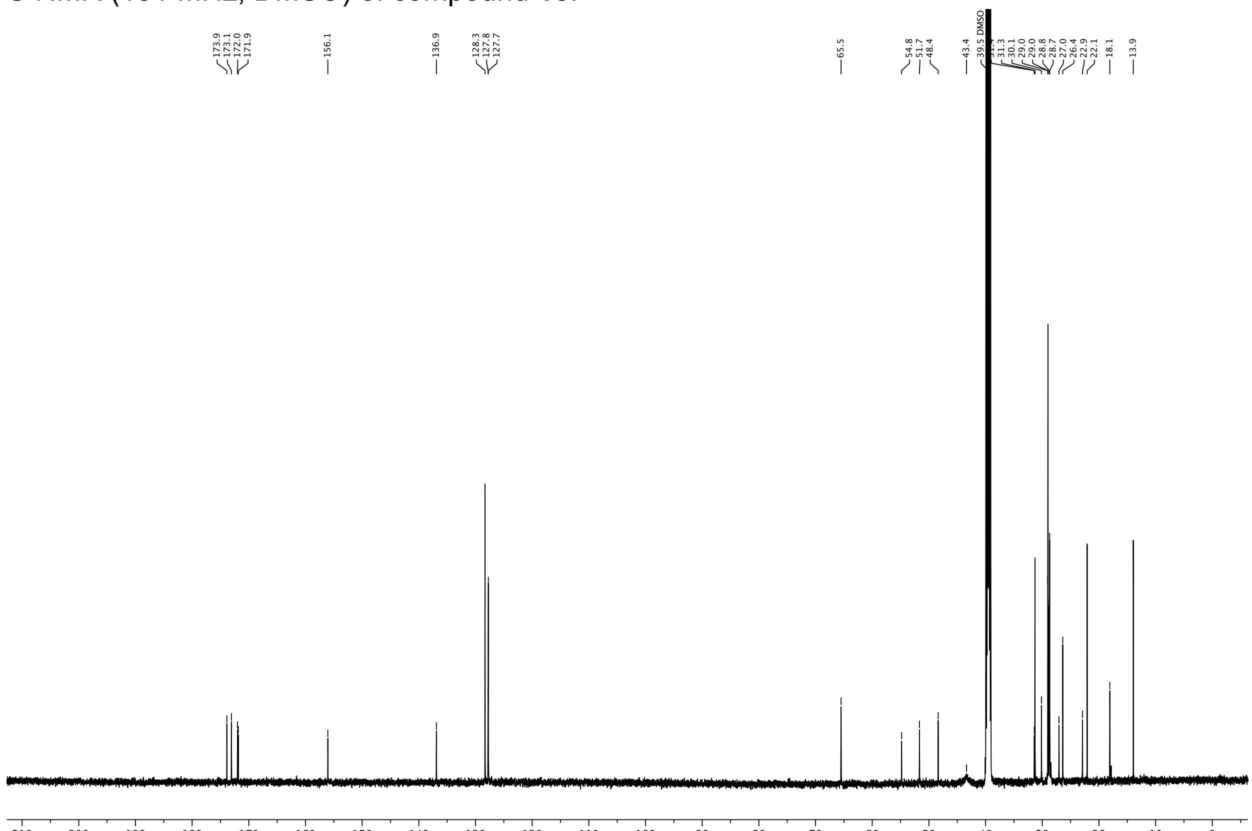

$^{1}H$ NMR (600 MHz, DMSO-$d_6$) of compound **16**.

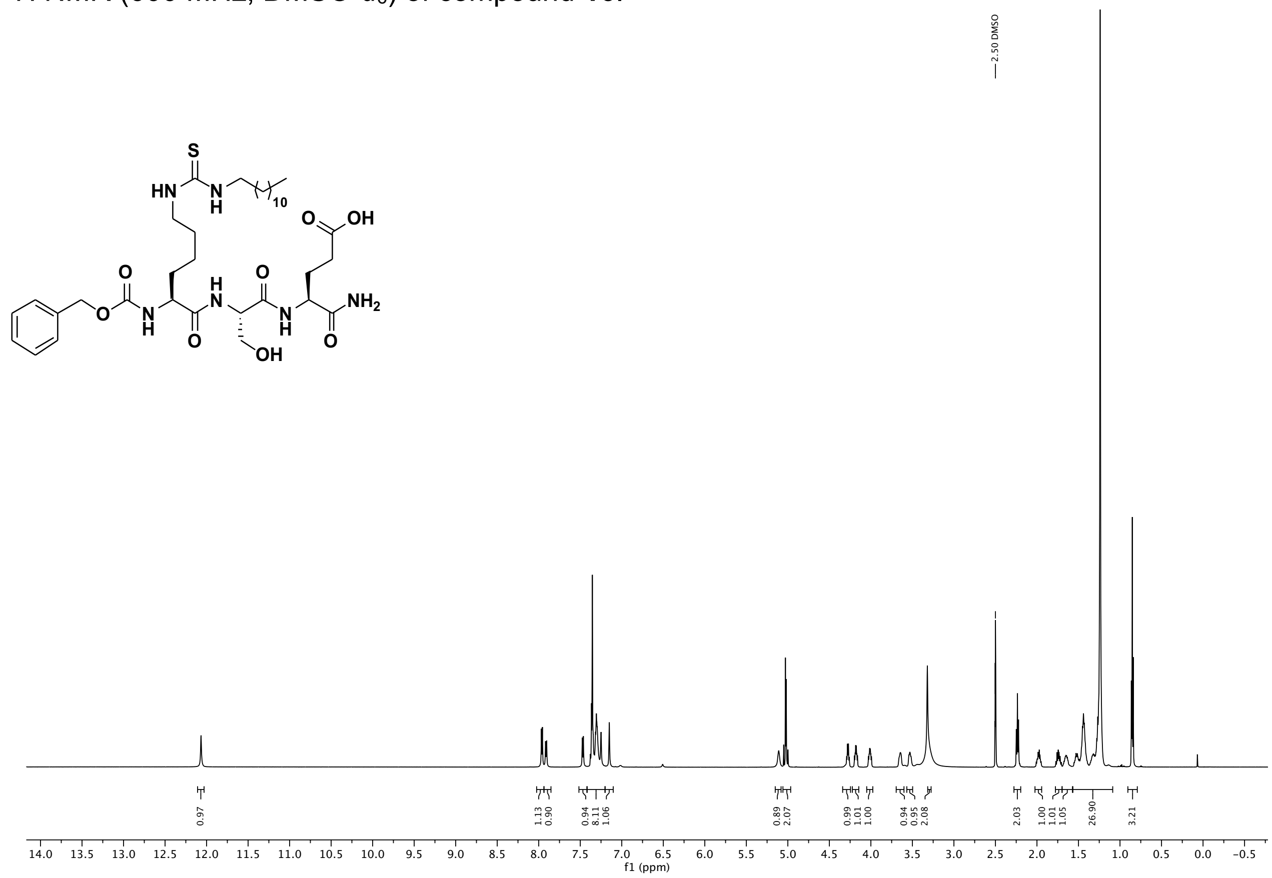

$^{13}C$ NMR (151 MHz, DMSO) of compound **16**.

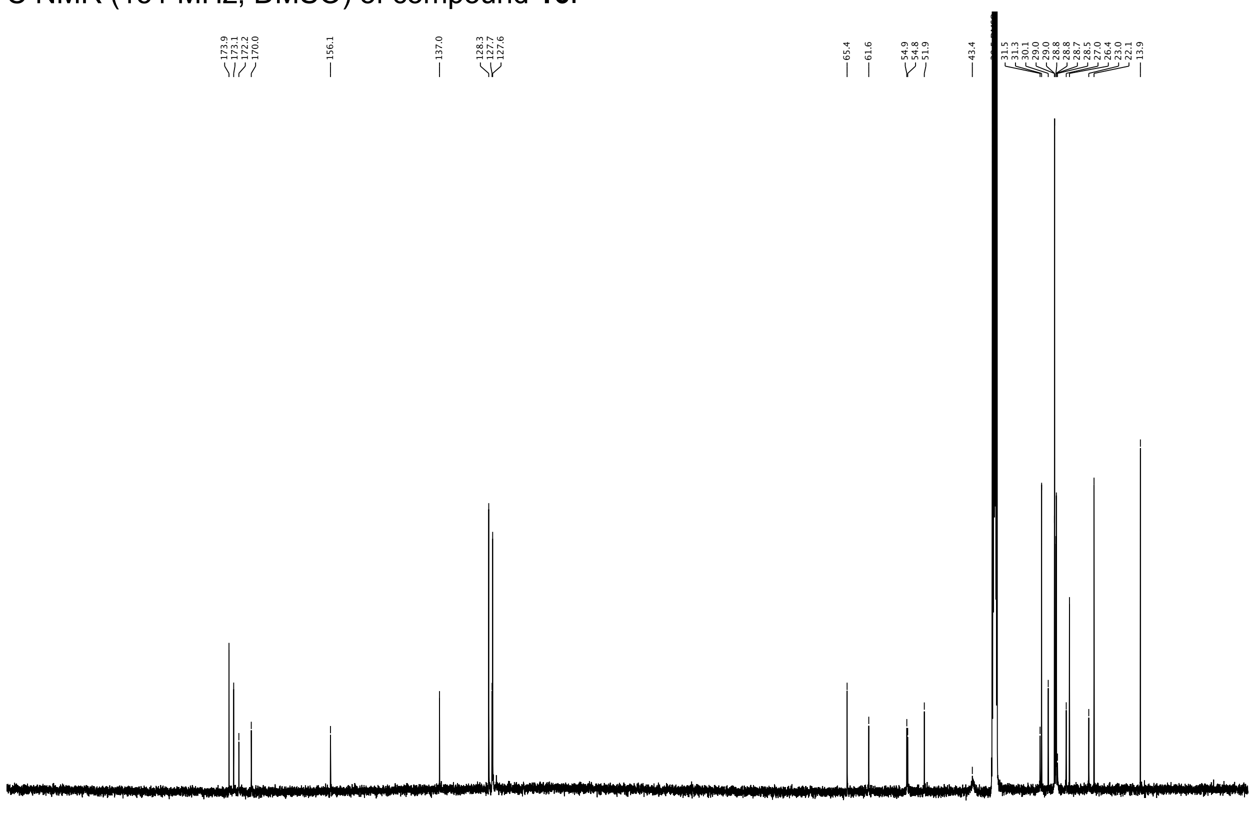

$^{1}$H NMR (600 MHz, DMSO-$d_6$) of compound **17**.

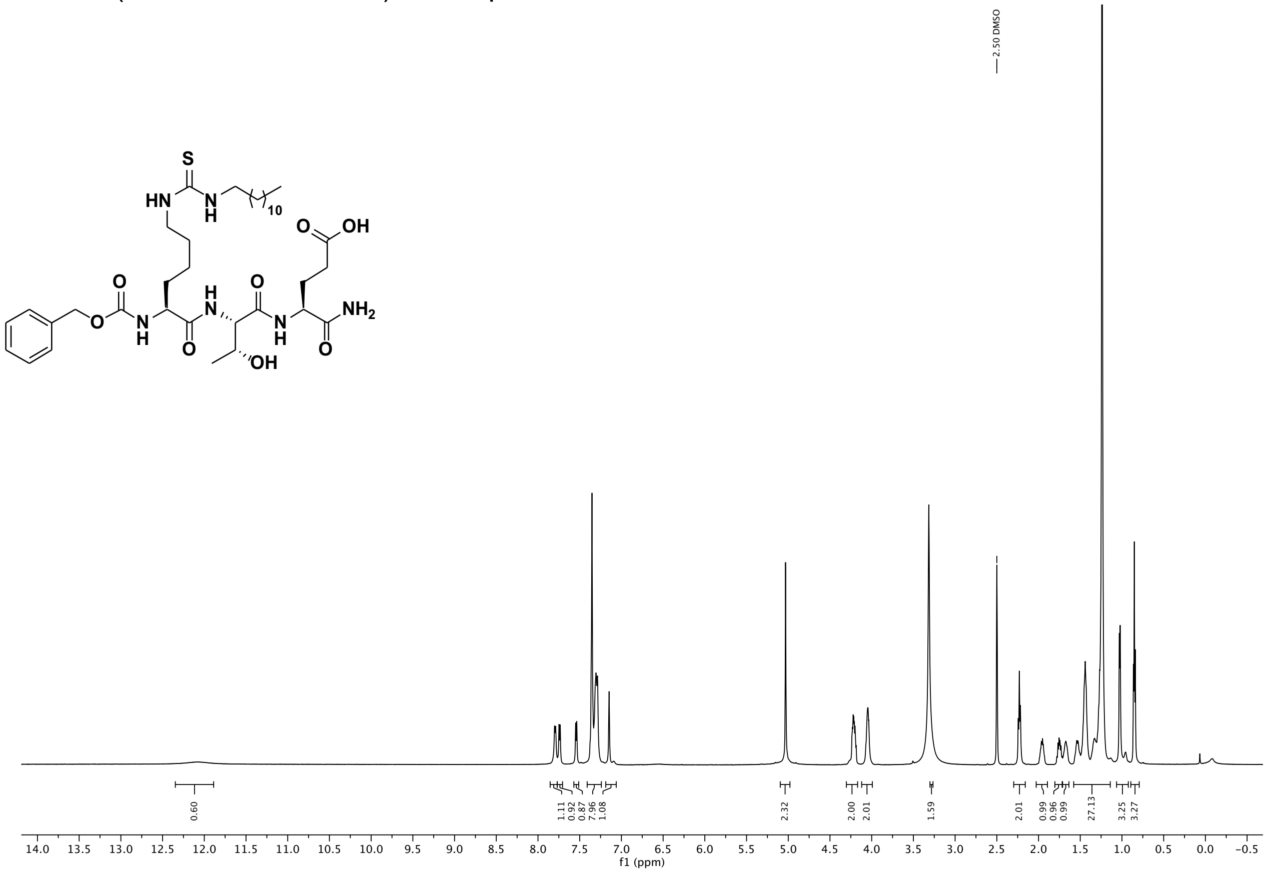

$^{13}$C NMR (151 MHz, DMSO) of compound **17**.

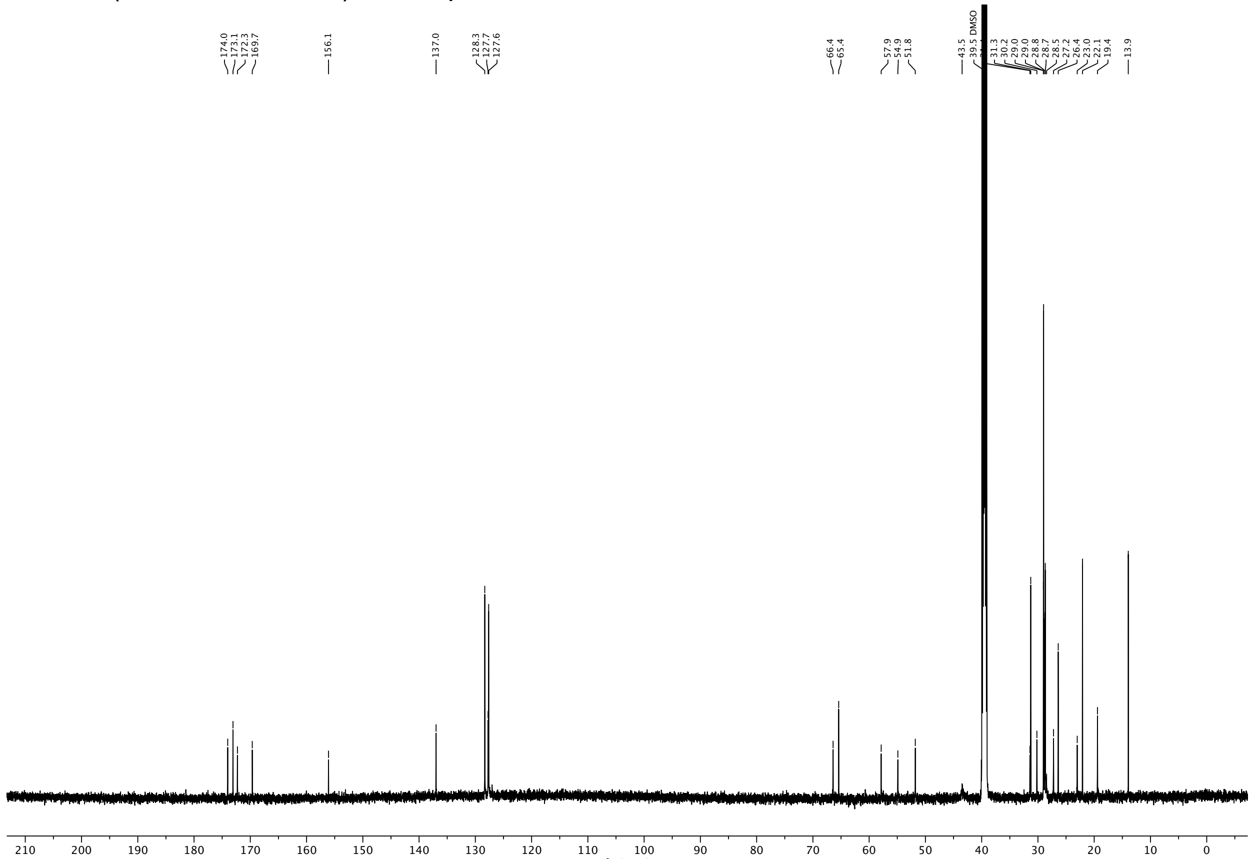

[1]H NMR (600 MHz, DMSO-$d_6$) of compound **18**.

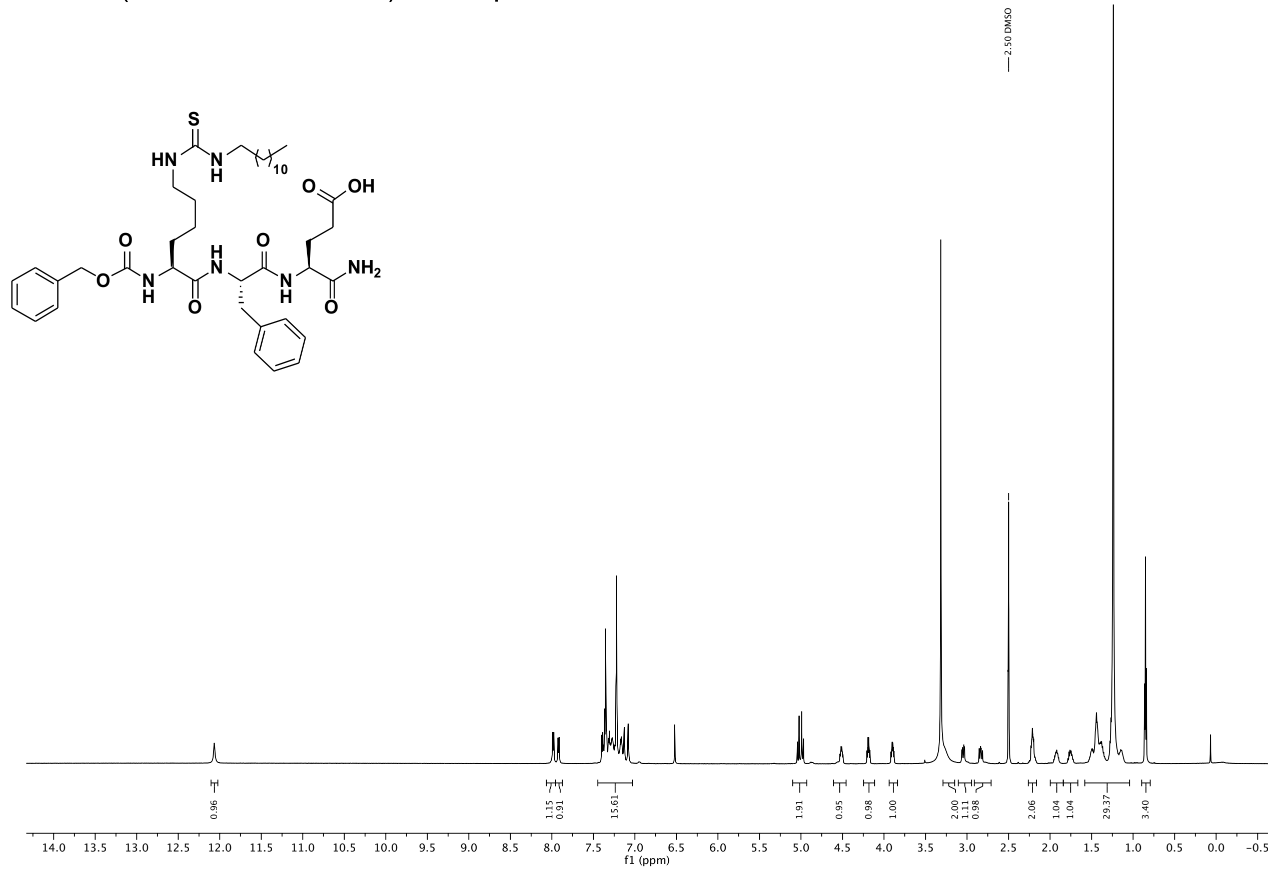

[13]C NMR (151 MHz, DMSO) of compound **18**.

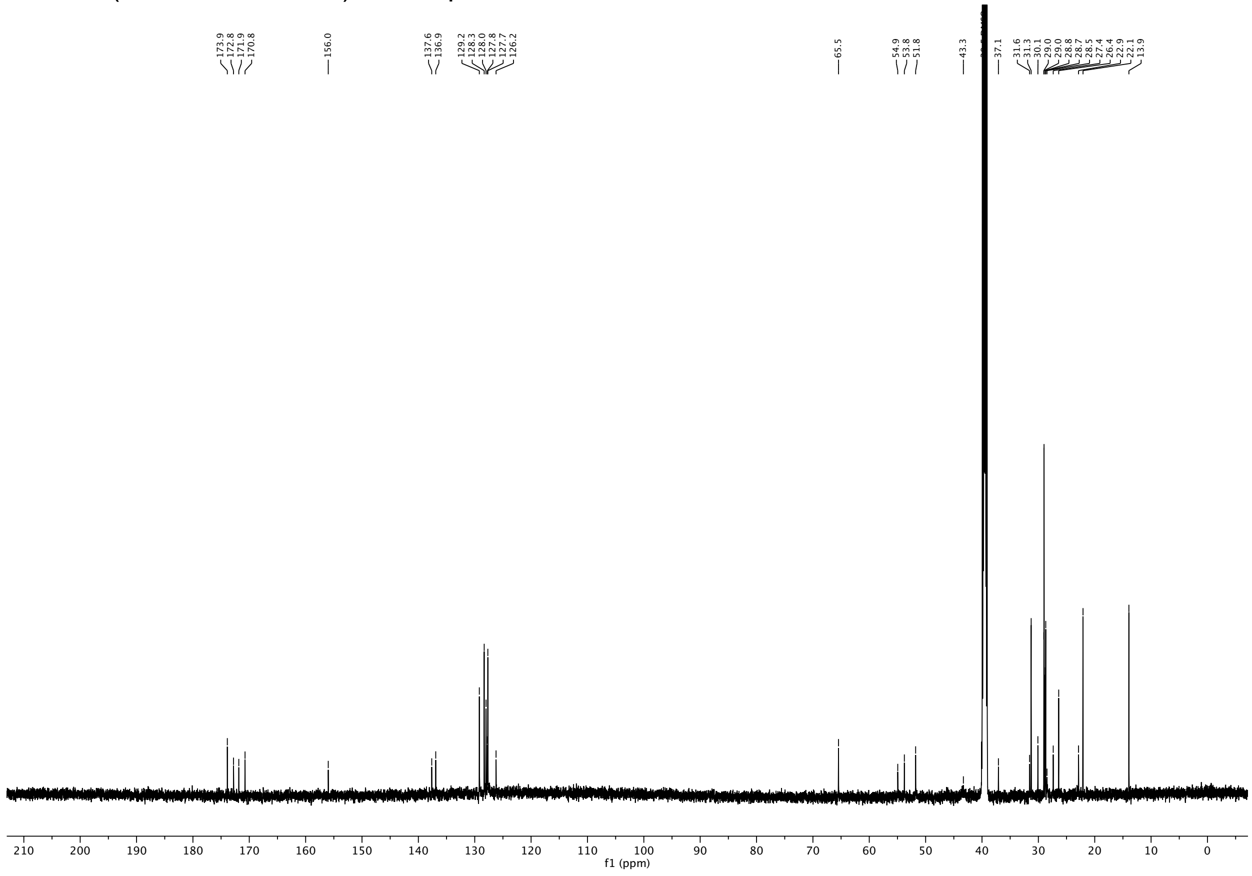

[1]H NMR (600 MHz, DMSO-$d_6$) of compound **19**.

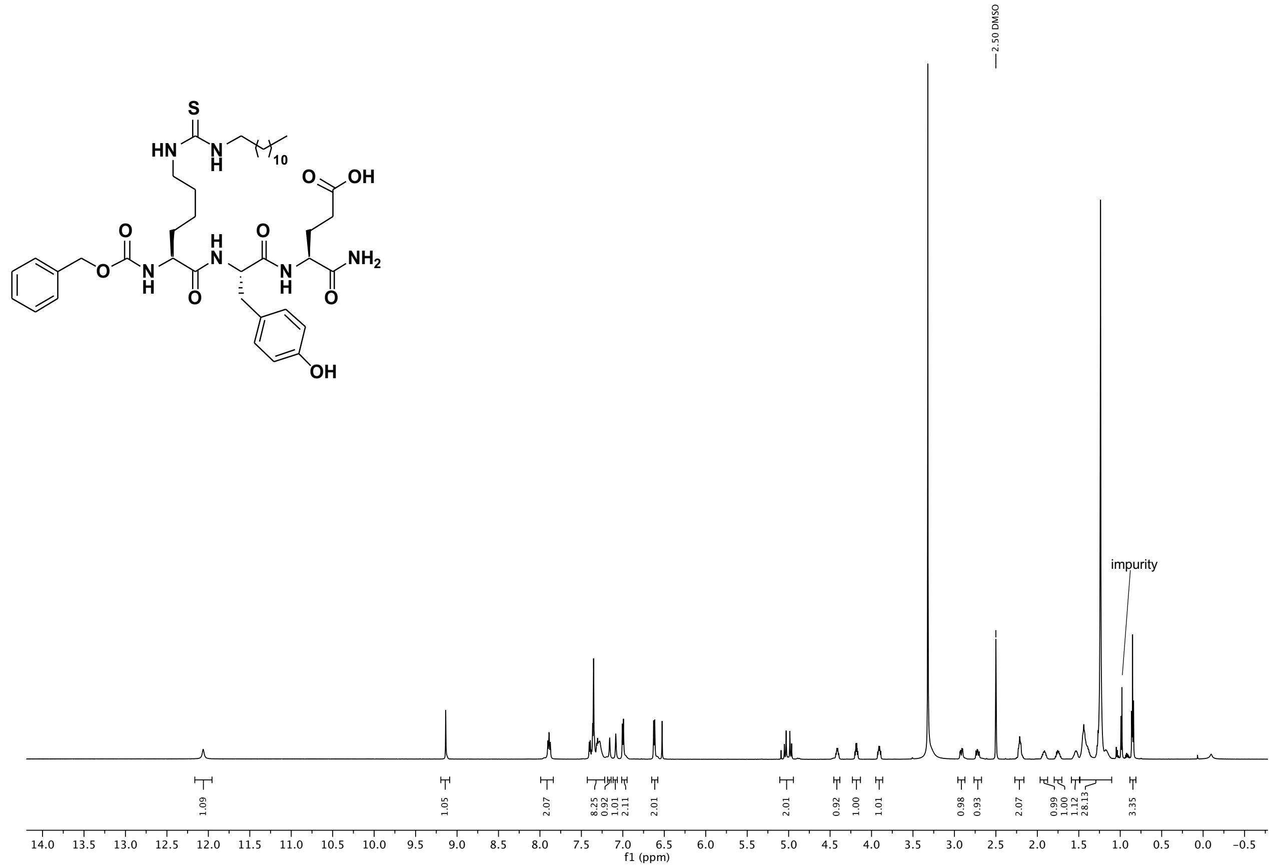

[13]C NMR (151 MHz, DMSO) of compound **19**.

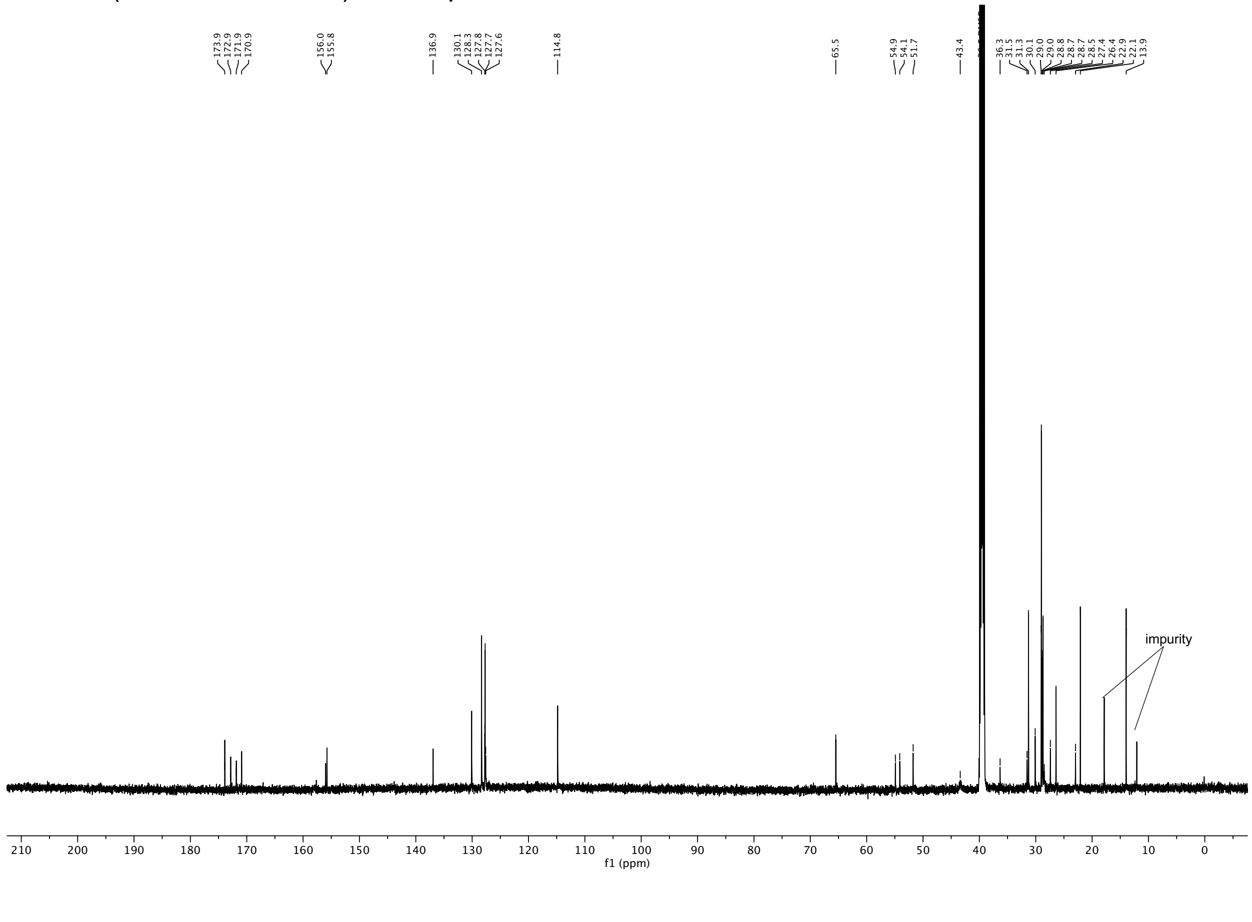

$^{1}H$ NMR (600 MHz, DMSO-$d_6$) of compound **20**.

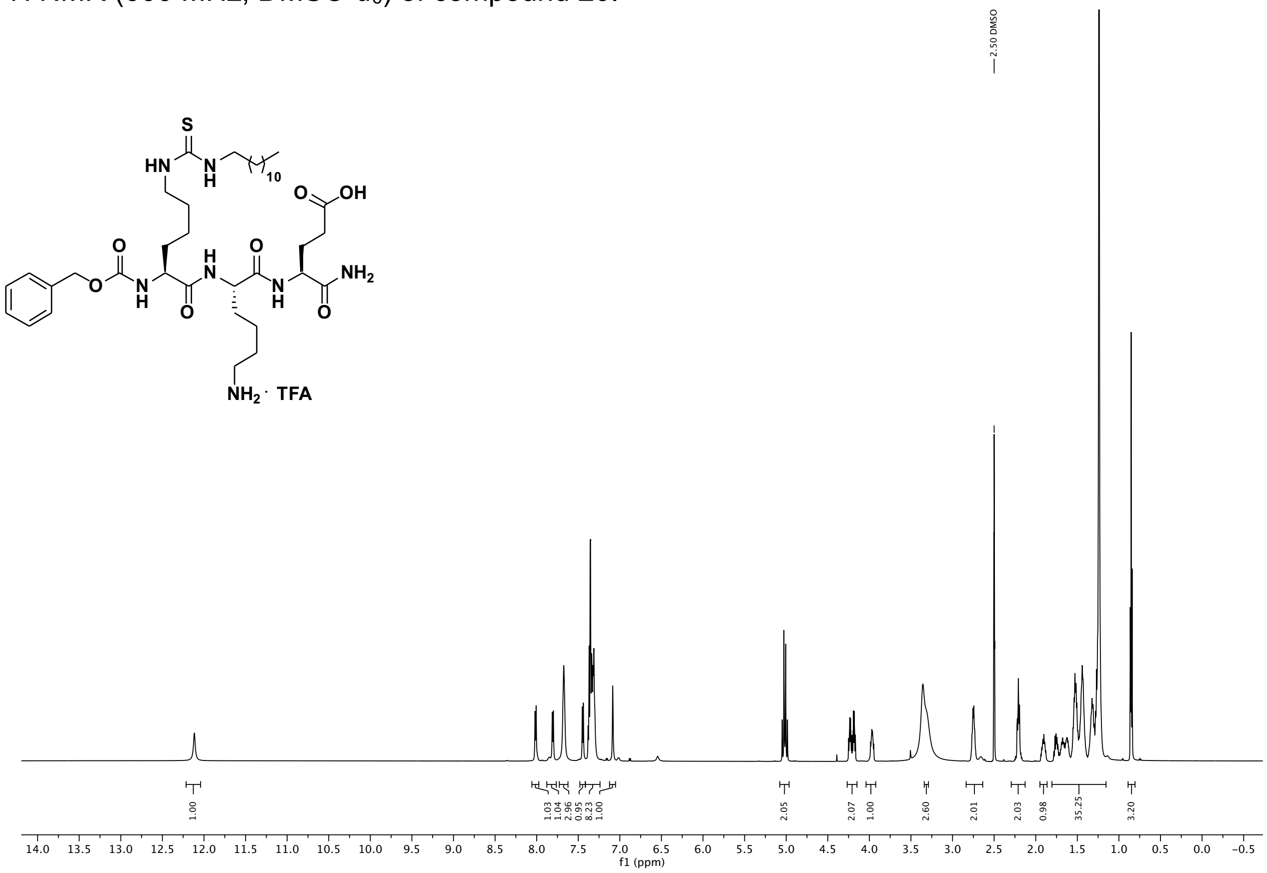

$^{13}C$ NMR (151 MHz, DMSO) of compound **20**.

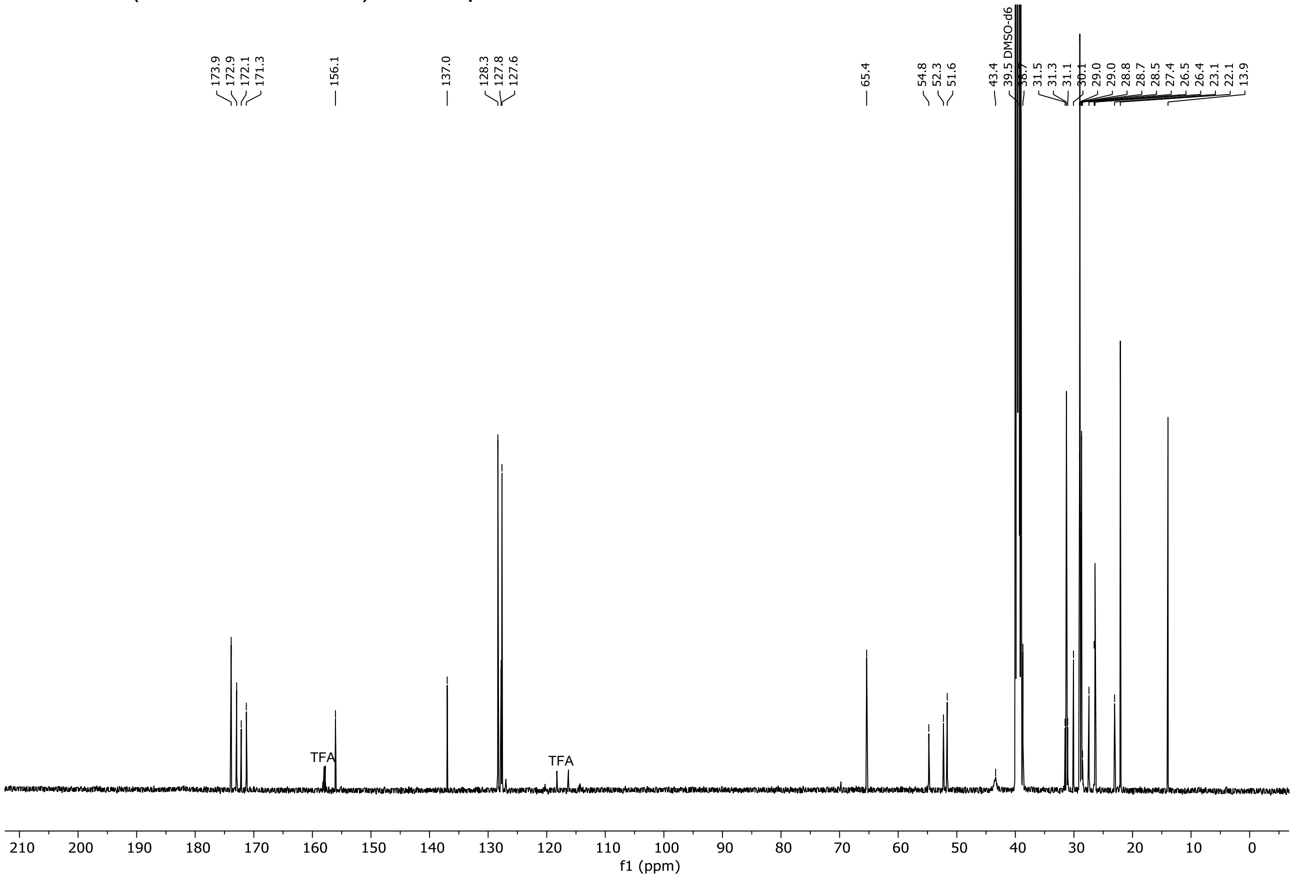

[1]H NMR (600 MHz, DMSO-$d_6$) of compound **21**.

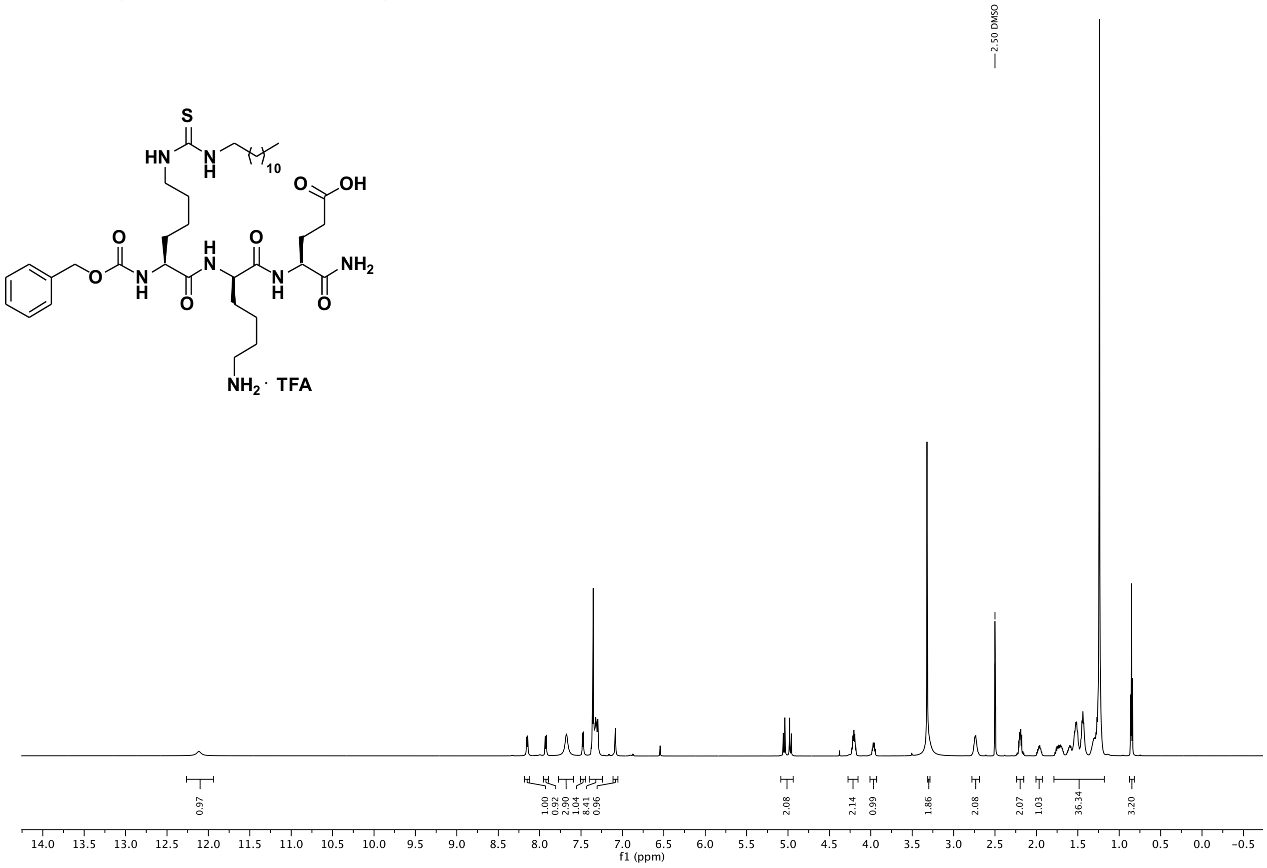

[13]C NMR (151 MHz, DMSO) of compound **21**.

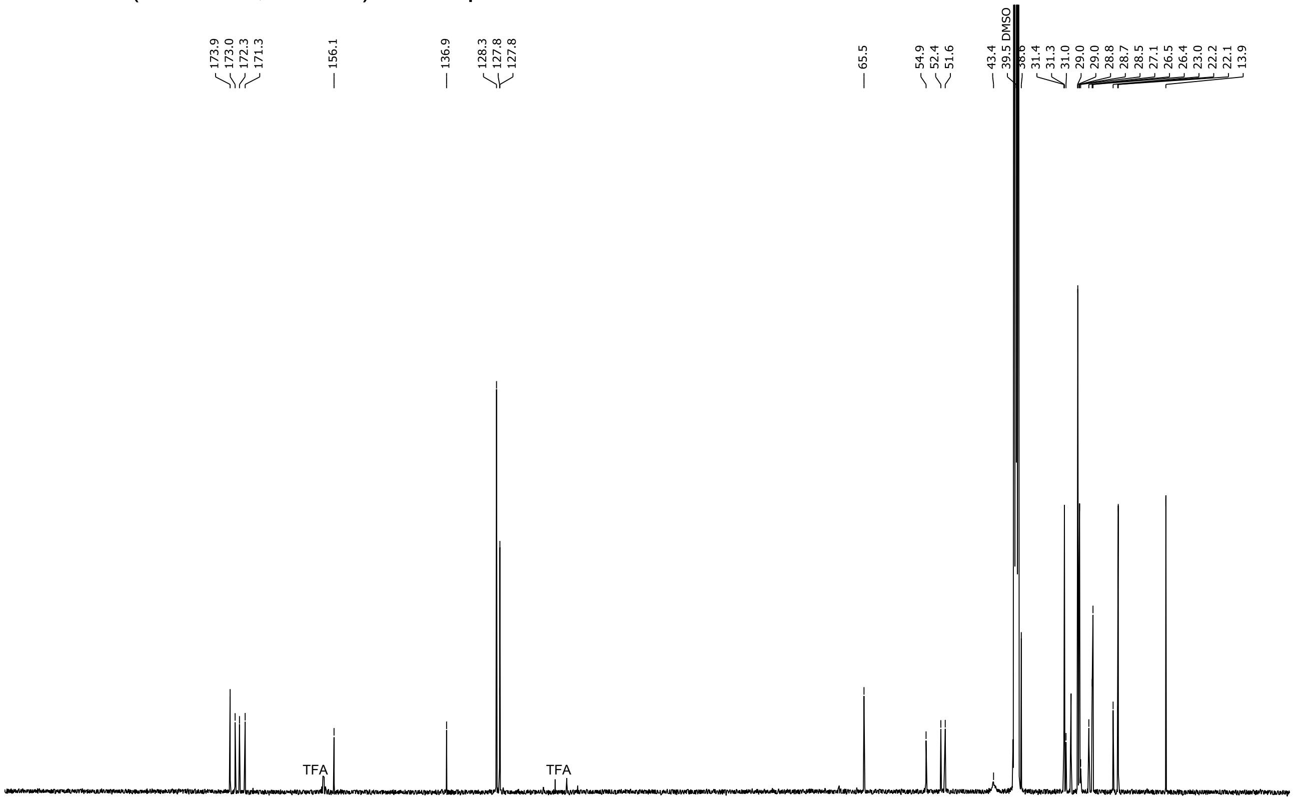

$^{1}H$ NMR (600 MHz, DMSO-$d_6$) of compound **22**.

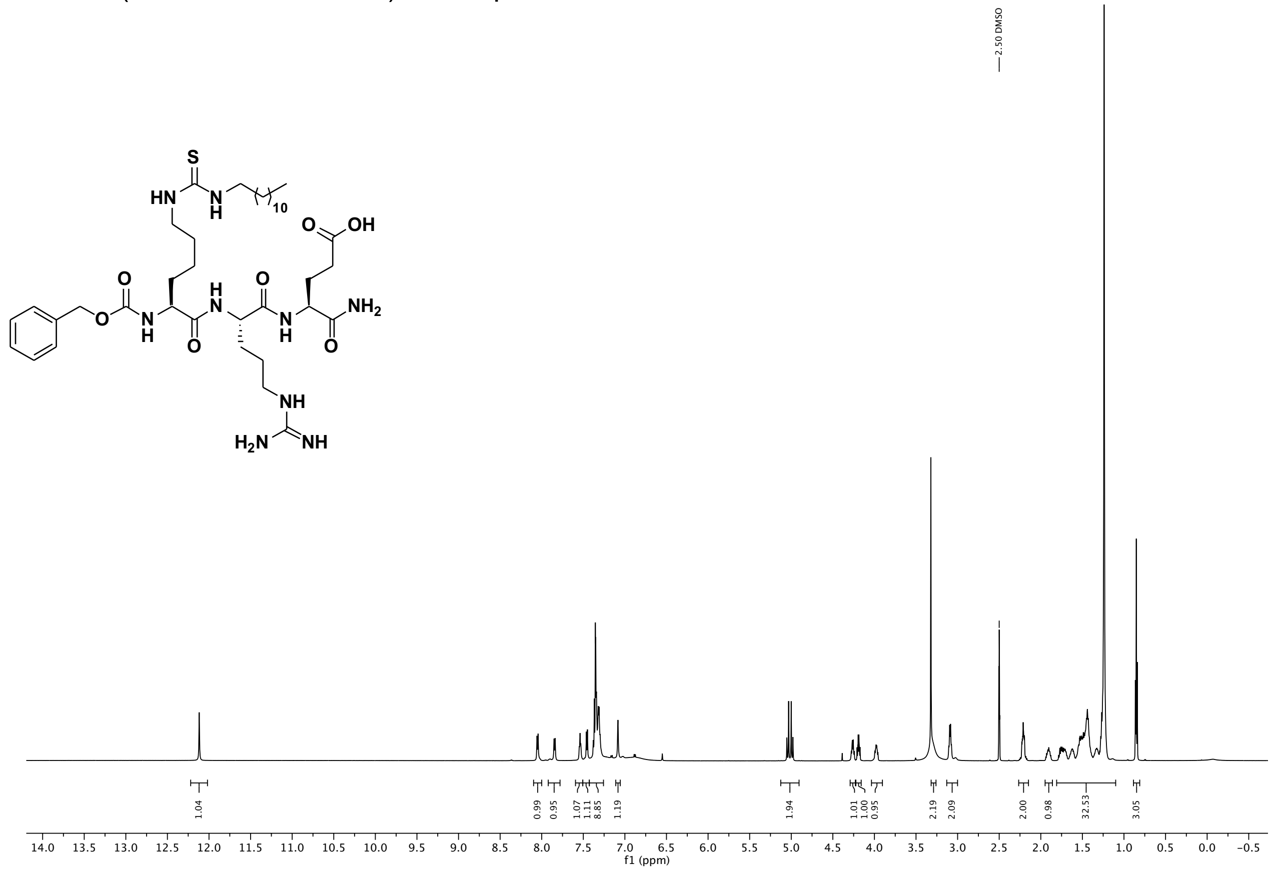

$^{13}C$ NMR (151 MHz, DMSO) of compound **22**.

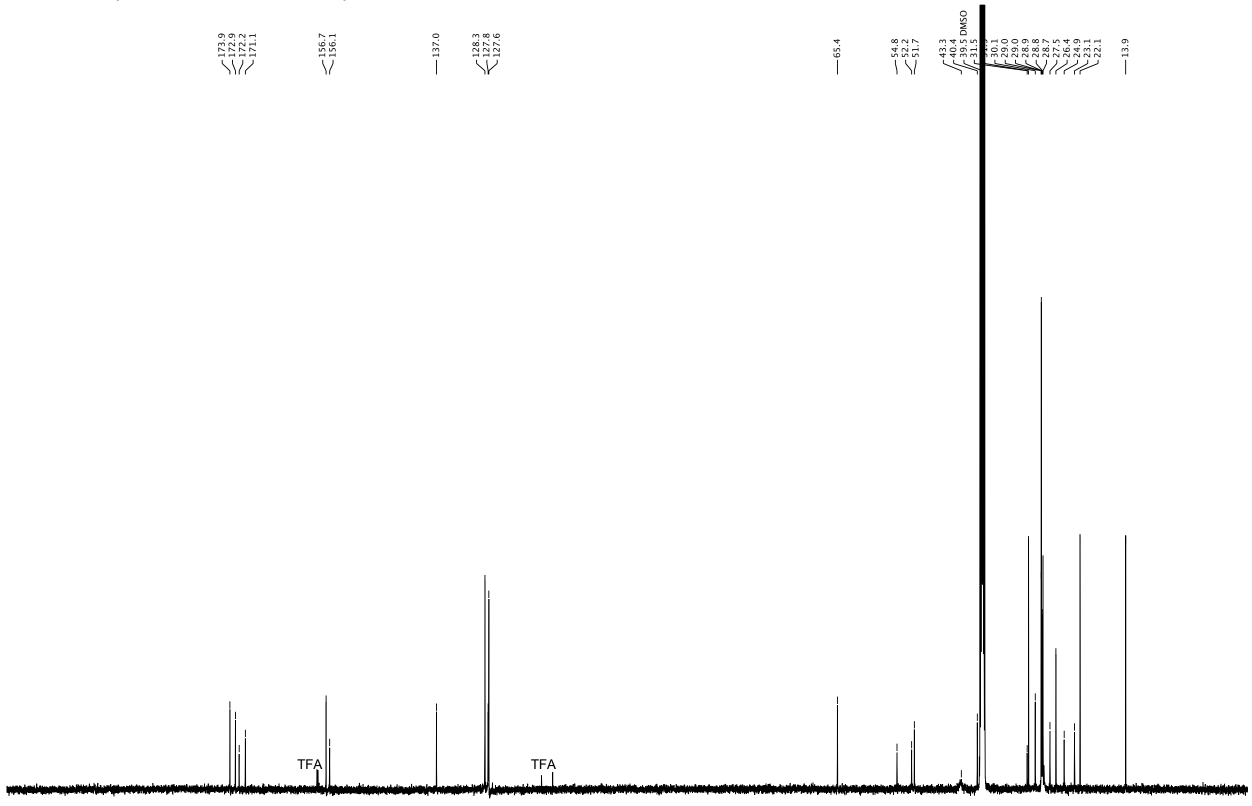

$^{1}H$ NMR (600 MHz, DMSO-$d_6$) of compound **23**.

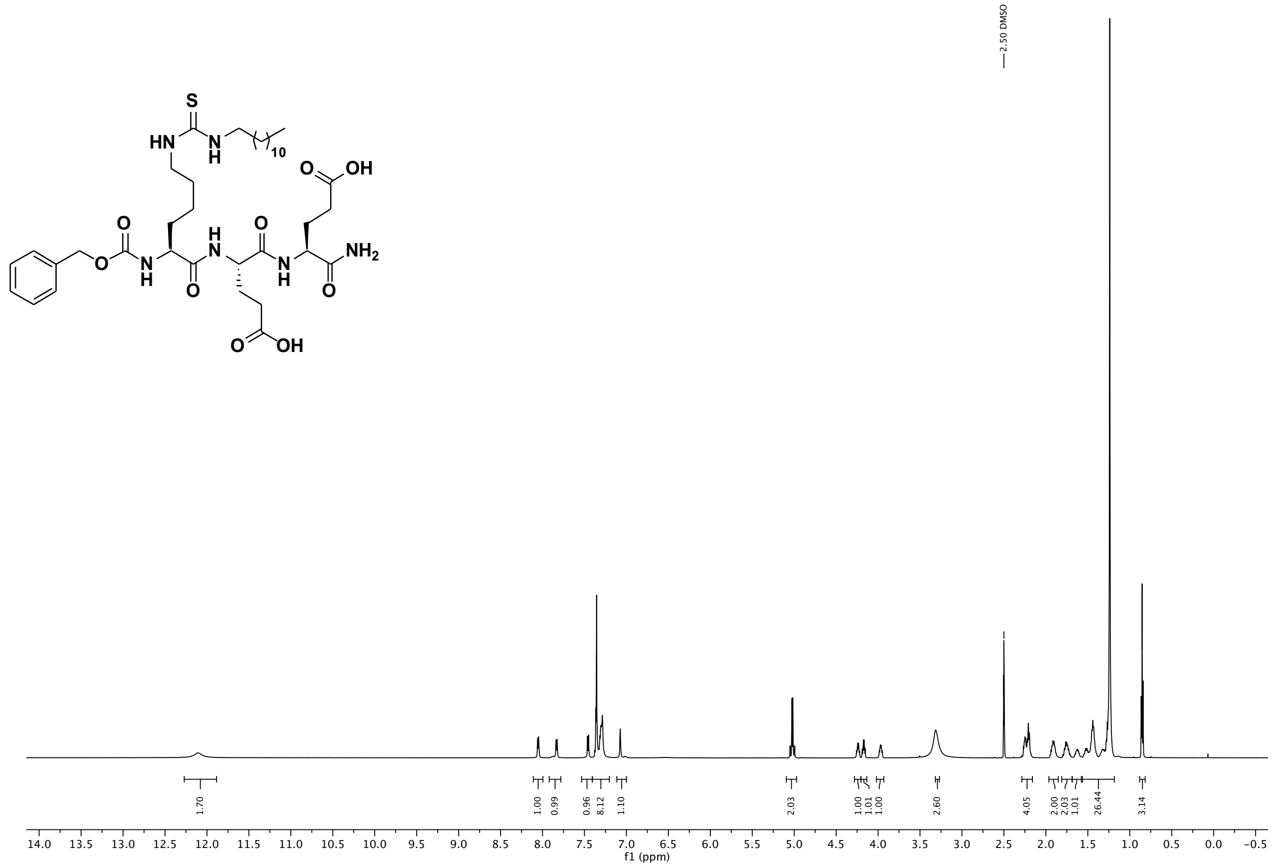

$^{13}C$ NMR (151 MHz, DMSO) of compound **23**.

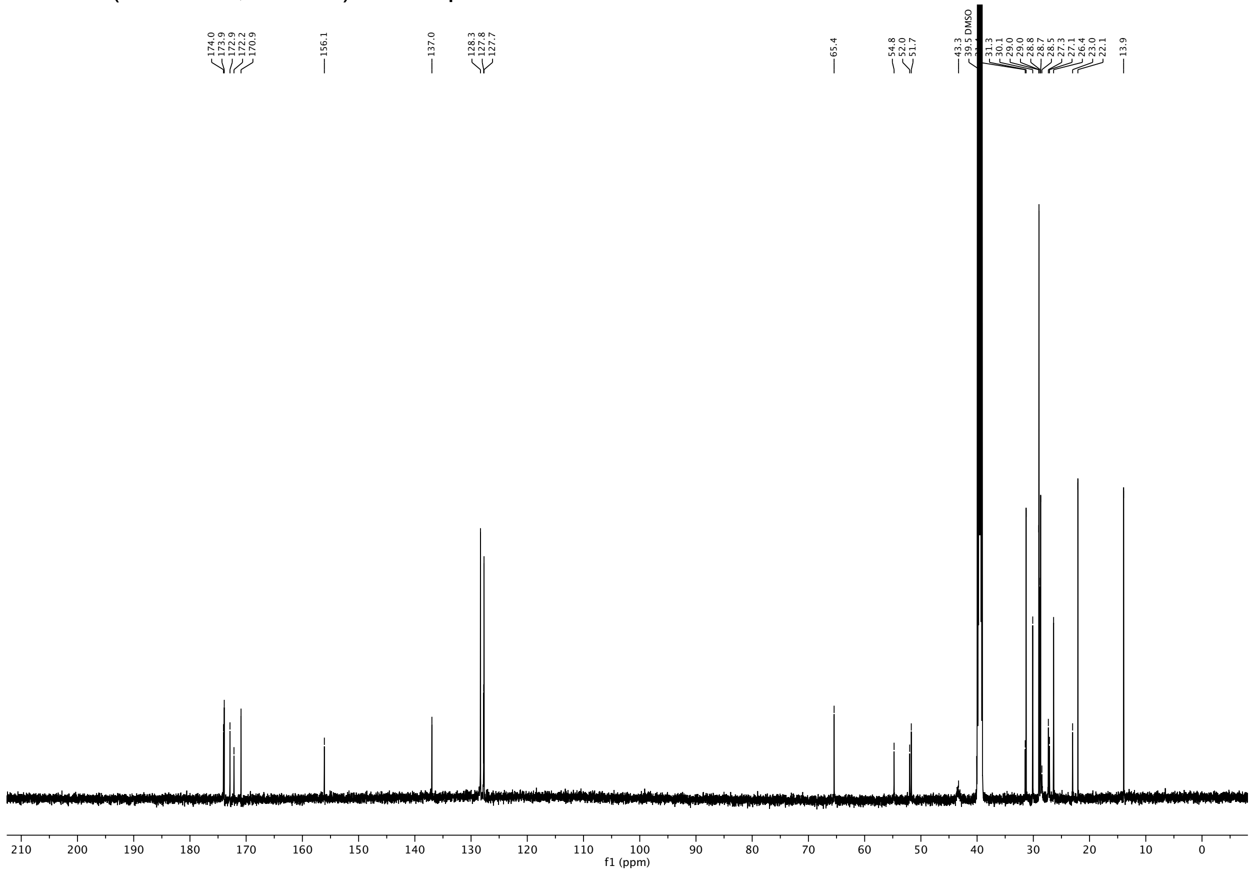

[1]H NMR (600 MHz, DMSO-$d_6$) of compound **24**.

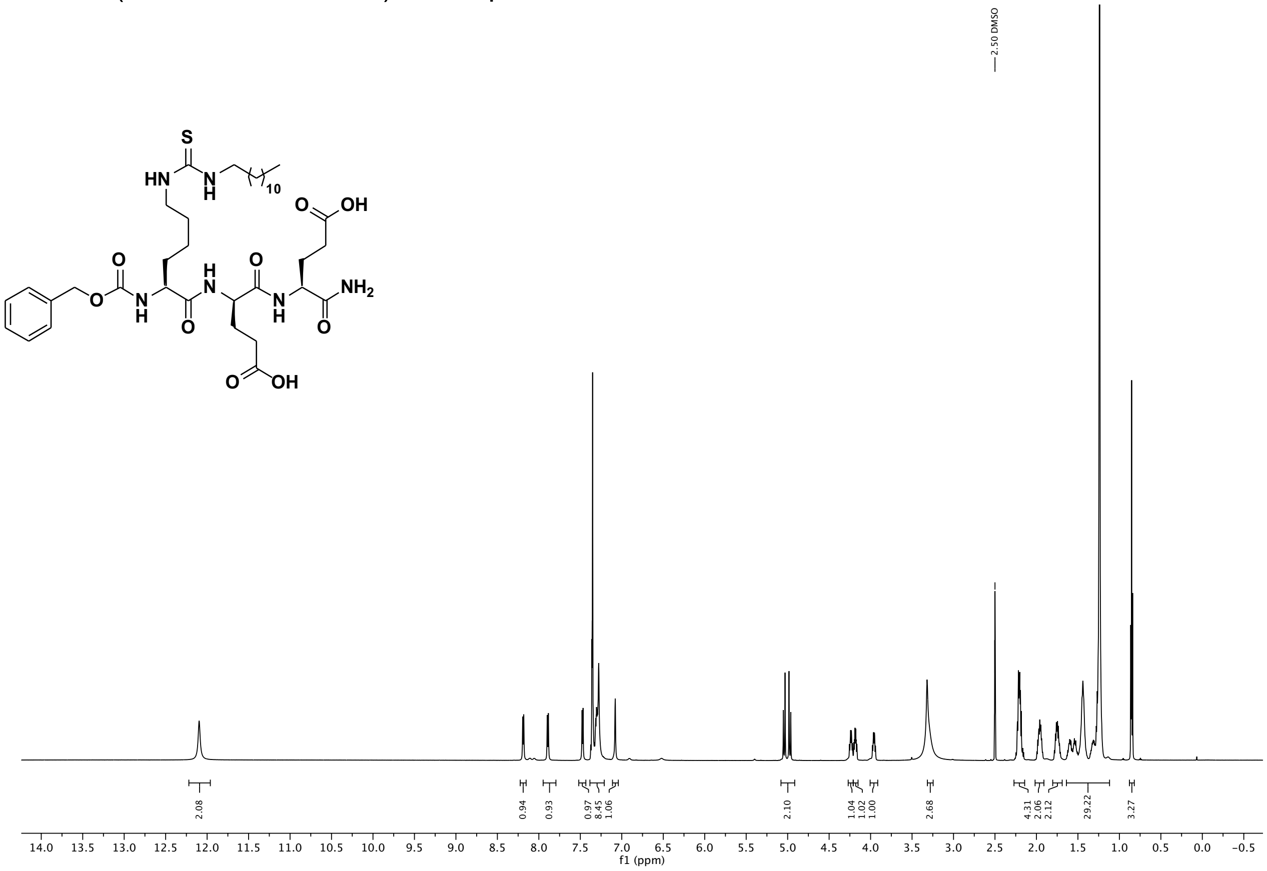

[13]C NMR (151 MHz, DMSO) of compound **24**.

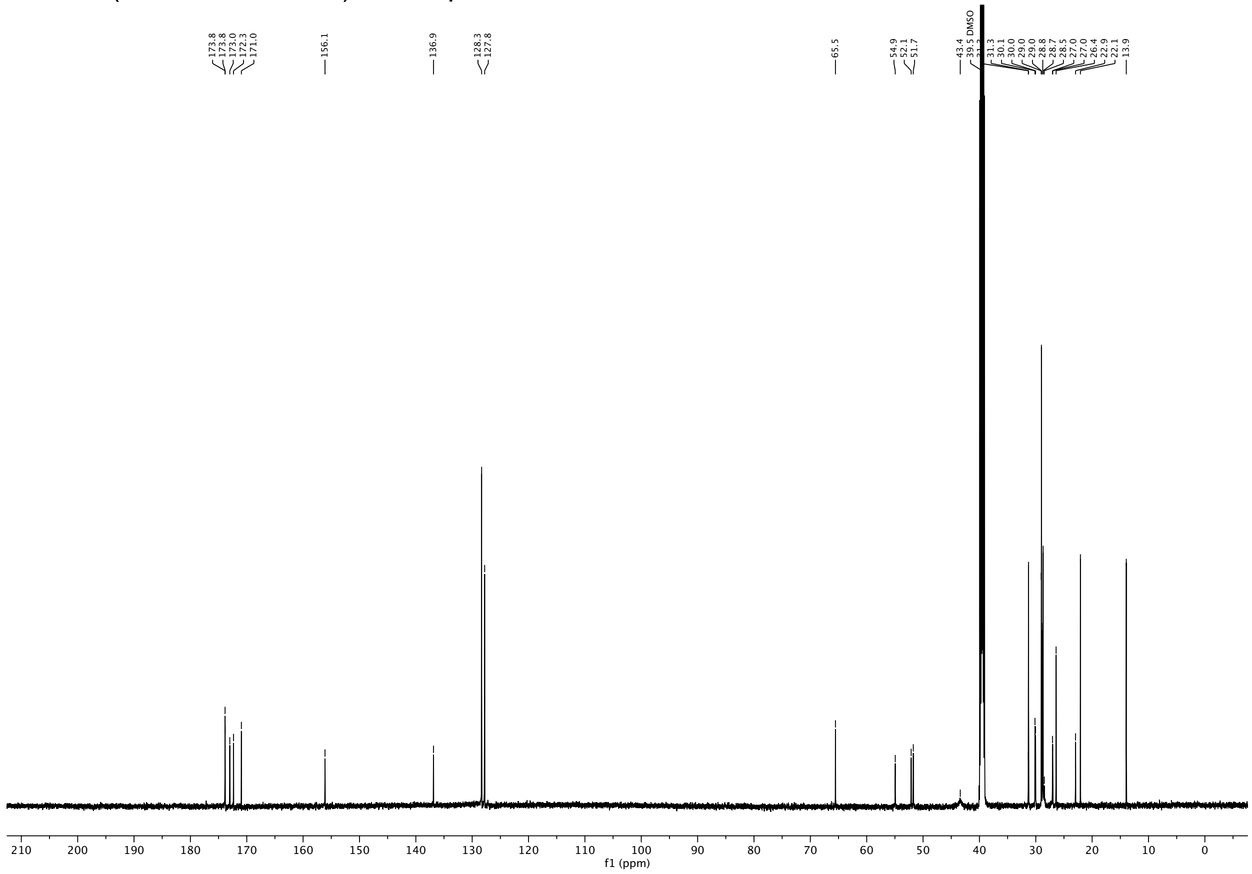

$^{1}$H NMR (600 MHz, DMSO-$d_6$) of compound **25**.

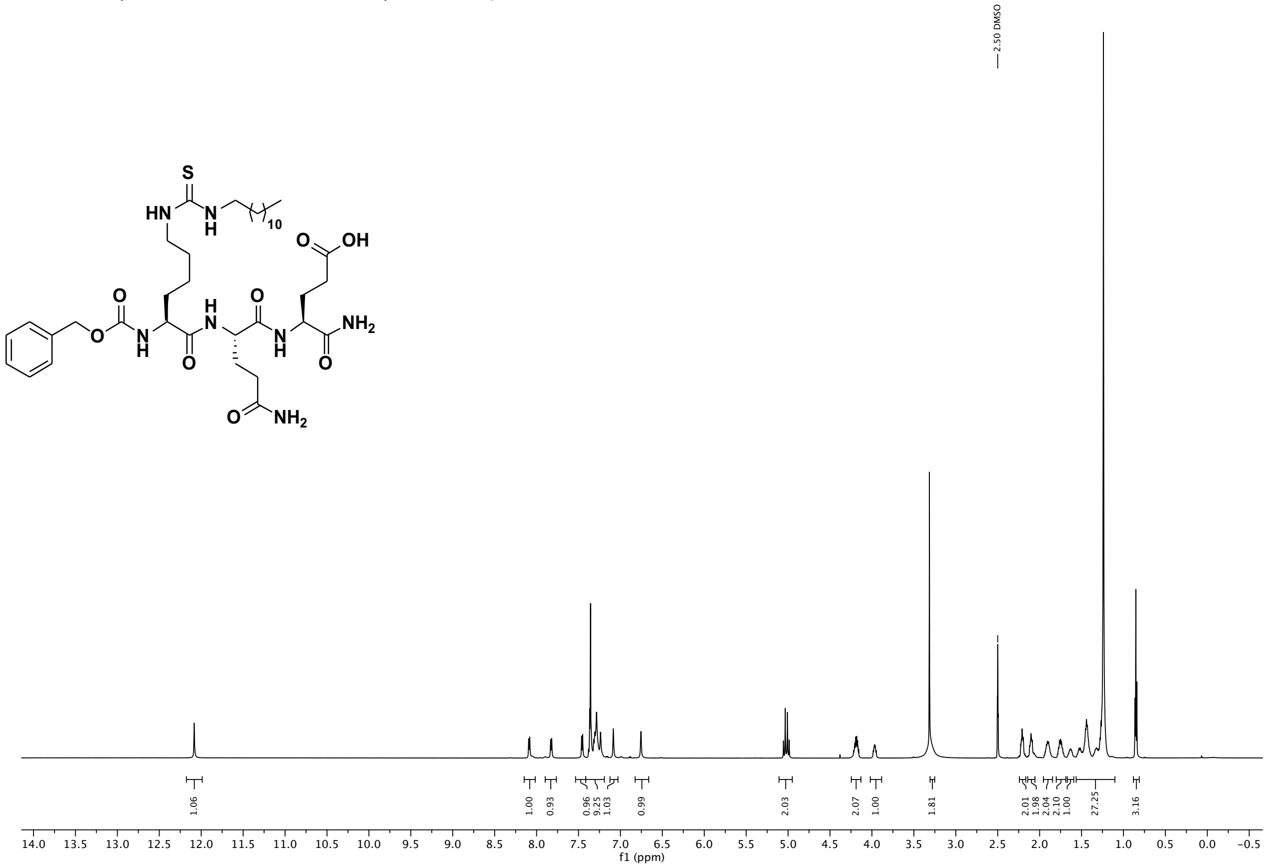

$^{13}$C NMR (151 MHz, DMSO) of compound **25**.

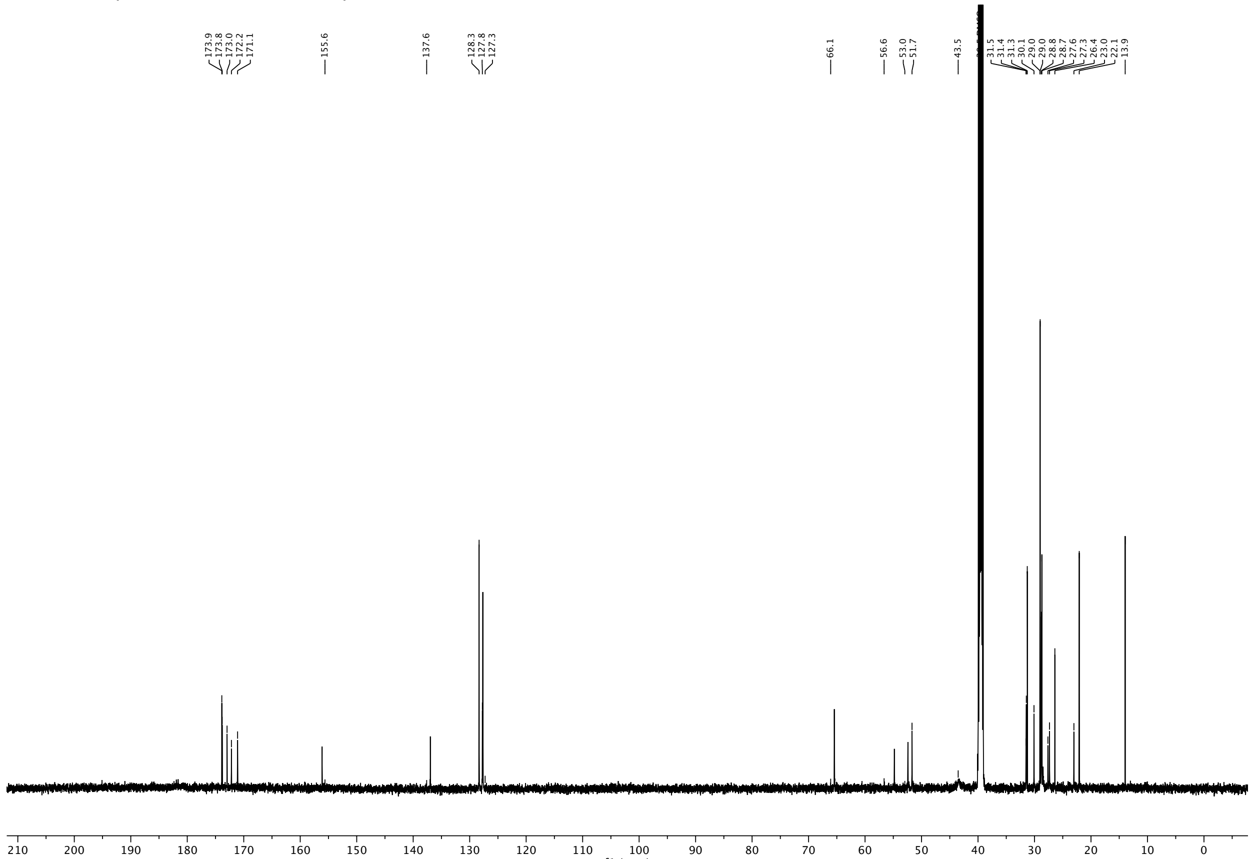

[1]H NMR (600 MHz, DMSO-$d_6$) of compound **26**.

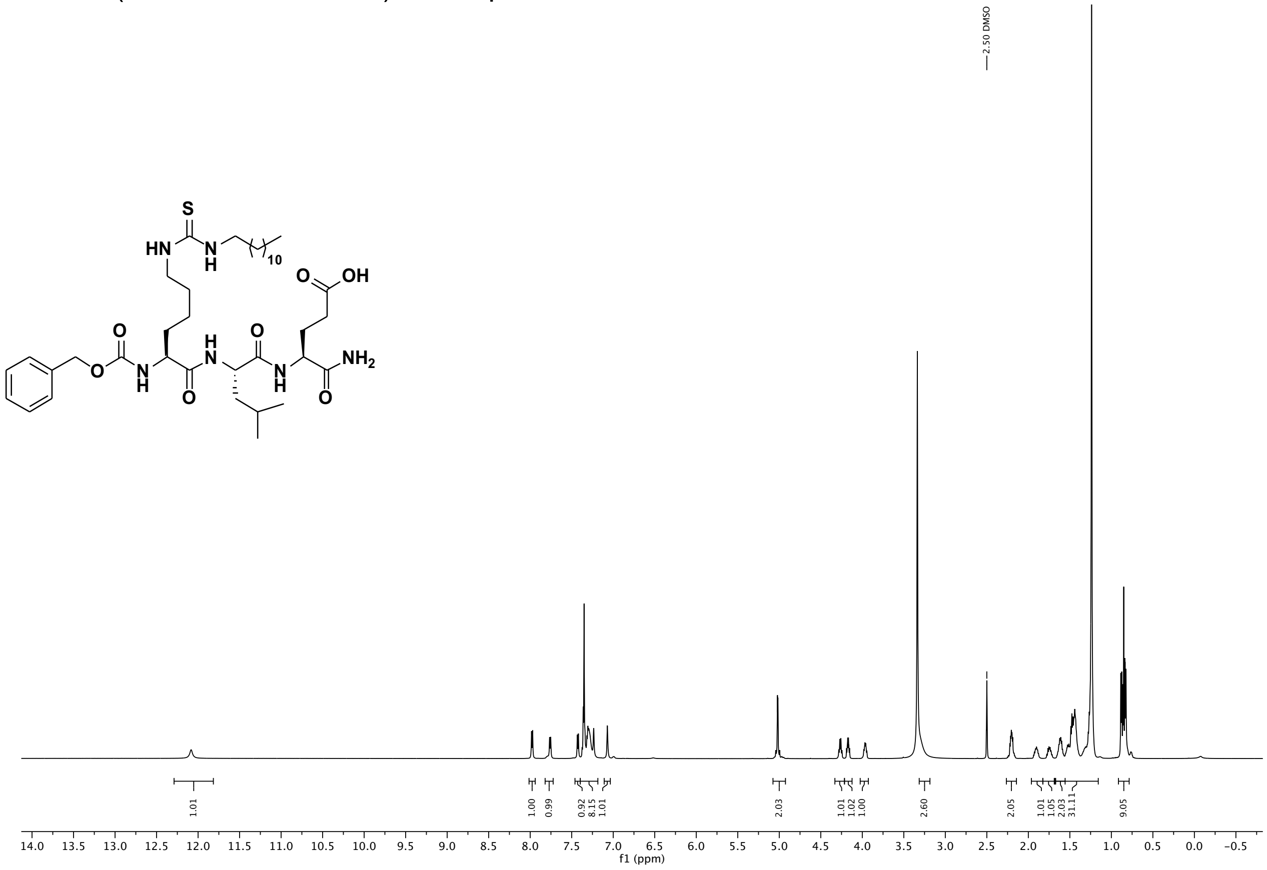

[13]C NMR (151 MHz, DMSO) of compound **26**.

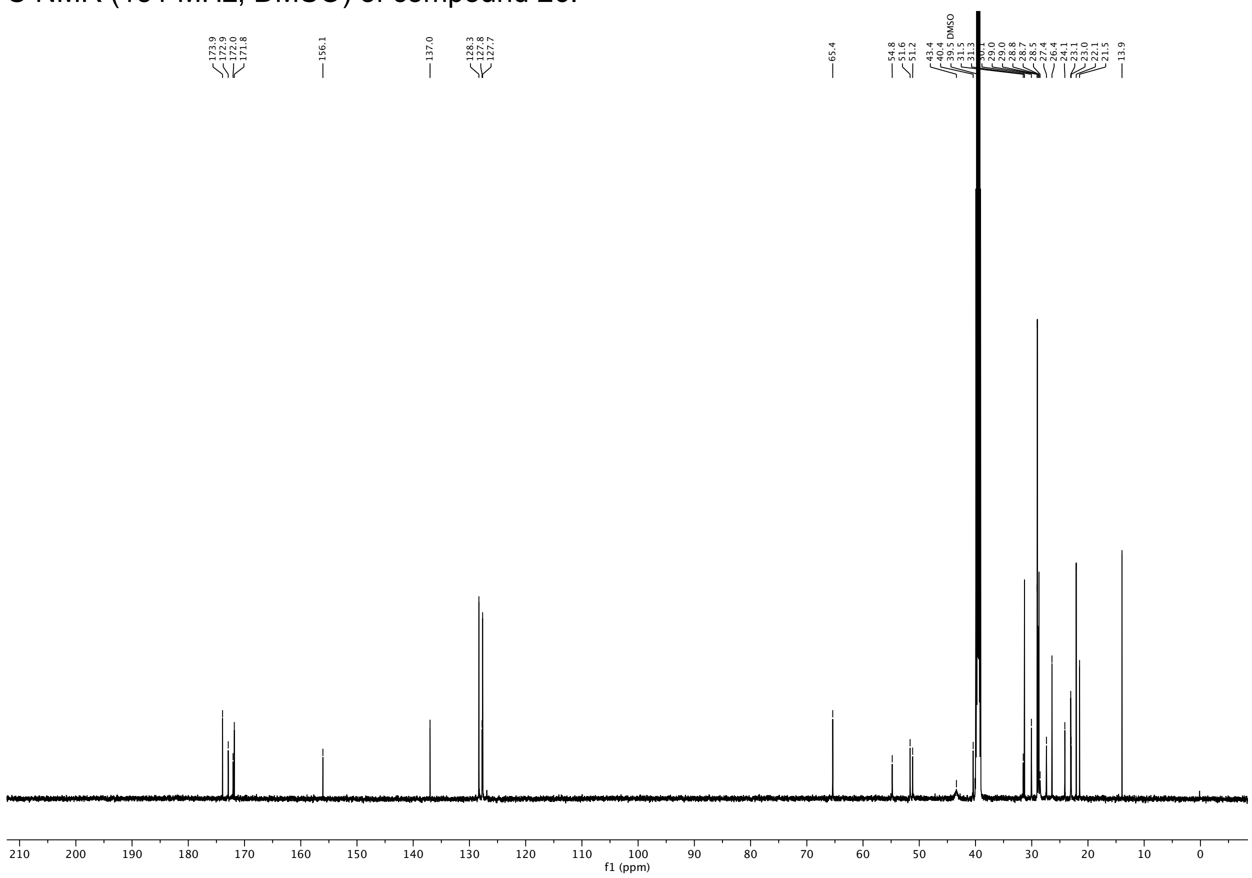

[1]H NMR (600 MHz, DMSO-$d_6$) of compound **27**.

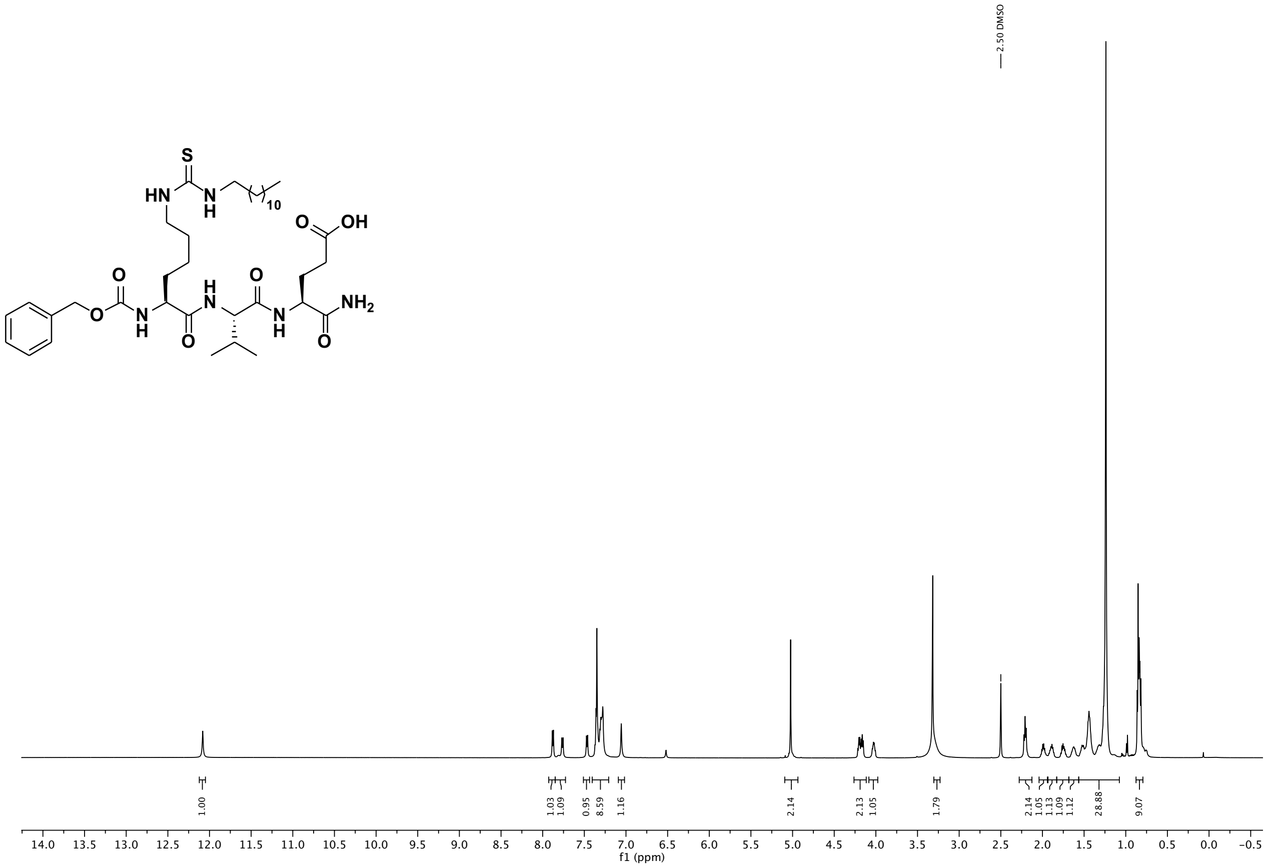

[13]C NMR (151 MHz, DMSO) of compound **27**.

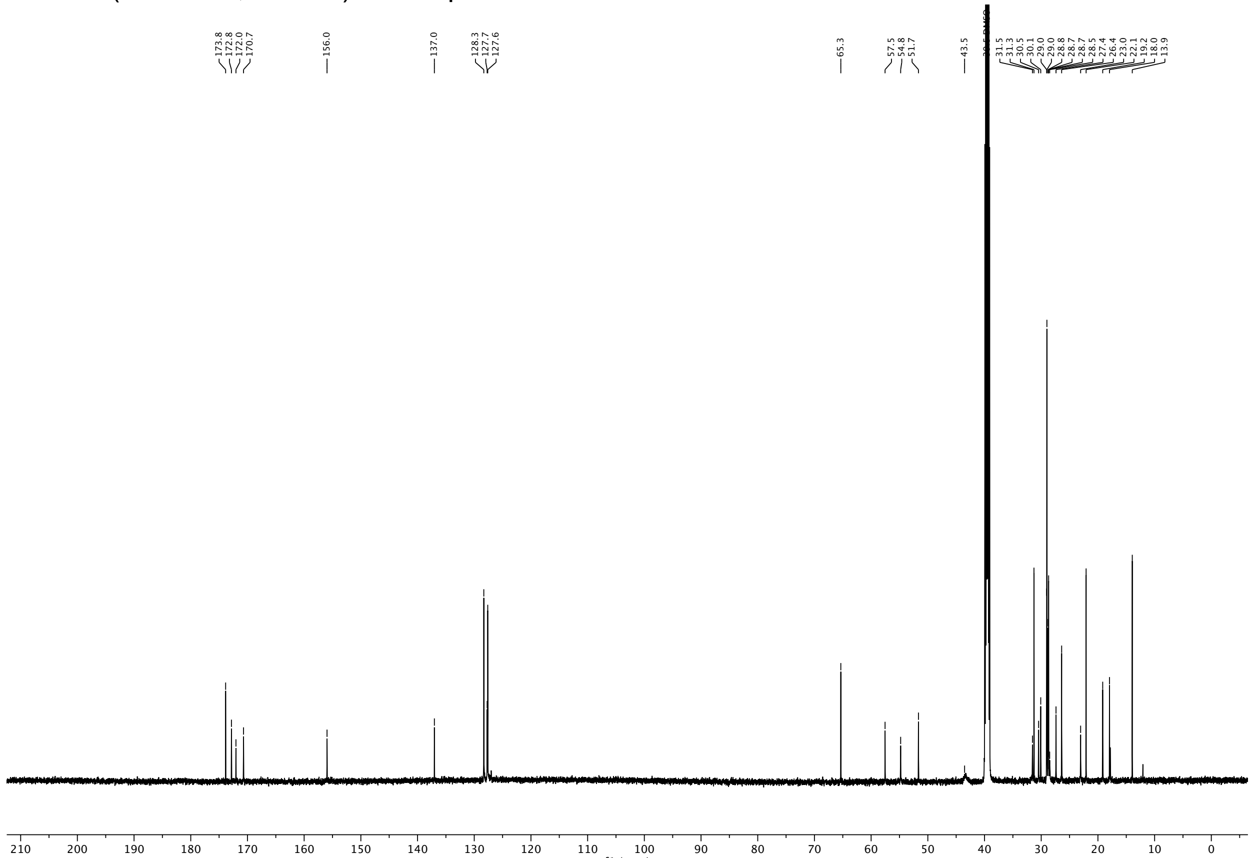

$^{1}H$ NMR (600 MHz, DMSO-$d_6$) of compound **28**.

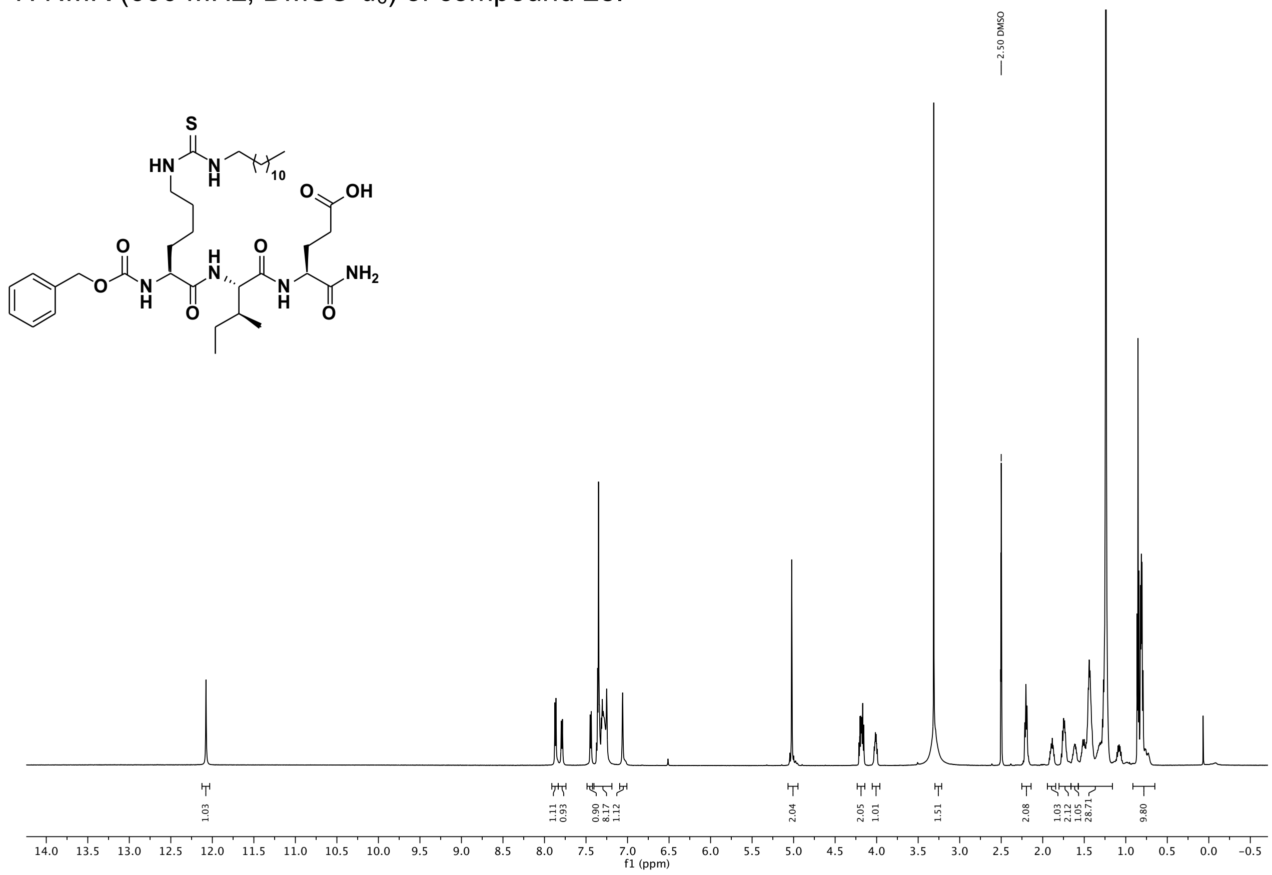

$^{13}C$ NMR (151 MHz, DMSO) of compound **28**.

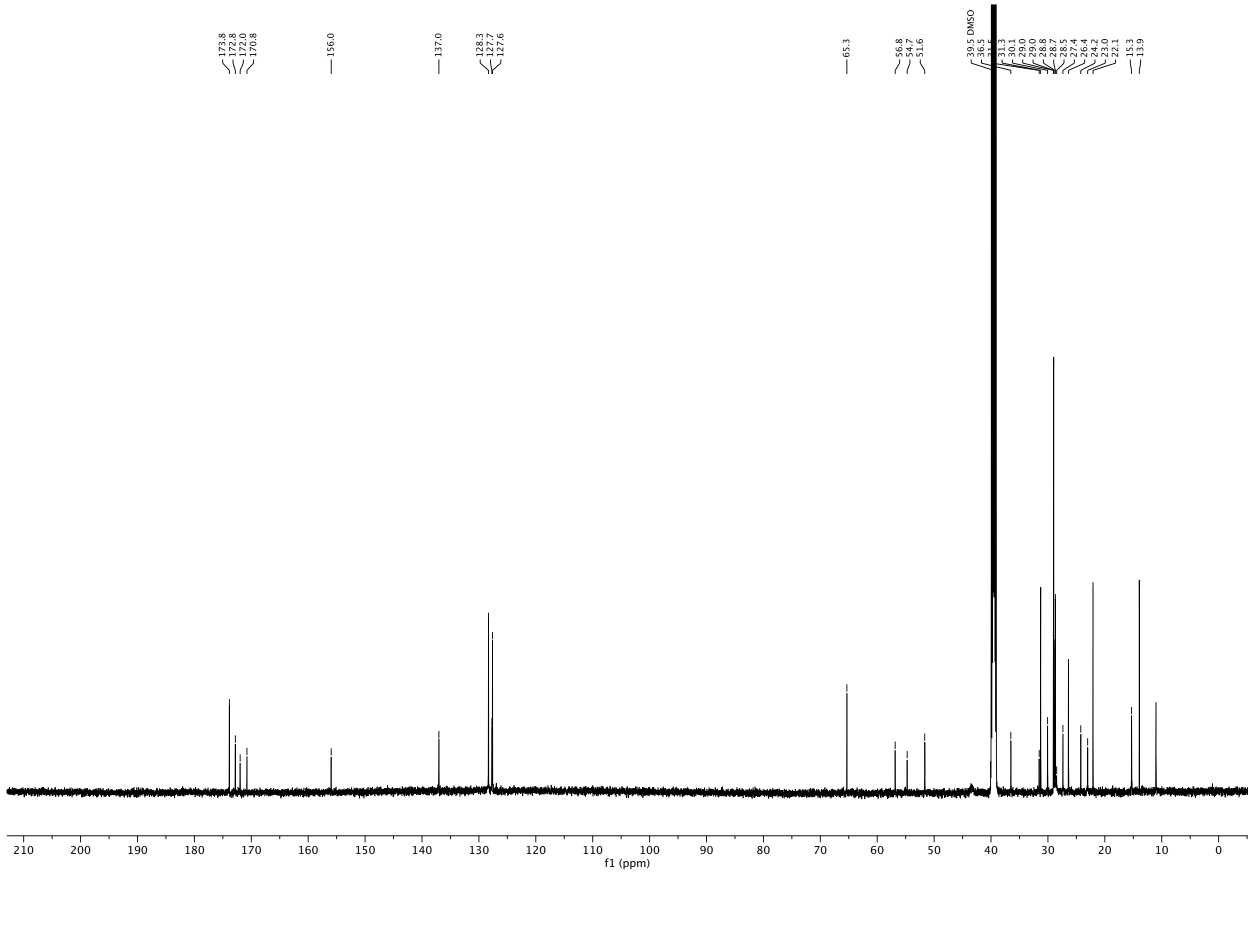

$^{1}H$ NMR (600 MHz, DMSO-$d_6$) of compound **29**.

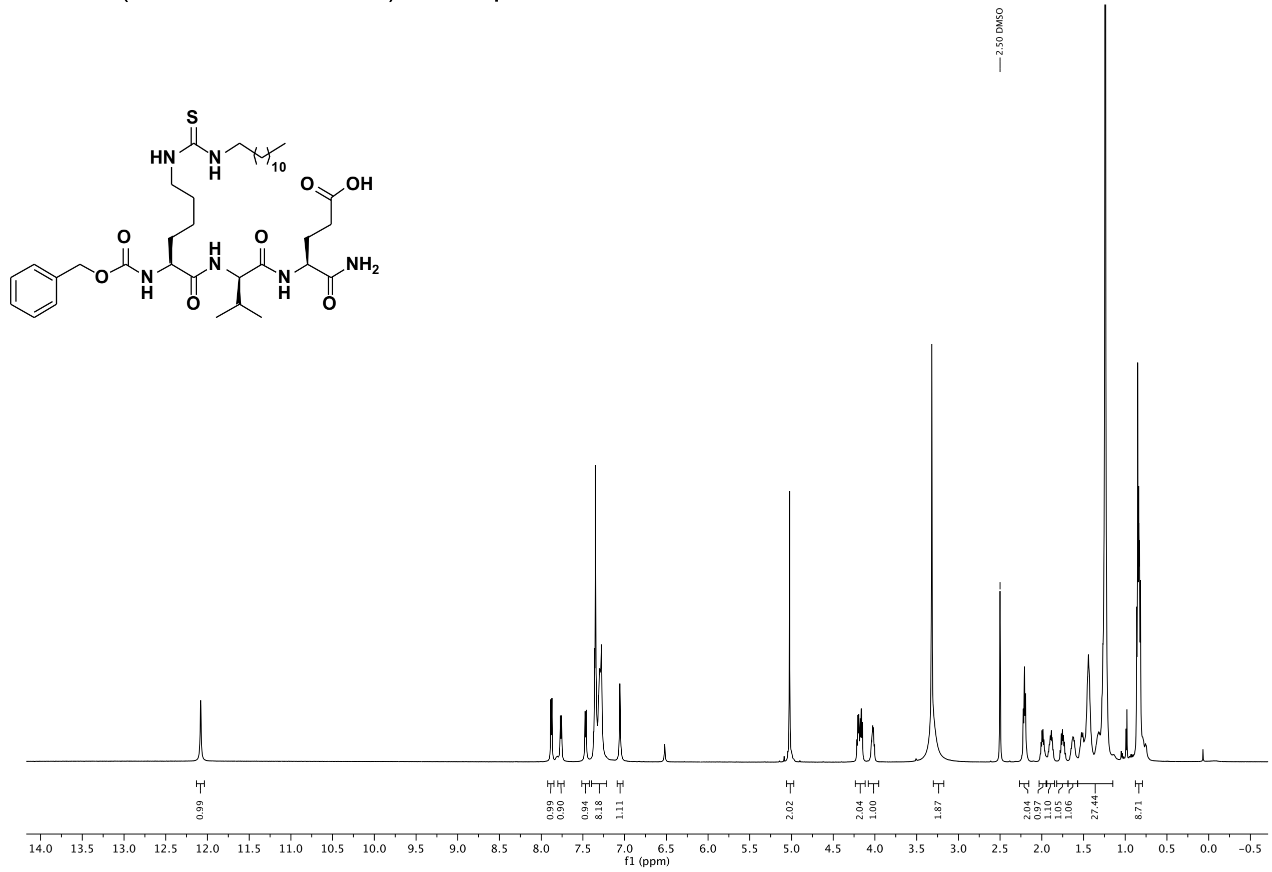

$^{13}C$ NMR (151 MHz, DMSO) of compound **29**.

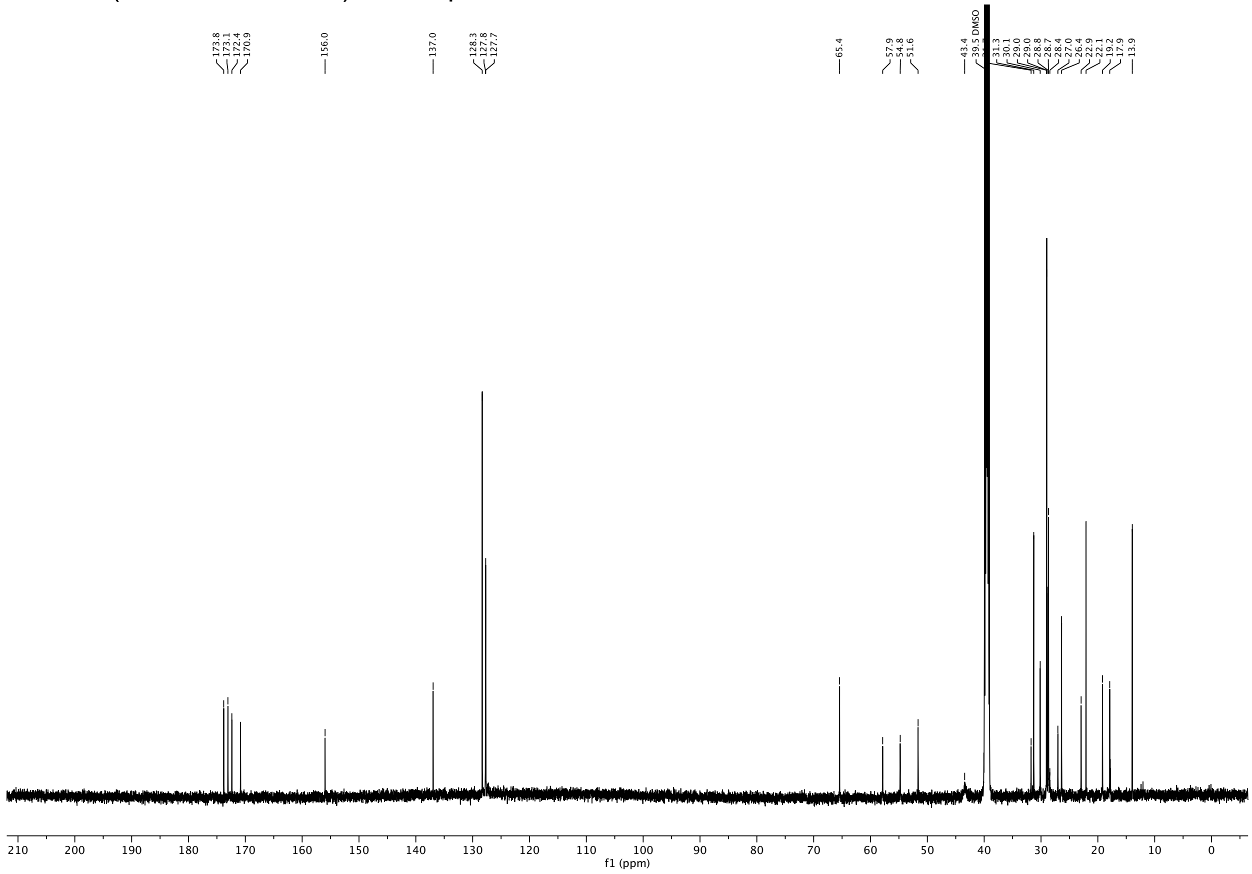

[1]H NMR (600 MHz, DMSO-$d_6$) of compound **30**

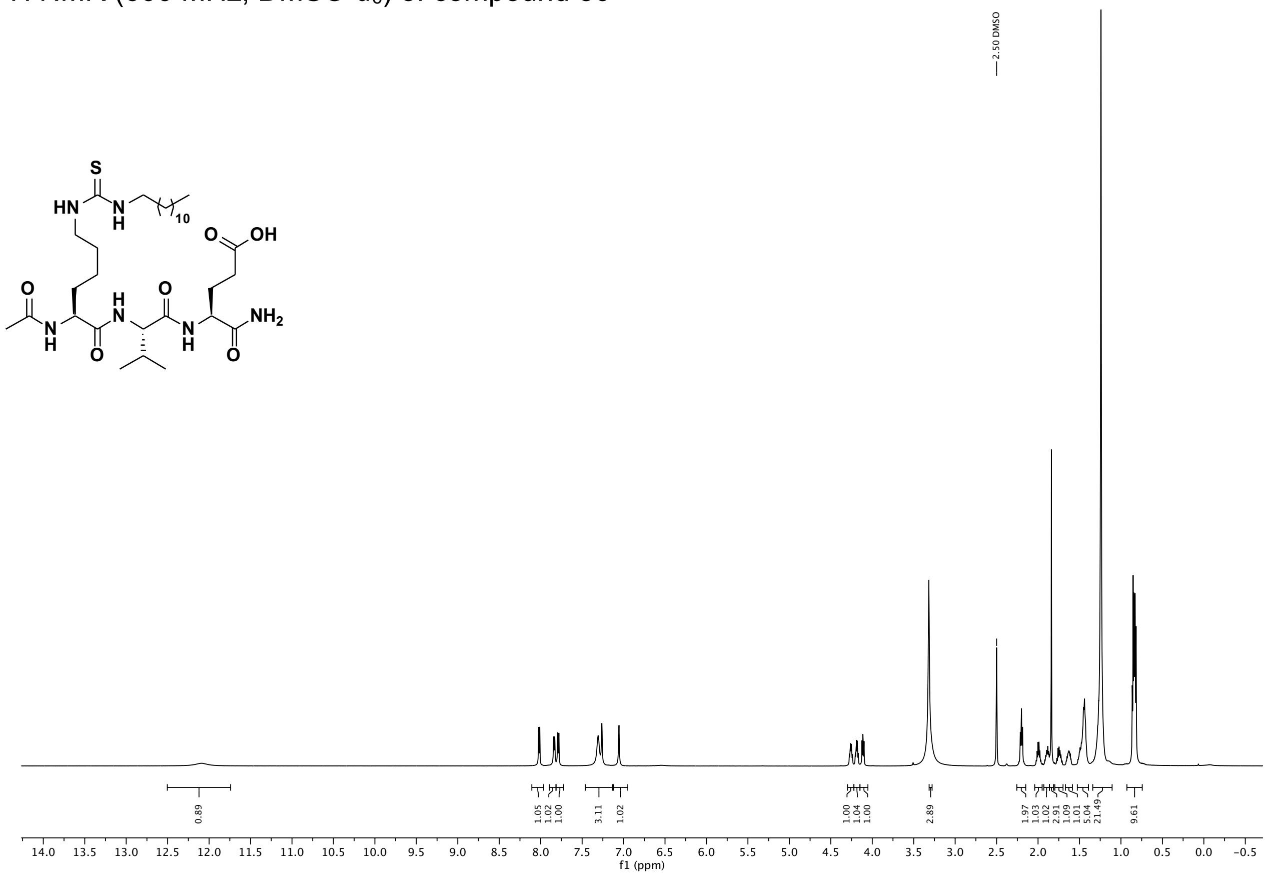

[13]C NMR (151 MHz, DMSO) of compound **30**.

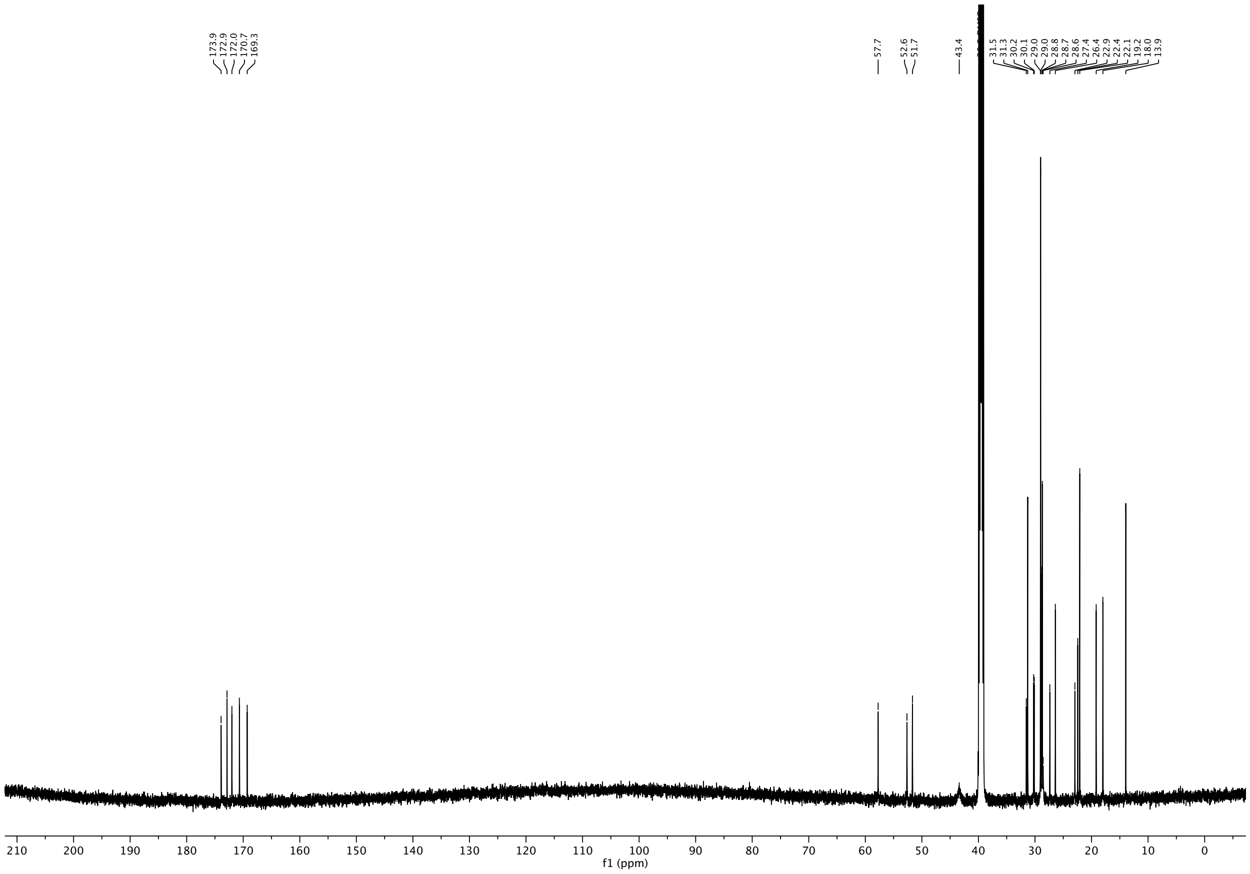

$^{1}H$ NMR (600 MHz, DMSO-$d_6$) of compound **31**.

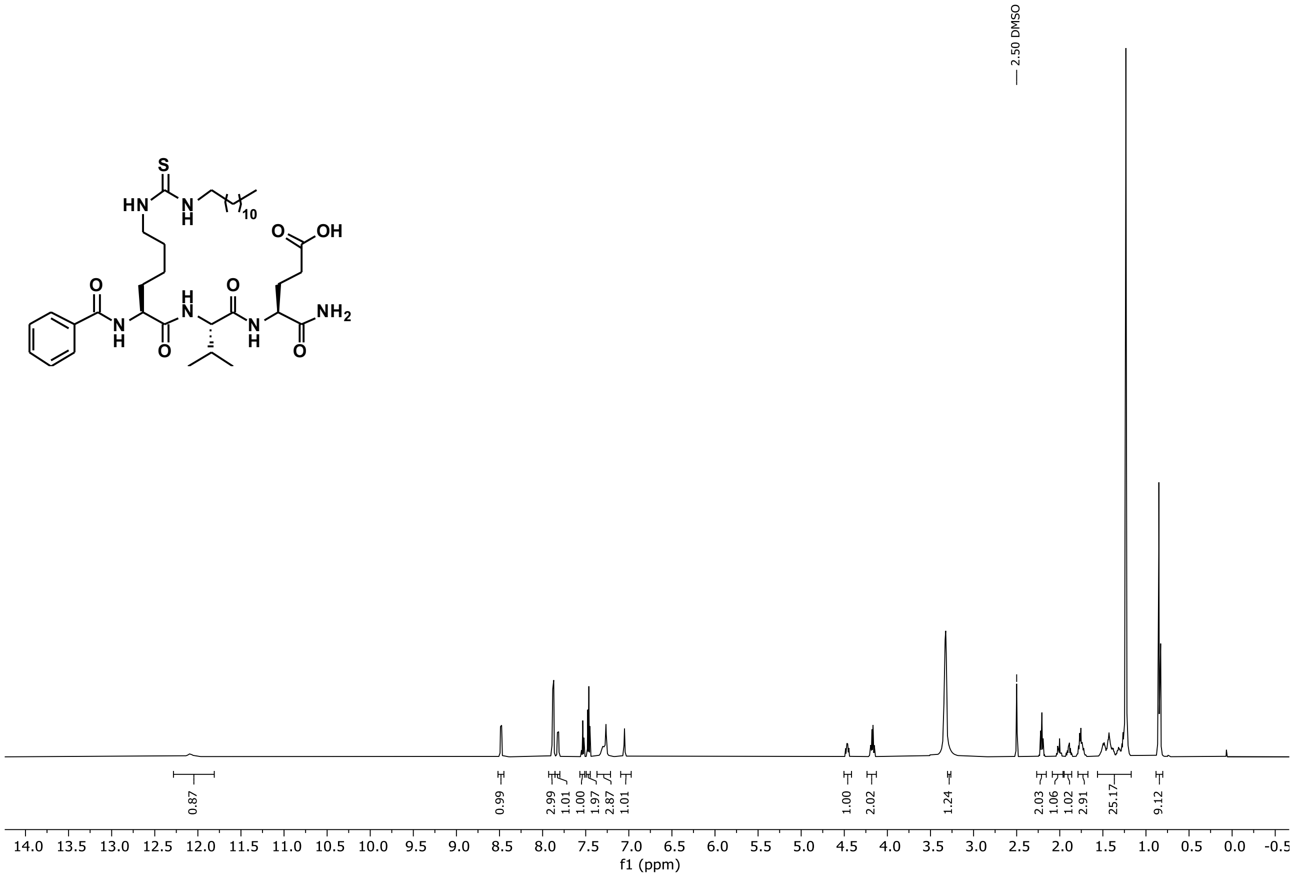

$^{13}C$ NMR (151 MHz, DMSO) of compound **31**.

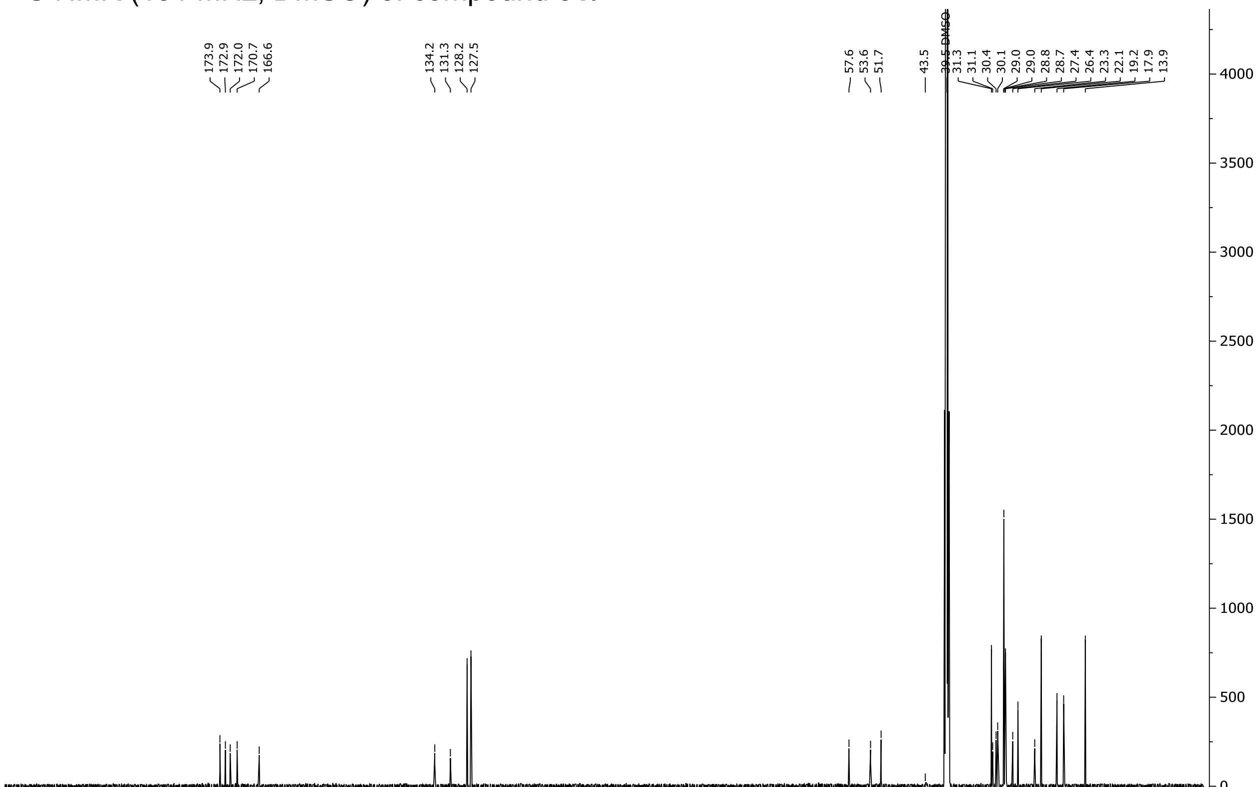

$^{1}H$ NMR (600 MHz, DMSO-$d_6$) of compound **32**.

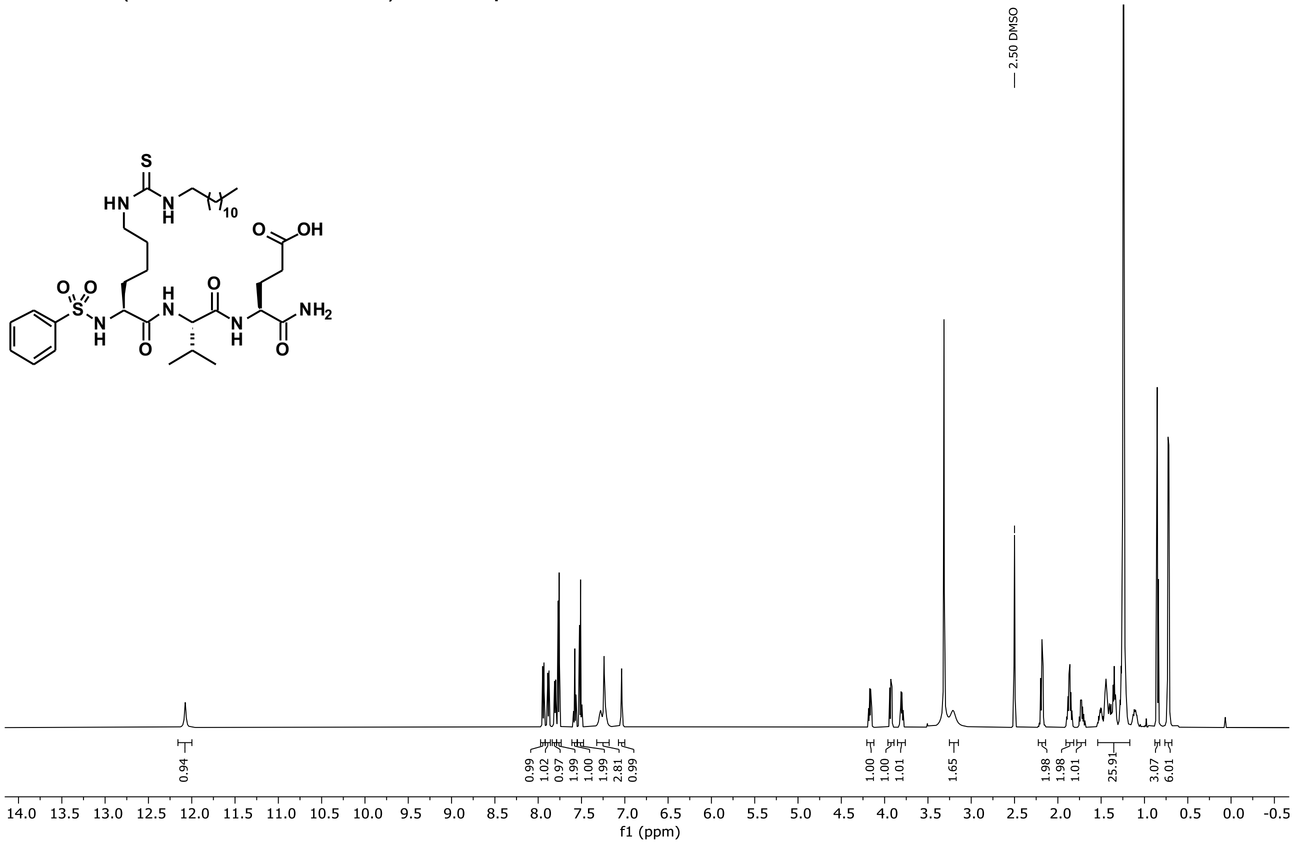

$^{13}C$ NMR (151 MHz, DMSO) of compound **32**.

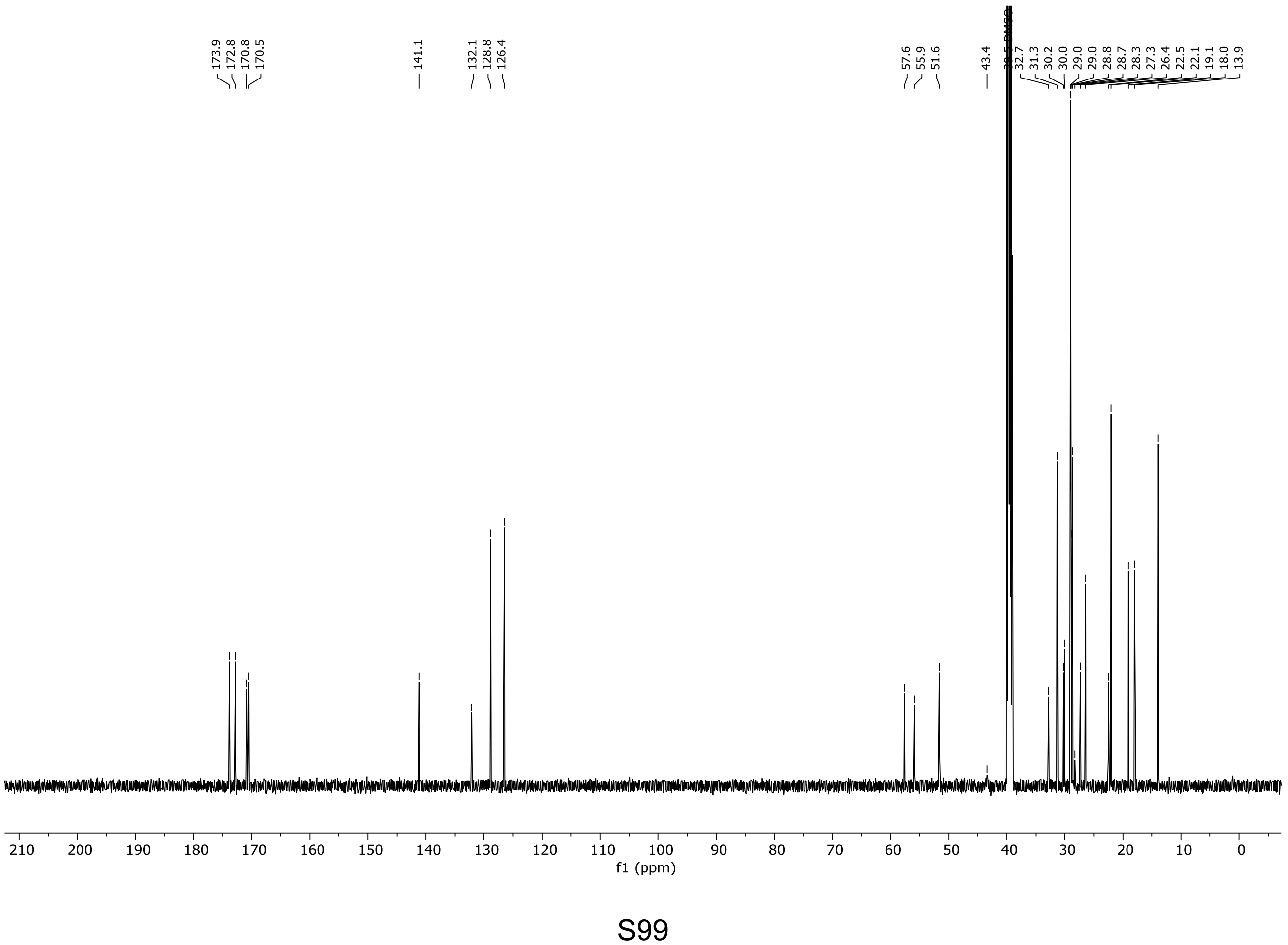

$^{1}$H NMR (600 MHz, DMSO-$d_6$) of compound **33**.

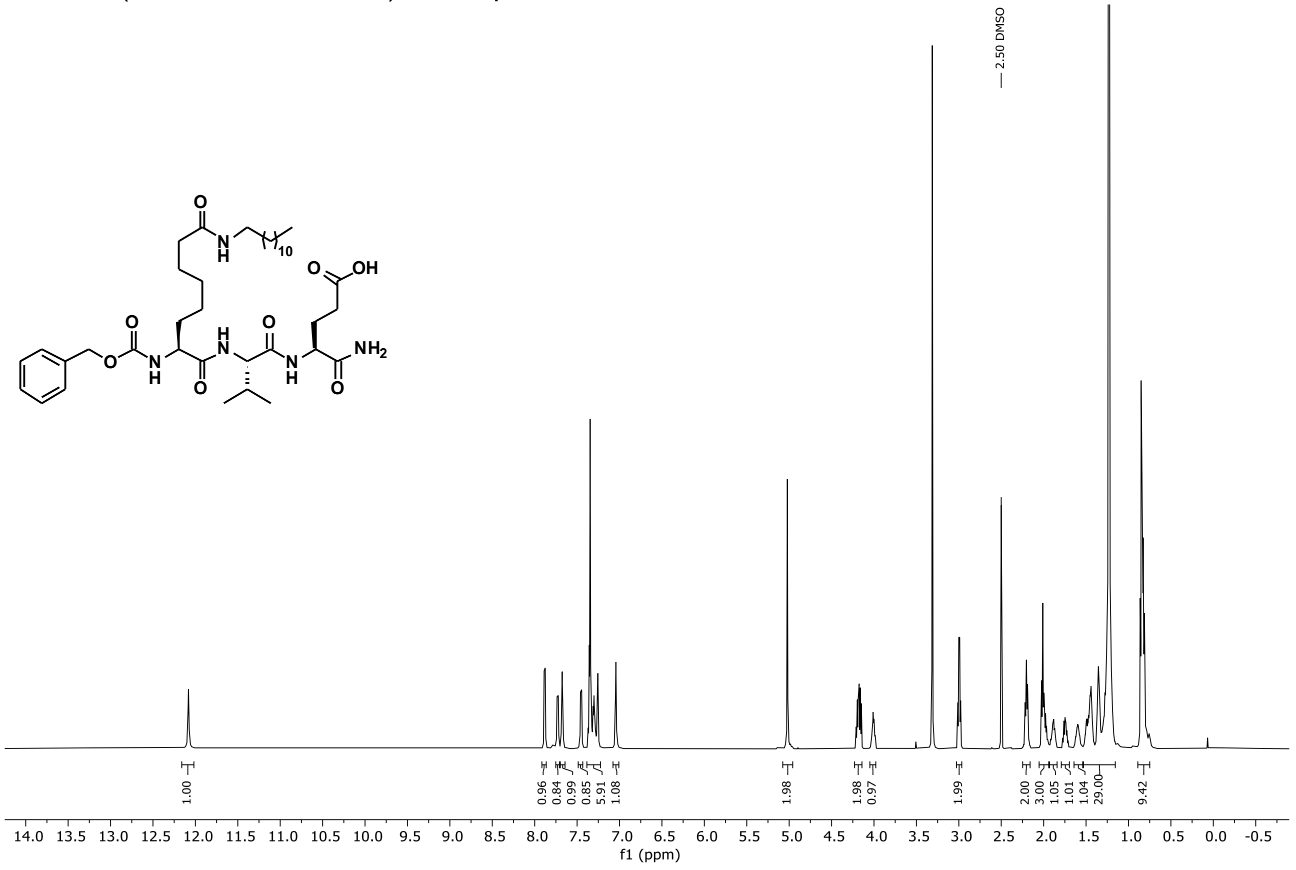

$^{13}$C NMR (151 MHz, DMSO) of compound **33**.

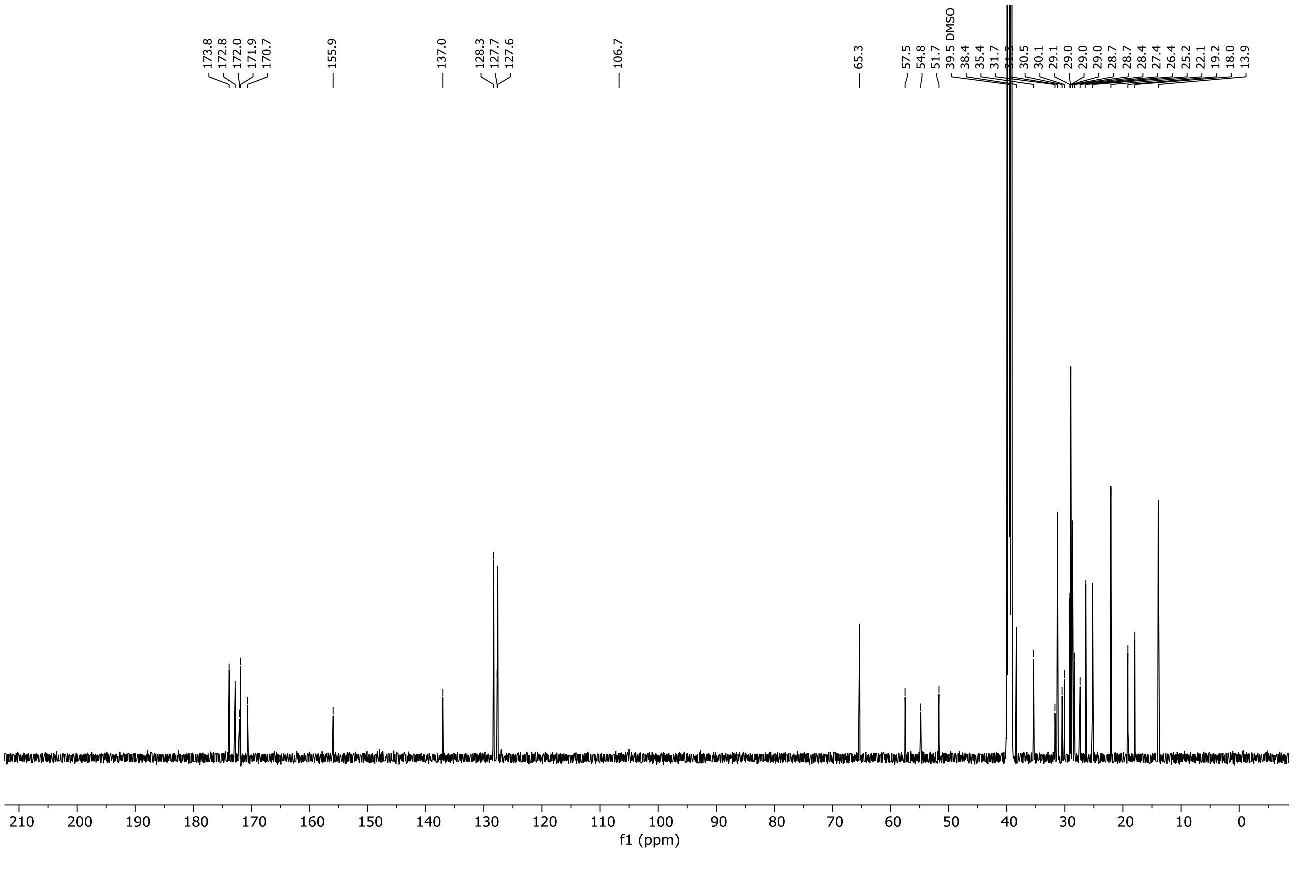

[1]H NMR (600 MHz, DMSO-$d_6$) of compound **34**.

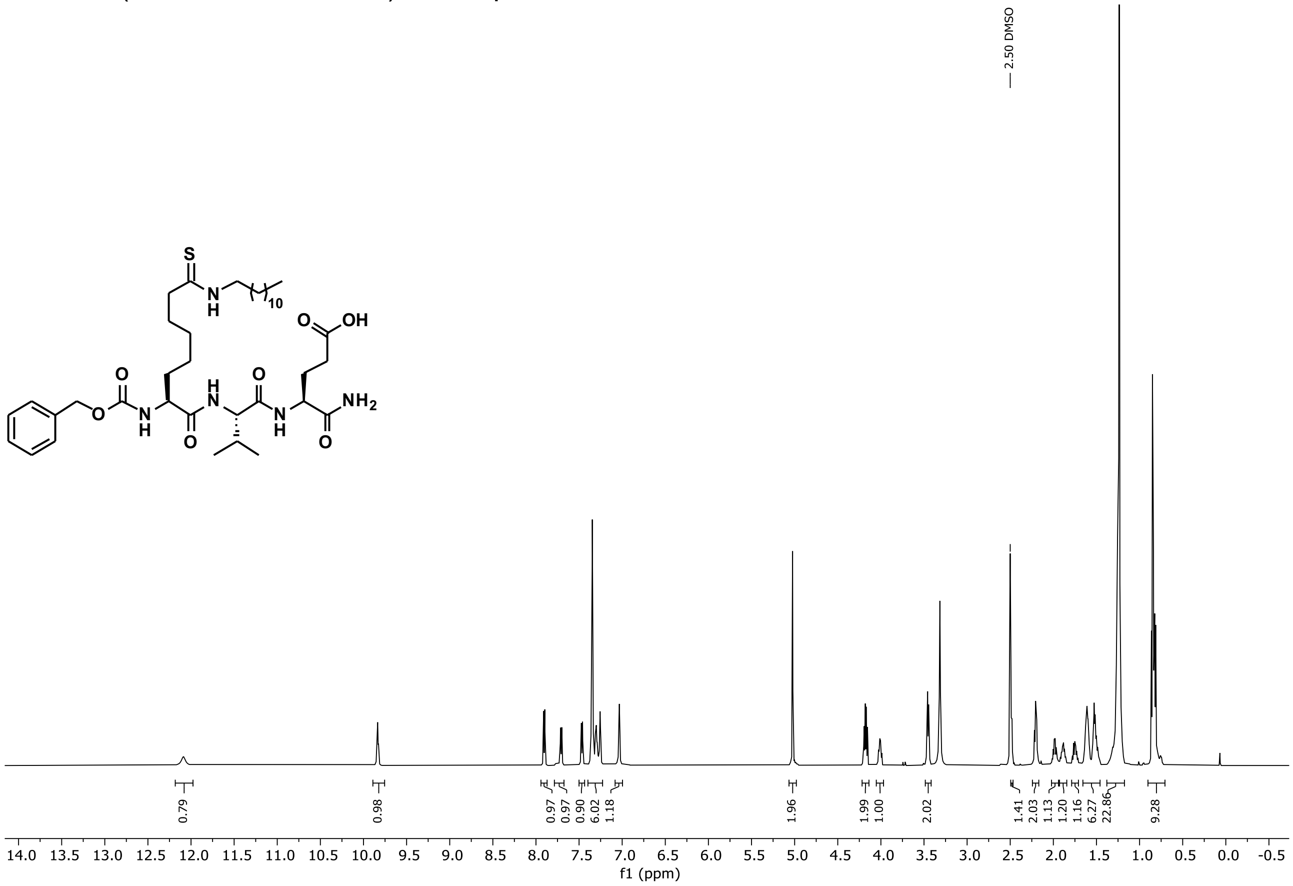

[13]C NMR (151 MHz, DMSO) of compound **34**.

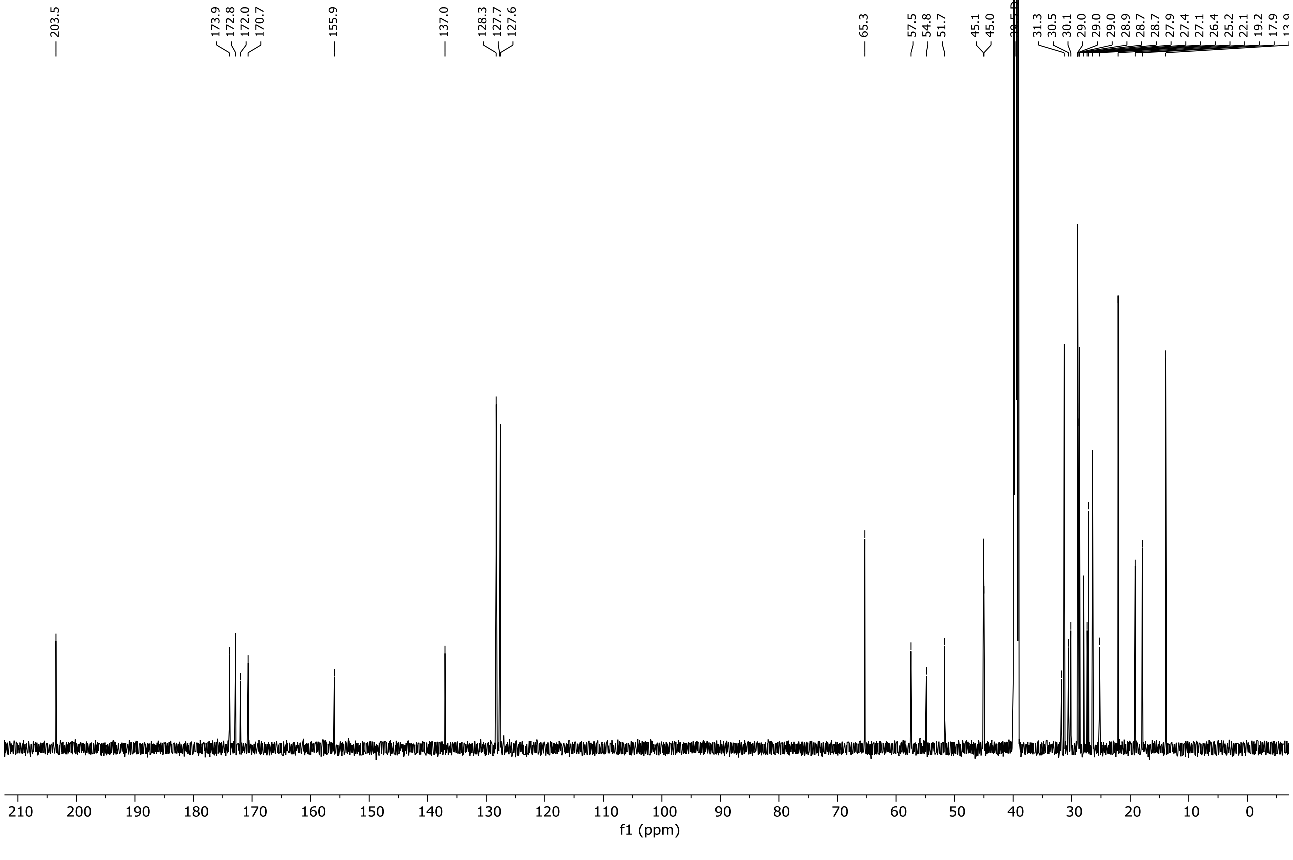

[1]H NMR (600 MHz, DMSO-$d_6$) of compound **35 (IJ4)**.

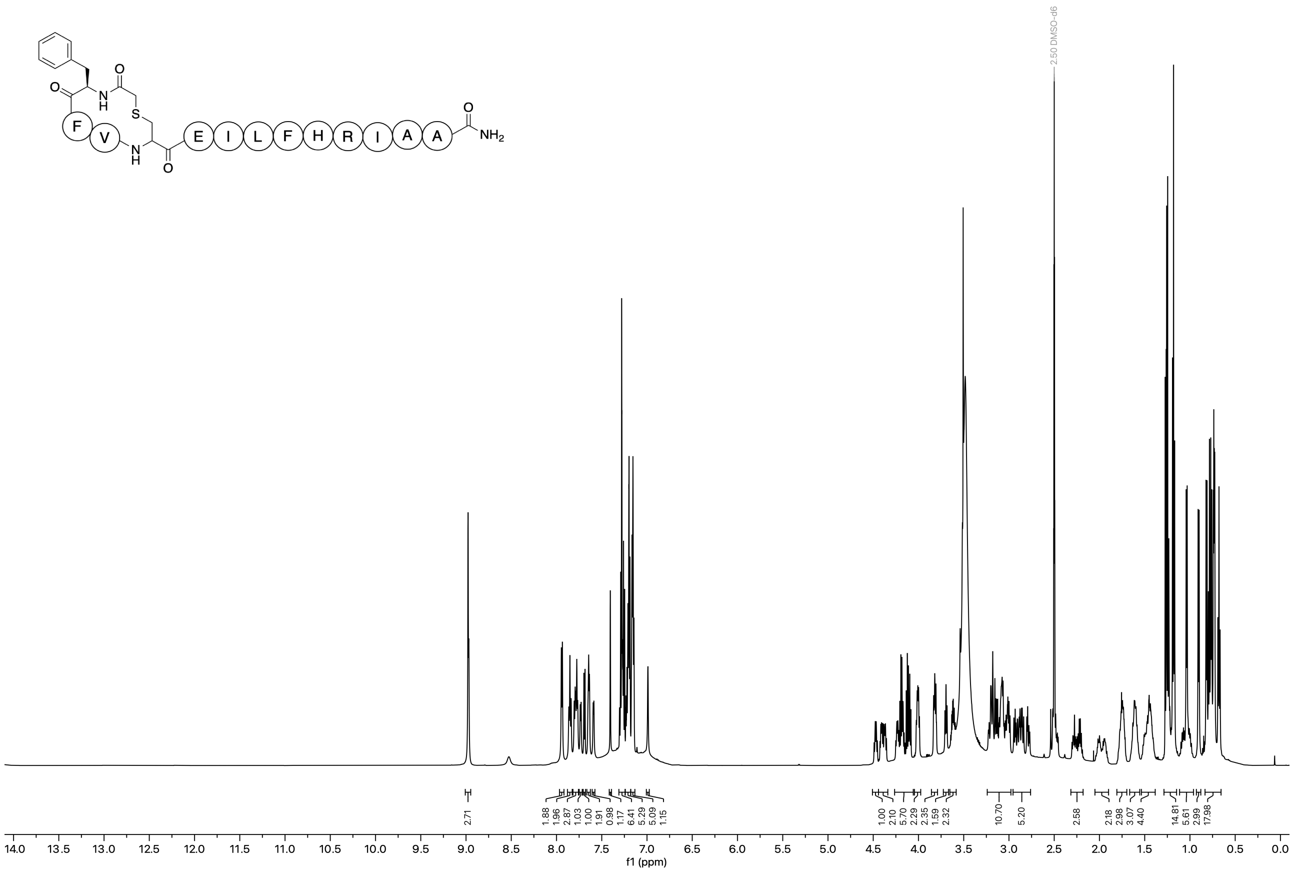

[13]C NMR (151 MHz, DMSO) of compound **35 (IJ4)**.

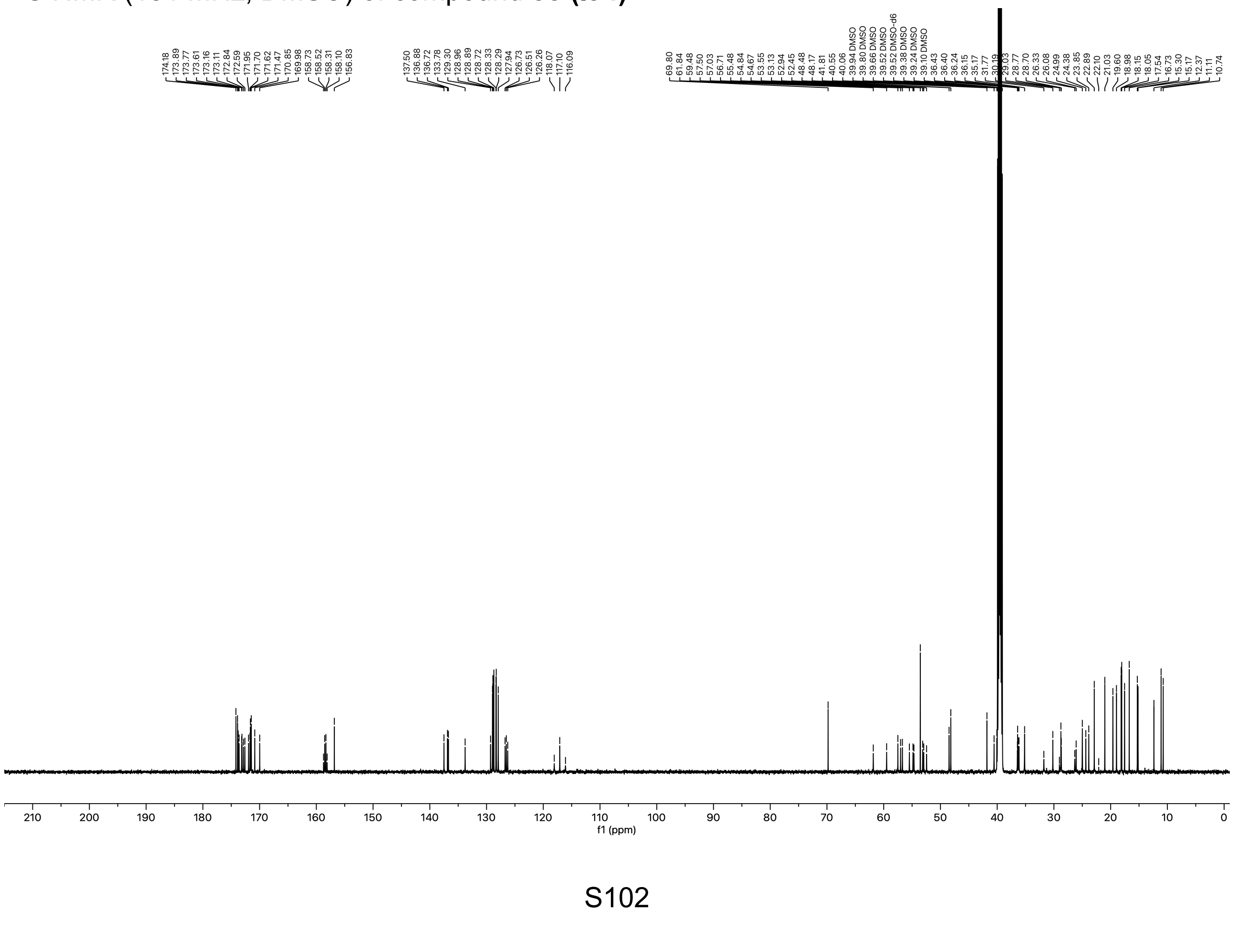

$^{1}H$ NMR (600 MHz, DMSO-$d_6$) of compound **36**.

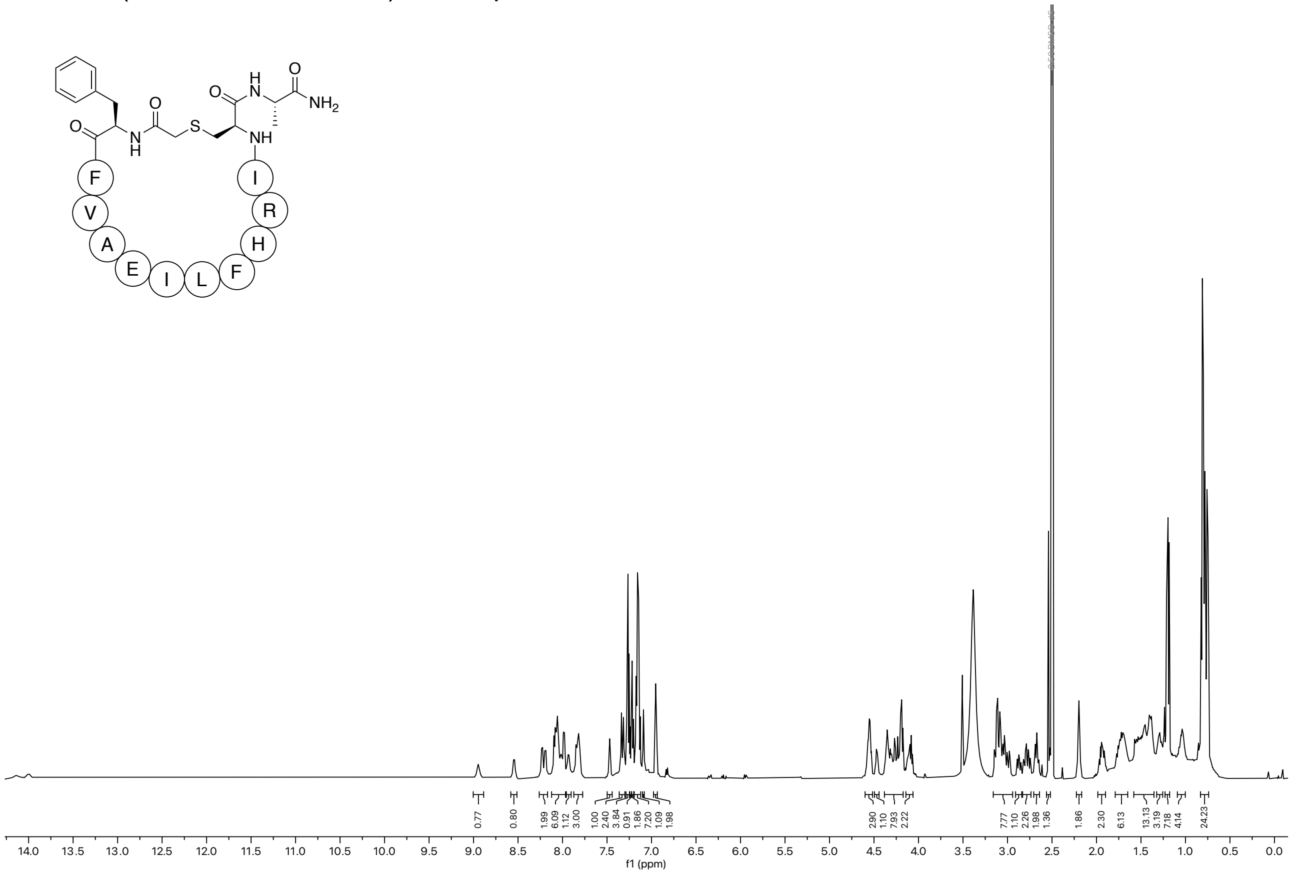

$^{13}C$ NMR (151 MHz, DMSO) of compound **36**.

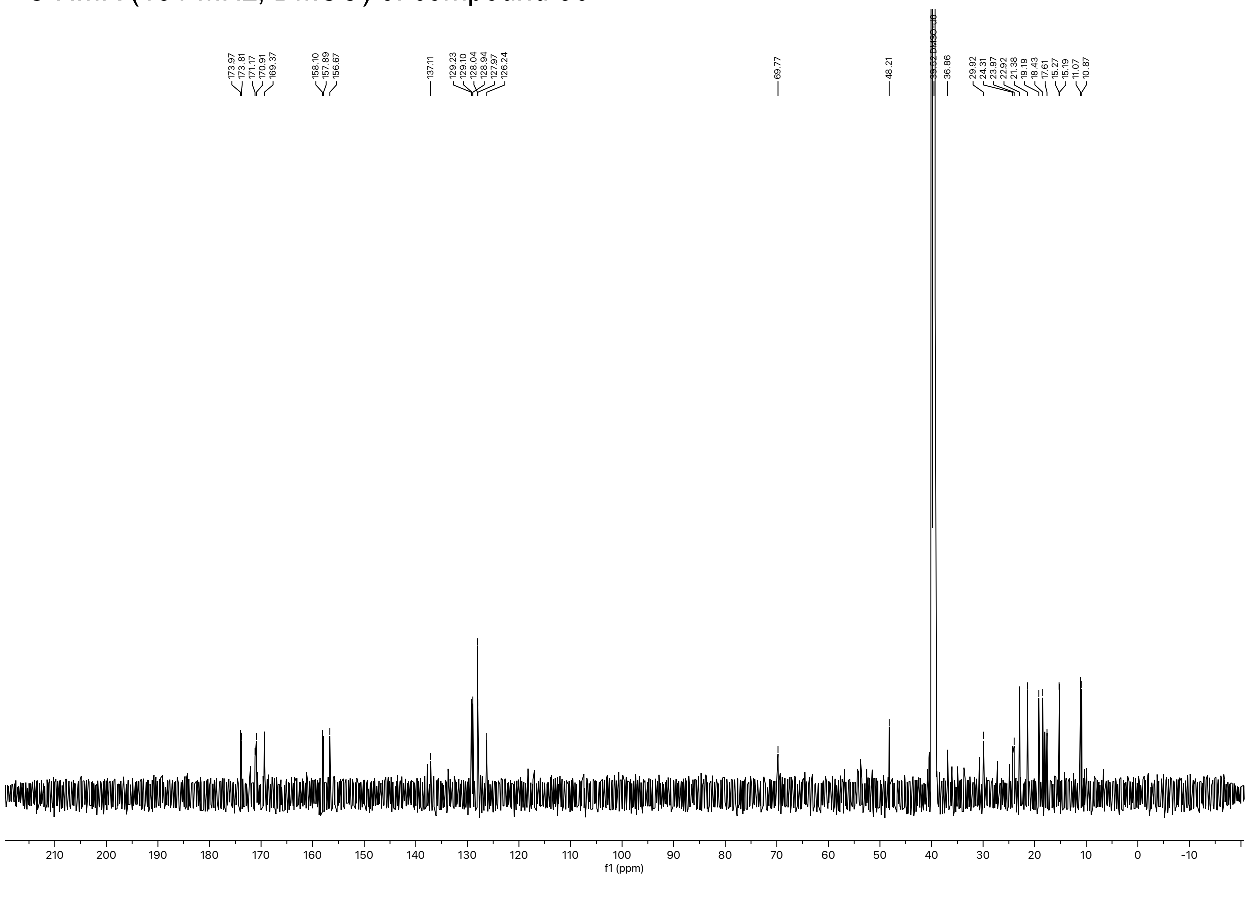

[1]H NMR (600 MHz, DMSO-$d_6$) of compound **37**.

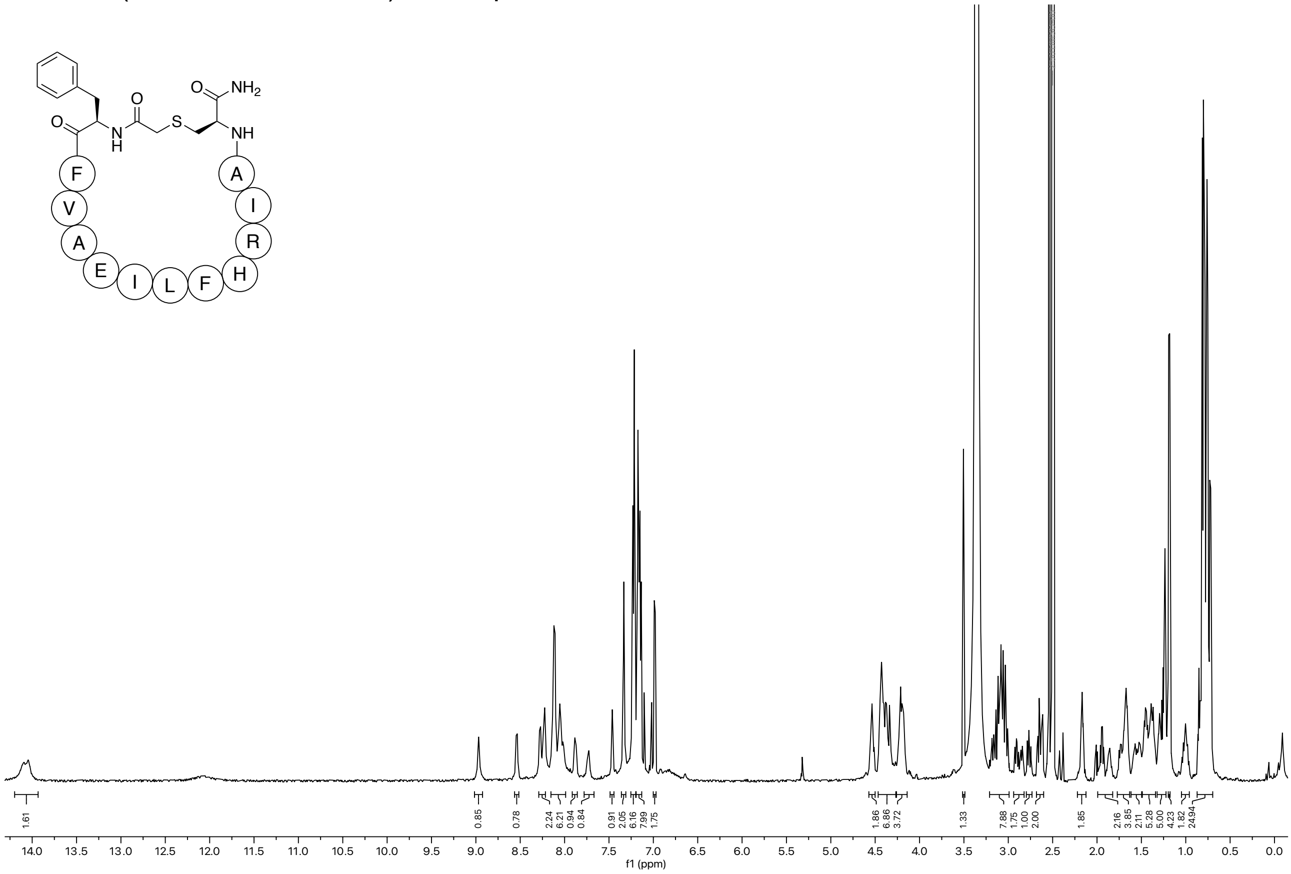

[13]C NMR (151 MHz, DMSO) of compound **37**.

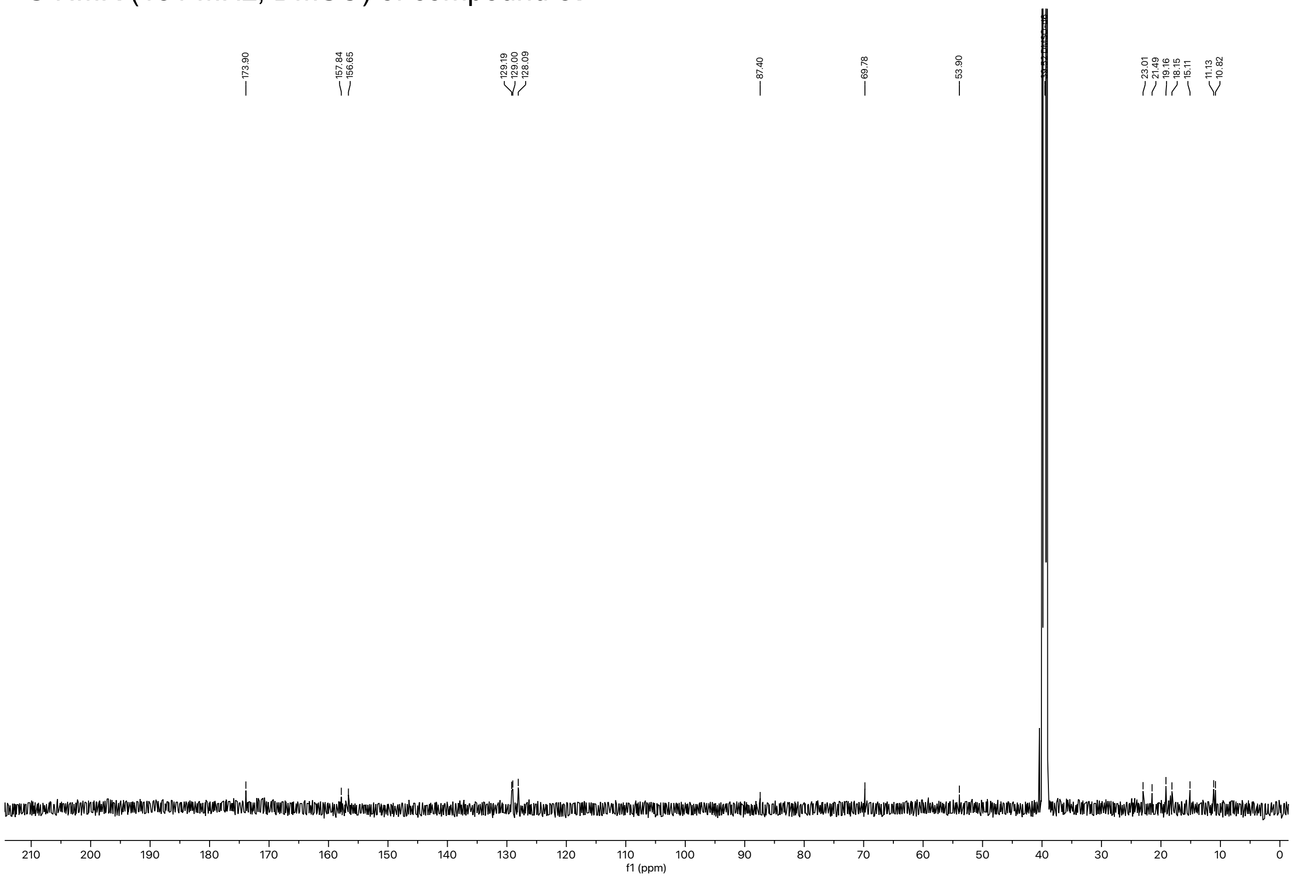

$^{1}H$ NMR (600 MHz, DMSO-$d_6$) of compound **38**.

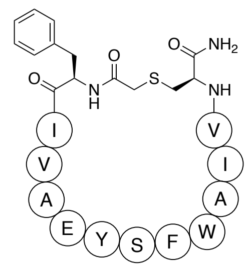

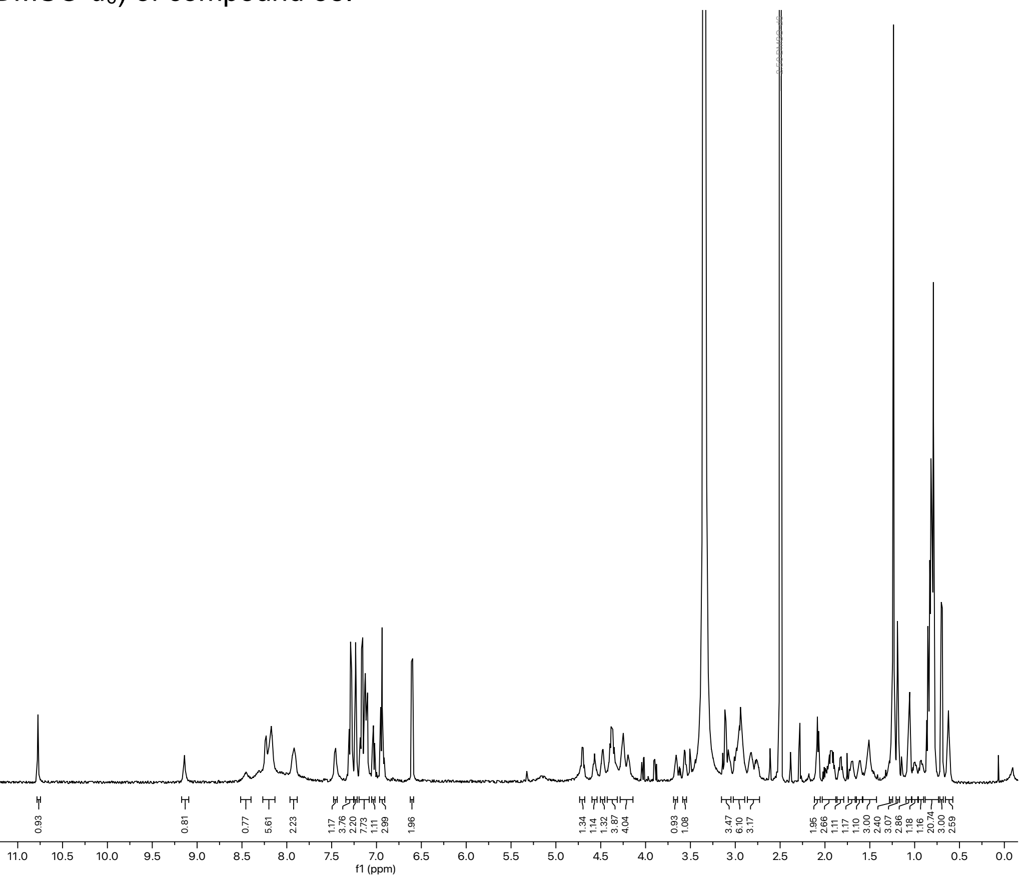

$^{13}C$ NMR (151 MHz, DMSO) of compound **38**.

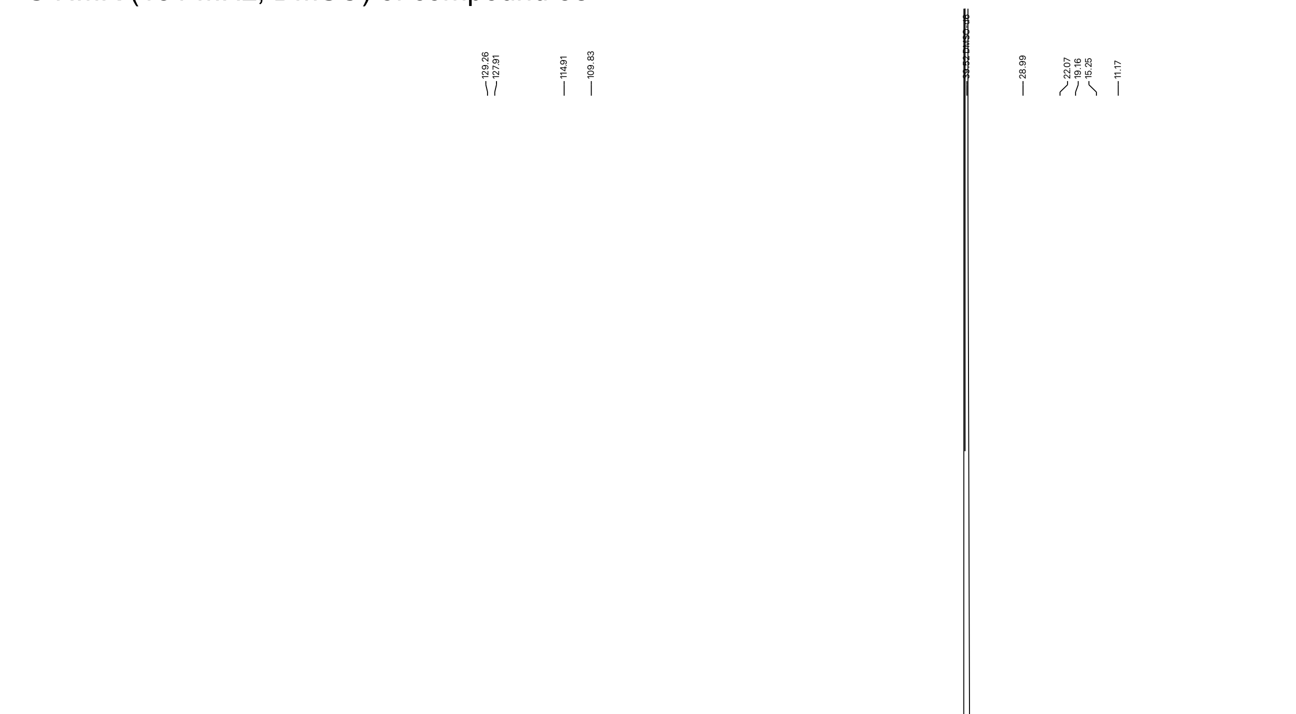

[1]H NMR (600 MHz, DMSO-$d_6$) of compound **39** (**IJ5)**.

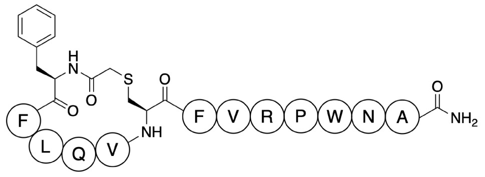

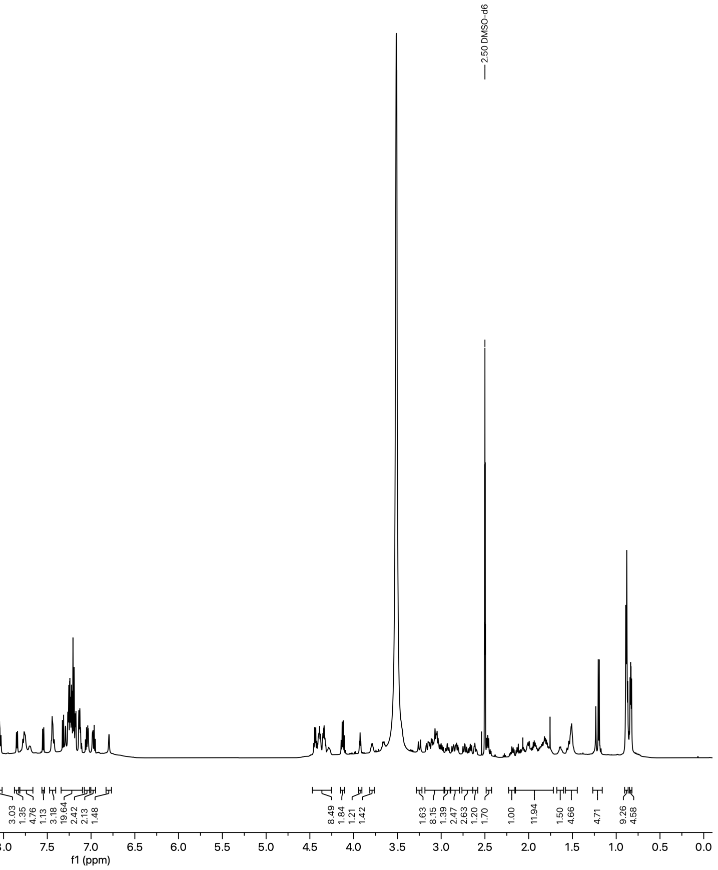

[13]C NMR (151 MHz, DMSO) of compound **39** (**IJ5)**.

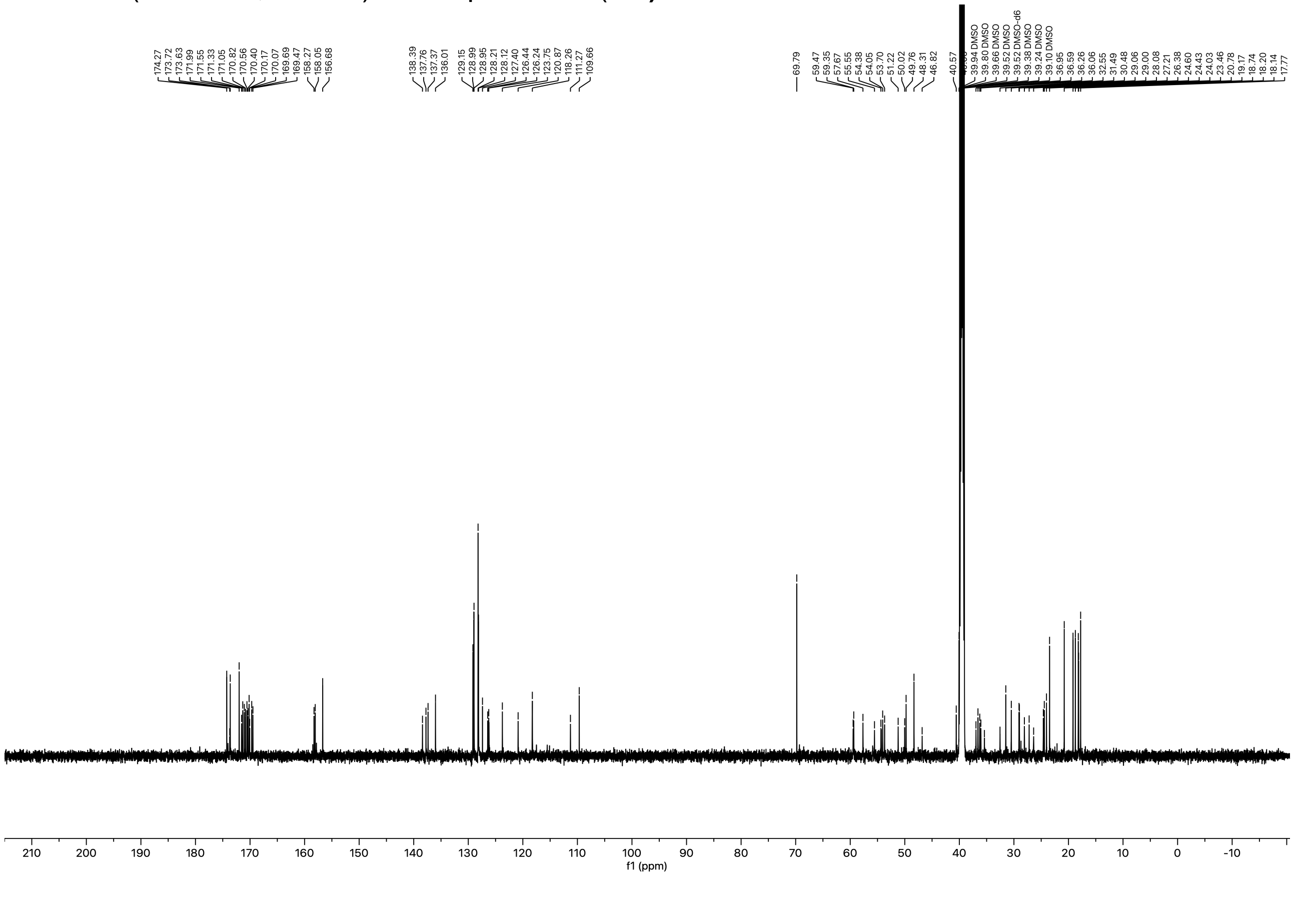

$^{1}H$ NMR (600 MHz, DMSO-$d_6$) of compound **40**.

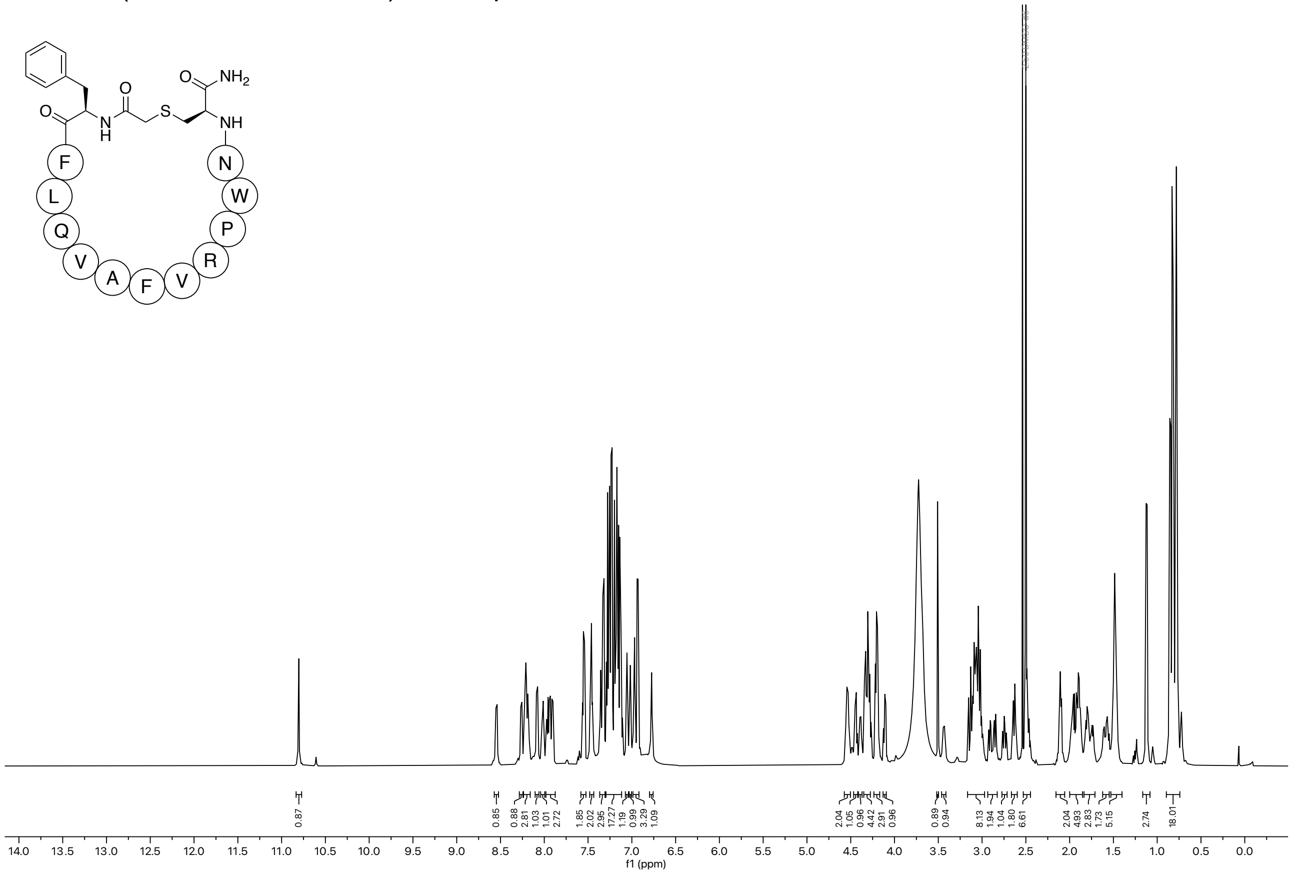

$^{13}C$ NMR (151 MHz, DMSO) of compound **40**.

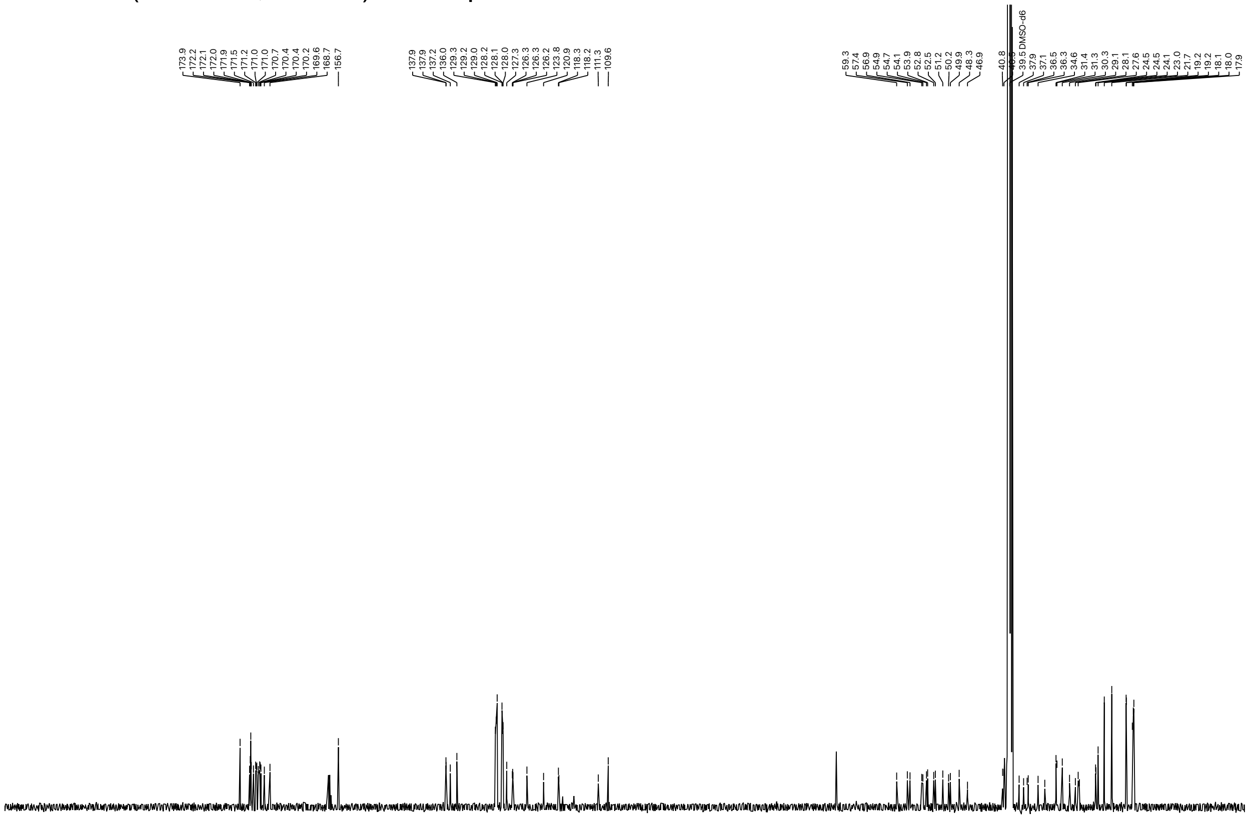

$^{1}$H NMR (600 MHz, DMSO-$d_6$) of compound **41**.

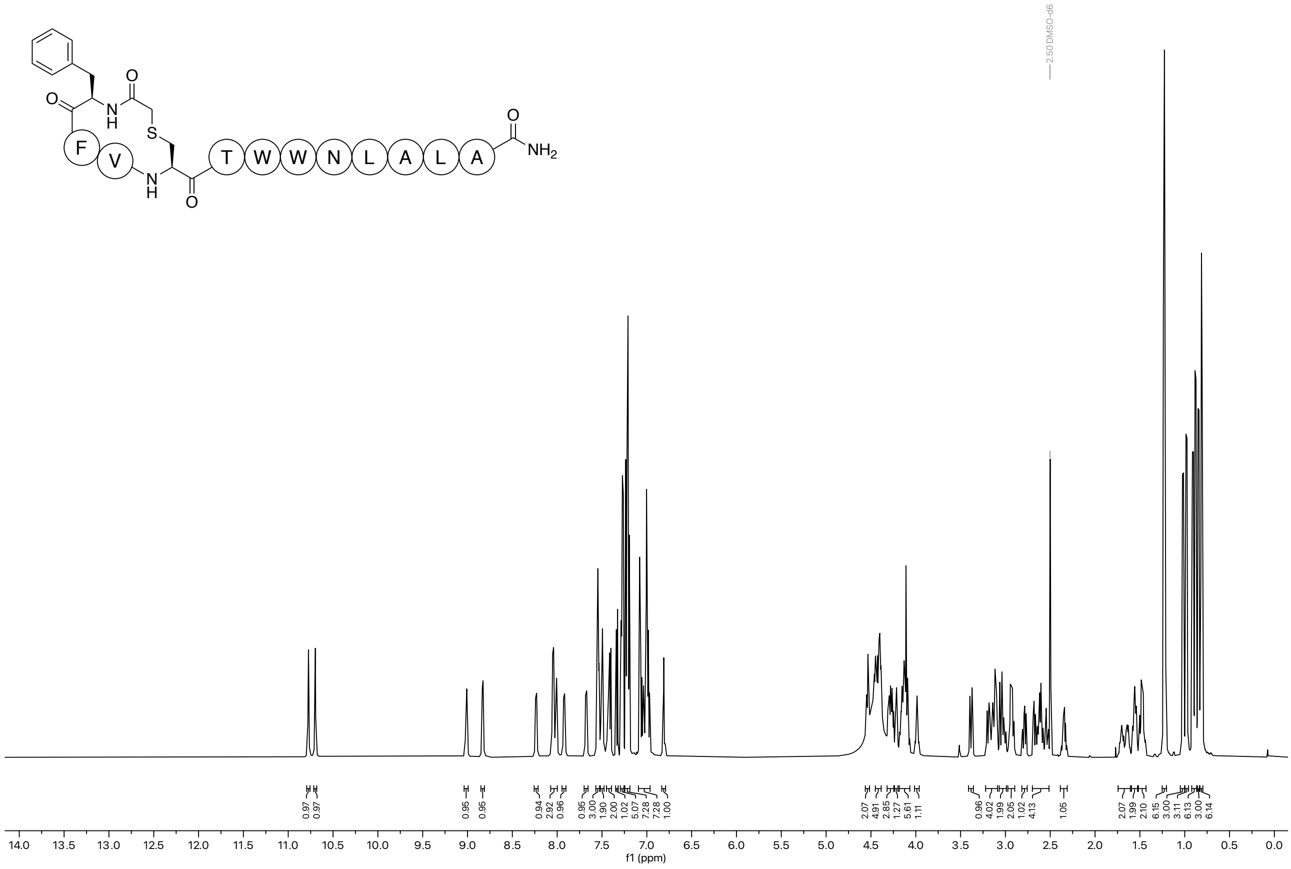

$^{13}$C NMR (151 MHz, DMSO) of compound **41**.

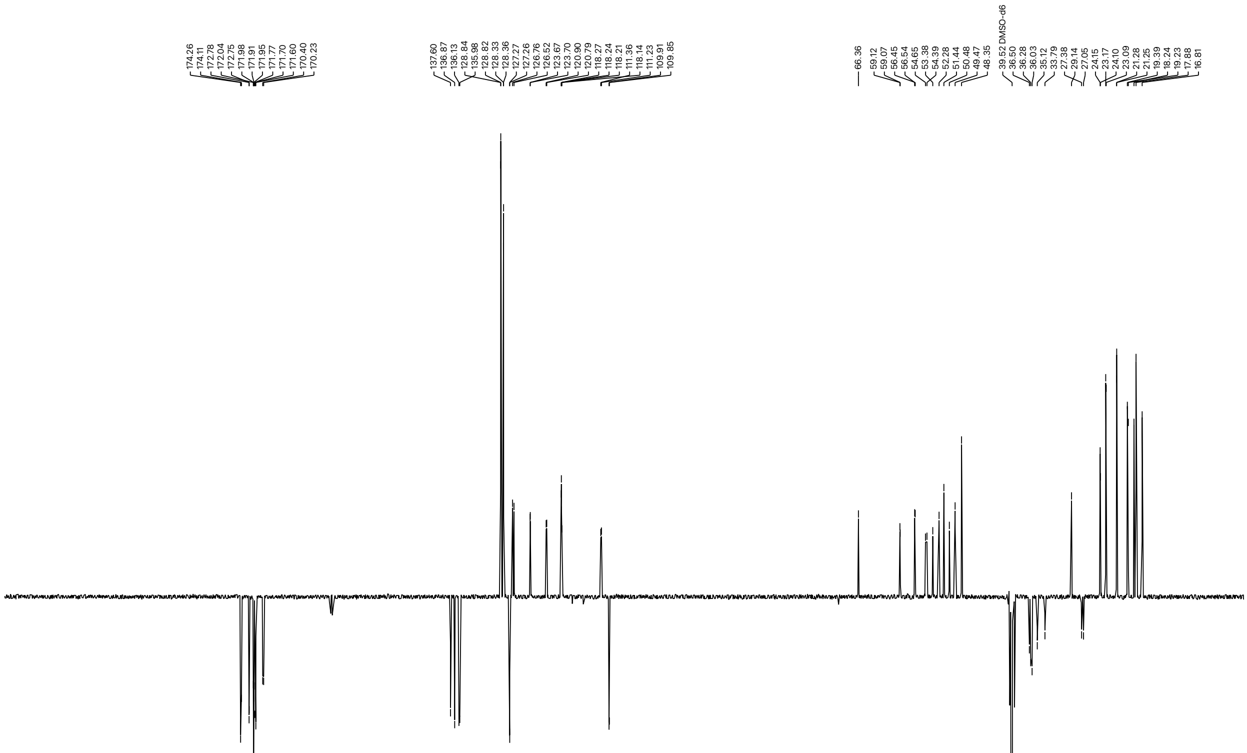

$^{1}$H NMR (600 MHz, DMSO-$d_6$) of compound **42**.

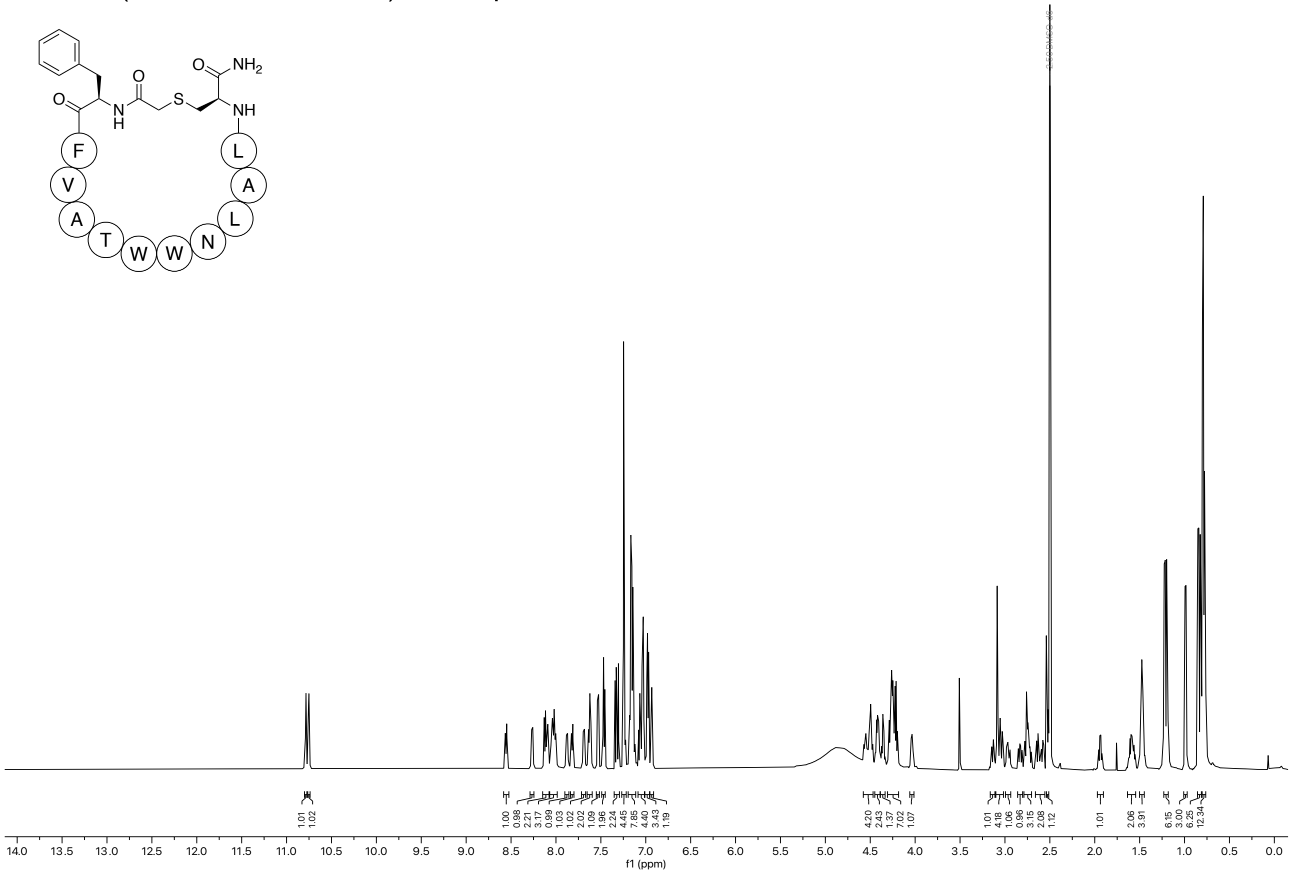

$^{13}$C NMR (151 MHz, DMSO) of compound **42**.

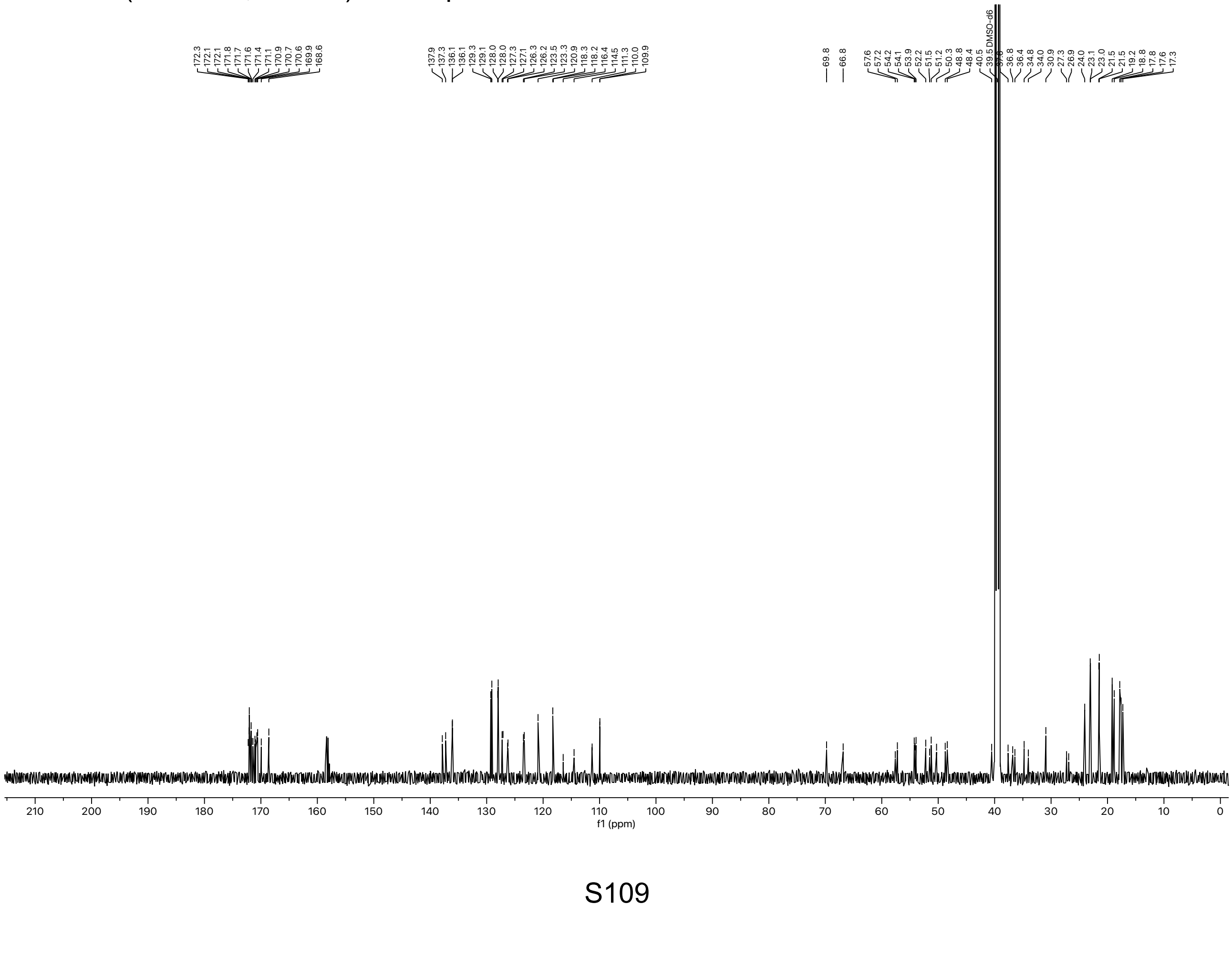

$^{1}$H NMR (600 MHz, DMSO-$d_6$) of compound **43**.

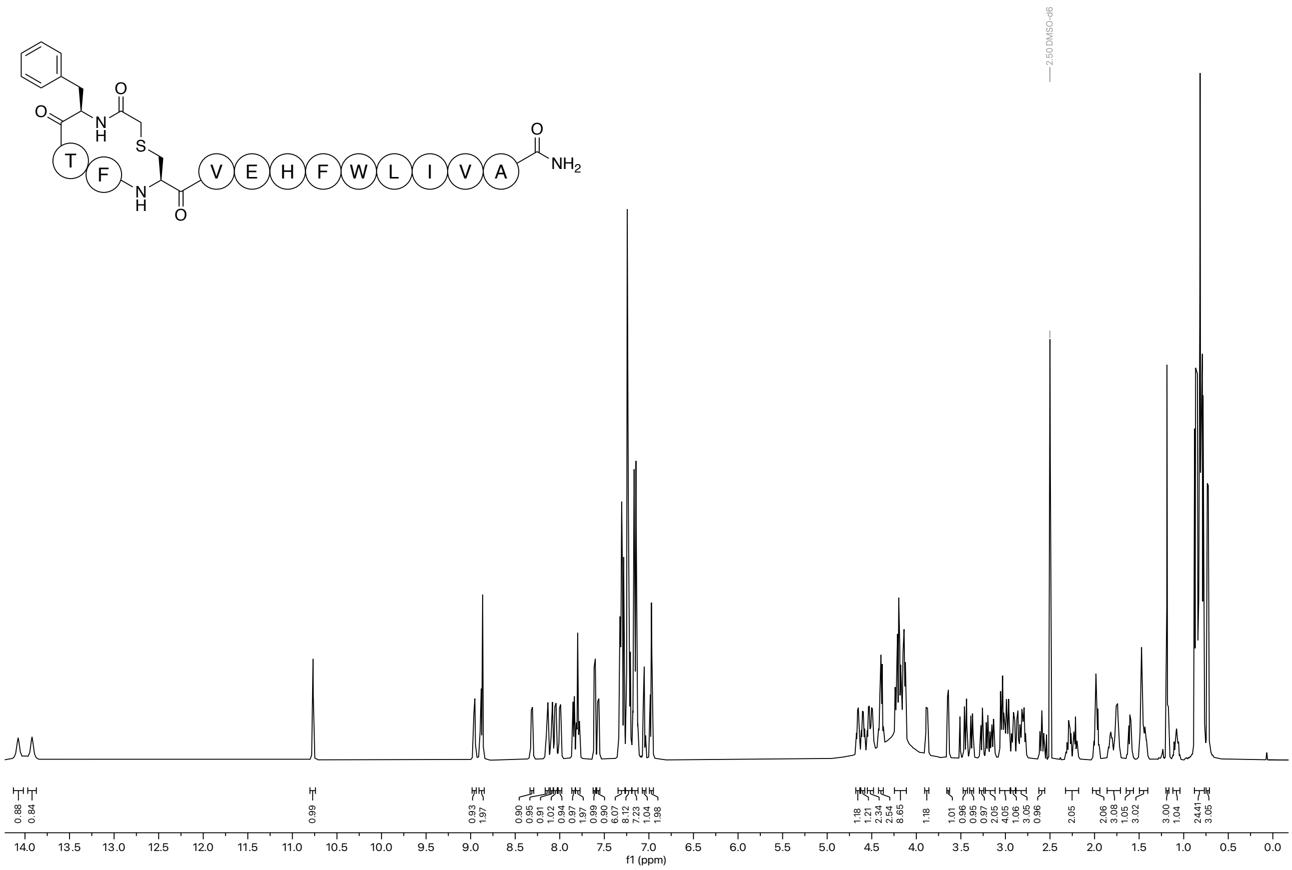

$^{13}$C NMR (151 MHz, DMSO) of compound **43**.

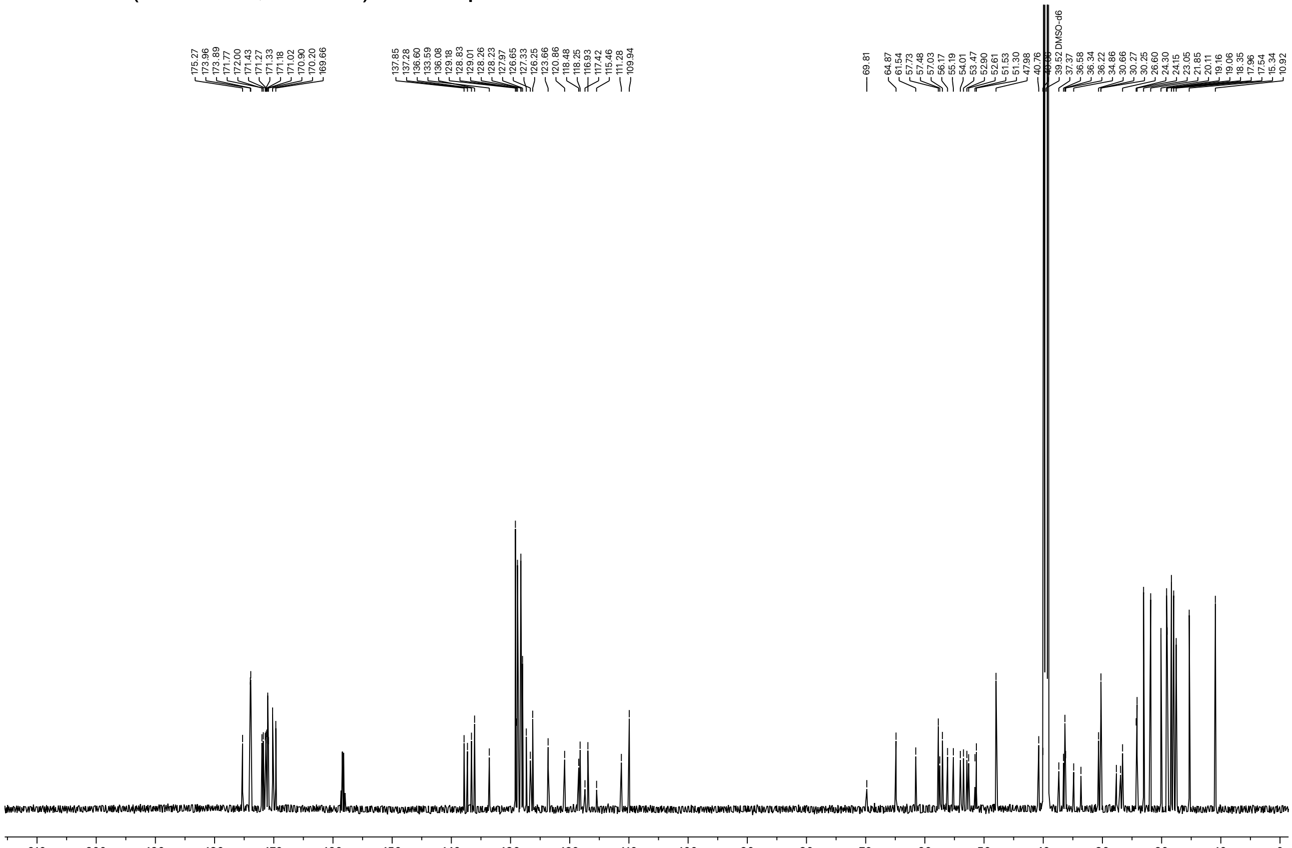

$^{1}H$ NMR (600 MHz, DMSO-$d_6$) of compound **44**.

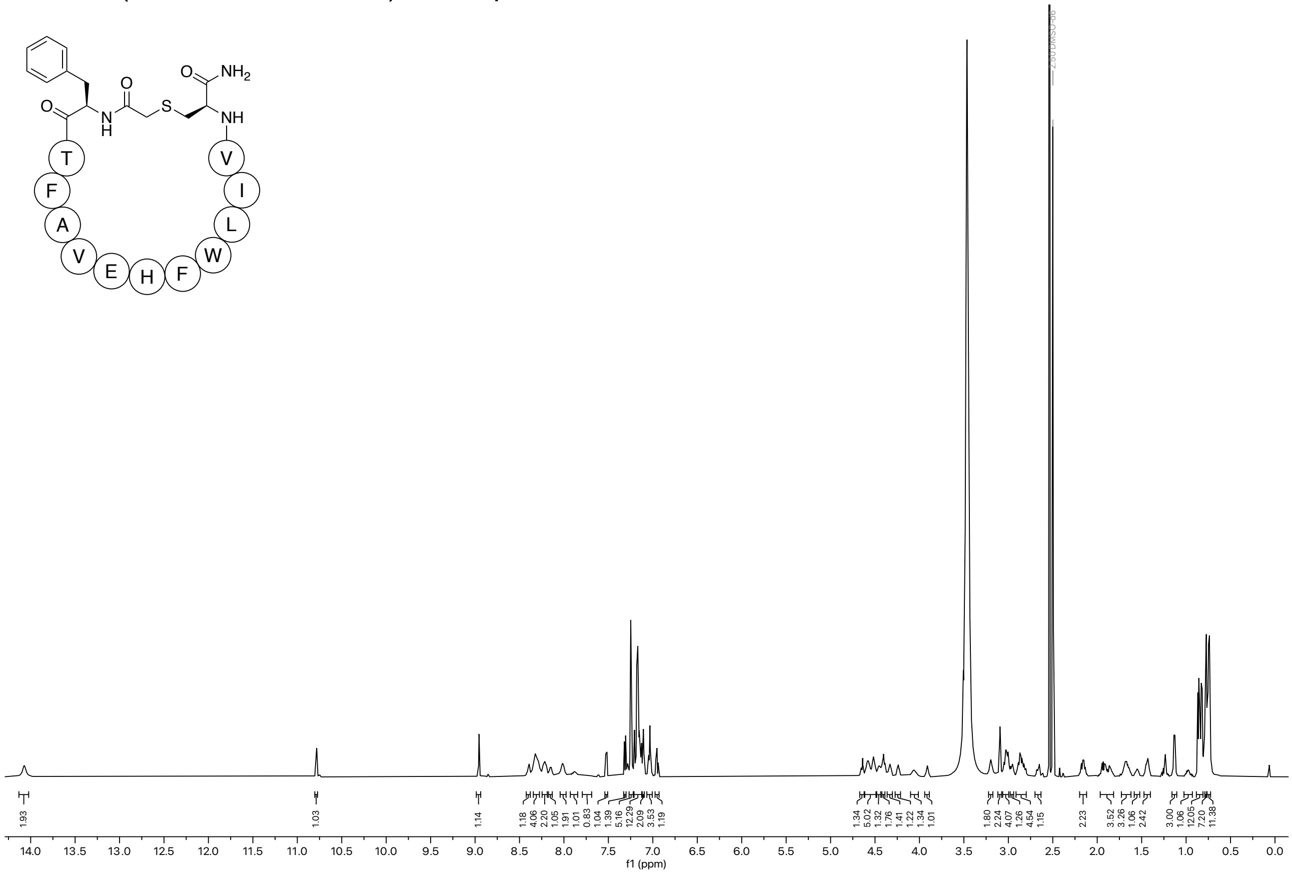

$^{13}C$ NMR (151 MHz, DMSO) of compound **44**.

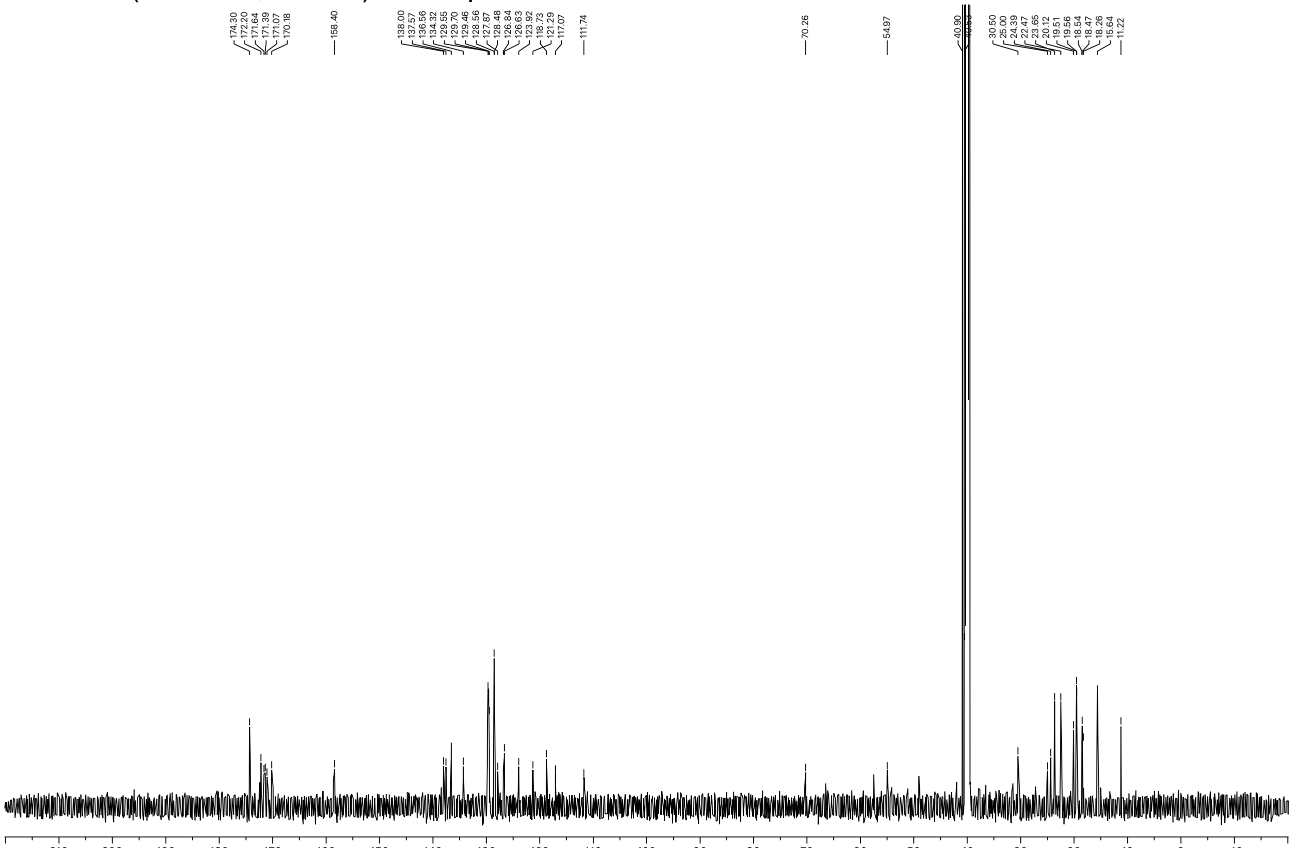

[1]H NMR (600 MHz, DMSO-$d_6$) of compound **45**.

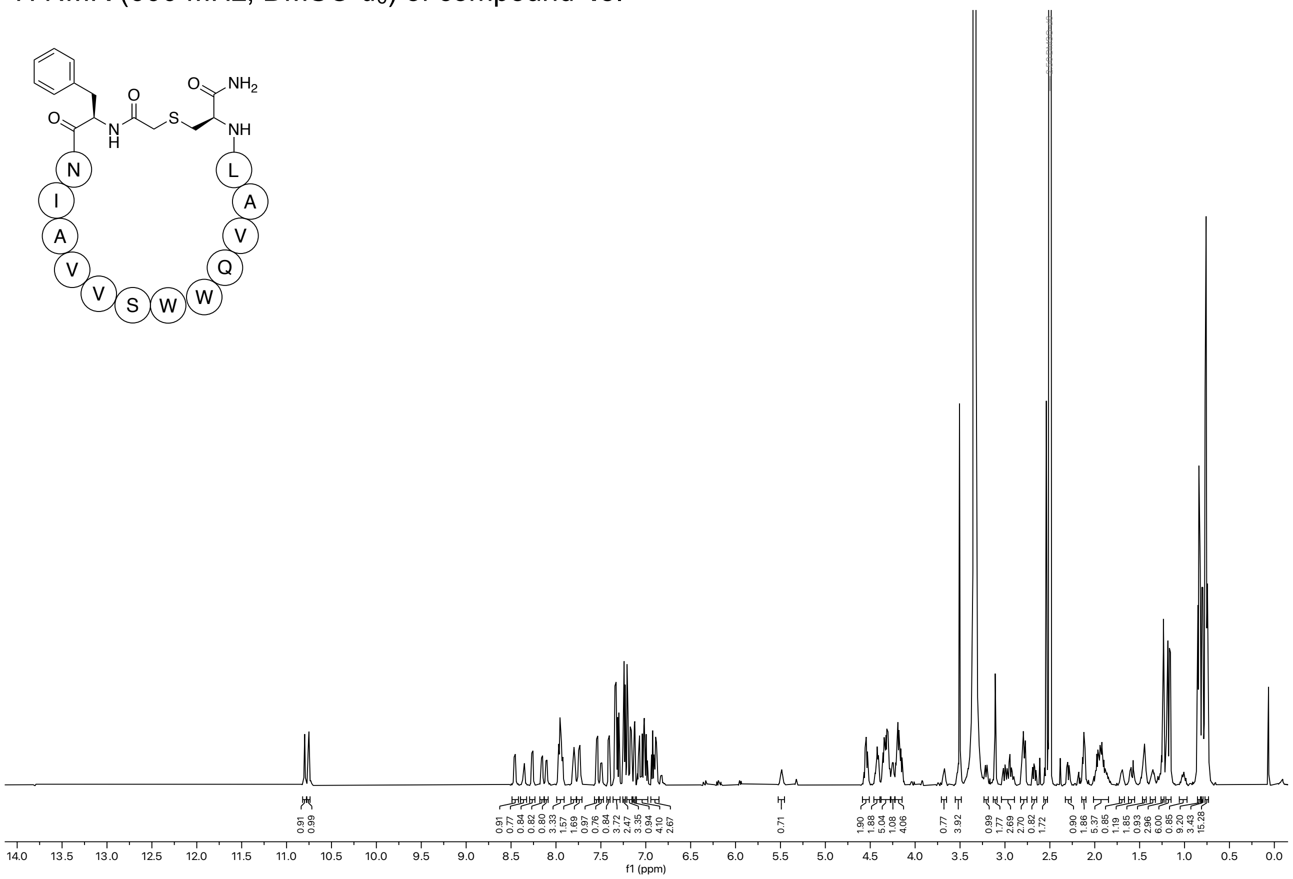

[13]C NMR (151 MHz, DMSO) of compound **45**.

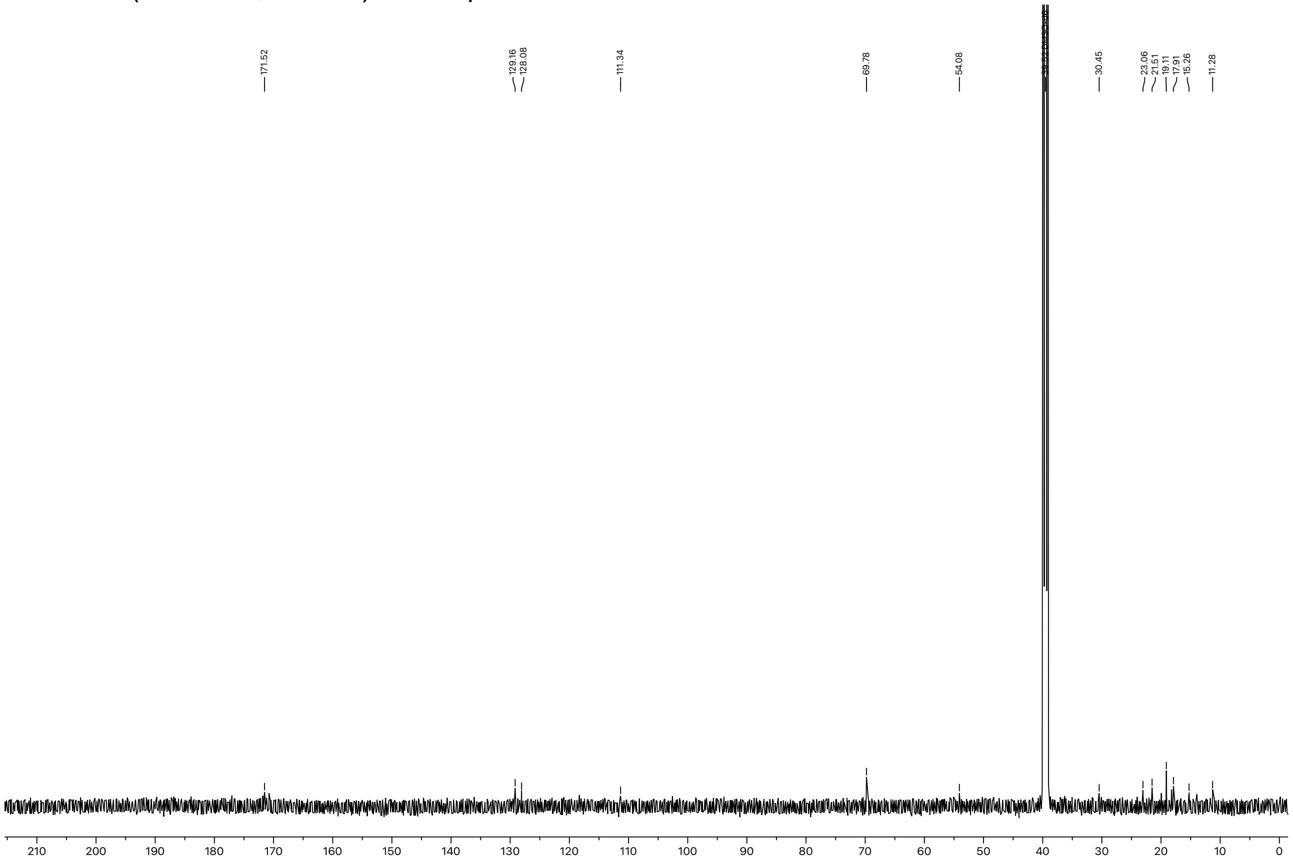

$^{1}$H NMR (600 MHz, DMSO-$d_6$) of compound **46**.

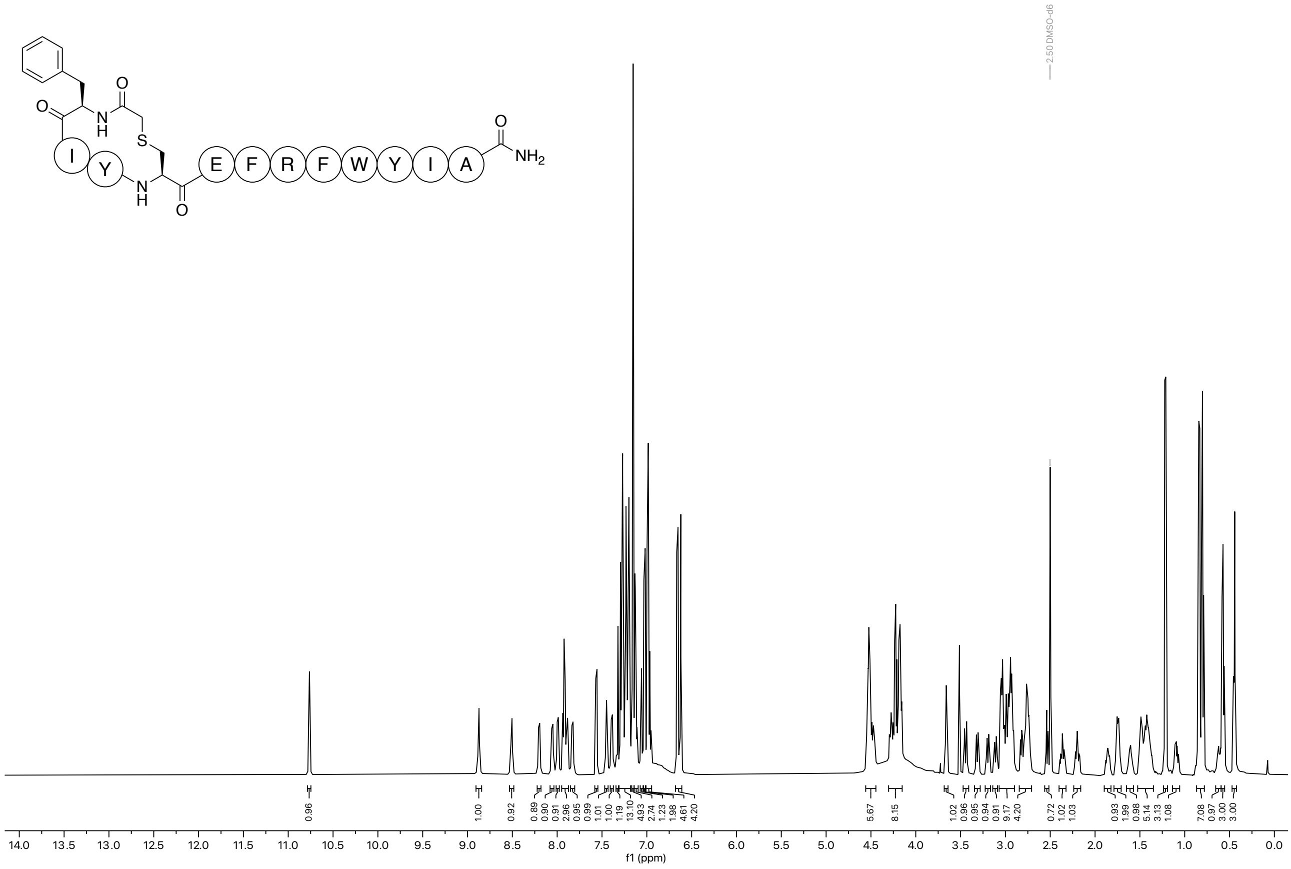

$^{13}$C NMR (151 MHz, DMSO) of compound **46**.

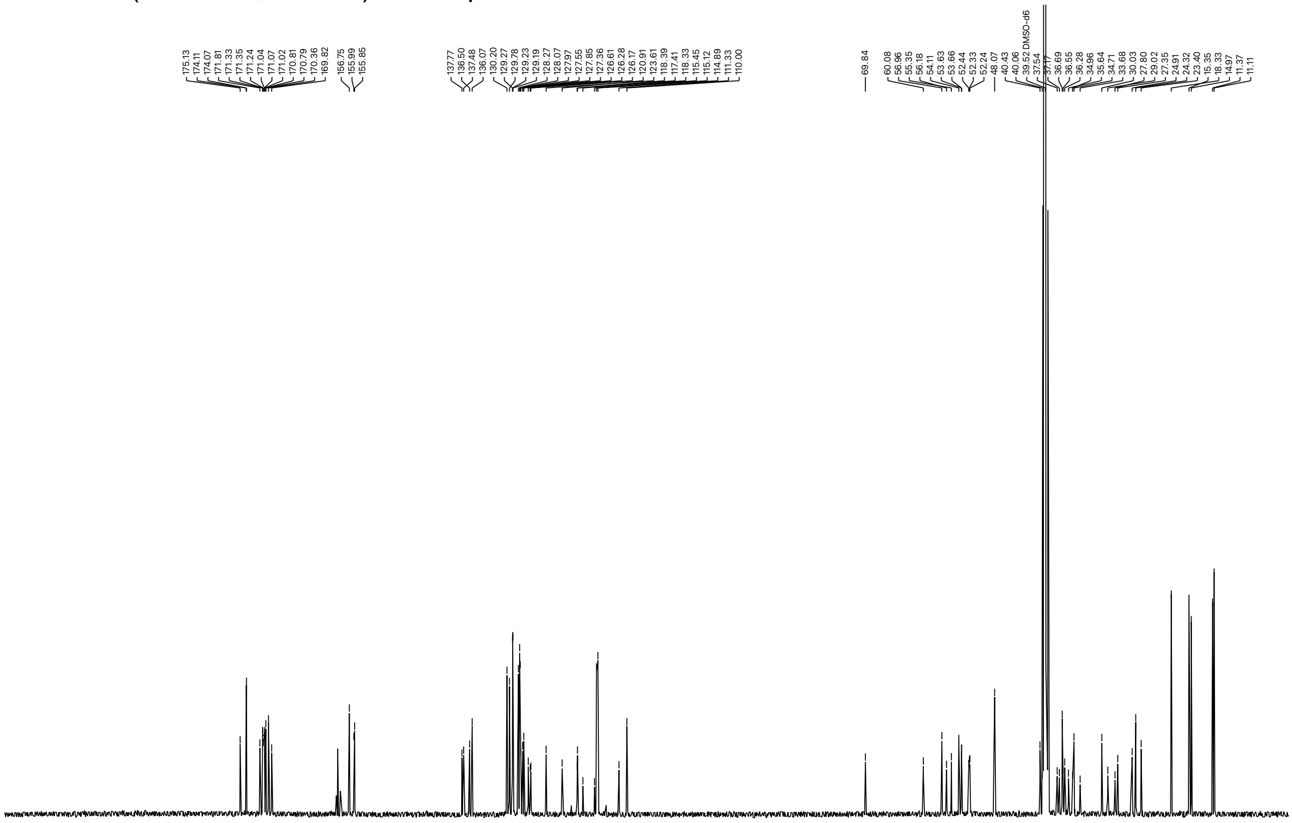

[1]H NMR (600 MHz, DMSO-$d_6$) of compound **47**.

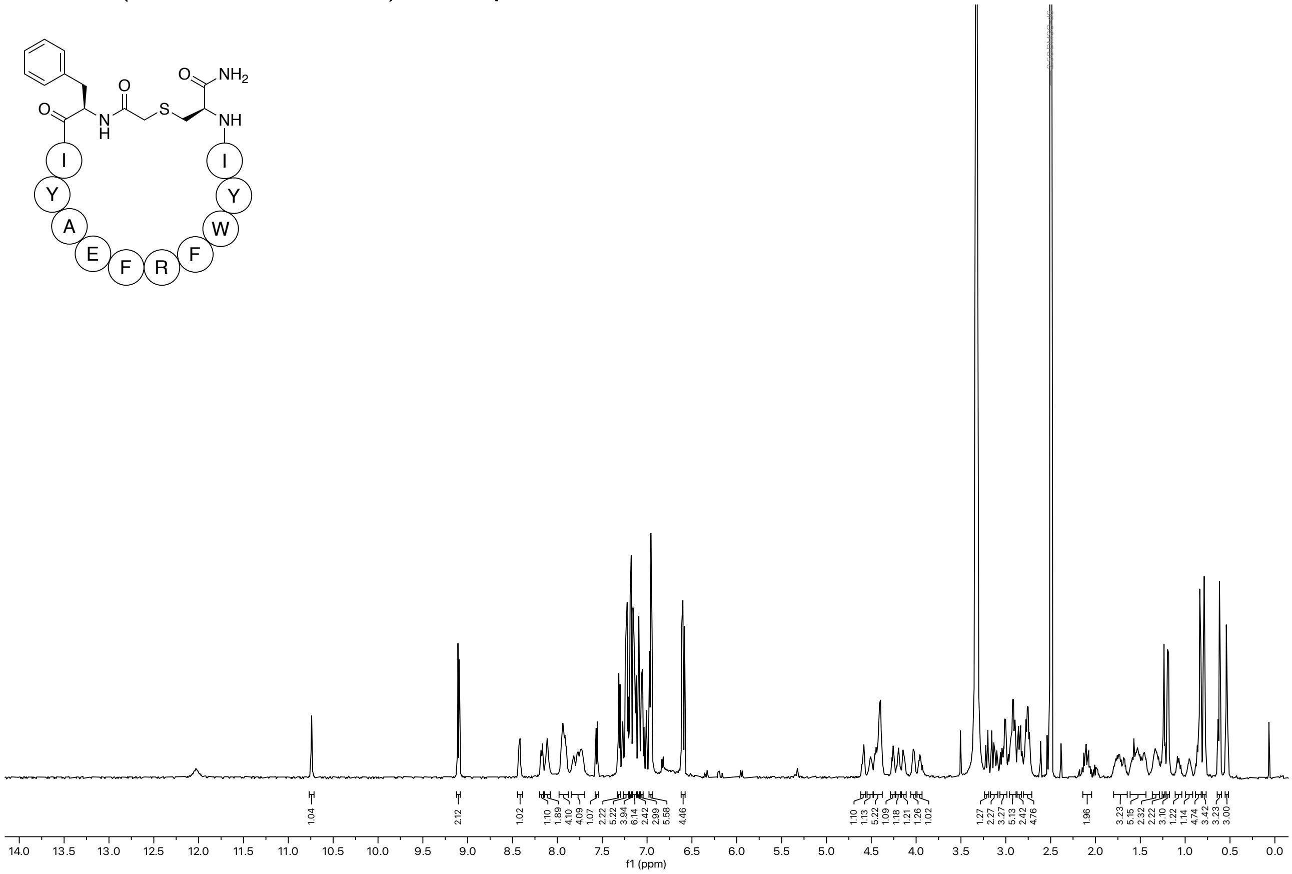

[13]C NMR (151 MHz, DMSO) of compound **47**.

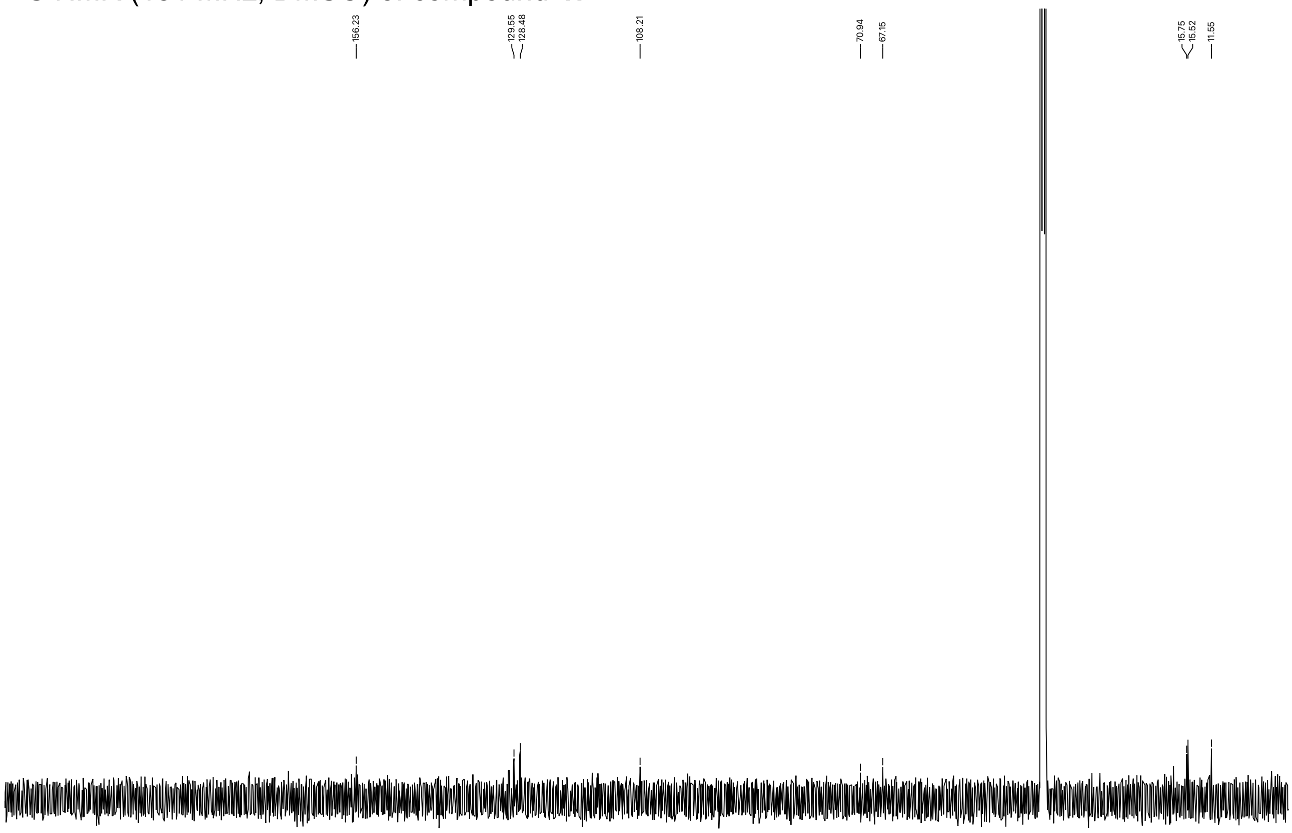

$^1$H NMR (600 MHz, DMSO-$d_6$) of compound **S6**

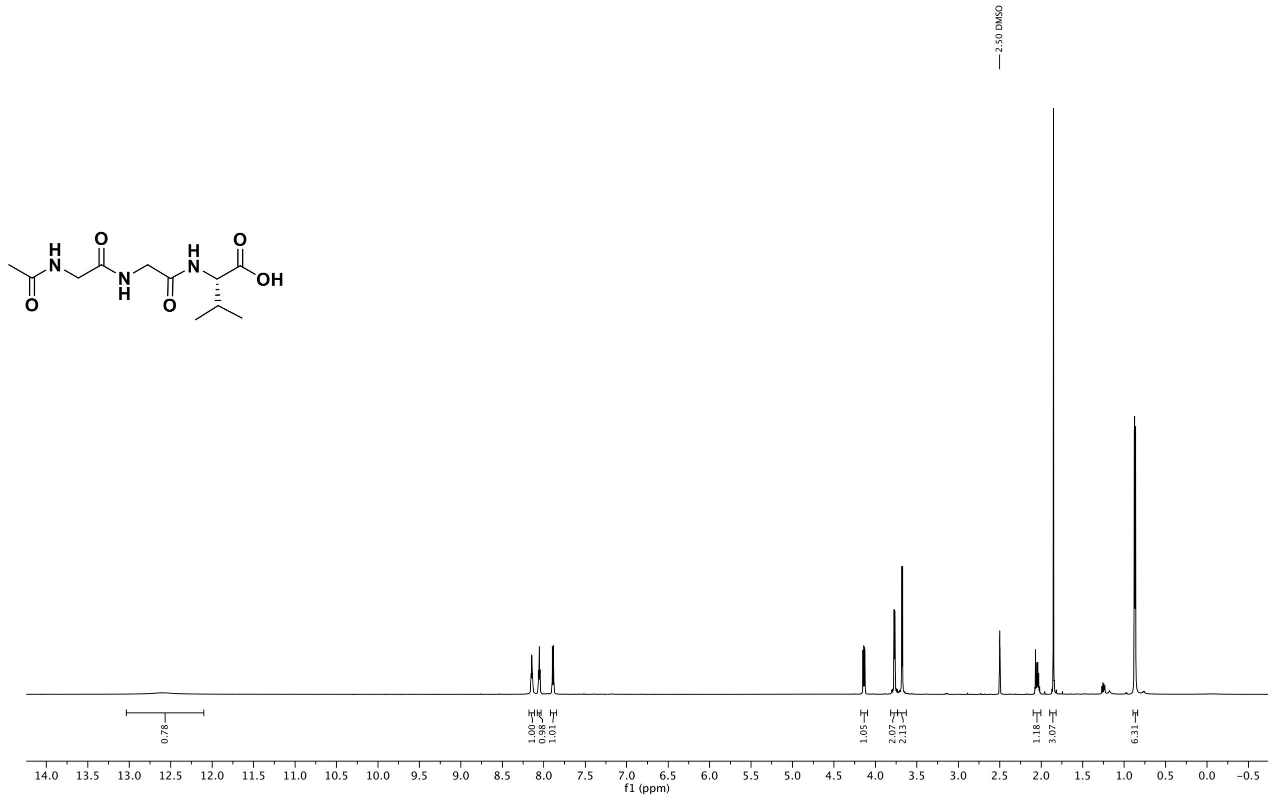

$^{13}$C NMR (151 MHz, DMSO) of compound **S6**

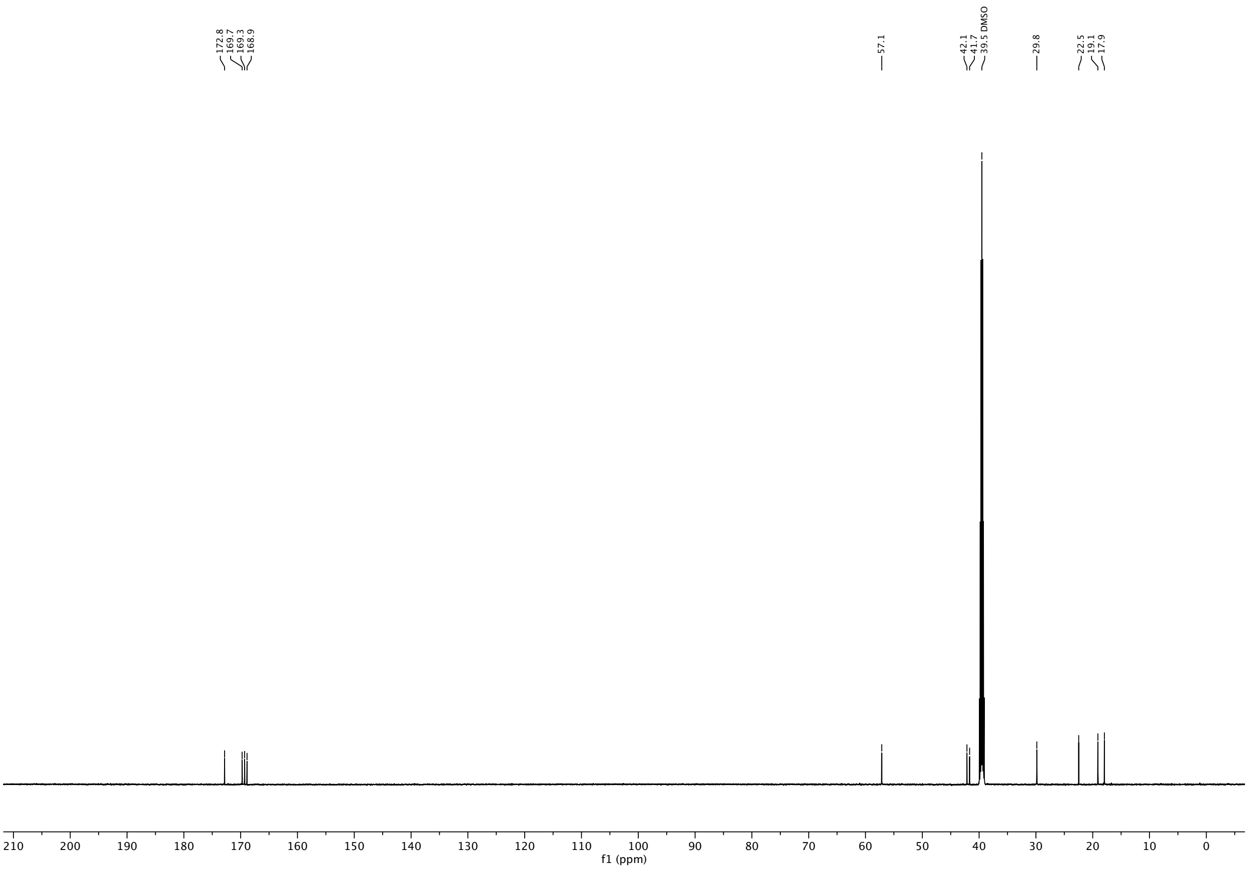

$^{1}$H NMR (600 MHz, DMSO-$d_6$) of compound **S7**

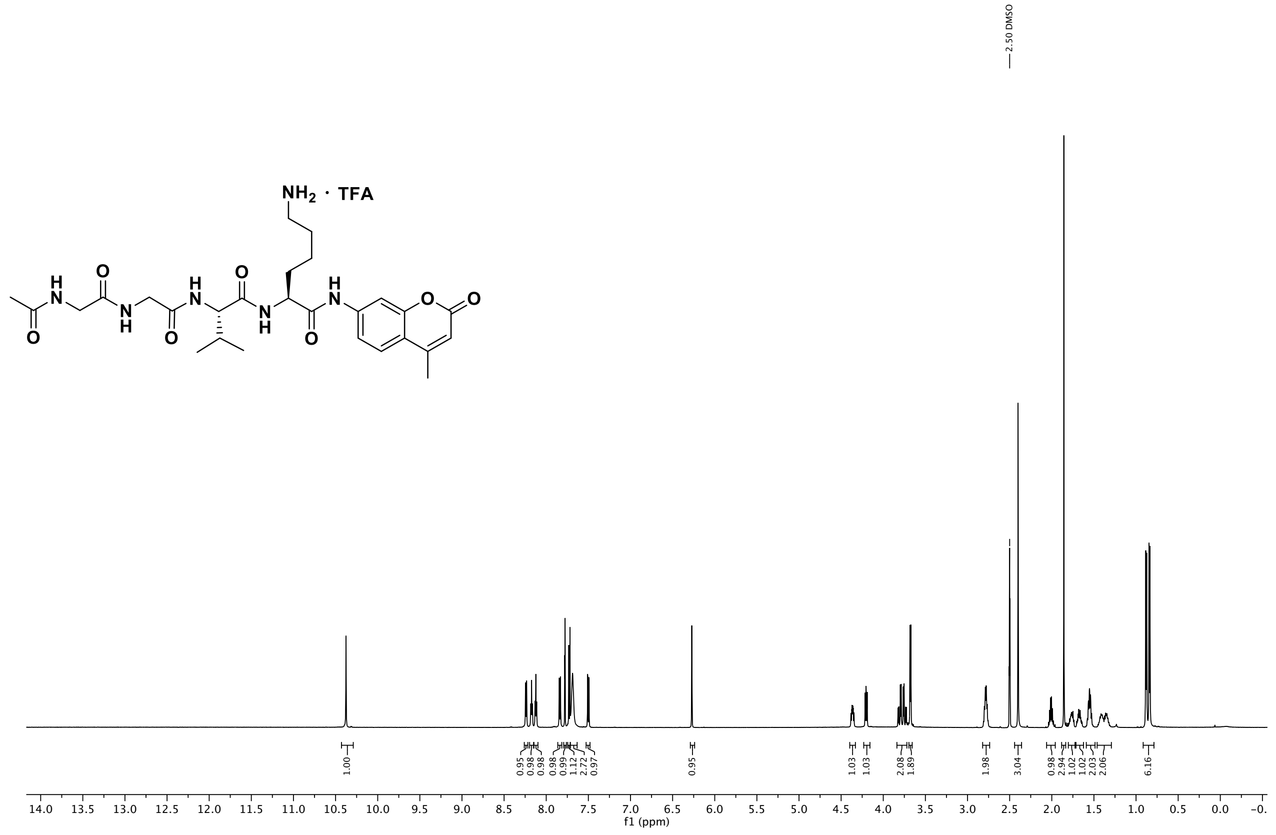

$^{13}$C NMR (151 MHz, DMSO) of compound **S7**

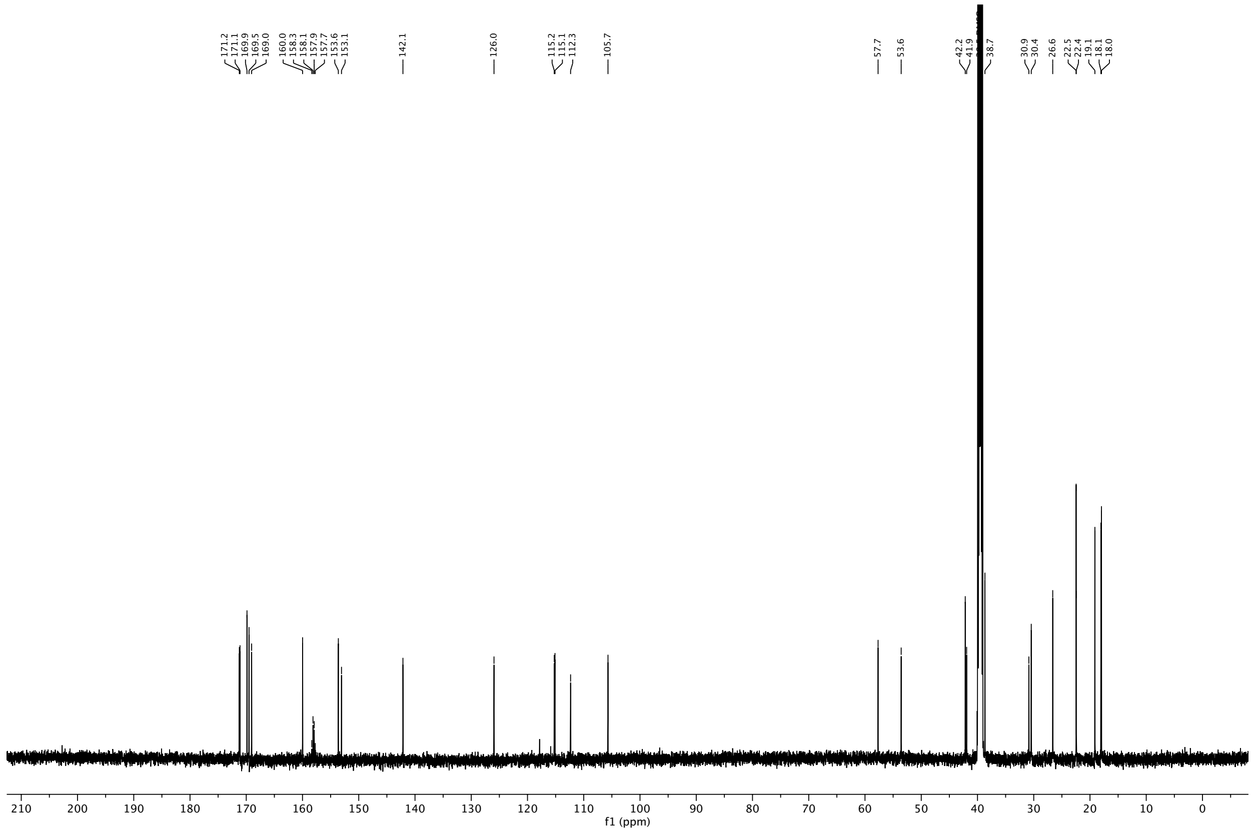

$^{1}H$ NMR (400 MHz, DMSO-$d_6$) of compound **S8**.

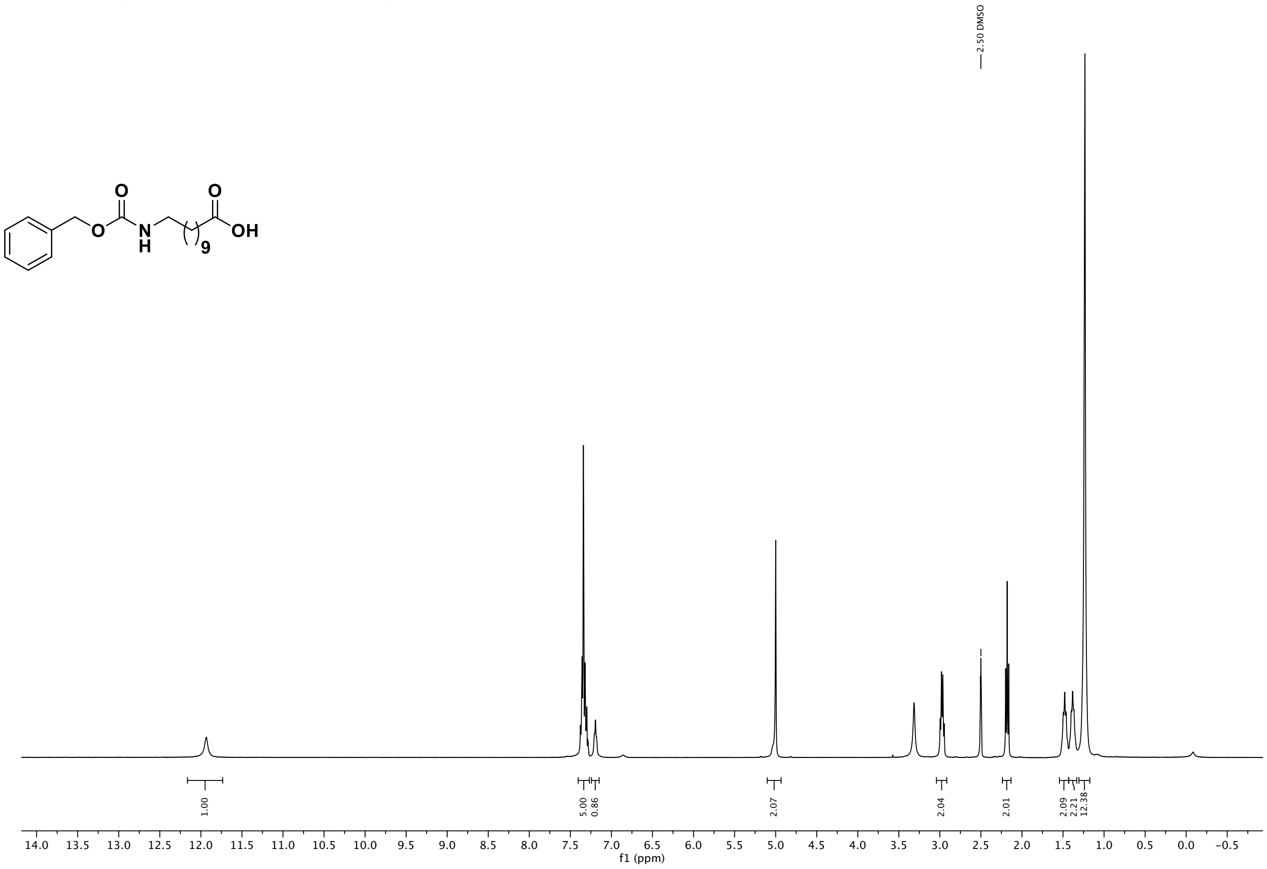

$^{13}C$ NMR (101 MHz, DMSO) of compound **S8**.

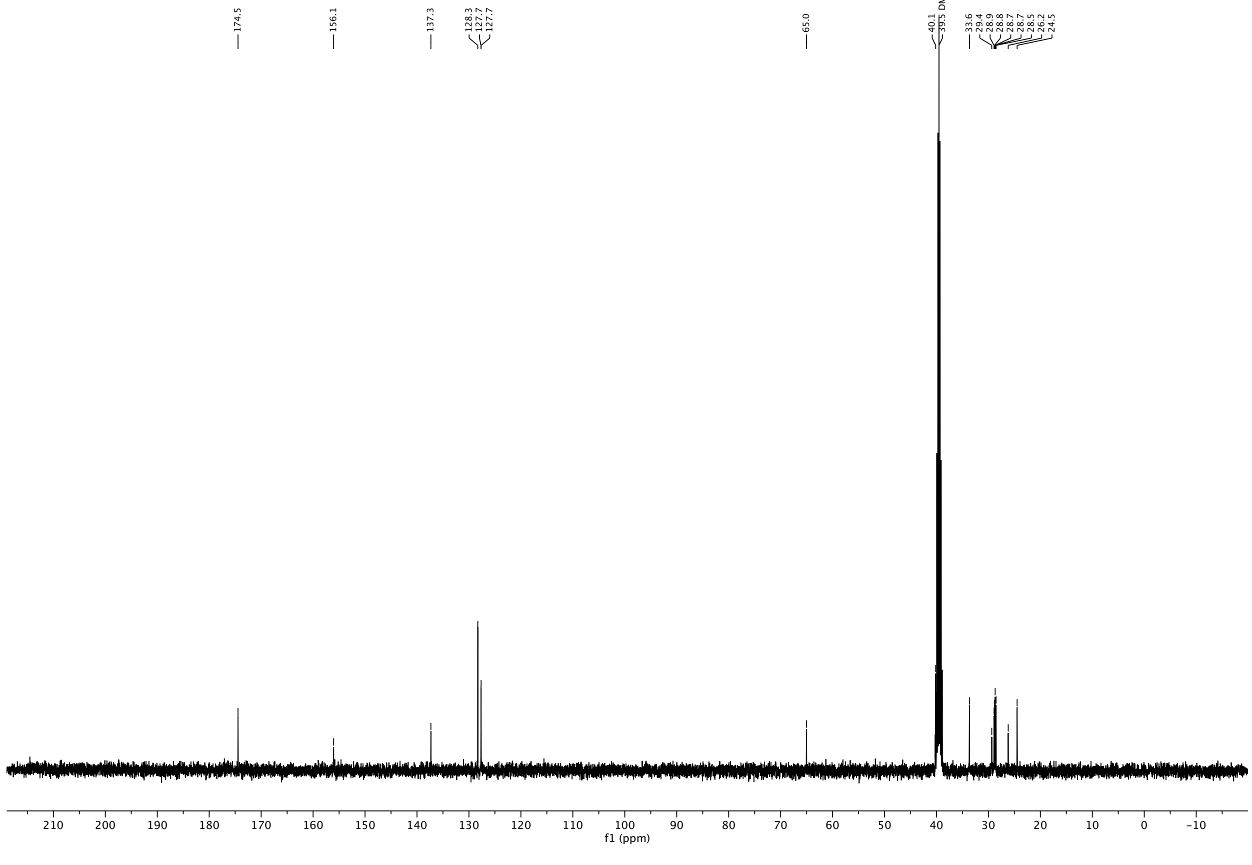

$^{1}H$ NMR (600 MHz, $CDCl_3$) of compound **S9**.

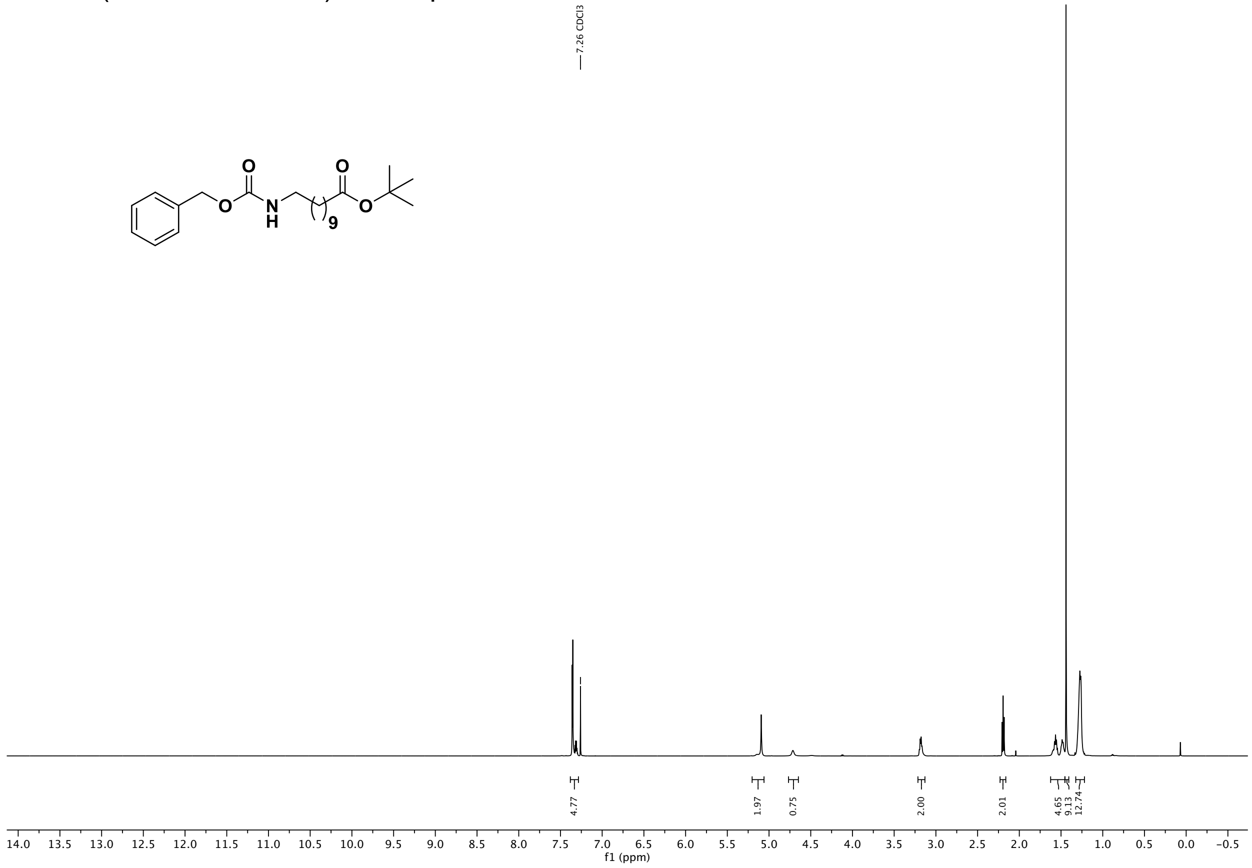

$^{13}C$ NMR (151 MHz, $CDCl_3$) of compound **S9**.

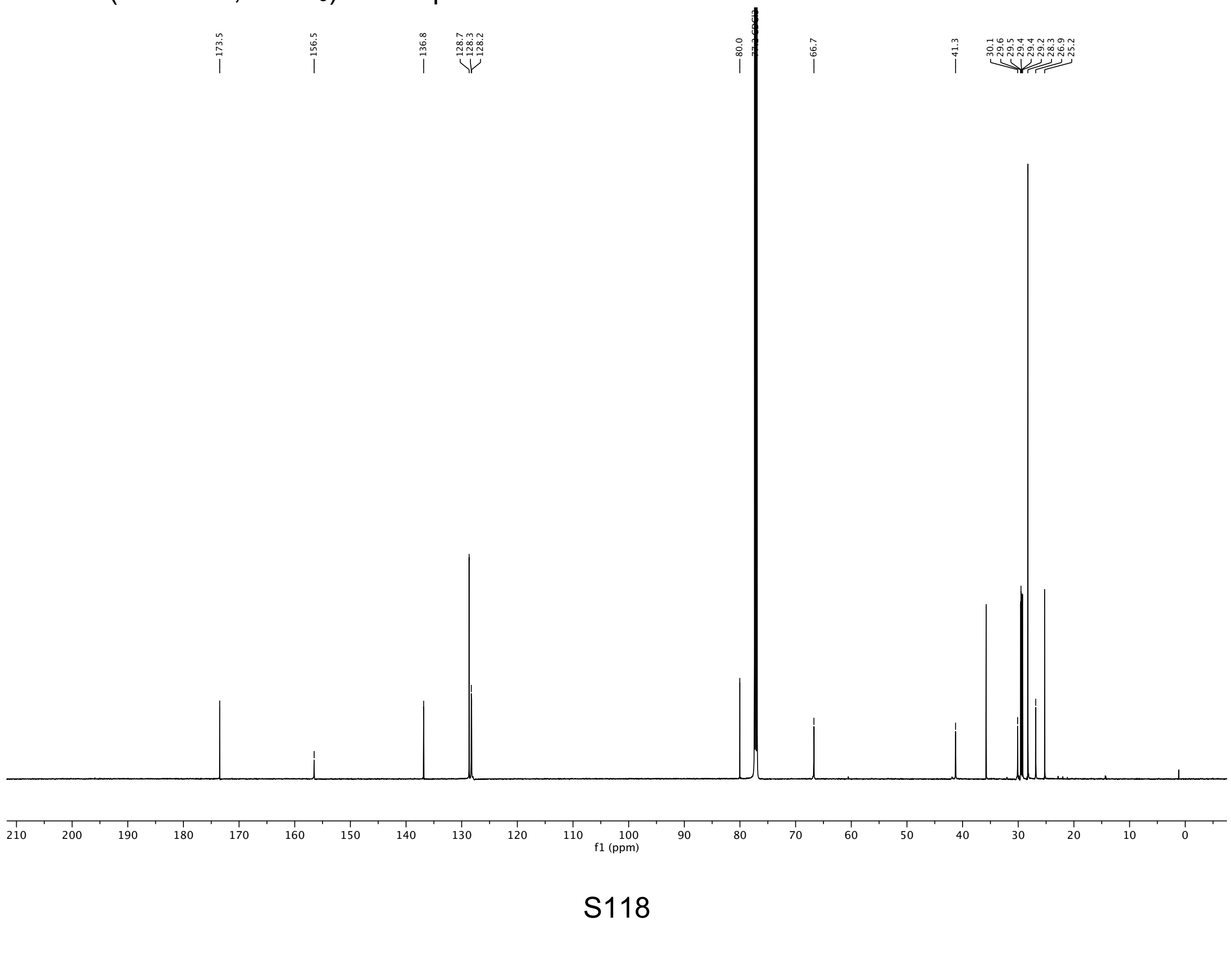

[1]H NMR (400 MHz, $CDCl_3$) of compound **S10**.

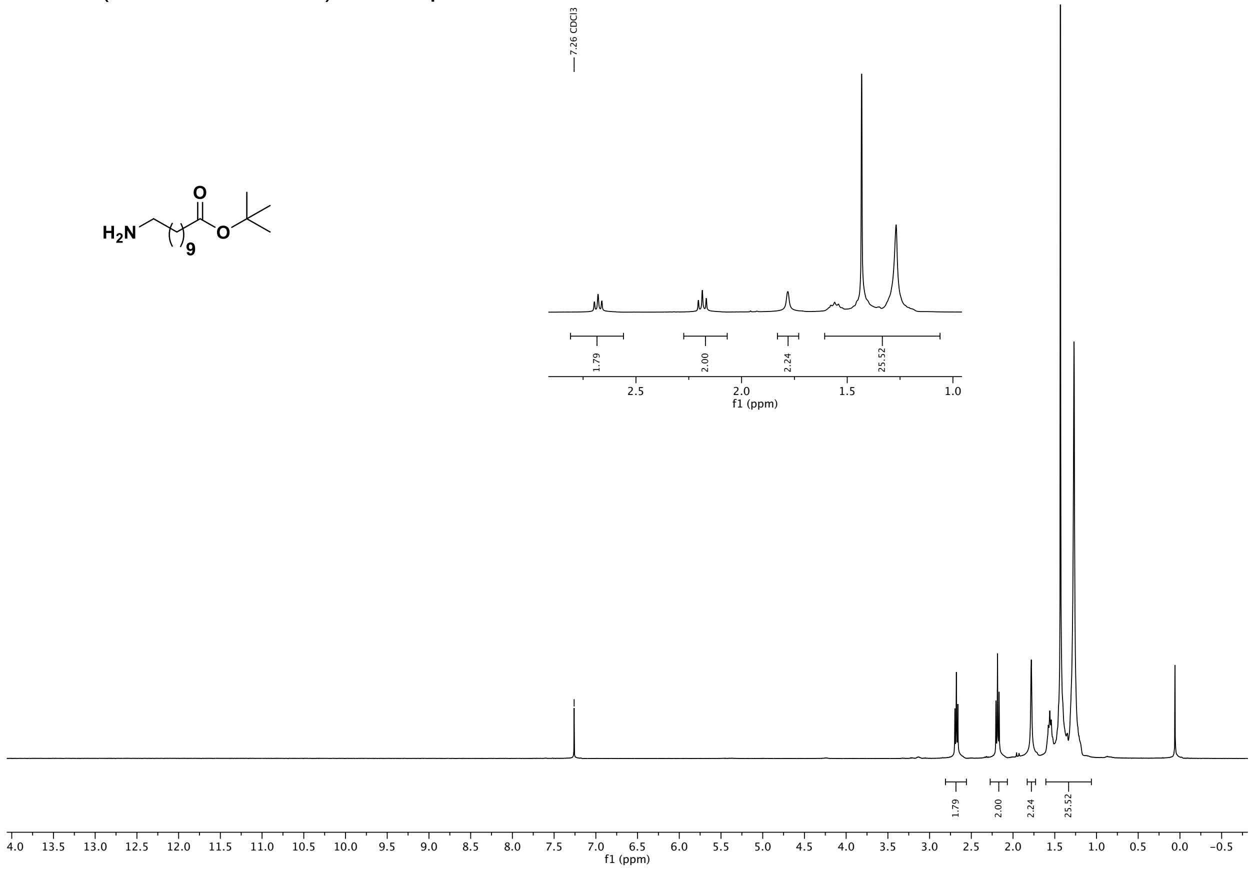

[13]C NMR (101 MHz, $CDCl_3$) of compound **S10**.

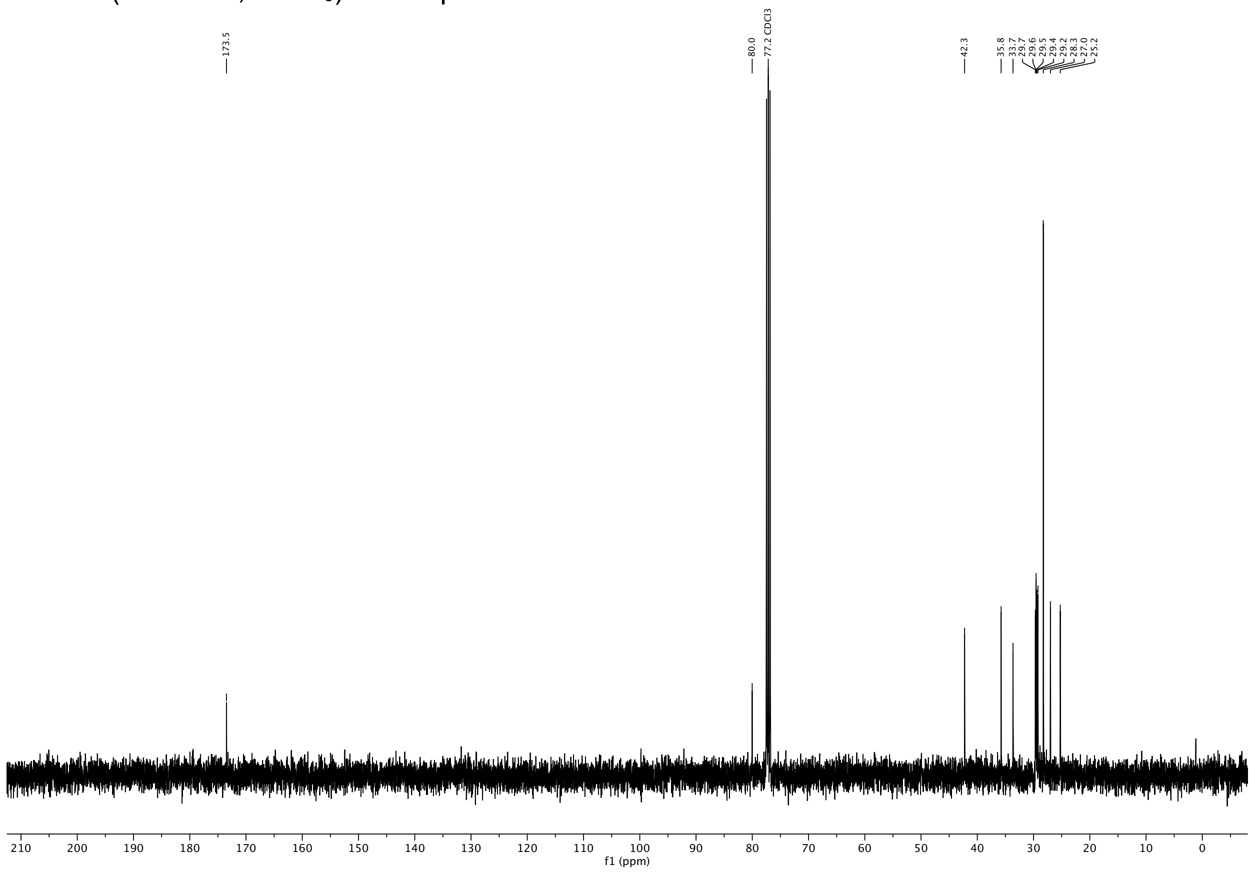

[1]H NMR (600 MHz, DMSO-$d_6$) of compound **S11**.

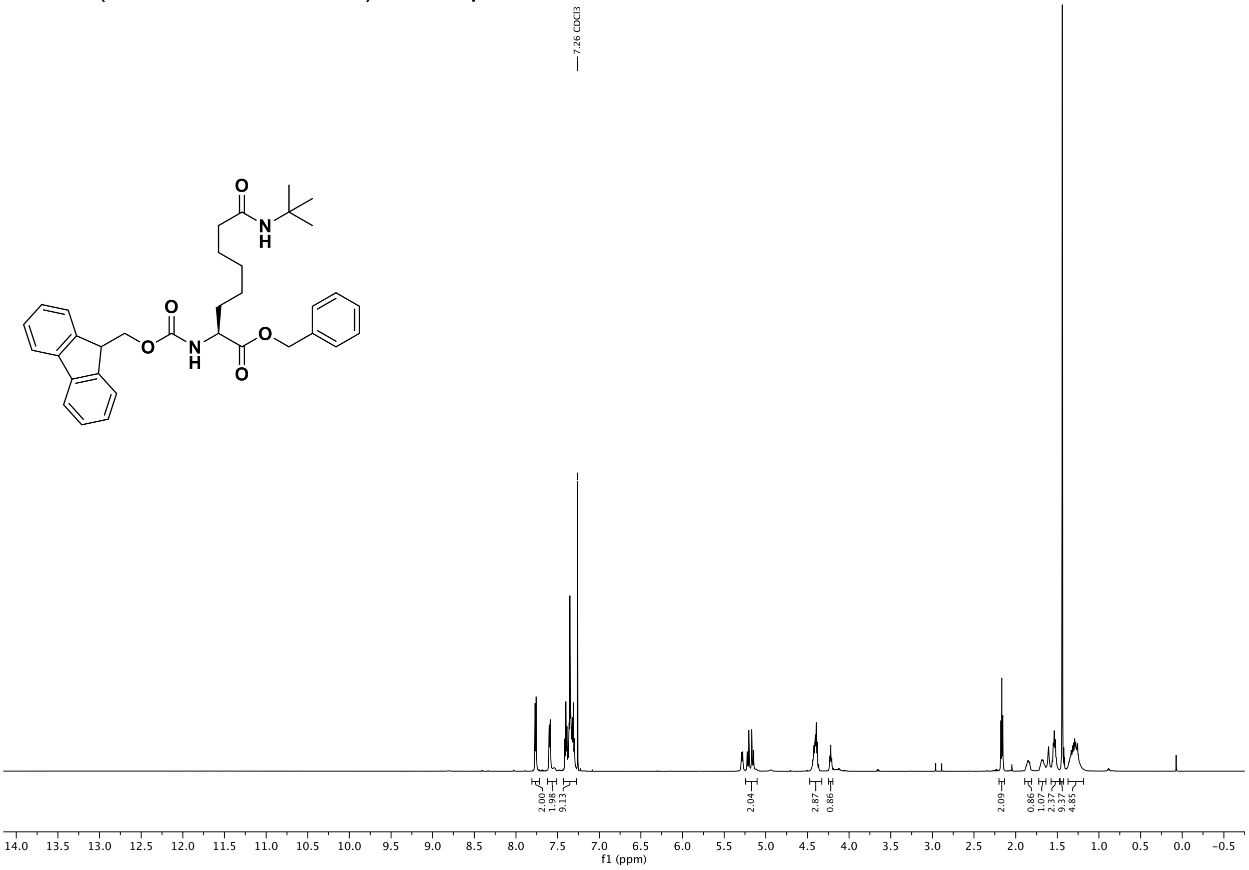

[13]C NMR (151 MHz, DMSO) of compound **S11**.

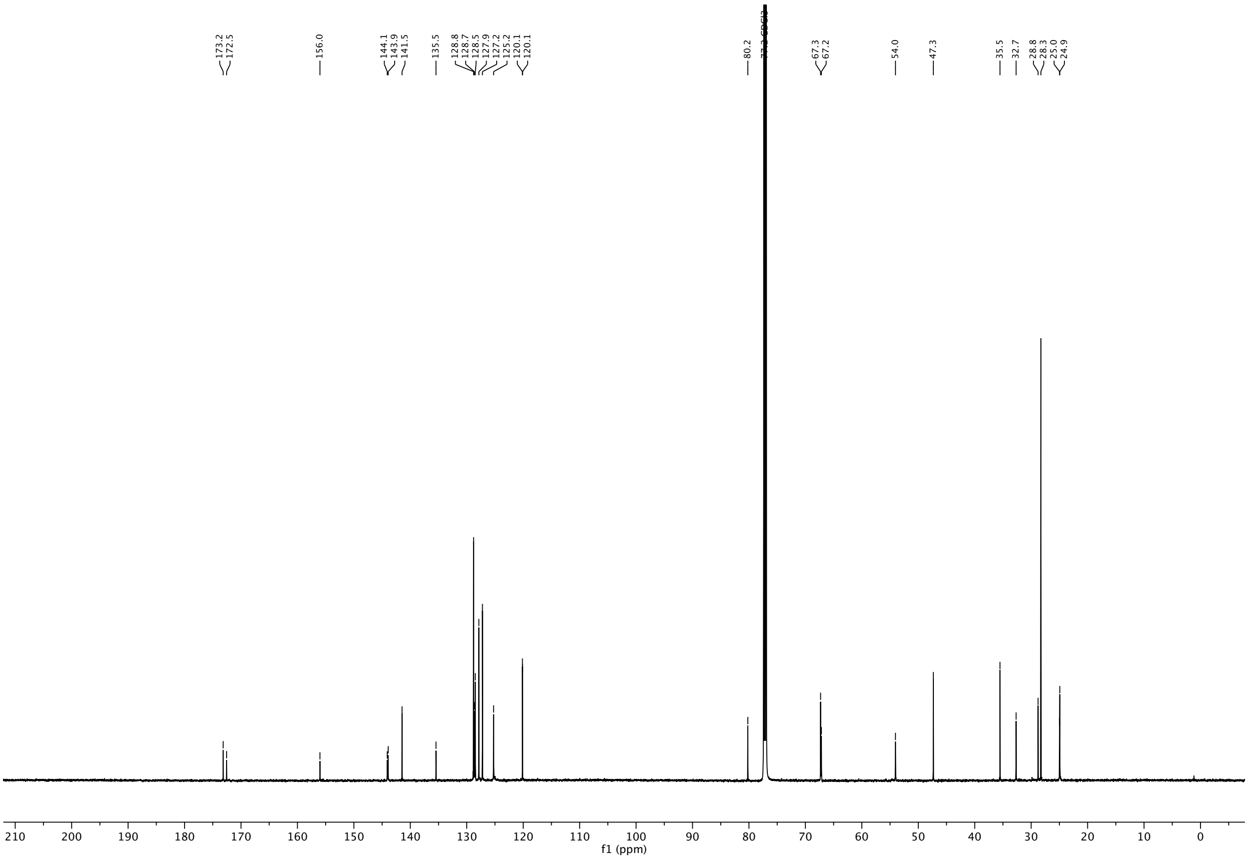

$^{1}$H NMR (600 MHz, DMSO-$d_6$) of compound **S12**.

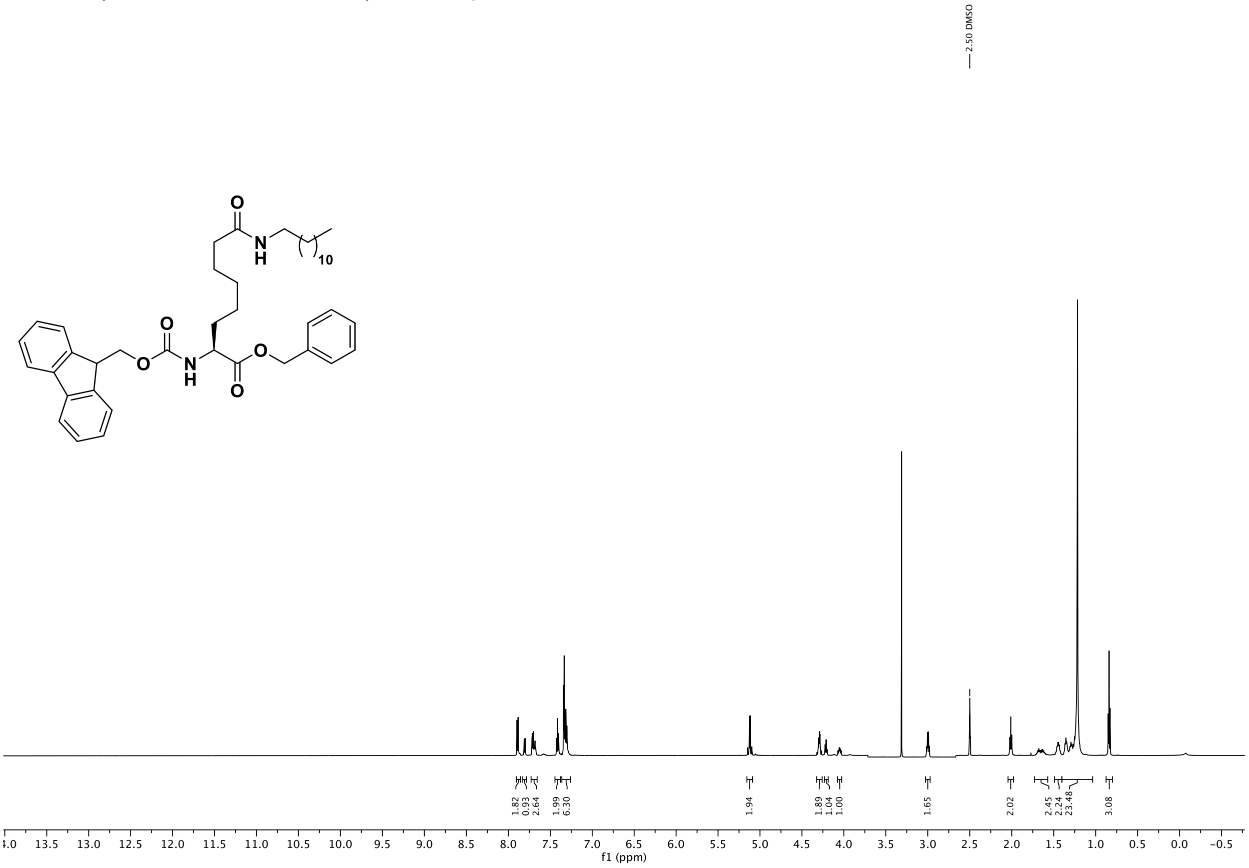

$^{13}$C NMR (151 MHz, DMSO) of compound **S12**.

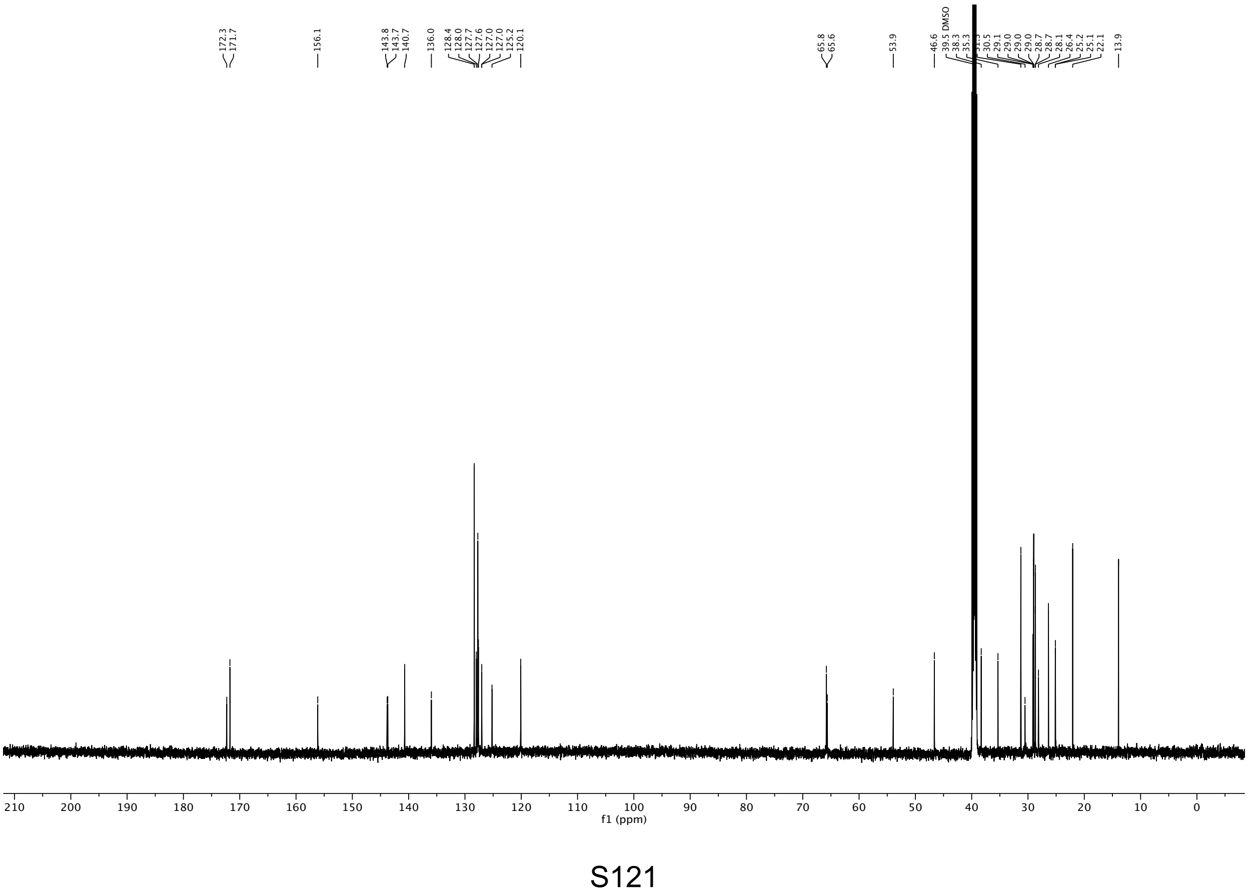

[1]H NMR (600 MHz, DMSO-$d_6$) of compound **S13**.

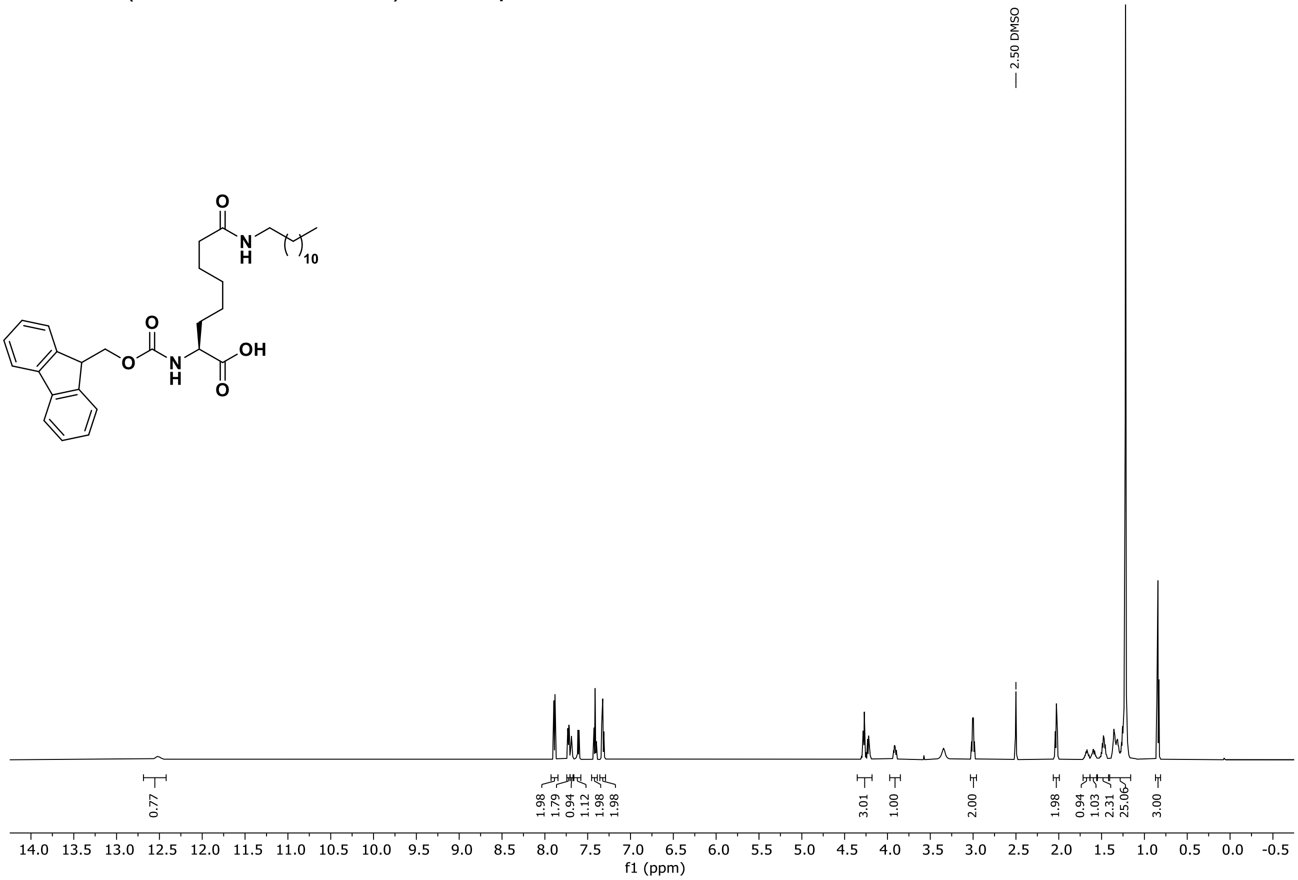

[13]C NMR (151 MHz, DMSO) of compound **S13**.

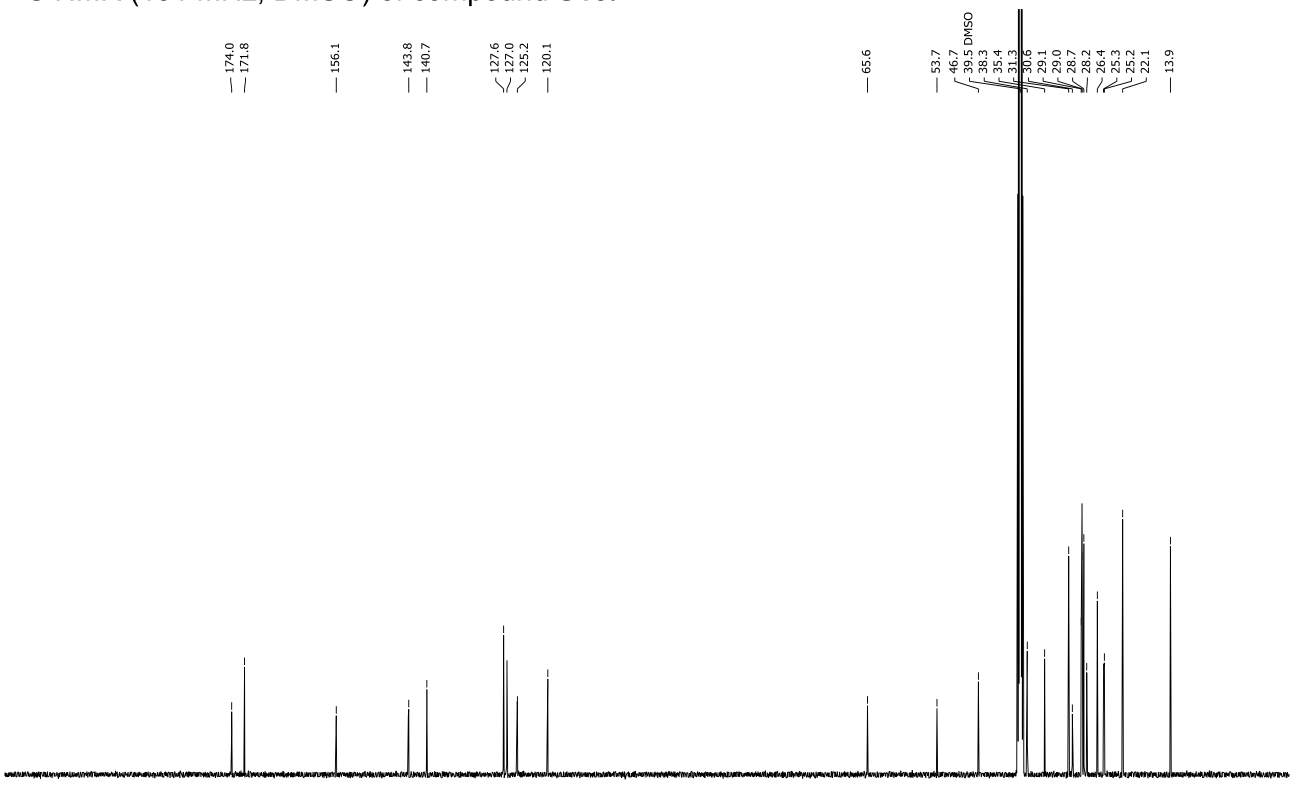

$^{1}H$ NMR (600 MHz, DMSO-$d_6$) of compound **S14**.

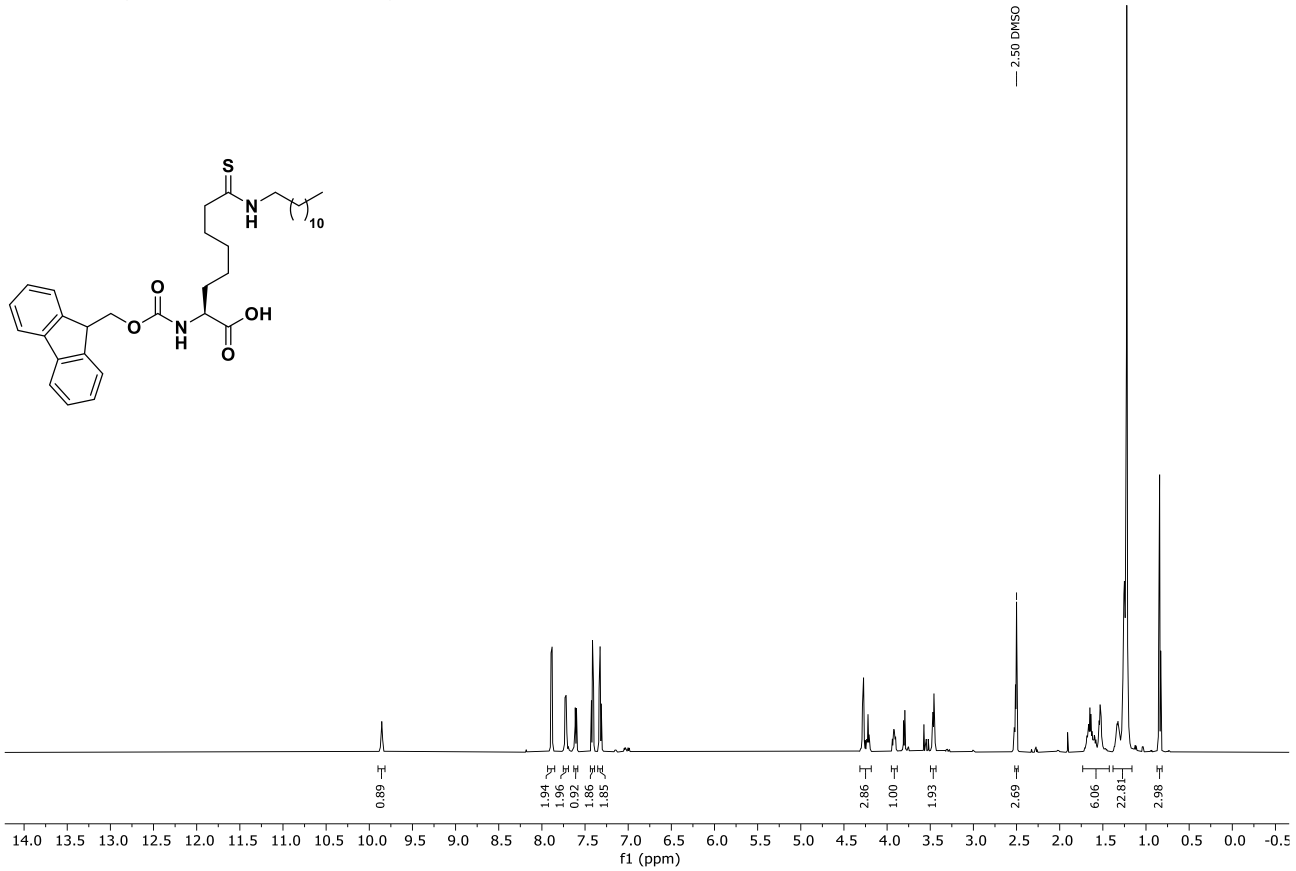

$^{13}C$ NMR (151 MHz, DMSO) of compound **S14**.

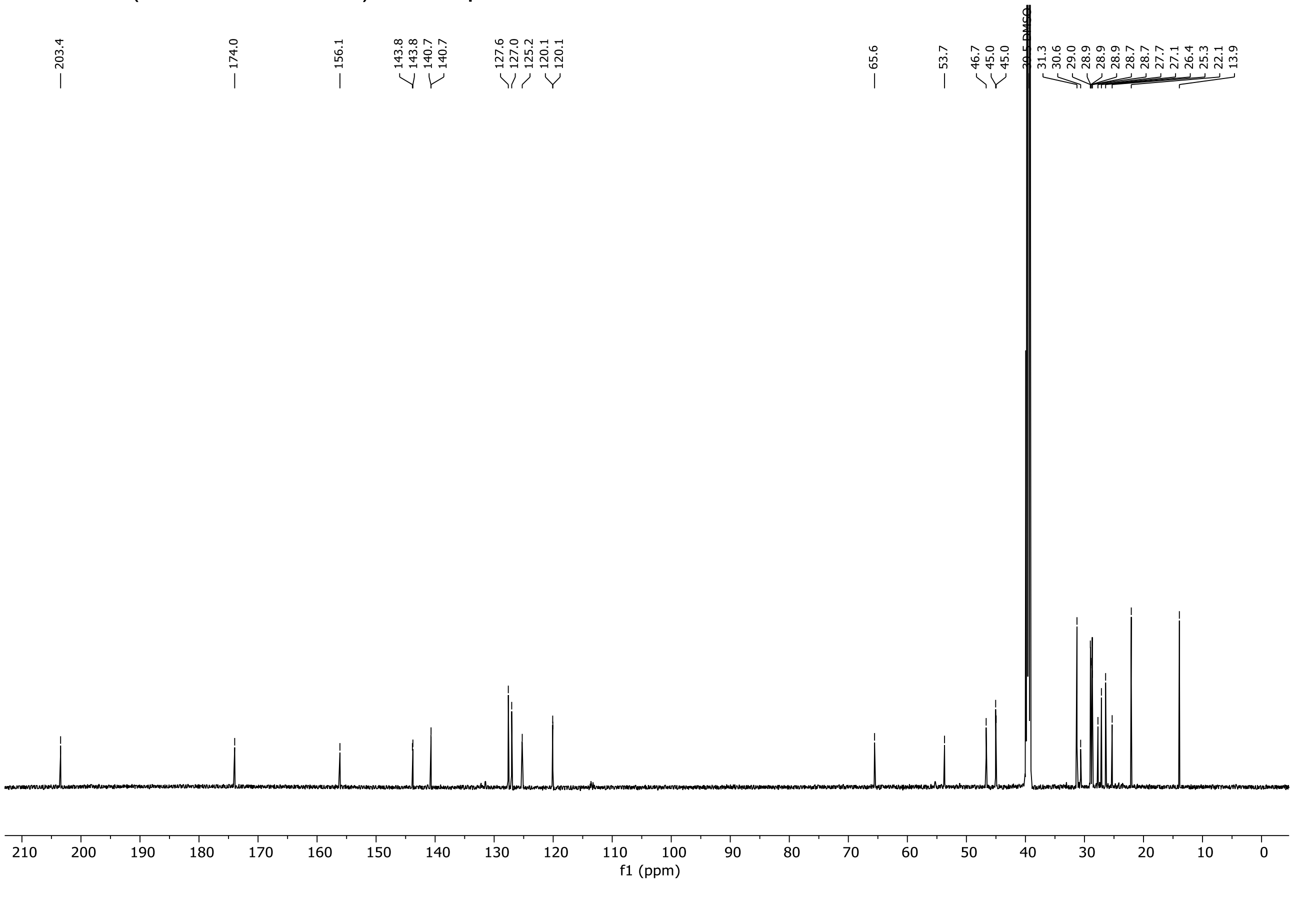
